# Leveraging Developmental Landscapes for Model Selection in Boolean Gene Regulatory Networks

**DOI:** 10.1101/2023.01.08.523151

**Authors:** Ajay Subbaroyan, Priyotosh Sil, Olivier C. Martin, Areejit Samal

**Affiliations:** The Institute of Mathematical Sciences (IMSc), Chennai 600113, India; Hormi, Bhabha National Institute (HBNI), Mumbai 400094, India; Université Paris-Saclay, CNRS, INRAE, Univ Evry, Institute of Plant Sciences Paris-Saclay (IPS2), 91405 Orsay, France; Université Paris-Cité, CNRS, INRAE, Institute of Plant Sciences Paris-Saclay (IPS2), 91405 Orsay, France

**Keywords:** Gene regulatory networks, Boolean modeling, Developmental landscape, Relative stability, Model selection

## Abstract

Boolean models are a well-established framework to model developmental gene regulatory networks (DGRN) for acquisition of cellular identity. During the reconstruction of Boolean DGRNs, even if the network *structure* is given, there is generally a very large number of combinations of Boolean functions (BFs) that will reproduce the different cell fates (biological attractors). Here we leverage the developmental landscape to enable model selection on such ensembles using the *relative stability* of the attractors. First we show that 5 previously proposed measures of relative stability are strongly correlated and we stress the usefulness of the one that captures best the cell state *transitions* via the mean first passage time (MFPT) as it also allows the construction of a cellular lineage tree. A property of great computational convenience is the relative insensitivity of the different measures to changes in noise intensities. That allows us to use stochastic approaches to estimate the MFPT and thus to scale up the computations to large networks. Given this methodology, we study the landscape of 3 Boolean models of *Arabidopsis thaliana* root development and find that the latest one (a 2020 model) does not respect the biologically expected hierarchy of cell states based on their relative stabilities. Therefore we developed an iterative greedy algorithm that searches for models which satisfy the expected hierarchy of cell states. By applying our algorithm to the 2020 model, we find many Boolean models that do satisfy the expected hierarchy. Our methodology thus provides new tools that can enable reconstruction of more realistic and accurate Boolean models of DGRNs.

## I. INTRODUCTION

The cells of a multicellular organism have the same genotype yet the associated tissues and organs manifest a plethora of different cell types. This phenotypic diversity has long been expected to arise from the multistability of gene expression patterns of the underlying dynamical system attached to the Gene Regulatory Network (GRN) [1, 2]. Boolean networks are a mathematical framework to model *developmental* GRNs (DGRNs) where gene expression levels are coarse-grained to take binary values [3–5], “0” corresponding to lack of expression (gene “off”) and “1” corresponding to presence of expression (gene “on”). Boolean networks often exhibit multistability, having more than one attractor state [5], and a number of studies have confirmed that Boolean models can have attractor states that correspond well to gene expression patterns specific to various cell types [6, 7]. In fact, since its inception, the Boolean approach has been quite successful in modeling a wide range of developmental processes. Some of these include: blood cell development [8], myeloid progenitor differentiation [9], pancreas cell differentiation [10, 11], *Arabidopsis thaliana* root stem cell niche [12, 13] and flower development [14]. Reconstructions of such Boolean models of GRNs require the specification of three components: network structure (which are the regulators of each target gene), logic rules (how do regulators of a gene affect its expression level) and an update scheme (e.g., update all genes synchronously or not) [15, 16]. Current efforts to reconstruct Boolean DGRNs are generally underdetermined, *i.e*., there exists many combinations of regulatory logic rules that can recover the desired gene expression patterns [17], even for a given network structure [10]. Without additional information, modelers typically have no other choice than to fix somewhat arbitrarily certain logic rules, a process that introduces hidden biases and preferences that are never made explicit. This contribution, in the spirit of earlier ones by some of us [10, 17], aims to address this unmet need of providing a systematic framework for *model selection* of dynamical GRNs from an ensemble of models that are equally plausible at the level of their logic rules. To do so, we work with the *hierarchy* of cell types emerging from the *relative stability* associated with the developmental landscape and demonstrate how that information can be used to select between otherwise equivalent models.

Many decades ago, Waddington introduced the so-called “developmental landscape” as a convenient metaphor of processes driving cell fate determination [1]. In his representation, cells are analogous to balls rolling downhill, committing to cell fates based on trajectories that encounter branch points on the landscape. Though Waddington’s landscape was originally meant to illustrate the differentiation paths for going from embryonic to terminally differentiated cells [1], it is now used to represent arbitrary differentiation cascades leading to various differentiated cell types. Note that if errors or uphill motion can happen - even if rarely -, this picture allows non standard transitions wherein cells will switch to another type, via de-differentiation (a mature cell type transmutes to a progenitor in the same lineage tree), trans-differentiation (a mature cell in one lineage transmutes to a mature cell type of another lineage) and reprogramming (a differentiated cell reverts to an induced pluripotent stem cell) [18–20]. Cell state transitions can be driven by many processes and of course are almost always strongly regulated to occur in a particular direction, but at the branching points one can expect noise [21, 22] to play a significant role. Such noise can come from the inherent stochasticity associated with biochemical processes controlling gene expression within cells or from noise coming from outside of the cells [22–24]. It is appropriate to keep in mind that the landscape metaphor is a bit misleading because in the biological context there is no such thing as height or potential energy; only very restricted dynamical systems allow for a potential energy function. Notwithstanding those limitations, a good feature of the landscape picture is that it enforces a strong (downhill) orientation preference for transitions. In other words, downhill changes are much more probable than the corresponding uphill ones, which amounts to saying there is not that much noise in these systems. This forms the conceptual basis of the *relative stability* of cell types, or the ‘relative ease’ of transitioning from one cell type to another [10, 25, 26].

Though the genesis of using developmental landscape for Boolean model selection was proposed earlier by Zhou et *al*. [10], the idea has been explored only at a small scale. In what follows we concretize and extend the ideas in [10, 26] into a systematic framework that leverages the developmental landscape to perform model selection for larger networks. First, we revisit the 5 measures of relative stability considered by Joo et *al*. [26], based on size of basin of attraction (BOA), steady state probability (SSP), mean first passage time (MFPT), basin transition rate (BTR) and stability index (SIND). We quantify the concordance between these measures of relative stability in a structured manner using different ensembles derived from a Root Stem Cell Niche (RSCN) network [12] and a Pancreas differentiation network [10]. We find that all 5 measures are strongly concordant and pick for subsequent analysis the MFPT as it emulates cell state transitions more closely than the other measures. Using the MFPT, we show how to construct an associated cellular lineage tree. To illustrate its usefulness, we determine the frequency of occurrence of different lineage trees in the above mentioned ensembles. In addition, because the matrix formalism to calculate MFPT as proposed in [10] cannot be scaled up computationally, we take a stochastic approach to compute the MFPT. With this method, we compute the relative orderings and cellular lineage trees for the successive root development models of Alvarez-Buylla’s group [13, 27, 28] that have increasing complexity. We find that the latest model proposed by that group does not satisfy the expected hierarchy, which indicates that relative stability between cell types was not one of their (conscious or not) criteria for selecting the logic rules. Lastly, we propose an iterative greedy algorithm that leverages the expected developmental landscape (or hierarchies) to perform model selection from an ensemble of models that recover the biologically desired gene expression patterns. Thanks to these conceptual and computational developments, we provide a systematic framework to perform model selection within a biologically plausible ensemble of Boolean models using the associated developmental landscapes.

## II. MATHEMATICAL FRAMEWORK

### A. Boolean network models of gene regulatory networks

A Boolean model of a gene regulatory network (GRN) consists of nodes and directed edges where nodes correspond to genes and directed edges correspond to interactions between them [3–5]. Boolean networks assume only two expression states for genes, ‘on’ or ‘off’, akin to switches. The state of a node can therefore be represented by a Boolean variable that can take values 1 (‘on’) or 0 (‘off’). For a Boolean network with *N* nodes, we denote by *x_i_*(*t*) the state of node *i* at time *t*, where *i* ∈ {1,*N*} and *x_i_* ∈ {0,1}. We use a vector **X**(*t*) to denote the state of all variables *x_i_*(*t*) of the network. **X**(*t*) is the the gene expression pattern of the network at time t. As each node can assume 2 states, there are 2^*N*^ possible gene expression patterns, defining the state space of the Boolean network. Boolean functions (BFs) (or *logical update rules*) along with an update scheme (*synchronous* [5] or *asynchronous* [29]) determine the temporal dynamics of the network, taking it from time *t* to time *t* + 1. In this work, each node is assigned one BF and the network is updated synchronously. A BF governs the dynamics at the level of a node by performing logical operations on the inputs to that node and returning an output. More succinctly, 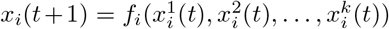, where *f_i_* is the BF that acts on the *k* inputs to node *i*, 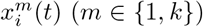, to return an output *x_i_*(*t* + 1). If all BFs **F** = (*f*_1_, *f*_2_,…, *f_N_*) act on all nodes simultaneously, the update is said to be synchronous and is expressed by the equation:

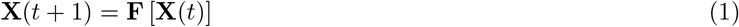

Starting from an initial state **X**(0), a state space trajectory can be traced out by the recursive application of Eq. (1). The collection of all such trajectories constitutes the *state transition graph* of the Boolean network. Under synchronous dynamics, a trajectory can meet with 2 possible fates. One, it reaches a state which on further update remains unaltered, in which case the trajectory is said to have converged to a *fixed point attractor* or simply, a *fixed point*. Two, it keeps cycling through a set of states, in which case the trajectory is said to have converged to a *cyclic attractor* or simply, a *cycle*. All the states which converge to an attractor (including the attractor itself), constitute its basin of attraction. We remark that the dynamics presented above is purely deterministic and generates a state transition graph with multiple disconnected components, each one corresponding to one attractor. In DGRNs which are the focus of this contribution, fixed points are of biological significance as they represent steady state expression patterns that specify cell types.

### B. Relative stability of fixed points

In developmental dynamics, the propensity of a less differentiated cell type to become a more differentiated one is higher than the converse. This inherent asymmetry in cell state transitions forms the conceptual basis of *relative stability* [10, 25]. The schematic illustration in Fig. 1(a) of a developmental landscape, inspired by Waddington, captures this asymmetry. In this particular landscape, the ‘Green’ cell type can take 2 possible routes downhill shown by the solid arrows, either to become a ‘Red’ cell type or a ‘Blue’ cell type. The reverse transitions are also possible as illustrated in Fig. 1(a) via the dashed arrows originating from the Blue or Red cell type and directed uphill to terminate at the ‘Green’ cell type. Of course, just as in the actual biological systems, such uphill trajectories, though possible, are less likely to occur spontaneously than the corresponding downhill trajectories. The relative stability between a pair of cell types *u* and *v* is determined by the ‘relative ease’ of transitioning from one type to the other [10, 25]. In other words, if it is relatively easier to transition from v to u than the other way around, then u is relatively more stable than v. Such cell state transitions cannot be modeled using the deterministic Boolean framework since attractors form disconnected components there (see section II A).

**FIG. 1.**
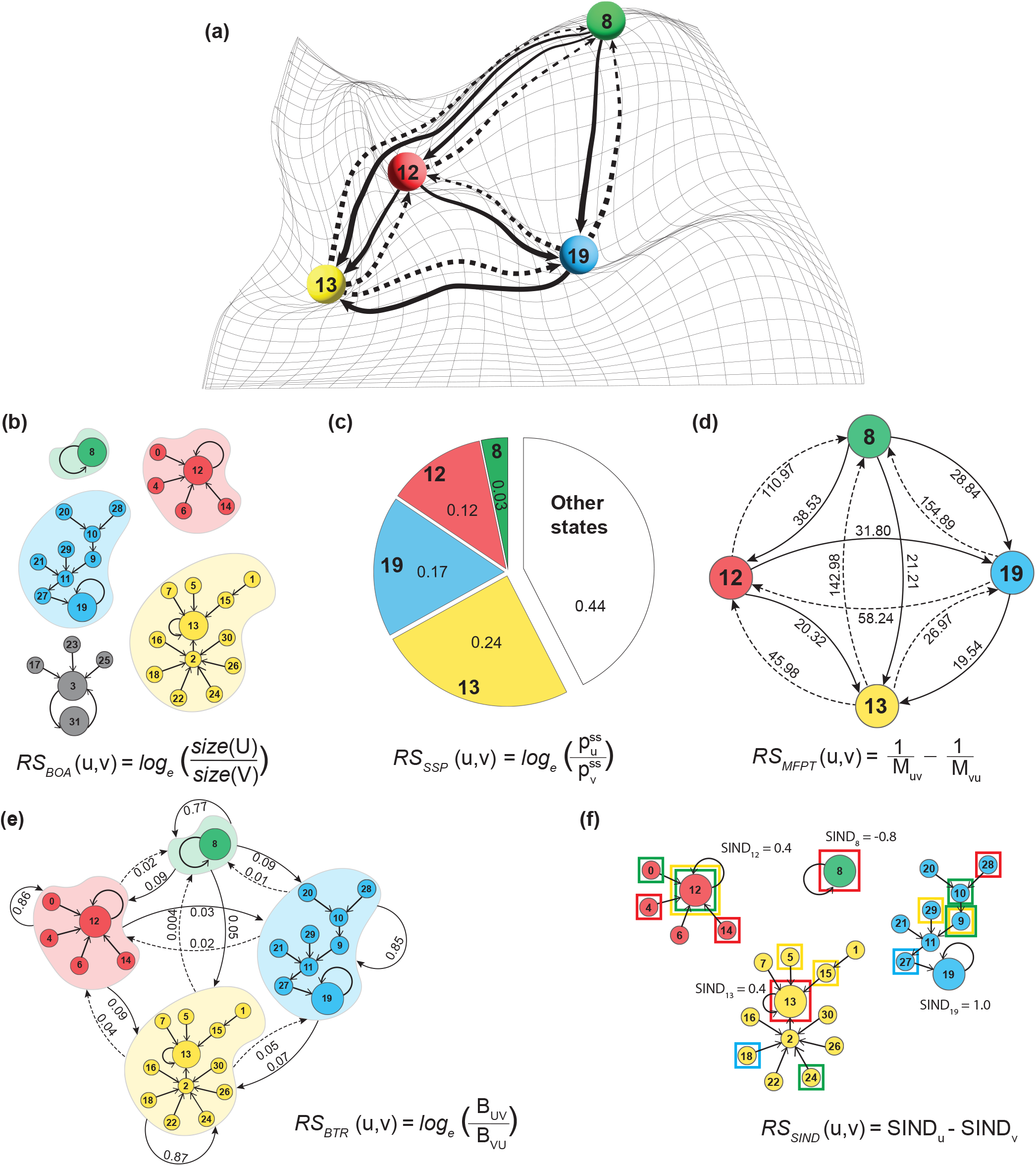
Developmental landscape inspired by Waddington and relative stability measures. The epigenetic landscape and its measures of relative stability are depicted here for the toy model displayed in SI Fig. S1. **(a)** The colored balls and their numeric labels represent different cell types and their corresponding fixed point states in the toy model. The balls are trapped in local minima but at different “altitudes”. Figuratively, the difference in altitude between two cell types indicates the *relative stability* between them. **(b) Basin of attraction (BOA).** The disconnected components represent the basins of attraction for the different attractors of the toy model. The colors of these basins are those of the cell states shown on the landscape. **(c) Steady state probability (SSP).** The pie chart gives the steady state probabilities of each fixed point (shown in color) and of all the other states (non fixed points, shown in white). **(d) Mean First Passage Time (MFPT).** The complete digraph provides the MFPT between all pairs of fixed points along both directions. Solid and dashed lines indicate smaller and larger MFPTs respectively. **(e) Basin Transition Rate (BTR).** A complete digraph where the nodes are the basins of attraction of fixed points and the edges are the BTR from one basin to the other. Solid and dashed lines indicate larger and smaller BTRs respectively. **(f) Stability Index (SIND).** A colored square indicates the 1-Hamming neighbors of the fixed point associated with that color. This information is used to compute the SIND (see section II B).

Zhou *et al*. [10] circumvented this problem by adding stochasticity to the dynamics. As a result, the state transition graph became an ergodic Markov chain [30]. Zhou *et al*. applied methods developed for ergodic Markov chains to Boolean modeling and used them to define the relative stability between a pair of cell types. Their formalism is briefly described below.

Let the *deterministic* dynamics be defined via a matrix **T** whose elements *T_ij_* = 1 if the update of ***X**_j_* via Boolean functions **F** gives **X**_*i*_. That is,

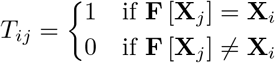

where *i, j* ∈ {0, 2^*N*^ – 1}. Stochastic dynamics is introduced via a noise parameter *η* that flips the state of each gene independently with a probability *η*. Let **P** be a matrix whose entries *P_ij_* are the probability that *η* alone drives the transition from **X**_*j*_ to **X**_*i*_. The number of flips necessary to drive a transition from state **X**_*j*_ to **X**_*i*_ in a single time step is the Hamming distance d(*i,j*) between **X**_*i*_ and **X**_*j*_. The entries *P_ij_* are thus defined as a function of *η* and d(*i, j*):

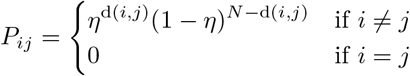

The deterministic **T** and the noise-dependent (*perturbation*) matrix **P** are then combined to give a state transition matrix **T**\* as follows:

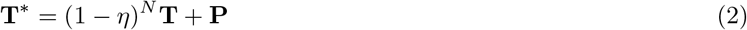

Note that 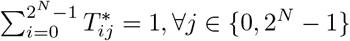 since:

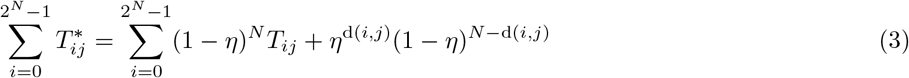

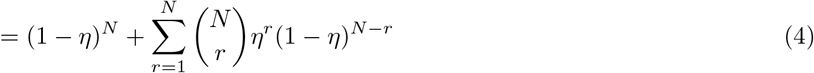

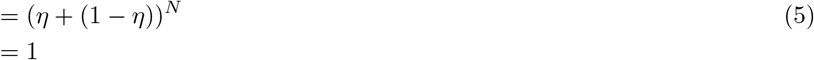

*T\** is thus a so called *stochastic matrix* and from it one can define different measures of relative stability [10, 26]. In the ensuing text, *u* and *v* symbolize biological fixed points, and *U* and *V* their basins of attraction, respectively.

#### Basin of Attraction (BOA)

A fixed point attractor with a larger basin size is expected to be more robust to perturbations and will exhibit greater stability compared to a biological fixed point with a smaller basin size. Fig. 1(b) illustrates the basins of attraction of the toy model detailed in SI Fig. S1. We define the relative stability between u and v using the sizes of basins of attraction *U* and *V* as follows:

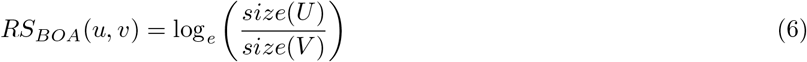

#### Steady State Probability (SSP)

Under the ergodic stochastic dynamics given by **T**\*, there is a unique steady state, equal to the vector of probabilities of finding the network in any particular state at long times. For any specified initial condition, let **p**(*t*) be the vector whose entry *p_j_*(*t*) is the probability of being in state **X**_*j*_ at time *t*. Note 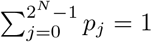. Then the probability vector at the next time step is given by:

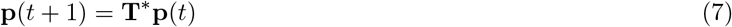

If **p**(*t* +1) = **p**(*t*), then **p**(*t*) is necessarily the (unique) steady state probability distribution **p**^*ss*^ of the network states. Fig. 1(c) illustrates the steady state probability distribution of the toy model shown in SI Fig. S1. From that we define the relative stability between *u* and *v* using **p**^*ss*^ as follows:

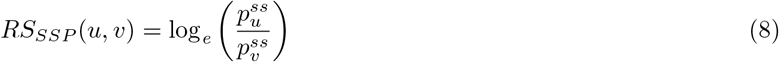

where 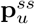 is the steady state probability of the fixed point *u* and similarly for *v*.

#### Mean First Passage Time (MFPT)

The number of steps along a state space trajectory starting at **X**_*j*_ and terminating at the first occurrence of **X**_*i*_ in a stochastic process is called the first passage time from **X**_*j*_ to **X**_*i*_. It’s mean is then the mean first passage time from **X**_*j*_ to **X**_*i*_ and is denoted by *M_ij_*. The MFPT of an ergodic Markov chain can be calculated analytically using the fundamental matrix **Z** [31] as follows:

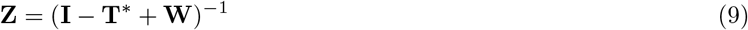

where **I** is the identity matrix and **W** is a matrix whose columns are the vector **p**^*ss*^. *M_ij_* is given by:

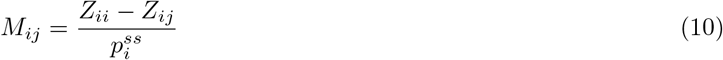

Fig. 1(d) illustrates the MFPT values from one cell type to another for the toy model shown in SI Fig. S1. The relative stability between the fixed points u and v defined using *M_uv_* [10] is given by :

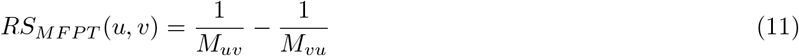

The basin transition rate is the probability to transition from any state in the basin of attraction of one fixed point *v* to the basin of attraction of another fixed point *u* when applying the stochastic dynamics for one time step. Mathematically, it is defined [26] via the formula:

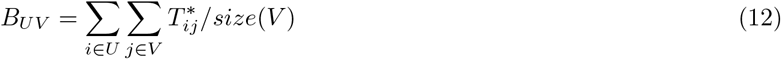

where *i* and *j* denote the states in the basins *U* and *V* respectively of fixed points u and v. Fig. 1(e) illustrates the BTR between all pairs of basins of attraction for the toy model shown in SI Fig. S1. We define the relative stability between *u* and *v* using *B_UV_* as follows:

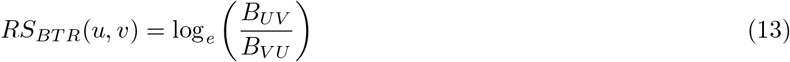

#### Stability Index (SIND)

The stability index *SIND_u_* of a fixed point *u* is defined as follows [26]:

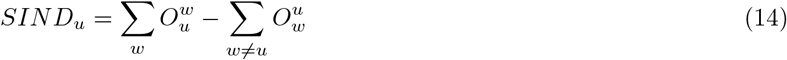

where the first sum is over all fixed points and the second over all fixed points other than *u* itself. In this formula, the quantity 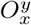 is a ratio, whose numerator is the number of 1-Hamming neighbors of fixed point *y* belonging to the basin of fixed point *x*, and whose denominator is the total number of 1-Hamming neighbors of fixed point *y*. Fig. 1(f) illustrates the SIND of the fixed points for the toy model shown in SI Fig. S1. We define the relative stability between *u* and *v* using *SIND_u_* as follows:

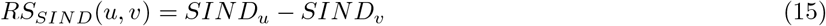

Note that if *RS_X_*(*u,v*) > 0, where *X* ∈ (*BOA, SSP, MFPT, BTR, SIND*), then the cell type corresponding to biological fixed point *u* is more stable than the cell type corresponding to biological fixed point *v*.

### C. Minimum Spanning Arborescence

A spanning tree of a connected undirected graph is a subgraph that is a tree and contains all the vertices of the graph. A minimum spanning tree of an undirected graph with weighted edges is a spanning tree that has the minimum sum over it’s edge weights. An *arborescence* is a rooted, directed tree in which all edges are oriented away from the root. A minimum spanning arborescence is the directed analog of the minimum spanning tree constructed from a directed graph with weighted edges. The total number of spanning arborescences for a complete digraph with *n* vertices (having distinctly labelled nodes) is *n*^*n*−1^. This can be reasoned as follows. From Cayley’s theorem [32], the number of distinct (undirected) trees of *n* labeled vertices is *n*^*n*−2^. To get an arborescence from an undirected tree, one simply has to specify the root, which gives a directed tree with edges that point away from the root. Since there are *n* ways to choose the root, there are *n* × *n*^*n*−2^ = *n*^*n*−1^ arborescences for *n* nodes.

## III. METHODS

### A. Deriving biologically plausible ensembles for model selection

Here we describe a model selection framework where, by successively imposing different biologically motivated constraints on a Boolean model with a fixed network structure, it is possible to converge to a smaller subset of models that are biologically relevant [10]. Let *k_i_* be the number of inputs to a gene *i* ∈ {1, *N*} in the Boolean network. Keeping the network structure fixed (i.e., list of input genes for each output gene), without imposing any constraints on the logical update rules or truth tables, there are 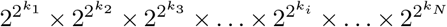 possible combinations of BFs (or Boolean models), which is often times astronomical. The first constraint is to restrict the truth tables to respect the fixed point condition for each of the desired biological fixed points. Every fixed point will constrain the output of 1 row in all of the truth tables, so *N* fixed points will constrain at most *N* rows of every truth table. Note that multiple fixed points may lead to redundant constraints. This constraint guarantees the recovery of all the biological fixed points but nevertheless does not exclude the presence of other (possibly, irrelevant) attractors. When feasible, one could also demand that there be no irrelevant attractors (i.e., attractors that do not correspond to biological fixed points). The Boolean GRNs then recover *only* the desired biological attractors. Secondly, we impose that the BF at each node conforms to the activatory or inhibitory signs of it’s regulators. In other words, we ensure that the BFs are *sign conforming* with respect to the network structure. Third, the choice of BFs is restricted to nested canalyzing functions (NCFs) (or some other choice such as effective function (EFs), or unate functions (UFs), or effective and unate functions (EUFs) (see SI text, section 1). As a motivation for this constraint, it has recently been shown that NCFs possess the minimum average sensitivity [33] for BFs and confer critical dynamics to the model, a hallmark of the dynamics of GRNs [5, 30, 33, 34]. Finally, the relative stability constraints on the biologically fixed points can be used to select for models that conform to the expected developmental landscape.

We quantitatively illustrate the above methodology using a Boolean model of *Arabidopsis thaliana* root stem cell niche (RSCN) [12] whose network structure is provided in SI Fig. S2(a). This RSCN model (*model A* in [12]) has 9 nodes, 19 edges and 4 fixed points that correspond to the cell types: Quiescent center (QC), Vascular initials (VI), Cortex-Endodermis initials (CEI) and Columella epidermis initials (CEpI) (SI Fig. S2(a)). In what follows, the order of nodes of this RSCN network is: PLT, AUXIN, ARF, AUXIAA, SHR, SCR, JKD, MGP, WOX5. The total number of models possible for this network without any constraints on the truth tables is 4 × 4 × 4 × 4 × ×4 × 65536 × 16 × 256 × 4294967296 ≈ 1.18 × 10^21^. By imposing the fixed point constraints on the truth tables we get 2×2×2×2×1×4096×2×16×268435456 = 562949953421312 ≈ 5.63×10^14^. Next, on imposing the sign conforming constraint, we get 2 × 2 × 2 × 2 × 1 × 70 × 2 × 2 × 848 = 3799040 ≈ 3.8 × 10^6^. Further imposing the NCF constraint, we get 1 × 1 × 1 × 1 × 1 × 17 × 1 × 1 × 75 = 1275 possible DGRNs. Demanding that only the desired fixed points be recovered, that number is further reduced to 170. Finally, imposing relative stability constraints from the expected developmental landscape, namely, that QC (Quiescent center) be the least stable of all the fixed points [35], we are left with 80 models. Note that these constraints are imposed successively, one after the other in the order presented above. Thus from approximately 10^21^ models, the space of viable models can be shrunk to just 80 models.

Since the last 2 constraints are imposed at the level of the model (not truth table), it may be computationally cumbersome to apply them to networks where the number of models are typically large even after imposing constraints on the truth tables. This necessitates the development of stochastic methods to enable model selection on larger ensembles of biologically plausible models.

### B. Two biological models and their ensembles of GRNs

Statistical analyses presented in this work are performed on ensembles of GRNs derived from the network structure of two biological models. The first is a root stem cell niche (RSCN) model of *Arabidopsis thaliana* proposed by Azpeitia *et al*. [12]. The other is a pancreatic cell differentiation model proposed by Zhou *et al*. [10]. The RSCN model is described in the previous section (see section III A), and has 4 attractors as shown in SI Fig. S2(a). The pancreatic cell differentiation model has 5 genes, 13 edges and 3 fixed points that correspond to the cell types: Exocrine, α/PP progenitor and *β/δ* progenitor (SI Fig. S2(b)).

Our nomenclature for these different ensembles is then as follows (two per model system):

- *Root_sc–NCF_*: GRN models that recover *at least* the biological attractors using sc-NCFs on the RSCN network.
- 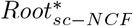: GRN models that recover *only* the biological attractors using sc-NCFs on the RSCN network.
- *Panc_sc–NCF_*: GRN models that recover *at least* the biological attractors using sc-NCFs on the pancreas cell differentiation network.
- 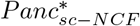: GRN models that recover *only* the biological attractors using sc-NCFs on the pancreas cell differentiation network.

The *Root_sc–NCF_* and 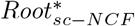 ensembles have 1275 and 170 Boolean models respectively. We remark here that the BF at the AUX node alone was fixed to the choice made in the original model. The *Panc_sc–NCF_* and 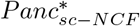 ensembles have 3600 and 109 Boolean models respectively. We also constructed similar ensembles by imposing that the BFs at each node be sc-EUFs rather than sc-NCFs. This led to 4 other ensembles: *Root_sc–EUF_*, 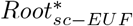, *Panc_sc–EUF_* and 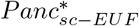. The *Root_sc–EUF_* and 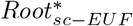 ensembles have 36600 and 1400 Boolean models respectively. The *Panc_sc–EUF_* and 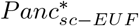 ensembles have 7056 and 159 Boolean models respectively.

### C. Relative stability and ordering of fixed points

As mentioned in section III A, the relative stability can be used as a constraint to select for models that conform to the developmental landscape. Relative stability is defined for pairs of fixed points. So for *n* fixed points, there exist n(n – 1)/2 possible comparisons. The associated inequalities provide a partial view of the landscape. Note that experimental data may not give us all the n(n – 1)/2 inequalities. In case all such inequalities are available, it is desirable to combine them to obtain the complete picture of the landscape. Consider for instance the *RS_MFPT_* for the toy model shown in Fig. 1(d). There are 4 fixed points, *R*, *B*, *G* and *Y*, leading to the following 6 pairwise relations: *G < R, G < B, G <Y, R < B, R < Y, B <Y*. All these inequalities can be combined into the linear hierarchy, *G < R < B < Y*, corresponding to a *total* order. But finding such a total order may not always be possible, there may be *inconsistencies* amongst the inequalities. For instance, if we had instead the inequalities *G < R, G < B, G<Y, B < R, R <Y* and *Y < B*, clearly the last three make it impossible to find a total order. Such a situation leads us to go beyond linear hierarchies by using tree-based (partial) hierarchies as we now discuss.

### D. Constructing cellular lineage trees using MFPT

Developmental trajectories are expected to follow paths of least resistance on the epigenetic landscape, taking one from undifferentiated to differentiated cells. Because of the possibility of branching during these differentiation processes, we can think of cell lineages as being associated with oriented trees. In terms of MFPT, we expect that a transition from an undifferentiated state to a more differentiated state should be more probable and take less time than a transition in the opposite direction. This then motivates the construction of cellular lineage trees in any given model by using the MFPTs between the model’s different fixed points. Let **M** be a matrix whose entries *M_ij_* are the MFPTs from attractor *j* to attractor *i*. Note that **M** is an asymmetric matrix and all its principal diagonal elements are 0 (i.e *M_ii_* = 0). **M** can thus be represented as a complete weighted directed graph G(**M**) whose nodes are biological attractors and edges carry the weights *M_ij_* (for edges from *j* to *i*). The expected cell lineages should then follow from a directed rooted tree that minimizes the sum of the MFPTs over it’s edges. Such a tree is precisely the minimum spanning arborescence (MSA) of G(**M**) (see section II B). To construct a MSA from G(**M**), we use the implementation of Edmond’s algorithm from the NetworkX package [36], namely, the *minimum_spanning _arborescence* module. Such trees could potentially serve as a model selection criterion in case they are available from experimental data for the cell types under consideration.

### E. A stochastic method to estimate the Mean First Passage Time (MFPT)

The matrix method of computing the MFPT as given in section IIB [10, 26] is not scalable to larger network sizes. It requires storing a 2^*N*^ × 2^*N*^ matrix (where *N* is the number of nodes in the network), which is computationally infeasible even for networks of size 16. This restricts it’s use to evaluate the relative stability and perform model selection to smaller networks. Therefore, it is imperative to devise numerical or stochastic approaches to compute the MFPT from one state to another. We propose such a method here which incorporates stochastic and deterministic dynamics to mimic the MFPT obtained via the matrix method provided in section II B. Let *η* be the noise intensity parameter that acts on each gene, flipping it’s state with a probability *η*. At every time step, this noise *η* is applied to each gene independently (stochastic dynamics). If the noise does not alter the state of the network, then the state is updated via the assigned BFs (deterministic dynamics). More concisely, *perform deterministic dynamics only if stochastic dynamics does not alter the state of the network*. The following equation captures the aforementioned dynamics:

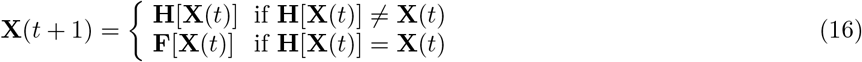

where, **X**(*t*) is the state of the network at time *t*, **H** and **F** are the stochastic and deterministic dynamics respectively. To obtain the first passage time from a state **X***j* to a state **X**_*i*_, let **X**(*t* = 0) = **X**_*j*_. Recursively apply Eq. (16) on **X**(0) until **X**(*t* = *t*’) = **X**_*i*_ such that *t* = *t*’ is the first instance at which state **X**_*i*_ is reached. Note that this gives us one trajectory from **X**_*j*_ to **X**_*i*_ and the number of steps in this trajectory is the first passage time. The mean first passage time from **X**_*j*_ to **X**_*i*_ is the mean of the first passage times over a large number of trajectories from **X**_*j*_ to **X**_*i*_.

### F. A greedy algorithm for Boolean model selection

Section IIIA highlights the challenge of model selection in larger models, even with a fixed network structure. Exhaustive exploration of ensembles, even after imposing constraints, is not feasible computationally and demands alternative approaches to apply our model selection framework. Here, we present an iterative greedy algorithm (Algorithm 1) to search for Boolean models that respect predefined constraints on the developmental landscape. The landscape constraints comprise of a set of pairwise inequalities or orderings. Our algorithm is inspired in part by the Potts model (a generalization of the Ising model) [37] because for each gene we assign a given set of Potts-like states corresponding to the possible Boolean functions specifying its output as a function of its inputs. The algorithm requires two types of information. The first is an initial Boolean model that generally will not respect the expected pairwise orderings. The second is a dictionary of the allowed BFs for each gene, the genes being sorted in ascending order according to their number of possible BFs. Genes associated with a single BF are removed from this dictionary since there is no decision to make for their Boolean rule to use. Note that the BFs for each output gene are obtained by imposing multiple constraints on their truth tables (see section III A).

#### Algorithm1 Greedy algorithm to find Boolean models that obey a specified developmental landscape

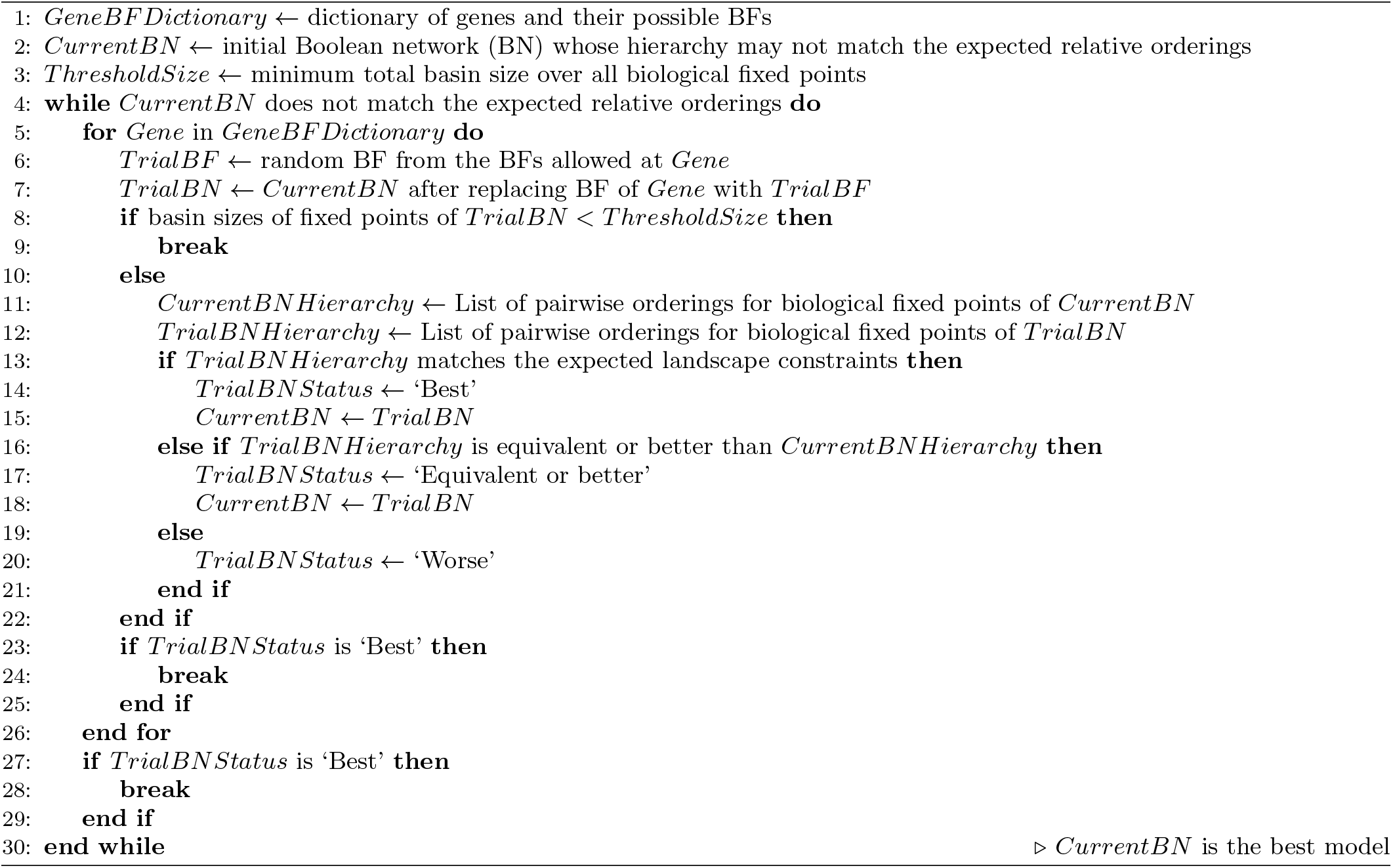

Our algorithm is iterative, consisting of repeatedly sweeping through the whole list of genes. For each gene the dictionary provides a corresponding list of possible *Potts-like states*, i.e., choices of BFs that are allowed based on the fixed point conditions. Note that this number of BFs varies from gene to gene. For each such choice of function, the algorithm replaces the current BF by this new choice and determines for the modified model all the pairwise orderings. If the resulting model is worse with respect to the developmental landscape, the change is rejected and the algorithm continues without implementing the modification. If instead the resulting model is better, then the algorithm makes it the current model. A model *A* is said to be worse than model *B* if *B* has a greater number of pairwise orderings that respect the predefined pairwise orderings than *A*. When using basin sizes, one can refine this notion further, especially when all inequalities are already satisfied, by imposing the inequalities to be satisfied *more strongly* (there are multiple ways of doing so) or by setting some appropriately motivated target values on those sizes. The algorithm continues until a satisfactory improved model is found.

## IV. RESULTS

### A. The five measures of relative stability are strongly correlated with each other

In this section, we compare how the 5 measures of relative stability presented in section IIB are correlated with each other. The Pearson correlation between these measures is computed and portrayed as a heatmap in Fig. 2 and shows that all measures are strongly correlated with one another. The measures are computed via exact means (matrix formulation) across all pairs of biological fixed points, for all 1275 models in the ensemble *Root_sc–NCF_* under a noise intensity parameter value of 1% (Fig. 2). Similar correlation heatmaps are constructed for the other *sc – NCF* ensembles described in section III B, namely, 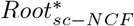 (SI Fig. S3), *Panc_sc–NCF_* (SI Fig. S4) and 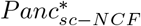 (SI Fig. S5), and the same conclusion ensues from all of them: the relative stability measures are all strongly correlated. Note that from Fig. 2 and SI Figs. S3, S4 and S5, *RS_BOA_* and *RS_BTR_* are perfectly correlated, a result we prove in the following subsection. The scatter plots for all pairs of measures (excluding *RS_BTR_* since it is equivalent to *RS_BOA_*) are provided as supplementary information for each of the 4 ensembles: 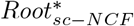 (SI Fig. S6), 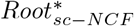 (SI Fig. S7), *Panc_sc–NCF_* (SI Fig. S8) and 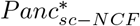 (SI Fig. S9).

**FIG. 2.**
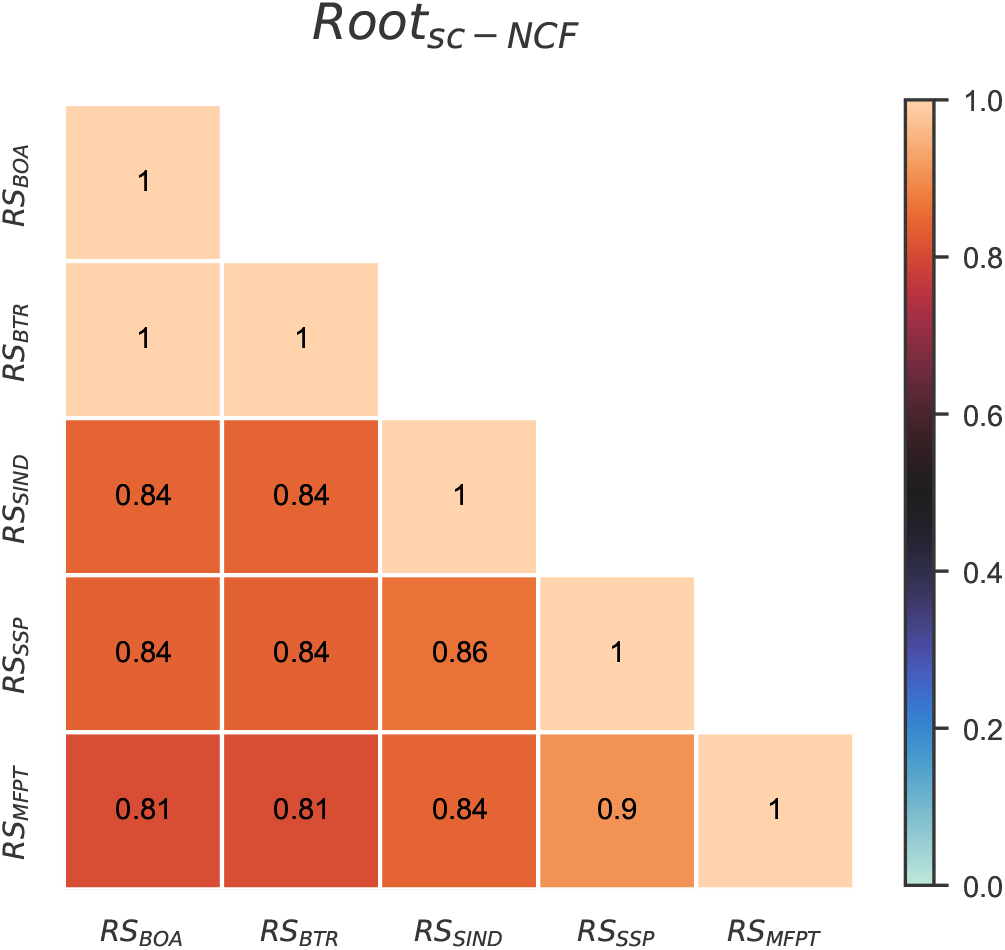
Pearson correlation between different pairs of relative stability measures for the ensemble *Root_sc–NCF_*. The rows and columns correspond to choices for the relative stability measures. These 5 measures are based on size of basin of attraction (*RS_BOA_*), basin transition rates (*RS_BTR_*), a stability index (*RS_SIND_*), steady state probabilities (*RS_SSP_*) and mean first passage times (*RS_MFPT_*). The heatmap indicates the value of the Pearson correlation coefficient between pairs of these measures. Note that these measures are computed by exact means across all pairs of biological fixed points, for all 1275 models in this ensemble *Root_sc–NCF_* using a noise intensity parameter value of 1%. The upper triangular portion of the heatmap is not displayed as the heatmap entries constitute a symmetric matrix. Furthermore, *RS_BOA_* and *RS_BTR_* are perfectly correlated, an observation which we prove theoretically by showing that *RS_BOA_* and *RS_BTR_* are in fact equivalent.

Next we show that this high degree of correlation is observed in particular when different pairs of fixed points are considered separately. The correlation heatmaps for the pairwise treatment of fixed points are provided as supplementary information for the different ensembles: *Root_sc–NCF_* (SI Fig. S10), 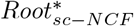 (SI Fig. S11), *Panc_sc–NCF_* (SI Fig. S12) and 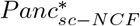 (SI Fig. S13). As before, scatter plots between the measures for different pairs of biological fixed points for the ensemble *Root_sc–NCF_* are provided as supplementary information. The pairs of biological fixed points are: QC and VI (SI Fig. S14), QC and CEI (SI Fig. S15), QC and CEpI (SI Fig. S16), VI and CEI (SI Fig. S17), VI and CEpI (SI Fig. S18) and, CEI and CEpI (SI Fig. S19), where QC corresponds to Quiescent center, VI to Vascular initials, CEI to Cortex-Endodermis initials, and CEpI to Columella epidermis initials.

Note that the preceding results were based on ensembles where the BF at each node is a *sc* – *NCF*. We have also tested whether the correlations are dependent on the type of BFs assigned to the nodes. So we studied the 4 ensembles following the procedure outlined in section IIIA for the *Arabidposis thaliana* RSCN and pancreas differentiation models, except that in this case, we restricted the BFs to sign conforming effective and unate functions (*sc* – *EUFs*) instead of *sc* – *NCFs*. The correlation heatmaps (using Pearson’s correlation coefficient) for the 4 ensembles are shown in the supplementary information: *Root_sc–EUF_* (SI Fig. S20), 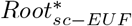 (SI Fig. S21), *Panc_sc–EUF_* (SI Fig. S22) and 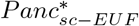 (SI Fig. S23). They reveal that despite using a different type of BF, the correlations between measures remains very high.

In sum, all the measures are in good agreement when comparing the relative stability of different pairs of fixed points; as a result, they will usually provide quite similar hierarchies in the landscapes. In case of disagreement, a preference for one measure over others must be rooted in biophysical or biological principles. As we shall see in the subsequent sections, the MFPT captures cell state transitions more naturally and also offers a richer representation of the landscapes in the form of trees.

#### The relative stability measure for the Basin Transition Rate (RS_BTR_) is identical to that of the Basin of Attraction (RS_BOA_)

As alluded to in the first paragraph of this section, we analytically prove here that *RS_BOA_* and *RS_BTR_* are in fact identical. Recall the definition of *RS_BTR_*(*u,v*) from section II B.

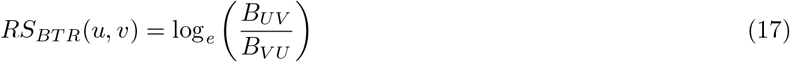

where 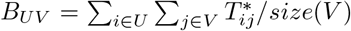 is the transition rate from basin *V* to basin *U*. In this definition *i* and *j* are states that belong to the basins *U* and *V* respectively. Note that transitions between states belonging to different basins are caused by the presence of noise, specified via the *perturbation* matrix **P** (see section II B). Hence 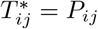 (if *i ∈ U* and *j ∈ V*), where *P_ij_* are the entries of the perturbation matrix **P**. Since **P** is a symmetric matrix, 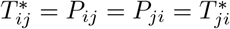. Now consider the expression for *V B_UV_*:

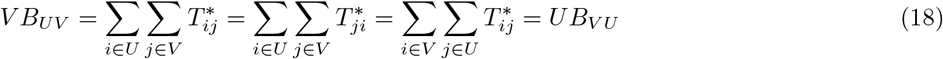

From Eq. (18), *B_UV_ /B_V U_* = *U/V*. Hence *RS_BTR_*(*u,v*) = *RS_BOA_*(*u, v*).

### B. Distribution of minimum spanning arborescences

In section IIID a prescription to generate a minimum spanning arborescence (MSA) from a MFPT matrix was provided. We apply this method to the 4 *sc* – *NCF* ensembles in section III B. Fig. 3 shows the distribution of MSAs when using the ensemble *Root_sc–NCF_* (where MFPT is computed using the ‘exact’ scheme with 1% noise). An immediate observation is that of 64 possible trees and 5 possible tree topologies, only 13 trees and 4 tree topologies occur in the ensemble. Not all of these 13 trees respect the relative stability conditions suggested by the underlying biology. For instance, the *QC* cell type is expected to be the least stable compared to the other cell types and is therefore expected to be the *root* of the tree. Therefore only 4 trees out of the 13 and 2 topologies out of 5 are likely to reflect the biological reality. Of these 4 trees, note that 2 of them appear most frequently in the distribution. For the ensemble 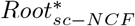 (SI Fig. S24), only 2 topologies are obtained, both of which occur more or less equally. The distribution of MSA for the Pancreas cell differentiation model is provided as supplementary figures: *Panc_sc–NCF_* (SI Fig. S25) and 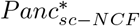 (SI Fig. S26).

**FIG. 3.**
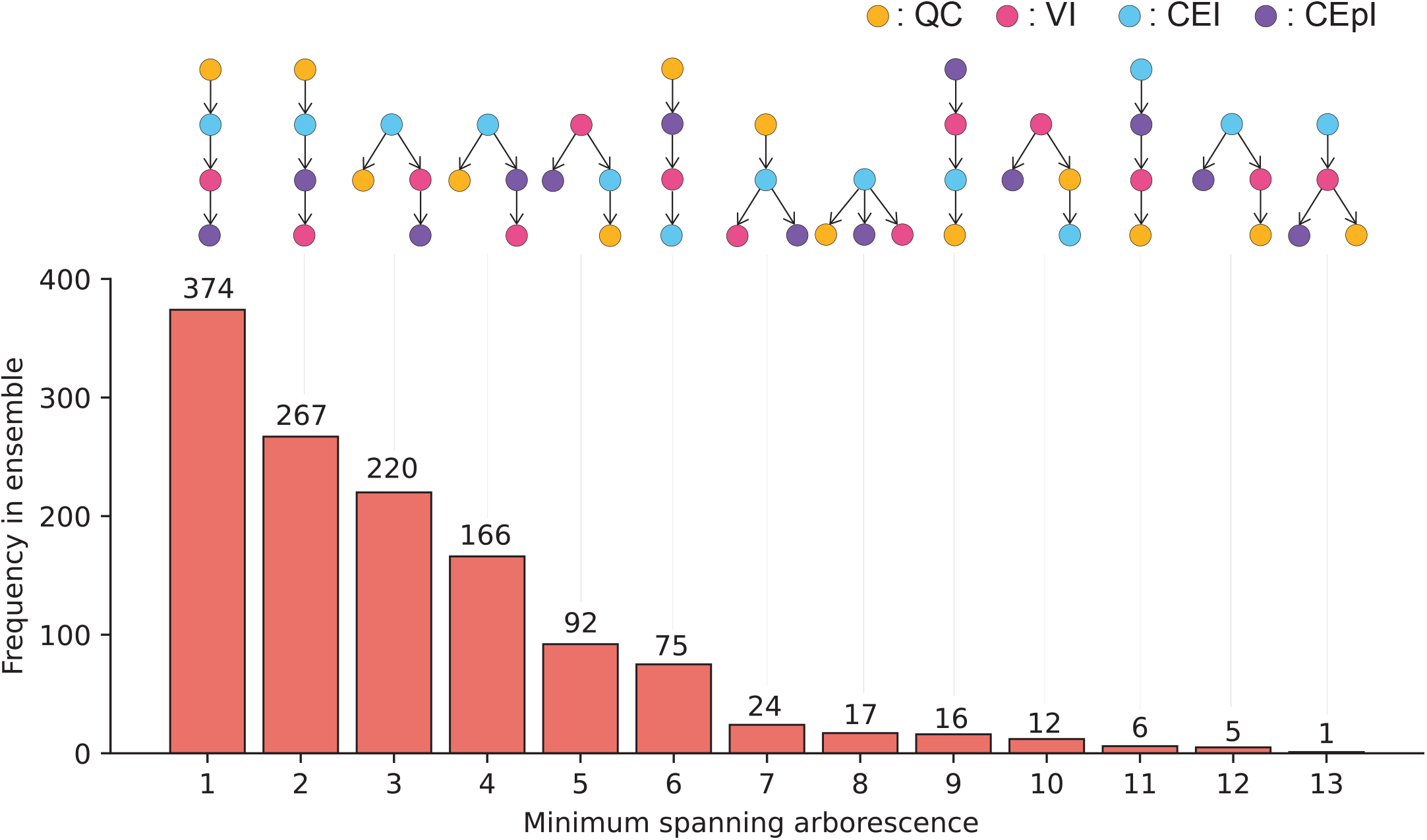
Frequency distribution of the minimum spanning arborescences (MSA) for the ensemble *Root_sc-NCF_*. The MSA for a Boolean model is constructed from a complete digraph whose vertices are biological fixed points and directed edges are the MFPTs (see section IIC for more details). The *x* axis labels the different MSAs that occur in the ensemble *Root_sc–NCF_*. Of the 64 possible (labeled and oriented) trees for 4 fixed points, only 13 occur in the *Root_sc–NCF_*. The *y* axis is the frequency of each of these trees among the 1275 models of the *Root_sc–NCF_* ensemble. The biological fixed points of the *Root_sc–NCF_* ensemble are as follows. QC: Quiescent center, VI: Vascular initials, CEI: Cortex-Endodermis initials and CEpI: Columella epidermis initials.

### C. Inferences drawn from MFPT are insensitive to changes in noise intensities

Before applying the MFPT to obtain a hierarchy of states for larger models, it is necessary to test it on smaller ones. First, we compute the *RS_MFPT_* values (for all pairs of biological fixed points) using the ‘exact’ method (see section II B), for different noise intensities ranging from 1% to 10%. Then for different pairs of noise values, we calculate the Pearson correlation coefficient between these *RS_MFPT_* values. The correlations are displayed via a heatmap in SI Fig. S24 for the ensemble *Root_sc–NCF_*. We find that for all pairs of noise values, *RS_MFPT_* values are strongly correlated for all 4 ensembles, namely, *Root_sc–NCF_* (SI Fig. S27), 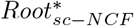 (SI Fig. S28), *Panc_sc–NCF_* (SI Fig. S29) and 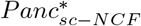 (SI Fig. S30). Note that as the difference between the pair of noise values increases, the correlation decreases, but only slightly.

Next, we obtain the set of partial orders (inequalities for all pairs of fixed points) using *RS_MFPT_* (computed using the ‘exact’ method) for different noise intensities ranging from 1% to 10%. For each pair of noise values, we compute the number and fraction of disagreements of their (partial) orders using the ensemble *Root_sc–NCF_*, and plot them as a heatmap in Fig. 4(a) and Fig. 4(b) respectively. Two observations follow from Fig. 4(b). First, the fraction of disagreements are quite low even for large differences in noise values. Second, the number and fraction of disagreements in the partial ordering increases with increasing differences in noise levels. These observations are recapitulated in other ensembles as well and are provided as the following supplementary figures: 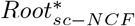 (SI Fig. S31), *Panc_sc–NCF_* (SI Fig. S32) and 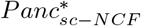 (SI Fig. S33).

**FIG. 4.**
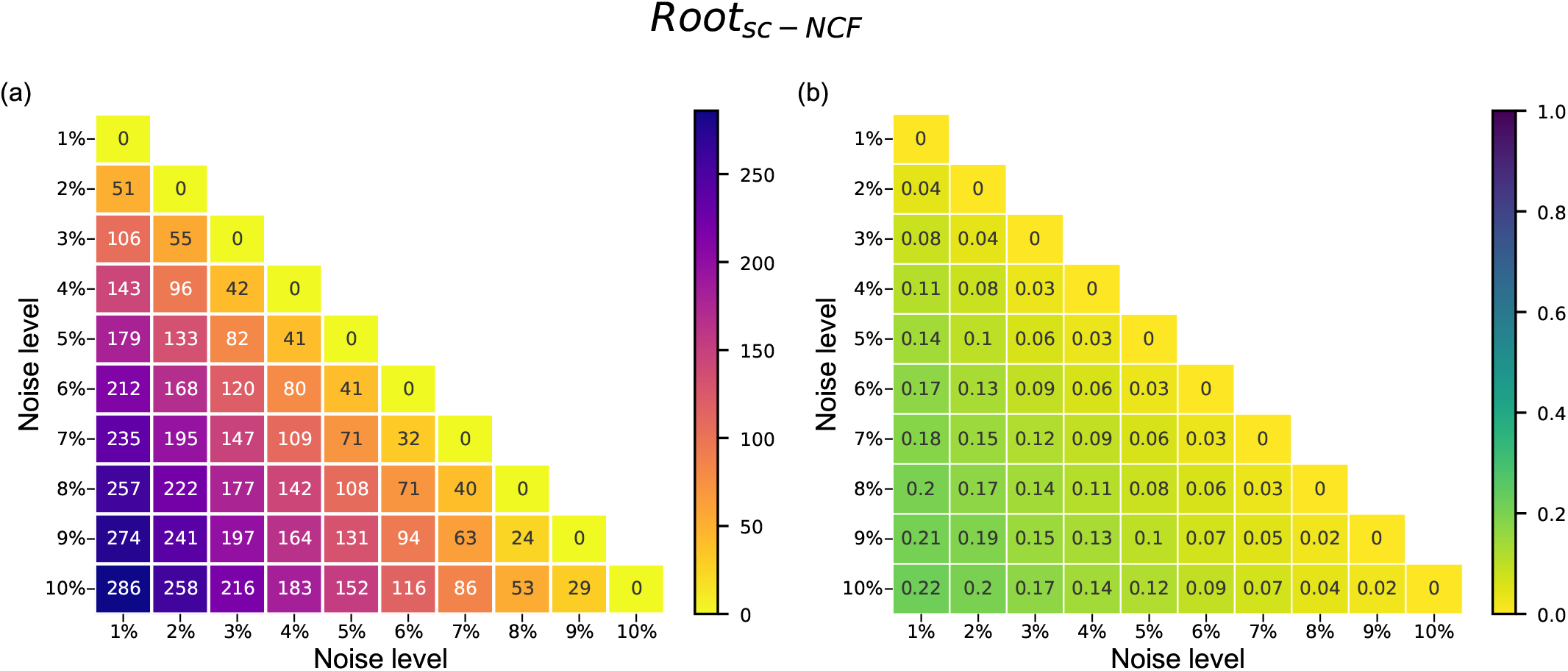
Number and fraction of models which differ in at least one comparison of partial ordering of the different biological fixed points when considering two different noise values, in the ensemble *Root_sc–NCF_*. The (partial) order of two fixed points is specified via the MFPT values for going from one to the other, computed here using an exact method. Rows and columns correspond to the noise intensity. The heatmap (a) gives the number of models (out of a total of 1275 models in the ensemble *Root_sc–NCF_*) that differ in at least one (partial) order across pairs of biological fixed points. The heatmap (b) provides the same information but using the fraction of such models. The upper triangular portions of the heatmaps are not displayed because they constitute a symmetric matrix.

These results reveal that the outcome of using a noise intensity of 5% will not differ much from using a noise intensity of 1%. This result is of practical importance for the stochastic simulations we perform using the MFPT in the later sections. Indeed, using a larger noise intensity greatly speeds up the stochastic simulations, while returning results compatible with smaller noise intensities.

### D. Comparison of the stochastic approach to the exact method of computing MFPT

Here we provide a comparison of the stochastic approach to compute MFPT described in section IIIE to it’s exact counterpart in section II B. First, for a given *sc* – *NCF* ensemble (in section III B), we compute *M_ij_* (MFPT from a biological fixed point *j* to another biological fixed point *i*) using both methods for various noise intensities (3%, 4%, 5%) and number of trajectories (500,1500, 2500) for the stochastic case. For a given noise and number of trajectories, the result is presented as a scatter plot (see SI Fig. S34) and as a table with Spearman and Kendall rank correlation coefficients (see SI Table S1) for the ensemble *Root_sc–NCF_*. Similar scatter plots and tables are provided for the other ensembles: 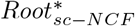 (SI Fig. S35, SI Table S2), *Panc_sc–NCF_* (SI Fig. S36, SI Table S3) and 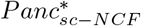 (SI Fig. S37, SI Table S4). As expected, these plots reveal that the stochastic method is in excellent agreement with the exact one, all the more so that one adds more and more trajectories.

To make this last claim more quantitative, we have used one model (chosen at random) from each of the 4 *sc* – *NCF* ensembles (in section III B) to test whether the *M_ij_* values obtained from the stochastic method are statistically reliable. In other words, we ensure that by choosing a sufficiently large number of trajectories, the statistical error associated with the stochastic method is within acceptable limits. This is clear from the bar plots shown as supplementary figures for each model from the 4 ensembles: *Root_sc–NCF_* (SI Fig. S38), 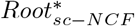 (SI Fig. S39), *Panc_sc–NCF_* (SI Fig. S40) and 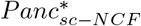 (SI Fig. S41).

Having established that the stochastic approach to compute MFPT is in concordance with the exact computation approach, we expect that the former can serve as a tool to derive developmental landscapes for larger Boolean network models.

### E. The *Arabidopsis thaliana* root development: A case study

We demonstrate on Boolean models of *Arabidopsis thaliana* root development GRNs [13, 27, 28] how our methods can be used to obtain landscapes and subsequently enable model selection using as constraints the expected developmental landscape of root cell differentiation. The models are given in the BoolNet format ([38]) in the supplementary information (2013 model [27] (SI Table S5), 2017 model [28] (SI Table S6) and 2020 model [13] (SI Table S7)). The network structure and biological fixed points (high auxin level) for the 2013 and 2017 models are shown in SI Figs. S42 (a) and (b) respectively. The 2020 model network structure and the biological fixed points at AUX=1 are shown in SI Fig. S43. Details of these model are provided in the supplementary information (see SI text, section 2).

#### Relative ordering of the biological fixed points by using different measures

Though the published Boolean models [13, 27, 28] of *Arabidopsis thaliana* DGRNs recover the expression patterns for the cell types observed in the root, it is appropriate to ask whether they conform to the expected developmental landscape. Indeed, a proposed Boolean model may reproduce the fixed points corresponding to cell types but their relative stabilities may not obey the hierarchies derived from the expected developmental landscape. In view of this, we compute the landscape hierarchies in the 2013 RSCN model, 2017 RAM (Root Apical Meristem) model and 2020 RSCN Boolean models of *Arabidopsis thaliana*. Fig. 5 shows the hierarchies obtained via 2 methods, first, the basin of attraction, and second, the MSA using MFPTs. The MSA was constructed using MFPTs obtained via the stochastic method at 5% noise using 10000 trajectories. From those values we can ascertain whether a computed hierarchy conforms to biological expectations. Experiments have shown that QCs undergo asymmetric cell division wherein one of the daughter cells differentiates to a another cell type and the other does not (maintaining the pool of QC cells in the RSCN) [35]. It is also known that the de-differentiation of other cell types to QC is rare. With this information we can construct the expected partial landscape of root development: QC is relatively less stable compared to all other cell types. We find that the 2013 and 2017 models are able to recover this expected partial landscape while using as relative stability measures basin of attraction and MFPT (to obtain MSAs) (see Fig. 5). However the 2020 model, though the most recent (and all these models have been reconstructed by the same group of scientists), does not recover the partial landscape, be-it via basin size or via MFPT as can be seen from Fig. 5. Specifically, the QC is more stable than the CEI/Endodermis, a hierarchy that is incompatible with experimental findings. It is also known that the cell type *Transition domain* (C. PTD2) is expected to more stable than the cell type *Central Pro-vascular initials* (C. PPD) [13]. For this case it is worth noting that relative stability associated with the basin of attraction of these cell types (C. PTD2 *<* C. PPD) violates the expected hierarchy (C. PPD *<* C. PTD2) whereas the relative stability associated with MFPT is in concordance with the expected hierarchy. This suggests that the developmental landscape may at least not have been considered during the reconstruction of this last Boolean model published in 2020. It also leads one to ask: how do we search for models that recapitulate the expected landscape?

**FIG. 5.**
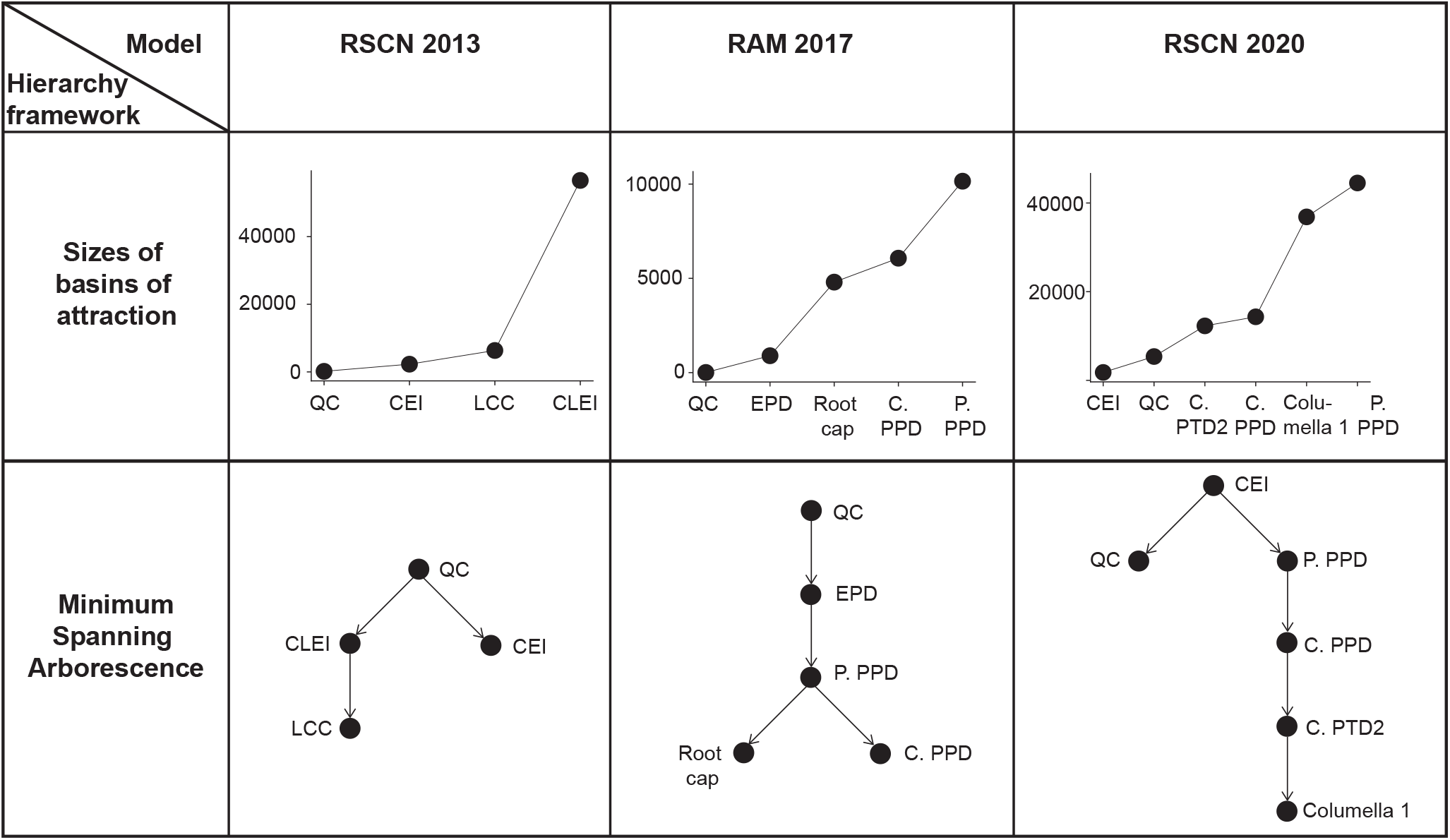
Hierarchy of fixed points based on basin sizes and minimum spanning arborescence for the 2013, 2017 and 2020 models. The first row of this table shows the hierarchy of fixed points according to the basin size. For the 2013 and 2017 models, the QC has the smallest basin size whereas for the 2020 model, the CEI has the smallest basin size. The second row gives the hierarchy of states according to the MSA. The MSA was constructed using MFPTs obtained using the stochastic method under a noise intensity parameter value of 5% and averaging over 10000 trajectories. For the 2013 and 2017 models, the QC is at the root of the tree whereas for the 2020 model it is the CEI that is at the root of the tree. The expansions of the abbreviations for the 2013 model are as follows: QC: Quiescent center, CEI: Cortex-endodermis initials, LCC: Lateral root-cap and CLEI: Columella and lateral root-cap-epidermis initials. The expansions of the abbreviations for the 2017 model are as follows: EPD: Endodermis Proliferation Domain, P. PPD: Peripheral Pro-vascular Proliferation Domain, C. PPD: Central Pro-vascular Proliferation Domain. The expansions of the abbreviations for the 2020 model are as follows: CEI: Cortex/endodermis initial cell, P. PPD: Peripheral Pro-vascular initials, C. PPD: Central Pro-vascular initials, C. PTD2: Transition domain, Columella 1: Columella initials.

#### A greedy algorithm proposes many models satisfying the expected developmental landscape

Here, we address the impending question of model selection using the developmental landscape constraint by implementing an iterative greedy algorithm (section III F, Algorithm 1) on the 2020 model of the *Arabidopsis thaliana* RSCN. To begin with, we define an ensemble of Boolean models using the procedure depicted in Fig. 6(a) (see section IIIA for details). In particular, we impose that the type of BF assigned to a gene be the same as the one assigned by García-Gómez *et. al* [13] in their 2020 Boolean model [13] (see SI Table S7 for the BFs). With that procedure, all genes were constrained to be sc-NCFs except for ARF5 which was of the sc-RoF type. After applying all the constraints in Fig. 6(a) we find that of 18 genes, 8 allowed a single BF and the remaining allowed for more. Fig. 6(b) shows the number of allowed BFs for each gene. Combining the BFs at each node results in an ensemble of size of the order 10^10^, too large for an exhaustive search. Thus, we propose an alternate approach to efficiently explore this ensemble based on an iterative greedy process.

**FIG. 6.**
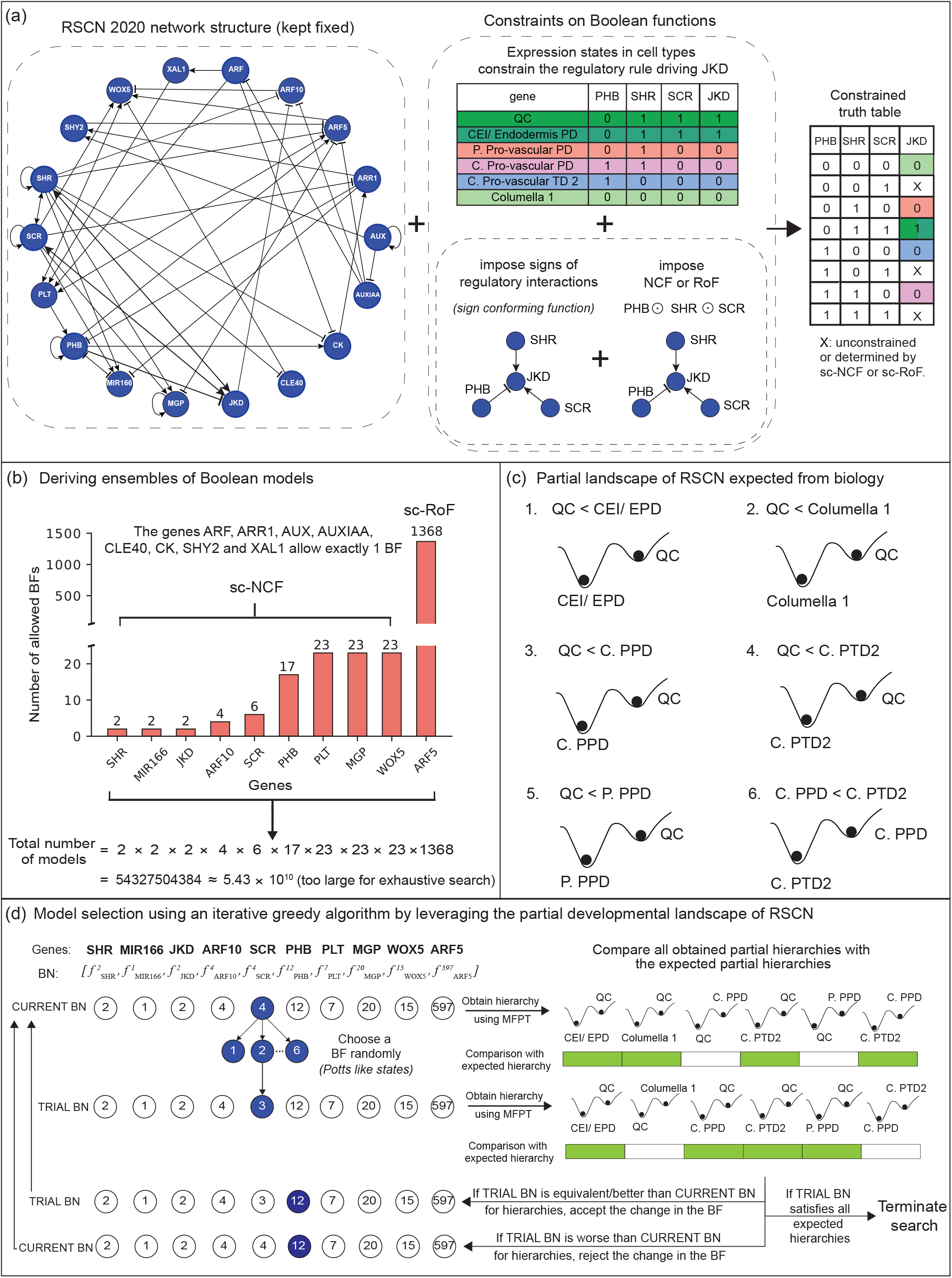
Workflow of our methodology for model selection, illustrated on the 2020 model of the Boolean GRN of *Arabidopsis thaliana* RSCN. **(a)** Procedure to generate ensembles of Boolean models that recover the desired attractors. Keeping the network structure of the reconstructed GRN fixed, impose the biological fixed point constraints on the truth tables of all genes. Next, impose the constraint that the BF should conform to the signs of regulatory interactions that are input to the gene under consideration. Lastly, impose the constraint on the type of BF, preferably minimally complex BFs. In sum, we get a constrained truth table as shown. Note that ‘X’ may or may not be determined when imposing the type of minimally complex BF. **(b)** The bar plot shows for each gene the number of BFs that have at least two allowed BFs after imposing all the constraints mentioned in (a). For this 2020 model, all genes are constrained to be sc-NCF except for ARF5 which is constrained to be of the sc-RoF type. The ensemble of models that recover the biological fixed points is obtained by generating all combinations of these BFs over all the genes. **(c)** The partial expected landscape represented as inequalities between different pairs of fixed points. Here QC is the least stable cell type among all the other cell types. **(d)** Schema of the iterative greedy algorithm to perform model selection within the ensemble. Each Boolean network is represented as a vector of BFs with each entry given by *f* where *i* ∈ {1, *m*}, *g* is the gene and *m* is the number of allowed BFs for that gene. The vector shown represents an instance of the Boolean network (BN) that we denote as CURRENT BN. From the CURRENT BN, we generate a TRIAL BN by assigning to a node a BF randomly chosen from the allowed BFs at that node (in this case, gene SCR). The partial hierarchies of CURRENT BN and TRIAL BN are obtained using the MFPT and are compared with the expected hierarchies provided in (c) which is shown on the right panel. The green colored rectangles indicate that the obtained hierarchy is in agreement with the expected hierarchy and white rectangles denote otherwise. In this instance, the TRIAL BN is as good as the CURRENT BN and so the change in the BF is accepted. But in a scenario where the TRIAL BN is worse than the CURRENT BN, the change in BF is rejected. For both these cases, the iterative scheme is continued by generating a TRIAL BF for the following gene (PHB in the case illustrated in (d)). The remaining scenario is one where TRIAL BN satisfies all the expected hierarchies, in which case the algorithm is terminated.

Our greedy approach requires as inputs the set of expected hierarchies and an initial model (which may satisfy some of the expected hierarchies). A partial landscape of the RSCN is available (as mentioned in the previous section) and is provided in Fig. 6(c). For our initial condition, we choose the model proposed in García-Gómez *et. al* [13]. Starting from this model we iterate through each gene, each time randomly selecting a BF from the set of allowed BFs at that gene and rejecting or accepting the BF in case the resulting model is worse or equivalent (or better) than the current model, respectively (see Fig. 6). The hierarchies (partial landscape) for each model were obtained using the MFPT computed via the stochastic method (at 5% noise intensity with 2500 trajectories). We terminate the algorithm as soon as a model satisfies all the expected partial hierarchies. Out of 1000 runs starting from the initial condition, we obtain 990 distinct models which conform to the expected partial landscape. These 990 models are provided in the supplementary information, each one as a list of BFs, one for each gene (see SI Table S8).

To summarize, the developmental landscape coupled with stochastic methods for computing relative stability can be used to perform model selection, and is applicable even to large networks.

## V. DISCUSSION AND CONCLUSIONS

The principal findings of our work can be stated as follows. First, the 5 different measures of relative stability [26], based on size of Basin of Attraction (BOA), Steady State Probability (SSP), Mean First Passage Time (MFPT), Basin Transition Rate (BTR) and Stability Index (SIND), are strongly correlated with each other. Second, noise intensities need not be particularly small in order to reliably identify the hierarchies between cell types (fixed points of the developmental GRN (DGRN)). Third, MFPT can be used to generate cellular lineage trees in addition to providing a hierarchy of cell states. Fourth, the ‘exact’ method to calculate MPFT based on matrix algebra is not feasible for large networks, whereas our alternative stochastic approach does not suffer from such limitations, and in addition, is particularly simple to implement. Fifth, we developed an iterative greedy search algorithm that can identify models conforming to the expected developmental landscape from a large *ensemble* of biologically plausible models. Last, multiple Boolean models were produced that satisfied the expected developmental landscape of the *Arabidopsis* root stem cell niche.

The inference of Boolean networks, both their structure and dynamics, has been a major problem in Boolean modeling and has been heavily researched for at least two decades [39–42]. More recently, there is a rising interest in inferring the Boolean functions using biological data as dynamical constraints [43–45]. The epigenetic landscape of Boolean models has also been quantified [46], particularly in some model systems such as the flower specification GRN of *Arabidopsis thaliana* [47, 48] and RSCN [49]. However not much endeavor has gone into leveraging the epigenetic landscape to infer viable Boolean models, barring few exceptions such as [10]. Furthermore, the relative stability has hardly been explored in the context of serving as a criteria for model selection, the main purpose of the work presented in this paper.

During the reconstruction of Boolean models of DGRNs from biological data, modelers are often forced to make arbitrary choices. Specifically, for a number of target genes, they will have to select just one of the many equally plausible logic rules allowing the dynamical system to recover the desired fixed points, thereby introducing their subjective proclivities into the reconstruction workflow. The present work advocates a more streamlined process to assign logic rules to genes by leveraging relative stability constraints derived from biological developmental landscapes. This idea had genesis in [10] where it was only applied to a minimal 5-gene Boolean DGRN of Pancreas cell differentiation. Since then, no effort has gone into scaling such a methodology to larger models or refining it. Here we build upon that work, taking as a case study an 18 gene Boolean model of Arabidopsis thaliana RSCN [13]. Although that model is quite recent, it does not satisfy the expected hierarchies between cell types. With the help of a simple but very effective search and select algorithm, we were able to improve that model to obtain multiple DGRNs having satisfactory developmental landscapes.

The addition of further constraints on the DGRNs, such as robustness to noise, will restrict even more the space of relevant models. Indeed, our methodology is very flexible and scales up to larger networks, the only real caveat being that any such additional constraints must be computationally tractable.

Another novel aspect of this contribution is that MFPTs can be exploited to obtain a minimum spanning arborescence (MSA), whose structure offers a far richer picture of developmental hierarchies than just a linear order on the fixed points (see Fig. 5). Surely such lineage trees could serve as a strong constraint for model selection, subject to their availability *a priori*. Note that some biological situations may require going beyond trees, that is, allowing for loops. Such cases arise when a cell type is the end result of multiple lineages, a phenomenon referred to as convergence of cell fates. Accounting for loops is also sometimes necessary in evolutionary phylogenies [50]. The associated methods allow one to infer reticulated graphs rather than tree graphs. Another context where loops are key is when a system responds to a transient stress: different cell types trace out different trajectories but finally return to their initial state. Our measures of stability may also be extended to other modeling frameworks where gene expression states could take multiple discrete or continuous values, and where temporal dynamics is continuous (as in ODEs). Finally our work also has practical consequences in the field of cellular reprogramming [10, 26, 51–53] to design rational approaches to alter the developmental landscape to facilitate desired cell state transitions.

In conclusion, these results hold promise for developing standardized workflows for Boolean model reconstruction of DGRNs by leveraging biological constraints and computational methods.

## DATA AND CODE AVAILABILITY

All data and codes needed to reproduce the results in this manuscript are deposited in GitHub and are available at: https://github.com/asamallab/LDLM.

## ACKNOWLEDGMENTS

Areejit Samal acknowledges support from the Max Planck Society, Germany, through the award of a Max Planck Partner Group in Mathematical Biology, and the Department of Atomic Energy, Government of India. IPS2 benefits from the support of Saclay Plant Sciences-SPS (ANR-17-EUR-0007).

## AUTHORS’ CONTRIBUTIONS

Designed the research: Aj.S., P.S., O.C.M. and Ar.S. Performed the research: Aj.S., P.S., O.C.M. and Ar.S. Performed the computations: Aj.S. and P.S. Wrote the paper: Aj.S., P.S., O.C.M. and Ar.S.

## COMPETING INTEREST

The authors declare no competing interest.

## Notes

https://github.com/asamallab/LDLM

### Competing Interest Statement

The authors have declared no competing interest.

