## Supplementary Information for "Leveraging Developmental Landscapes for Model Selection in Boolean Gene Regulatory Networks"

### 1. BIOLOGICALLY MEANINGFUL BOOLEAN FUNCTIONS

#### Effective functions

A BF $f$ with $k$ inputs is said to be *effective* if each input $i$ elicits a change to the output under some context of all other inputs [54]. Mathematically,

$$\forall\ i \in [1,k],\ \exists\ \mathbf{x} \in \{0,1\}^k \text{ with } x_i = 0,\ f(\mathbf{x}) \neq f(\mathbf{x} + \mathbf{e}_i). \tag{1}$$

Here $\mathbf{e}_i \in \{0,1\}^k$ denotes the unit vector associated to the component of index $i$ having all entries set to 0 except for entry $i$.

#### Unate functions

A BF $f$ with $k$ inputs is said to be *unate* if each input is either increasing monotone (activatory) or decreasing monotone (inhibitory) [55]. An input $i$ (or variable $x_i$) is increasing monotone (or activatory) if:

$$\forall\ \mathbf{x} \in \{0,1\}^k \text{ with } x_i = 0,\ f(\mathbf{x}) \leq f(\mathbf{x} + \mathbf{e}_i), \tag{2}$$

and decreasing monotone (or inhibitory) if:

$$\forall\ \mathbf{x} \in \{0,1\}^k \text{ with } x_i = 0,\ f(\mathbf{x}) \geq f(\mathbf{x} + \mathbf{e}_i). \tag{3}$$

#### Read-once functions

A BF of $k$ variables is said to be *read-once* if it can be represented by a Boolean expression, using the operations of conjunction, disjunction and negation, in which every variable appears exactly once [56]. Mathematically, a BF $f = f(x_1, x_2, \ldots, x_k)$ is a RoF if and only if there is a permutation $\sigma$ on its inputs $\{1, 2, \ldots, k\}$ such that

$$f = x_{\sigma(1)} \odot x_{\sigma(2)} \odot x_{\sigma(3)} \ldots \odot x_{\sigma(k)}, \tag{4}$$

where $\odot$ represents the AND ($\wedge$) or OR ($\vee$) operator and in this expression we have omitted all parentheses that may need to be placed to define the function.

---

**Nested canalyzing functions**

A BF $f$ with $k$ inputs is *nested canalyzing* [57, 58] with respect to a permutation $\sigma$ on its inputs $\{1, 2, \ldots, k\}$ if:

$$f(\mathbf{x}) = \begin{cases} b_1 & \text{if } x_{\sigma(1)} = a_1, \\ b_2 & \text{if } x_{\sigma(1)} \neq a_1, x_{\sigma(2)} = a_2, \\ b_3 & \text{if } x_{\sigma(1)} \neq a_1, x_{\sigma(2)} \neq a_2, x_{\sigma(3)} = a_3, \\ \vdots & \\ b_k & \text{if } x_{\sigma(1)} \neq a_1, x_{\sigma(2)} \neq a_2, \ldots, x_{\sigma(k)} = a_k, \\ \overline{b}_k & \text{if } x_{\sigma(1)} \neq a_1, x_{\sigma(2)} \neq a_2, \ldots, x_{\sigma(k)} = \overline{a}_k. \end{cases} \tag{5}$$

In the above equation, $a_1, a_2, \ldots, a_k$ are the canalyzing input values and $b_1, b_2, \ldots, b_k$ are the canalyzed output values for inputs $\sigma(1), \sigma(2), \ldots, \sigma(k)$ in the permutation $\sigma$ of the $k$ inputs. Here, $\overline{a}_k$ and $\overline{b}_k$ are the complements of the Boolean values $a_k$ and $b_k$, respectively.

### 2. A BRIEF OVERVIEW OF BOOLEAN MODELS OF ROOT DEVELOPMENT

As case studies, we considered 3 Boolean models of the *Arabidopsis thaliana* root development that have been reconstructed and published between the years 2013 and 2020 [13, 27, 28]. These include: a 2013 RSCN (Root Stem Cell Niche) model [27], a 2017 RAM (Root Apical Meristem) model [28] and a 2020 RSCN model [13]. The 2013 [27] and 2017 [28] studies presented multiple Boolean models from which we chose one per published article. Our choice was based on 2 simple criteria. One, that the model should recover most of the expected biological fixed points with the levels of the phytohormone auxin being high (see [13]). Other, that the fraction of state space occupied by the basins of biological fixed points having high auxin levels be the largest among all models proposed in that article. Of the 10 models in the 2013 [27] publication, those criteria led to choosing *model 4* (see SI Table S5). Of the 2 models in the 2017 [28] publication, it was the *GHRN1 model* that satisfied these criteria. In the 2020 article [13], only 1 model was provided. The models are given in the BoolNet format ([38]) in the supplementary information (2013 model (SI Table S5), 2017 model (SI Table S6) and 2020 model (SI Table S7)). The network structure and biological fixed points (high auxin level) for the 2013 and 2017 models are shown in SI Figs. S42(a) and (b) respectively. The 2020 model network and the biological fixed points at AUX=1 are shown in SI Fig. S43. Starting from 2010, the root development models have been refined over the years by the addition of new genes and interactions, resulting in more diverse cell types as can be seen in SI Figs. S42 and S43.

TABLE S1. **Correlation between the 'exact' and stochastic methods to compute MFPT using the $Root_{sc-NCF}$ ensemble**. The Spearman and Kendall rank correlation coefficients for MFPT values computed by the exact method and stochastic method for the ensemble $Root_{sc-NCF}$ for different combinations of number of trajectories and noise intensities is given. The associated p-values of the correlation coefficients are also provided. The last column of the table gives the number of MFPT values obtained from 'exact' computations that lie outside the error bar obtained via the stochastic method.

| Noise (%) | Trajectories | Spearman | | Kendall | | Number of analytical MFPT values outside error bar (Among 15300 values) |
|---|---|---|---|---|---|---|
| | | correlation ($\rho$) | p-value | correlation ($\tau$) | p-value | |
| 3 | 500 | 0.9969 | $< 10^{-323}$ | 0.9592 | $< 10^{-310}$ | 4950 |
| 3 | 1000 | 0.9982 | $< 10^{-323}$ | 0.9703 | $< 10^{-310}$ | 4825 |
| 3 | 1500 | 0.9987 | $< 10^{-323}$ | 0.9749 | $< 10^{-310}$ | 4935 |
| 3 | 2000 | 0.9990 | $< 10^{-323}$ | 0.9780 | $< 10^{-310}$ | 4938 |
| 3 | 2500 | 0.9991 | $< 10^{-323}$ | 0.9801 | $< 10^{-310}$ | 4884 |
| 4 | 500 | 0.9975 | $< 10^{-323}$ | 0.9615 | $< 10^{-310}$ | 4872 |
| 4 | 1000 | 0.9986 | $< 10^{-323}$ | 0.9718 | $< 10^{-310}$ | 4928 |
| 4 | 1500 | 0.9990 | $< 10^{-323}$ | 0.9768 | $< 10^{-310}$ | 4919 |
| 4 | 2000 | 0.9992 | $< 10^{-323}$ | 0.9796 | $< 10^{-310}$ | 4851 |
| 4 | 2500 | 0.9994 | $< 10^{-323}$ | 0.9816 | $< 10^{-310}$ | 4853 |
| 5 | 500 | 0.9976 | $< 10^{-323}$ | 0.9616 | $< 10^{-310}$ | 4891 |
| 5 | 1000 | 0.9988 | $< 10^{-323}$ | 0.9725 | $< 10^{-310}$ | 4885 |
| 5 | 1500 | 0.9991 | $< 10^{-323}$ | 0.9774 | $< 10^{-310}$ | 4861 |
| 5 | 2000 | 0.9993 | $< 10^{-323}$ | 0.9803 | $< 10^{-310}$ | 4851 |
| 5 | 2500 | 0.9994 | $< 10^{-323}$ | 0.9822 | $< 10^{-310}$ | 4895 |

TABLE S2. **Correlation between the 'exact' and stochastic methods to compute MFPT using the $Root^{*}_{sc-NCF}$ ensemble**. The Spearman and Kendall rank correlation coefficients for MFPT values computed by the exact method and stochastic method for the ensemble $Root^{*}_{sc-NCF}$ for different combinations of number of trajectories and noise intensities is given. The associated p-values of the correlation coefficients are also provided. The last column of the table gives the number of MFPT values obtained from 'exact' computations that lie outside the error bar obtained via the stochastic method.

| Noise (%) | Trajectories | Spearman | | Kendall | | Number of analytical MFPT values outside error bar (Among 2040 values) |
|---|---|---|---|---|---|---|
| | | correlation ($\rho$) | p-value | correlation ($\tau$) | p-value | |
| 3 | 500 | 0.9826 | $< 10^{-323}$ | 0.9154 | $< 10^{-310}$ | 623 |
| 3 | 1000 | 0.9878 | $< 10^{-323}$ | 0.9322 | $< 10^{-310}$ | 632 |
| 3 | 1500 | 0.9903 | $< 10^{-323}$ | 0.9406 | $< 10^{-310}$ | 655 |
| 3 | 2000 | 0.9926 | $< 10^{-323}$ | 0.9483 | $< 10^{-310}$ | 665 |
| 3 | 2500 | 0.9943 | $< 10^{-323}$ | 0.9548 | $< 10^{-310}$ | 613 |
| 4 | 500 | 0.9869 | $< 10^{-323}$ | 0.9260 | $< 10^{-310}$ | 628 |
| 4 | 1000 | 0.9913 | $< 10^{-323}$ | 0.9417 | $< 10^{-310}$ | 651 |
| 4 | 1500 | 0.9936 | $< 10^{-323}$ | 0.9510 | $< 10^{-310}$ | 631 |
| 4 | 2000 | 0.9946 | $< 10^{-323}$ | 0.9557 | $< 10^{-310}$ | 652 |
| 4 | 2500 | 0.9958 | $< 10^{-323}$ | 0.9610 | $< 10^{-310}$ | 640 |
| 5 | 500 | 0.9849 | $< 10^{-323}$ | 0.9204 | $< 10^{-310}$ | 669 |
| 5 | 1000 | 0.9902 | $< 10^{-323}$ | 0.9382 | $< 10^{-310}$ | 679 |
| 5 | 1500 | 0.9937 | $< 10^{-323}$ | 0.9504 | $< 10^{-310}$ | 632 |
| 5 | 2000 | 0.9948 | $< 10^{-323}$ | 0.9553 | $< 10^{-310}$ | 644 |
| 5 | 2500 | 0.9956 | $< 10^{-323}$ | 0.9597 | $< 10^{-310}$ | 629 |

TABLE S3. **Correlation between the 'exact' and stochastic methods to compute MFPT using the $Panc_{sc-NCF}$ ensemble**. The Spearman and Kendall rank correlation coefficients for MFPT values computed by the exact method and stochastic method for the ensemble $Panc_{sc-NCF}$ for different combinations of number of trajectories and noise intensities is given. The associated p-values of the correlation coefficients are also provided. The last column of the table gives the number of MFPT values obtained from 'exact' computations that lie outside the error bar obtained via the stochastic method.

| **Noise (%)** | **Trajectories** | **Spearman** | | **Kendall** | | **Number of analytical MFPT values outside error bar (Among 15600 values)** |
|---|---|---|---|---|---|---|
| | | **correlation ($\rho$)** | **p-value** | **correlation ($\tau$)** | **p-value** | |
| 3 | 500 | 0.9992 | $< 10^{-323}$ | 0.9757 | $< 10^{-310}$ | 6874 |
| 3 | 1000 | 0.9995 | $< 10^{-323}$ | 0.9828 | $< 10^{-310}$ | 6851 |
| 3 | 1500 | 0.9997 | $< 10^{-323}$ | 0.9859 | $< 10^{-310}$ | 6842 |
| 3 | 2000 | 0.9998 | $< 10^{-323}$ | 0.9878 | $< 10^{-310}$ | 6871 |
| 3 | 2500 | 0.9998 | $< 10^{-323}$ | 0.9892 | $< 10^{-310}$ | 6764 |
| 4 | 500 | 0.9990 | $< 10^{-323}$ | 0.9739 | $< 10^{-310}$ | 6961 |
| 4 | 1000 | 0.9995 | $< 10^{-323}$ | 0.9817 | $< 10^{-310}$ | 6854 |
| 4 | 1500 | 0.9997 | $< 10^{-323}$ | 0.9852 | $< 10^{-310}$ | 6738 |
| 4 | 2000 | 0.9998 | $< 10^{-323}$ | 0.9869 | $< 10^{-310}$ | 6938 |
| 4 | 2500 | 0.9998 | $< 10^{-323}$ | 0.9883 | $< 10^{-310}$ | 6958 |
| 5 | 500 | 0.9989 | $< 10^{-323}$ | 0.9723 | $< 10^{-310}$ | 6938 |
| 5 | 1000 | 0.9995 | $< 10^{-323}$ | 0.9806 | $< 10^{-310}$ | 6888 |
| 5 | 1500 | 0.9996 | $< 10^{-323}$ | 0.9841 | $< 10^{-310}$ | 6777 |
| 5 | 2000 | 0.9997 | $< 10^{-323}$ | 0.9860 | $< 10^{-310}$ | 6872 |
| 5 | 2500 | 0.9998 | $< 10^{-323}$ | 0.9875 | $< 10^{-310}$ | 6889 |

TABLE S4. **Correlation between the 'exact' and stochastic methods to compute MFPT using the $Panc^{*}_{sc-NCF}$ ensemble**. The Spearman and Kendall rank correlation coefficients for MFPT values computed by the exact method and stochastic method for the ensemble $Panc^{*}_{sc-NCF}$ for different combinations of number of trajectories and noise intensities is given. The associated p-values of the correlation coefficients are also provided. The last column of the table gives the number of MFPT values obtained from 'exact' computations that lie outside the error bar obtained via the stochastic method.

| **Noise (%)** | **Trajectories** | **Spearman** | | **Kendall** | | **Number of analytical MFPT values outside error bar (Among 654 values)** |
|---|---|---|---|---|---|---|
| | | **correlation ($\rho$)** | **p-value** | **correlation ($\tau$)** | **p-value** | |
| 3 | 500 | 0.9984 | $< 10^{-323}$ | 0.9715 | $< 10^{-310}$ | 213 |
| 3 | 1000 | 0.9990 | $< 10^{-323}$ | 0.9788 | $< 10^{-310}$ | 191 |
| 3 | 1500 | 0.9993 | $< 10^{-323}$ | 0.9829 | $< 10^{-310}$ | 210 |
| 3 | 2000 | 0.9995 | $< 10^{-323}$ | 0.9845 | $< 10^{-310}$ | 216 |
| 3 | 2500 | 0.9995 | $< 10^{-323}$ | 0.9859 | $< 10^{-310}$ | 227 |
| 4 | 500 | 0.9987 | $< 10^{-323}$ | 0.9730 | $< 10^{-310}$ | 187 |
| 4 | 1000 | 0.9992 | $< 10^{-323}$ | 0.9795 | $< 10^{-310}$ | 213 |
| 4 | 1500 | 0.9994 | $< 10^{-323}$ | 0.9828 | $< 10^{-310}$ | 214 |
| 4 | 2000 | 0.9996 | $< 10^{-323}$ | 0.9860 | $< 10^{-310}$ | 199 |
| 4 | 2500 | 0.9996 | $< 10^{-323}$ | 0.9867 | $< 10^{-310}$ | 206 |
| 5 | 500 | 0.9987 | $< 10^{-323}$ | 0.9732 | $< 10^{-310}$ | 217 |
| 5 | 1000 | 0.9993 | $< 10^{-323}$ | 0.9804 | $< 10^{-310}$ | 226 |
| 5 | 1500 | 0.9995 | $< 10^{-323}$ | 0.9838 | $< 10^{-310}$ | 212 |
| 5 | 2000 | 0.9996 | $< 10^{-323}$ | 0.9868 | $< 10^{-310}$ | 188 |
| 5 | 2500 | 0.9997 | $< 10^{-323}$ | 0.9872 | $< 10^{-310}$ | 203 |

TABLE S5. **Boolean functions for the 2013 RSCN model in the BoolNet format.** The column with header 'Target gene name' contains the list of genes whose regulation is captured by the corresponding row entry in the column 'Regulatory logic rule'. The symbols &, | and ! correspond to the logic operators AND, OR and NOT respectively.

| Serial Number | Target gene name | Regulatory logic rule |
|---|---|---|
| 1 | SHR | JKD \| (IAA5 & (!CYA \| !SHR)) |
| 2 | SCR | (JKD & (MGP \| WOX5)) \| (SYS & (MGP & WOX5)) |
| 3 | JKD | (SYS & JKD & (!PHB \| MGP)) \| (SYS & MGP) \| (!PHB & MGP) |
| 4 | MGP | JYI \| SYM |
| 5 | miRNA165 | SYS \| miRNA165 |
| 6 | PHB | !miRNA165 & (!CYA \| PYI52) |
| 7 | Auxin | Auxin |
| 8 | IAA5 | !Auxin & !CYA |
| 9 | WOX5 | PYI5 & !CYA & WOX5 |
| 10 | CLE | (CLE \| ! IAA5) & !SHR |
| 11 | ACR | CLE & Auxin |
| 12 | SYS | SHR & SCR |
| 13 | CYA | CLE & ACR |
| 14 | PYI5 | !PHB & !IAA5 |
| 15 | JYI | JKD & IAA5 |
| 16 | SYM | SYS & MGP |
| 17 | PYI52 | PHB & IAA5 |

TABLE S6. **Boolean functions for the 2017 RAM model in the BoolNet format.** The column with header 'Target gene name' contains the list of genes whose regulation is captured by the corresponding row entry in the column 'Regulatory logic rule'. The symbols &, | and ! correspond to the logic operators AND, OR and NOT respectively.

| Serial Number | Target gene name | Regulatory logic rule |
|---|---|---|
| 1 | CK | (PHB & !ARF) \| !SHR |
| 2 | ARR1 | !SCR & CK |
| 3 | SHY2 | ARR1 & !AUX |
| 4 | AUXIAA | !AUX |
| 5 | ARF | !AUXIAA |
| 6 | ARF10 | !(JKD & SHR) & !AUXIAA |
| 7 | ARF5 | !(SHR & MGP) & !SHY2 & !AUXIAA |
| 8 | AUX | WOX5 \| AUX |
| 9 | SCR | SHR & JKD & SCR |
| 10 | SHR | SHR \| (SCR & JKD) |
| 11 | MIR166 | (SHR & SCR & ! CK) \| !PHB |
| 12 | PHB | !MIR166 |
| 13 | JKD | !PHB & SHR & SCR |
| 14 | MGP | !WOX5 & SHR & SCR |
| 15 | WOX5 | !ARF10 & ARF5 & !CLE40 |
| 16 | CLE40 | !SHR |

TABLE S7. **Boolean functions for the 2020 RSCN model in the BoolNet format.** The column with header 'Target gene name' contains the list of genes whose regulation is captured by the corresponding row entry in the column 'Regulatory logic rule'. The symbols &, | and ! correspond to the logic operators AND, OR and NOT respectively.

| **Serial Number** | **Target gene name** | **Regulatory logic rule** |
|---|---|---|
| 1 | CK | (PHB & !ARF) \| !SHR |
| 2 | ARR1 | !SCR & CK |
| 3 | SHY2 | ARR1 & !AUX |
| 4 | AUXIAA | !AUX |
| 5 | ARF | !AUXIAA |
| 6 | ARF10 | !(JKD & SHR) & !AUXIAA |
| 7 | ARF5 | ((PHB \| PLT) & !(SHR & MGP)) & !SHY2 & !AUXIAA |
| 8 | XAL1 | ARF |
| 9 | PLT | ARF5 \| ARF \| WOX5 \| XAL1 |
| 10 | AUX | AUX |
| 11 | SCR | SHR & JKD & SCR |
| 12 | SHR | SHR \| (SCR & JKD) |
| 13 | MIR166 | (SCR & SHR & !ARR1) \| !PHB |
| 14 | PHB | ((!ARR1 & PLT) \| PHB) & !MIR166 |
| 15 | JKD | !PHB & SHR & SCR |
| 16 | MGP | !ARF5 & SHR & SCR & MGP |
| 17 | WOX5 | !ARF10 & ARF5 & !CLE40 & SCR & PLT |
| 18 | CLE40 | !SHR |

TABLE S8. **990 unique models that satisfy the expected landscape of the RSCN**. Provided below are the 990 Boolean models that are obtained using the iterative greedy algorithm described in section III F. Starting from 2020 *Arabidopsis thaliana* RSCN Boolean model (see SI Table S7) which does not satisfy the expected hierarchy, we implement our iterative greedy algorithm (Algorithm 1). Each row is a Boolean network model represented as a vector whose entries specify the Boolean functions assigned to the 18 genes of the network. The genes are ordered as CK, ARR1, SHY2, AUXIAA, ARF, ARF10, ARF5, XAL1, PLT, AUX, SCR, SHR, MIR166, PHB, JKD, MGP, WOX5, CLE40. Furthermore, each $k$-input BF is encoded as an integer in the following manner. Take the output column ($(k+1)^{th}$ column) of the truth table. Order the corresponding binary string so that the leftmost (most significant) bit comes from the first row (all input bits equal to 0) and the rightmost (least significant) bit comes from the last row (all input bits equal to 1). The integer equivalent of this binary string is the encoding of the BF. The inputs to each gene are taken in the following order in their truth tables (from left to right). CK: [PHB, ARF, SHR], ARR1: [SCR, CK], SHY2: [ARR1, AUX], AUXIAA: [AUX], ARF: [AUXIAA], ARF10: [JKD, SHR, AUXIAA], ARF5: [PHB, PLT, SHR, MGP, SHY2, AUXIAA], XAL1: [ARF], PLT: [ARF5, ARF, WOX5, XAL1], AUX: [AUX], SCR: [SHR, JKD, SCR], SHR: [SHR, SCR, JKD], MIR166: [SCR, SHR, ARR1, PHB], PHB: [ARR1, PLT, PHB, MIR166], JKD: [PHB, SHR, SCR], MGP: [ARF5, SHR, SCR, MGP], WOX5: [ARF10, ARF5, CLE40, SCR, PLT], CLE40: [SHR]

| Model No. | Boolean functions | | | | | | | | | | | | | | | | | |
|---|---|---|---|---|---|---|---|---|---|---|---|---|---|---|---|---|---|---|
| | CK | ARR1 | SHY2 | AUXIAA | ARF | ARF10 | ARF5 | XAL1 | PLT | AUX | SCR | SHR | MIR166 | PHB | JKD | MGP | WOX5 | CLE40 |
| 1 | 174 | 4 | 2 | 2 | 2 | 168 | 1152939152196298372 | 1 | 1807 | 1 | 1 | 31 | 43759 | 48043 | 16 | 5393 | 7536640 | 2 |
| 2 | 174 | 4 | 2 | 2 | 2 | 168 | 11529391657816465568 | 1 | 16255 | 1 | 1 | 31 | 43759 | 47931 | 81 | 4864 | 3342385 | 2 |
| 3 | 174 | 4 | 2 | 2 | 2 | 168 | 11529391657816465568 | 1 | 3871 | 1 | 1 | 31 | 43759 | 47931 | 16 | 4353 | 3211264 | 2 |
| 4 | 174 | 4 | 2 | 2 | 2 | 168 | 11529391657942296744 | 1 | 32767 | 1 | 1 | 31 | 43694 | 8963 | 16 | 13057 | 7340080 | 2 |
| 5 | 174 | 4 | 2 | 2 | 2 | 168 | 11529391659096534252 | 1 | 24447 | 1 | 1 | 31 | 43694 | 10786 | 16 | 4881 | 5308416 | 2 |
| 6 | 174 | 4 | 2 | 2 | 2 | 168 | 11529462164001710304 | 1 | 7999 | 1 | 1 | 31 | 43759 | 47931 | 81 | 4881 | 3211281 | 2 |
| 7 | 174 | 4 | 2 | 2 | 2 | 168 | 11529479963624144124 | 1 | 14143 | 1 | 1 | 31 | 43694 | 47931 | 16 | 4864 | 3342352 | 2 |
| 8 | 174 | 4 | 2 | 2 | 2 | 168 | 11565420317396478634 | 1 | 32767 | 1 | 1 | 31 | 43759 | 47931 | 81 | 769 | 7667712 | 2 |
| 9 | 174 | 4 | 2 | 2 | 2 | 168 | 11565420454837526688 | 1 | 3967 | 1 | 1 | 31 | 43759 | 13091 | 81 | 4864 | 7667712 | 2 |
| 10 | 174 | 4 | 2 | 2 | 2 | 168 | 11565420455003728554 | 1 | 32767 | 1 | 1 | 31 | 43694 | 8963 | 16 | 4881 | 3342352 | 2 |
| 11 | 174 | 4 | 2 | 2 | 2 | 168 | 11565508553372925900 | 1 | 8191 | 1 | 19 | 31 | 43694 | 48043 | 16 | 4864 | 1048576 | 2 |
| 12 | 174 | 4 | 2 | 2 | 2 | 168 | 11565508554806788095 | 1 | 3967 | 1 | 19 | 31 | 43759 | 47931 | 16 | 769 | 16711696 | 2 |
| 13 | 174 | 4 | 2 | 2 | 2 | 168 | 11565508759531356159 | 1 | 8031 | 1 | 1 | 31 | 43694 | 47931 | 16 | 769 | 16187392 | 2 |
| 14 | 174 | 4 | 2 | 2 | 2 | 168 | 11574427654092269728 | 1 | 24447 | 1 | 1 | 31 | 43759 | 13091 | 16 | 4881 | 7536691 | 2 |
| 15 | 174 | 4 | 2 | 2 | 2 | 168 | 11574427654092284064 | 1 | 21887 | 1 | 1 | 31 | 43759 | 13091 | 81 | 769 | 7667712 | 2 |
| 16 | 174 | 4 | 2 | 2 | 2 | 168 | 11574427655166029984 | 1 | 24447 | 1 | 1 | 31 | 43759 | 10786 | 81 | 769 | 1114128 | 2 |
| 17 | 174 | 4 | 2 | 2 | 2 | 168 | 11574427655367360511 | 1 | 14143 | 1 | 1 | 31 | 43759 | 47931 | 81 | 769 | 5242896 | 2 |
| 18 | 174 | 4 | 2 | 2 | 2 | 168 | 11574427655686127596 | 1 | 3967 | 1 | 1 | 31 | 43759 | 47931 | 16 | 4881 | 16711729 | 2 |
| 19 | 174 | 4 | 2 | 2 | 2 | 168 | 11574498022836461792 | 1 | 5631 | 1 | 1 | 31 | 43694 | 15155 | 16 | 769 | 7340032 | 2 |
| 20 | 174 | 4 | 2 | 2 | 2 | 168 | 11574511216975998188 | 1 | 3871 | 1 | 1 | 31 | 43759 | 13091 | 81 | 13057 | 3211281 | 2 |
| 21 | 174 | 4 | 2 | 2 | 2 | 168 | 11574511218256047340 | 1 | 21887 | 1 | 1 | 31 | 43694 | 13091 | 81 | 4881 | 15728656 | 2 |
| 22 | 174 | 4 | 2 | 2 | 2 | 168 | 11574515615157255336 | 1 | 21887 | 1 | 1 | 31 | 43694 | 47931 | 16 | 13057 | 3342385 | 2 |
| 23 | 174 | 4 | 2 | 2 | 2 | 168 | 11574515960218714108 | 1 | 8191 | 1 | 1 | 31 | 43759 | 48043 | 81 | 4881 | 3211312 | 2 |
| 24 | 174 | 4 | 2 | 2 | 2 | 168 | 11574532108971999136 | 1 | 3871 | 1 | 1 | 31 | 43694 | 47931 | 81 | 769 | 15728656 | 2 |
| 25 | 174 | 4 | 2 | 2 | 2 | 168 | 12150989906662916095 | 1 | 8191 | 1 | 1 | 31 | 43694 | 48043 | 16 | 65281 | 16187443 | 2 |
| 26 | 174 | 4 | 2 | 2 | 2 | 168 | 12249978457458540480 | 1 | 3871 | 1 | 1 | 31 | 43694 | 15155 | 16 | 4353 | 3342352 | 2 |
| 27 | 174 | 4 | 2 | 2 | 2 | 168 | 12249978593480321704 | 1 | 3871 | 1 | 1 | 31 | 43694 | 8706 | 81 | 769 | 15794176 | 2 |
| 28 | 174 | 4 | 2 | 2 | 2 | 168 | 12250049099654560496 | 1 | 5461 | 1 | 1 | 31 | 43694 | 15155 | 81 | 4881 | 16187392 | 2 |
| 29 | 174 | 4 | 2 | 2 | 2 | 168 | 12250049101080625088 | 1 | 1287 | 1 | 1 | 31 | 43694 | 8994 | 81 | 4353 | 1048576 | 2 |
| 30 | 174 | 4 | 2 | 2 | 2 | 168 | 12286007391929040880 | 1 | 32767 | 1 | 1 | 31 | 43759 | 43522 | 81 | 13057 | 3211281 | 2 |
| 31 | 174 | 4 | 2 | 2 | 2 | 168 | 12295014589755599528 | 1 | 8191 | 1 | 1 | 31 | 43694 | 15155 | 16 | 13057 | 7340032 | 2 |
| 32 | 174 | 4 | 2 | 2 | 2 | 168 | 12295014589755616992 | 1 | 1807 | 1 | 1 | 31 | 43694 | 10786 | 16 | 769 | 1114128 | 2 |
| 33 | 174 | 4 | 2 | 2 | 2 | 168 | 13835269714693115016 | 1 | 14335 | 1 | 1 | 31 | 43759 | 2819 | 16 | 4864 | 16711696 | 2 |
| 34 | 174 | 4 | 2 | 2 | 2 | 168 | 13835269715565477887 | 1 | 3871 | 1 | 1 | 31 | 43759 | 47931 | 81 | 769 | 7667712 | 2 |

| Model No. | Functions in integer form at the nodes of the network | | | | | | | | | | | | | | | | | |
|---|---|---|---|---|---|---|---|---|---|---|---|---|---|---|---|---|---|---|
| | CK | ARR1 | SHY2 | AUXIAA | ARF | ARF10 | ARF5 | XAL1 | PLT | AUX | SCR | SHR | MIR166 | PHB | JKD | MGP | WOX5 | CLE40 |
| 35 | 174 | 4 | 2 | 2 | 2 | 168 | 1383530503710814 6336 | 1 | 1807 | 1 | 1 | 31 | 43759 | 48043 | 81 | 4864 | 5570576 | 2 |
| 36 | 174 | 4 | 2 | 2 | 2 | 168 | 13835305037108146336 | 1 | 5461 | 1 | 1 | 31 | 43694 | 48043 | 16 | 4353 | 7536640 | 2 |
| 37 | 174 | 4 | 2 | 2 | 2 | 168 | 13835322972220488442 | 1 | 22527 | 1 | 1 | 31 | 43694 | 15155 | 81 | 4881 | 1048576 | 2 |
| 38 | 174 | 4 | 2 | 2 | 2 | 168 | 13835322972633624816 | 1 | 8031 | 1 | 1 | 31 | 43694 | 48043 | 81 | 4881 | 3342352 | 2 |
| 39 | 174 | 4 | 2 | 2 | 2 | 168 | 13871298511519150320 | 1 | 2047 | 1 | 1 | 31 | 43759 | 10786 | 16 | 4881 | 3211264 | 2 |
| 40 | 174 | 4 | 2 | 2 | 2 | 168 | 13871298511519150320 | 1 | 22527 | 1 | 1 | 31 | 43759 | 10786 | 81 | 4881 | 7340032 | 2 |
| 41 | 174 | 4 | 2 | 2 | 2 | 168 | 13871298511519154090 | 1 | 8031 | 1 | 1 | 31 | 43759 | 10786 | 16 | 4881 | 1114128 | 2 |
| 42 | 174 | 4 | 2 | 2 | 2 | 168 | 13871298786603355340 | 1 | 2047 | 1 | 1 | 31 | 43759 | 10786 | 16 | 13057 | 16187392 | 2 |
| 43 | 174 | 4 | 2 | 2 | 2 | 168 | 13871351769117815024 | 1 | 14199 | 1 | 1 | 31 | 43759 | 10786 | 16 | 769 | 15728656 | 2 |
| 44 | 174 | 4 | 2 | 2 | 2 | 168 | 13871351769117815024 | 1 | 5631 | 1 | 1 | 31 | 43759 | 10786 | 81 | 769 | 3342352 | 2 |
| 45 | 174 | 4 | 2 | 2 | 2 | 168 | 13871351769315475455 | 1 | 32767 | 1 | 1 | 31 | 43759 | 2562 | 16 | 769 | 3342352 | 2 |
| 46 | 174 | 4 | 2 | 2 | 2 | 168 | 13889313185112575180 | 1 | 30591 | 1 | 1 | 31 | 43759 | 10786 | 16 | 4881 | 3342352 | 2 |
| 47 | 174 | 4 | 2 | 2 | 2 | 168 | 13889313185615380479 | 1 | 14143 | 1 | 1 | 31 | 43694 | 47931 | 16 | 769 | 3342352 | 2 |
| 48 | 174 | 4 | 2 | 2 | 2 | 168 | 13889313185967701994 | 1 | 32767 | 1 | 1 | 31 | 43759 | 8706 | 16 | 13057 | 1114128 | 2 |
| 49 | 174 | 4 | 2 | 2 | 2 | 168 | 13889359364399098602 | 1 | 7999 | 1 | 1 | 31 | 43759 | 15155 | 16 | 4881 | 3211312 | 2 |
| 50 | 174 | 4 | 2 | 2 | 2 | 168 | 13889359365456068544 | 1 | 30591 | 1 | 1 | 31 | 43759 | 8706 | 16 | 769 | 1048576 | 2 |
| 51 | 174 | 4 | 2 | 2 | 2 | 168 | 13889365961468866784 | 1 | 8191 | 1 | 1 | 31 | 43759 | 15155 | 16 | 769 | 7536691 | 2 |
| 52 | 174 | 4 | 2 | 2 | 2 | 168 | 13889366099716272368 | 1 | 8031 | 1 | 1 | 31 | 43694 | 10786 | 16 | 13057 | 7536640 | 2 |
| 53 | 174 | 4 | 2 | 2 | 2 | 168 | 13889366168164167920 | 1 | 1287 | 1 | 1 | 31 | 43759 | 10786 | 16 | 769 | 15728656 | 2 |
| 54 | 174 | 4 | 2 | 2 | 2 | 168 | 13889366168331942640 | 1 | 22367 | 1 | 1 | 31 | 43759 | 8994 | 81 | 4881 | 7667712 | 2 |
| 55 | 174 | 4 | 2 | 2 | 2 | 168 | 141287244210304 | 1 | 21887 | 1 | 1 | 31 | 43759 | 8706 | 16 | 769 | 7340032 | 2 |
| 56 | 174 | 4 | 2 | 2 | 2 | 168 | 141287244210304 | 1 | 32767 | 1 | 1 | 31 | 43694 | 48043 | 81 | 4864 | 5570576 | 2 |
| 57 | 174 | 4 | 2 | 2 | 2 | 168 | 141287244212864 | 1 | 32767 | 1 | 1 | 31 | 43759 | 10786 | 81 | 4881 | 1048576 | 2 |
| 58 | 174 | 4 | 2 | 2 | 2 | 168 | 141290096339584 | 1 | 22367 | 1 | 1 | 31 | 43694 | 47931 | 16 | 13057 | 3211312 | 2 |
| 59 | 174 | 4 | 2 | 2 | 2 | 168 | 141290096339584 | 1 | 24447 | 1 | 1 | 31 | 43694 | 8994 | 16 | 4881 | 3145744 | 2 |
| 60 | 174 | 4 | 2 | 2 | 2 | 168 | 141290465443968 | 1 | 1287 | 1 | 1 | 31 | 43694 | 10786 | 16 | 13057 | 3211312 | 2 |
| 61 | 174 | 4 | 2 | 2 | 2 | 168 | 14465782733441399536 | 1 | 8031 | 1 | 1 | 31 | 43759 | 10786 | 81 | 4881 | 3211281 | 2 |
| 62 | 174 | 4 | 2 | 2 | 2 | 168 | 14465782733441400831 | 1 | 3871 | 1 | 1 | 31 | 43759 | 15155 | 16 | 769 | 15794176 | 2 |
| 63 | 174 | 4 | 2 | 2 | 2 | 168 | 14465837915181742842 | 1 | 30591 | 1 | 1 | 31 | 43694 | 13091 | 81 | 769 | 7536640 | 2 |
| 64 | 174 | 4 | 2 | 2 | 2 | 168 | 14465837915181742842 | 1 | 7999 | 1 | 1 | 31 | 43694 | 8994 | 81 | 4881 | 16187392 | 2 |
| 65 | 174 | 4 | 2 | 2 | 2 | 168 | 14699974037287783628 | 1 | 22527 | 1 | 1 | 31 | 43694 | 47931 | 81 | 4881 | 16711729 | 2 |
| 66 | 174 | 4 | 2 | 2 | 2 | 168 | 14699974037300366528 | 1 | 2047 | 1 | 1 | 31 | 43694 | 8994 | 81 | 13057 | 5242896 | 2 |
| 67 | 174 | 4 | 2 | 2 | 2 | 168 | 14699974037300366528 | 1 | 24447 | 1 | 1 | 31 | 43759 | 15155 | 16 | 4864 | 1048576 | 2 |
| 68 | 174 | 4 | 2 | 2 | 2 | 168 | 14699974312174603464 | 1 | 3967 | 1 | 1 | 31 | 43759 | 15155 | 81 | 769 | 16711729 | 2 |
| 69 | 174 | 4 | 2 | 2 | 2 | 168 | 14700009359098838256 | 1 | 7999 | 1 | 1 | 31 | 43694 | 10786 | 16 | 4864 | 7340032 | 2 |
| 70 | 174 | 4 | 2 | 2 | 2 | 168 | 14700009359954476960 | 1 | 3967 | 1 | 1 | 31 | 43759 | 13091 | 81 | 13057 | 5570576 | 2 |
| 71 | 174 | 4 | 2 | 2 | 2 | 168 | 14736003109197237440 | 1 | 24447 | 1 | 1 | 31 | 43759 | 8963 | 16 | 769 | 3342385 | 2 |
| 72 | 174 | 4 | 2 | 2 | 2 | 168 | 14736059047704854432 | 1 | 22367 | 1 | 1 | 31 | 43759 | 47931 | 16 | 13057 | 3211264 | 2 |
| 73 | 174 | 4 | 2 | 2 | 2 | 168 | 14754017507706719432 | 1 | 8031 | 1 | 19 | 31 | 43694 | 48043 | 16 | 1792 | 5308416 | 2 |
| 74 | 174 | 4 | 2 | 2 | 2 | 168 | 14754055029113548512 | 1 | 24447 | 1 | 1 | 31 | 43759 | 48043 | 16 | 4881 | 7536640 | 2 |
| 75 | 174 | 4 | 2 | 2 | 2 | 168 | 14754073789531357178 | 1 | 2047 | 1 | 1 | 31 | 43759 | 47931 | 81 | 769 | 3145744 | 2 |
| 76 | 174 | 4 | 2 | 2 | 2 | 168 | 14754073789531357178 | 1 | 5631 | 1 | 1 | 31 | 43694 | 47931 | 16 | 13057 | 15728656 | 2 |
| 77 | 174 | 4 | 2 | 2 | 2 | 168 | 150085627316352 | 1 | 1301 | 1 | 1 | 31 | 43694 | 10786 | 81 | 769 | 16711729 | 2 |
| 78 | 174 | 4 | 2 | 2 | 2 | 168 | 150085627316352 | 1 | 14143 | 1 | 1 | 31 | 43694 | 10786 | 16 | 13057 | 1048576 | 2 |
| 79 | 174 | 4 | 2 | 2 | 2 | 168 | 150085627316352 | 1 | 14143 | 1 | 1 | 31 | 43694 | 10786 | 81 | 4864 | 1048576 | 2 |

| Model No. | Functions in integer form at the nodes of the network | | | | | | | | | | | | | | | | | |
|---|---|---|---|---|---|---|---|---|---|---|---|---|---|---|---|---|---|---|
| | CK | ARR1 | SHY2 | AUXIAA | ARF | ARF10 | ARF5 | XAL1 | PLT | AUX | SCR | SHR | MIR166 | PHB | JKD | MGP | WOX5 | CLE40 |
| 80 | 174 | 4 | 2 | 2 | 2 | 168 | 150085627316352 | 1 | 14143 | 1 | 1 | 31 | 43759 | 10786 | 16 | 13057 | 1048576 | 2 |
| 81 | 174 | 4 | 2 | 2 | 2 | 168 | 150085627316352 | 1 | 14335 | 1 | 1 | 31 | 43759 | 10786 | 16 | 4881 | 7667712 | 2 |
| 82 | 174 | 4 | 2 | 2 | 2 | 168 | 150085627316352 | 1 | 16255 | 1 | 1 | 31 | 43694 | 10786 | 81 | 13057 | 1048576 | 2 |
| 83 | 174 | 4 | 2 | 2 | 2 | 168 | 150085627316352 | 1 | 16255 | 1 | 1 | 31 | 43694 | 10786 | 81 | 4881 | 1048576 | 2 |
| 84 | 174 | 4 | 2 | 2 | 2 | 168 | 150085627316352 | 1 | 16255 | 1 | 1 | 31 | 43759 | 10786 | 16 | 13057 | 1048576 | 2 |
| 85 | 174 | 4 | 2 | 2 | 2 | 168 | 150085627316352 | 1 | 16255 | 1 | 1 | 31 | 43759 | 15155 | 16 | 13057 | 1048576 | 2 |
| 86 | 174 | 4 | 2 | 2 | 2 | 168 | 150085627316352 | 1 | 16255 | 1 | 1 | 31 | 43759 | 48043 | 81 | 4353 | 1048576 | 2 |
| 87 | 174 | 4 | 2 | 2 | 2 | 168 | 150085627316352 | 1 | 16255 | 1 | 19 | 31 | 43759 | 48043 | 16 | 769 | 5242896 | 2 |
| 88 | 174 | 4 | 2 | 2 | 2 | 168 | 150085627316352 | 1 | 1807 | 1 | 1 | 31 | 43694 | 10786 | 81 | 13057 | 1048576 | 2 |
| 89 | 174 | 4 | 2 | 2 | 2 | 168 | 150085627316352 | 1 | 1807 | 1 | 1 | 31 | 43759 | 10786 | 16 | 4881 | 1048576 | 2 |
| 90 | 174 | 4 | 2 | 2 | 2 | 168 | 150085627316352 | 1 | 2047 | 1 | 1 | 31 | 43759 | 10786 | 16 | 769 | 1048576 | 2 |
| 91 | 174 | 4 | 2 | 2 | 2 | 168 | 150085627316352 | 1 | 22367 | 1 | 1 | 31 | 43759 | 8963 | 16 | 769 | 1048576 | 2 |
| 92 | 174 | 4 | 2 | 2 | 2 | 168 | 150085627316352 | 1 | 22527 | 1 | 1 | 31 | 43759 | 10786 | 16 | 4864 | 1048576 | 2 |
| 93 | 174 | 4 | 2 | 2 | 2 | 168 | 150085627316352 | 1 | 22527 | 1 | 1 | 31 | 43759 | 10786 | 81 | 769 | 1048576 | 2 |
| 94 | 174 | 4 | 2 | 2 | 2 | 168 | 150085627316352 | 1 | 22527 | 1 | 1 | 31 | 43759 | 10786 | 81 | 769 | 7536691 | 2 |
| 95 | 174 | 4 | 2 | 2 | 2 | 168 | 150085627316352 | 1 | 24447 | 1 | 1 | 31 | 43694 | 10786 | 16 | 4881 | 1048576 | 2 |
| 96 | 174 | 4 | 2 | 2 | 2 | 168 | 150085627316352 | 1 | 24447 | 1 | 1 | 31 | 43759 | 10786 | 16 | 4881 | 1048576 | 2 |
| 97 | 174 | 4 | 2 | 2 | 2 | 168 | 150085627316352 | 1 | 24447 | 1 | 1 | 31 | 43759 | 47931 | 16 | 769 | 3145744 | 2 |
| 98 | 174 | 4 | 2 | 2 | 2 | 168 | 150085627316352 | 1 | 24447 | 1 | 1 | 31 | 43759 | 48043 | 16 | 769 | 3145744 | 2 |
| 99 | 174 | 4 | 2 | 2 | 2 | 168 | 150085627316352 | 1 | 30591 | 1 | 1 | 31 | 43759 | 10786 | 16 | 4864 | 16187392 | 2 |
| 100 | 174 | 4 | 2 | 2 | 2 | 168 | 150085627316352 | 1 | 32767 | 1 | 1 | 31 | 43694 | 10786 | 16 | 13057 | 1048576 | 2 |
| 101 | 174 | 4 | 2 | 2 | 2 | 168 | 150085627316352 | 1 | 32767 | 1 | 1 | 31 | 43694 | 10786 | 16 | 4881 | 1048576 | 2 |
| 102 | 174 | 4 | 2 | 2 | 2 | 168 | 150085627316352 | 1 | 32767 | 1 | 1 | 31 | 43694 | 10786 | 16 | 769 | 1048576 | 2 |
| 103 | 174 | 4 | 2 | 2 | 2 | 168 | 150085627316352 | 1 | 32767 | 1 | 1 | 31 | 43694 | 10786 | 81 | 4881 | 1048576 | 2 |
| 104 | 174 | 4 | 2 | 2 | 2 | 168 | 150085627316352 | 1 | 32767 | 1 | 1 | 31 | 43694 | 13091 | 81 | 769 | 3342385 | 2 |
| 105 | 174 | 4 | 2 | 2 | 2 | 168 | 150085627316352 | 1 | 32767 | 1 | 1 | 31 | 43694 | 47931 | 16 | 769 | 1048576 | 2 |
| 106 | 174 | 4 | 2 | 2 | 2 | 168 | 150085627316352 | 1 | 32767 | 1 | 1 | 31 | 43694 | 8706 | 16 | 13057 | 1048576 | 2 |
| 107 | 174 | 4 | 2 | 2 | 2 | 168 | 150085627316352 | 1 | 32767 | 1 | 1 | 31 | 43759 | 10786 | 16 | 13057 | 1048576 | 2 |
| 108 | 174 | 4 | 2 | 2 | 2 | 168 | 150085627316352 | 1 | 32767 | 1 | 1 | 31 | 43759 | 10786 | 16 | 4864 | 1048576 | 2 |
| 109 | 174 | 4 | 2 | 2 | 2 | 168 | 150085627316352 | 1 | 32767 | 1 | 1 | 31 | 43759 | 10786 | 81 | 4881 | 1048576 | 2 |
| 110 | 174 | 4 | 2 | 2 | 2 | 168 | 150085627316352 | 1 | 32767 | 1 | 1 | 31 | 43759 | 15155 | 81 | 13057 | 1048576 | 2 |
| 111 | 174 | 4 | 2 | 2 | 2 | 168 | 150085627316352 | 1 | 32767 | 1 | 1 | 31 | 43759 | 47931 | 16 | 13057 | 1048576 | 2 |
| 112 | 174 | 4 | 2 | 2 | 2 | 168 | 150085627316352 | 1 | 32767 | 1 | 1 | 31 | 43759 | 48043 | 81 | 13057 | 1048576 | 2 |
| 113 | 174 | 4 | 2 | 2 | 2 | 168 | 150085627316352 | 1 | 3871 | 1 | 1 | 31 | 43759 | 10786 | 16 | 4353 | 1048576 | 2 |
| 114 | 174 | 4 | 2 | 2 | 2 | 168 | 150085627316352 | 1 | 3871 | 1 | 1 | 31 | 43759 | 10786 | 81 | 4881 | 1048576 | 2 |
| 115 | 174 | 4 | 2 | 2 | 2 | 168 | 150085627316352 | 1 | 3871 | 1 | 1 | 31 | 43759 | 10786 | 81 | 769 | 1048576 | 2 |
| 116 | 174 | 4 | 2 | 2 | 2 | 168 | 150085627316352 | 1 | 3967 | 1 | 1 | 31 | 43694 | 48043 | 16 | 13057 | 3211264 | 2 |
| 117 | 174 | 4 | 2 | 2 | 2 | 168 | 150085627316352 | 1 | 3967 | 1 | 1 | 31 | 43759 | 10786 | 81 | 13057 | 1048576 | 2 |
| 118 | 174 | 4 | 2 | 2 | 2 | 168 | 150085627316352 | 1 | 3967 | 1 | 1 | 31 | 43759 | 10786 | 81 | 13057 | 7667712 | 2 |
| 119 | 174 | 4 | 2 | 2 | 2 | 168 | 150085627316352 | 1 | 3967 | 1 | 1 | 31 | 43759 | 13091 | 81 | 4881 | 3342352 | 2 |
| 120 | 174 | 4 | 2 | 2 | 2 | 168 | 150085627316352 | 1 | 5631 | 1 | 1 | 31 | 43694 | 10786 | 16 | 4353 | 1048576 | 2 |
| 121 | 174 | 4 | 2 | 2 | 2 | 168 | 150085627316352 | 1 | 5631 | 1 | 1 | 31 | 43759 | 10786 | 16 | 4864 | 1048576 | 2 |
| 122 | 174 | 4 | 2 | 2 | 2 | 168 | 150085627316352 | 1 | 5631 | 1 | 1 | 31 | 43759 | 10786 | 81 | 13057 | 1048576 | 2 |
| 123 | 174 | 4 | 2 | 2 | 2 | 168 | 150085627316352 | 1 | 7999 | 1 | 1 | 31 | 43694 | 10786 | 16 | 4881 | 1048576 | 2 |
| 124 | 174 | 4 | 2 | 2 | 2 | 168 | 150085627316352 | 1 | 7999 | 1 | 1 | 31 | 43759 | 10786 | 81 | 4864 | 1048576 | 2 |

| Model No. | Functions in integer form at the nodes of the network | | | | | | | | | | | | | | | | | |
|---|---|---|---|---|---|---|---|---|---|---|---|---|---|---|---|---|---|---|
| | CK | ARR1 | SHY2 | AUXIAA | ARF | ARF10 | ARF5 | XAL1 | PLT | AUX | SCR | SHR | MIR166 | PHB | JKD | MGP | WOX5 | CLE40 |
| 125 | 174 | 4 | 2 | 2 | 2 | 168 | 150085627316352 | 1 | 7999 | 1 | 1 | 31 | 43759 | 48043 | 81 | 4353 | 1048576 | 2 |
| 126 | 174 | 4 | 2 | 2 | 2 | 168 | 150085627316352 | 1 | 8031 | 1 | 1 | 31 | 43694 | 48043 | 81 | 769 | 1048576 | 2 |
| 127 | 174 | 4 | 2 | 2 | 2 | 168 | 150085627316352 | 1 | 8031 | 1 | 1 | 31 | 43759 | 10786 | 81 | 769 | 1048576 | 2 |
| 128 | 174 | 4 | 2 | 2 | 2 | 168 | 150085627316352 | 1 | 8191 | 1 | 1 | 31 | 43694 | 10786 | 81 | 13057 | 1048576 | 2 |
| 129 | 174 | 4 | 2 | 2 | 2 | 168 | 150085627316352 | 1 | 8191 | 1 | 1 | 31 | 43759 | 10786 | 81 | 13057 | 1048576 | 2 |
| 130 | 174 | 4 | 2 | 2 | 2 | 168 | 150085627840648 | 1 | 14335 | 1 | 19 | 31 | 43759 | 48043 | 16 | 5393 | 3211264 | 2 |
| 131 | 174 | 4 | 2 | 2 | 2 | 168 | 150086199847584 | 1 | 22527 | 1 | 1 | 31 | 43759 | 13091 | 16 | 4864 | 15794176 | 2 |
| 132 | 174 | 4 | 2 | 2 | 2 | 168 | 16141165981851513072 | 1 | 5631 | 1 | 1 | 31 | 43759 | 10786 | 16 | 13057 | 5570576 | 2 |
| 133 | 174 | 4 | 2 | 2 | 2 | 168 | 16141165982120997104 | 1 | 8031 | 1 | 1 | 31 | 43694 | 8706 | 16 | 769 | 3211281 | 2 |
| 134 | 174 | 4 | 2 | 2 | 2 | 168 | 16186184042337591520 | 1 | 3871 | 1 | 1 | 31 | 43759 | 13091 | 81 | 4881 | 1114128 | 2 |
| 135 | 174 | 4 | 2 | 2 | 2 | 168 | 16186184317483938032 | 1 | 30591 | 1 | 1 | 31 | 43694 | 10786 | 81 | 4881 | 3145744 | 2 |
| 136 | 174 | 4 | 2 | 2 | 2 | 168 | 16186201634527834352 | 1 | 8191 | 1 | 1 | 31 | 43694 | 47931 | 81 | 769 | 5308433 | 2 |
| 137 | 174 | 4 | 2 | 2 | 2 | 168 | 16186201634792075424 | 1 | 32767 | 1 | 1 | 31 | 43694 | 47931 | 16 | 4881 | 3342352 | 2 |
| 138 | 174 | 4 | 2 | 2 | 2 | 168 | 16195191379033386700 | 1 | 32767 | 1 | 1 | 31 | 43694 | 13091 | 16 | 14080 | 1048576 | 2 |
| 139 | 174 | 4 | 2 | 2 | 2 | 168 | 16195191516708007662 | 1 | 32767 | 1 | 1 | 31 | 43759 | 8994 | 81 | 4881 | 15728656 | 2 |
| 140 | 174 | 4 | 2 | 2 | 2 | 168 | 16195208971455102924 | 1 | 5461 | 1 | 1 | 31 | 43759 | 10786 | 81 | 4881 | 5308416 | 2 |
| 141 | 174 | 4 | 2 | 2 | 2 | 168 | 16204214109414096895 | 1 | 3871 | 1 | 1 | 31 | 43694 | 10786 | 81 | 13057 | 5308416 | 2 |
| 142 | 174 | 4 | 2 | 2 | 2 | 168 | 16204216308183658720 | 1 | 7999 | 1 | 1 | 31 | 43694 | 15155 | 81 | 4881 | 5242896 | 2 |
| 143 | 174 | 4 | 2 | 2 | 2 | 168 | 16915778314696719088 | 1 | 3967 | 1 | 1 | 31 | 43759 | 10786 | 81 | 4881 | 3342385 | 2 |
| 144 | 174 | 4 | 2 | 2 | 2 | 168 | 17050888365133594623 | 1 | 3871 | 1 | 1 | 31 | 43694 | 10786 | 16 | 769 | 3211281 | 2 |
| 145 | 174 | 4 | 2 | 2 | 2 | 168 | 17149970030875569902 | 1 | 3871 | 1 | 1 | 31 | 43759 | 13091 | 16 | 4864 | 5242896 | 2 |
| 146 | 174 | 4 | 2 | 2 | 2 | 168 | 17149970031161901055 | 1 | 2047 | 1 | 19 | 31 | 43759 | 48043 | 16 | 4881 | 16187392 | 2 |
| 147 | 174 | 4 | 2 | 2 | 2 | 168 | 17149987691765497855 | 1 | 21887 | 1 | 1 | 31 | 43759 | 15155 | 16 | 13057 | 5570576 | 2 |
| 148 | 174 | 4 | 2 | 2 | 2 | 168 | 17149987692049661680 | 1 | 3871 | 1 | 1 | 31 | 43694 | 8994 | 81 | 4864 | 5570576 | 2 |
| 149 | 174 | 4 | 2 | 2 | 2 | 168 | 17294087005686263968 | 1 | 24447 | 1 | 1 | 31 | 43694 | 10786 | 16 | 13057 | 1114128 | 2 |
| 150 | 174 | 4 | 2 | 2 | 2 | 168 | 17294087005686263968 | 1 | 5631 | 1 | 1 | 31 | 43694 | 15155 | 81 | 4864 | 7667712 | 2 |
| 151 | 174 | 4 | 2 | 2 | 2 | 168 | 17294087280562073792 | 1 | 2047 | 1 | 1 | 31 | 43759 | 10786 | 81 | 4864 | 7340080 | 2 |
| 152 | 174 | 4 | 2 | 2 | 2 | 168 | 17294087418244300782 | 1 | 5631 | 1 | 1 | 31 | 43759 | 10786 | 16 | 769 | 7667712 | 2 |
| 153 | 174 | 4 | 2 | 2 | 2 | 168 | 17294087486846335224 | 1 | 24447 | 1 | 1 | 31 | 43694 | 48043 | 16 | 22272 | 3211312 | 2 |
| 154 | 174 | 4 | 2 | 2 | 2 | 168 | 17330115802703130816 | 1 | 5631 | 1 | 1 | 31 | 43694 | 2562 | 16 | 4881 | 15728656 | 2 |
| 155 | 174 | 4 | 2 | 2 | 2 | 168 | 17330116077588377840 | 1 | 5631 | 1 | 1 | 31 | 43759 | 48043 | 81 | 769 | 16187443 | 2 |
| 156 | 174 | 4 | 2 | 2 | 2 | 168 | 17330116077833224140 | 1 | 16255 | 1 | 1 | 31 | 43759 | 47931 | 16 | 769 | 16711696 | 2 |
| 157 | 174 | 4 | 2 | 2 | 2 | 168 | 17348130613789458431 | 1 | 1807 | 1 | 1 | 31 | 43759 | 47931 | 81 | 769 | 5242896 | 2 |
| 158 | 174 | 4 | 2 | 2 | 2 | 168 | 17348130682423671546 | 1 | 2047 | 1 | 1 | 31 | 43694 | 10786 | 81 | 4881 | 5308433 | 2 |
| 159 | 174 | 4 | 2 | 2 | 2 | 168 | 17348130682423671546 | 1 | 22527 | 1 | 1 | 31 | 43694 | 47931 | 16 | 769 | 3211264 | 2 |
| 160 | 174 | 4 | 2 | 2 | 2 | 168 | 17357137812790509567 | 1 | 24447 | 1 | 1 | 31 | 43694 | 8963 | 16 | 769 | 16711729 | 2 |
| 161 | 174 | 4 | 2 | 2 | 2 | 168 | 17357137812790509567 | 1 | 32767 | 1 | 1 | 31 | 43759 | 13091 | 16 | 4881 | 3211281 | 2 |
| 162 | 174 | 4 | 2 | 2 | 2 | 168 | 17361641481272621808 | 1 | 8191 | 1 | 1 | 31 | 43694 | 13058 | 81 | 4881 | 15728656 | 2 |
| 163 | 174 | 4 | 2 | 2 | 2 | 168 | 17361641481306700538 | 1 | 24447 | 1 | 1 | 31 | 43694 | 47931 | 16 | 3841 | 3211312 | 2 |
| 164 | 174 | 4 | 2 | 2 | 2 | 168 | 17361641481373286142 | 1 | 32767 | 1 | 1 | 31 | 43759 | 43522 | 16 | 13057 | 7667712 | 2 |
| 165 | 174 | 4 | 2 | 2 | 2 | 168 | 17361654675277938426 | 1 | 3871 | 1 | 1 | 31 | 43694 | 47931 | 16 | 65281 | 7536640 | 2 |
| 166 | 174 | 4 | 2 | 2 | 2 | 168 | 176471616319728 | 1 | 22367 | 1 | 1 | 31 | 43694 | 8706 | 16 | 4881 | 16187392 | 2 |
| 167 | 174 | 4 | 2 | 2 | 2 | 168 | 176475655434432 | 1 | 21887 | 1 | 1 | 31 | 43694 | 48043 | 81 | 4864 | 1048576 | 2 |
| 168 | 174 | 4 | 2 | 2 | 2 | 168 | 176609055254688 | 1 | 5631 | 1 | 1 | 31 | 43759 | 10786 | 81 | 4881 | 1114128 | 2 |
| 169 | 174 | 4 | 2 | 2 | 2 | 168 | 17938115427834003455 | 1 | 8191 | 1 | 1 | 31 | 43694 | 13091 | 16 | 769 | 1114128 | 2 |

| Model No. | Functions in integer form at the nodes of the network | | | | | | | | | | | | | | | | | |
|---|---|---|---|---|---|---|---|---|---|---|---|---|---|---|---|---|---|---|
| | CK | ARR1 | SHY2 | AUXIAA | ARF | ARF10 | ARF5 | XAL1 | PLT | AUX | SCR | SHR | MIR166 | PHB | JKD | MGP | WOX5 | CLE40 |
| 170 | 174 | 4 | 2 | 2 | 2 | 168 | 18082228416801734384 | 1 | 24447 | 1 | 1 | 31 | 43759 | 13091 | 81 | 13057 | 7667712 | 2 |
| 171 | 174 | 4 | 2 | 2 | 2 | 168 | 18082228416802257658 | 1 | 22527 | 1 | 1 | 31 | 43694 | 8994 | 81 | 769 | 15794176 | 2 |
| 172 | 174 | 4 | 2 | 2 | 2 | 168 | 18082232814848245744 | 1 | 1807 | 1 | 1 | 31 | 43694 | 10786 | 16 | 4881 | 7536640 | 2 |
| 173 | 174 | 4 | 2 | 2 | 2 | 168 | 18082232814915354352 | 1 | 14143 | 1 | 1 | 31 | 43694 | 15155 | 16 | 13057 | 5570576 | 2 |
| 174 | 174 | 4 | 2 | 2 | 2 | 168 | 18374967409170776012 | 1 | 8191 | 1 | 1 | 31 | 43759 | 48043 | 81 | 5393 | 15728656 | 2 |
| 175 | 174 | 4 | 2 | 2 | 2 | 168 | 18374967409170776063 | 1 | 22367 | 1 | 1 | 31 | 43759 | 47931 | 16 | 5393 | 16187392 | 2 |
| 176 | 174 | 4 | 2 | 2 | 2 | 168 | 18374967409179688840 | 1 | 2047 | 1 | 1 | 31 | 43759 | 13058 | 16 | 769 | 15728656 | 2 |
| 177 | 174 | 4 | 2 | 2 | 2 | 168 | 18374967409181261728 | 1 | 21847 | 1 | 1 | 31 | 43759 | 2819 | 81 | 13057 | 16187443 | 2 |
| 178 | 174 | 4 | 2 | 2 | 2 | 168 | 18374967409184145356 | 1 | 7999 | 1 | 1 | 31 | 43759 | 10786 | 81 | 4864 | 5308416 | 2 |
| 179 | 174 | 4 | 2 | 2 | 2 | 168 | 18374967684048682952 | 1 | 30591 | 1 | 1 | 31 | 43694 | 10786 | 81 | 769 | 15728656 | 2 |
| 180 | 174 | 4 | 2 | 2 | 2 | 168 | 18374967684059824106 | 1 | 32767 | 1 | 1 | 31 | 43759 | 15155 | 16 | 1792 | 7536691 | 2 |
| 181 | 174 | 4 | 2 | 2 | 2 | 168 | 18410996206198128639 | 1 | 22527 | 1 | 1 | 31 | 43759 | 48043 | 16 | 4881 | 15794176 | 2 |
| 182 | 174 | 4 | 2 | 2 | 2 | 168 | 18410996687236562928 | 1 | 5631 | 1 | 1 | 31 | 43759 | 15155 | 16 | 769 | 3211312 | 2 |
| 183 | 174 | 4 | 2 | 2 | 2 | 168 | 18420003542893920224 | 1 | 22367 | 1 | 1 | 31 | 43694 | 15155 | 16 | 13057 | 3145744 | 2 |
| 184 | 174 | 4 | 2 | 2 | 2 | 168 | 18420003542894444458 | 1 | 8191 | 1 | 1 | 31 | 43694 | 8963 | 81 | 4881 | 16711729 | 2 |
| 185 | 174 | 4 | 2 | 2 | 2 | 168 | 18429011085750239216 | 1 | 1807 | 1 | 1 | 31 | 43759 | 10786 | 16 | 769 | 3211264 | 2 |
| 186 | 174 | 4 | 2 | 2 | 2 | 168 | 185405148299263 | 1 | 14335 | 1 | 1 | 31 | 43759 | 13091 | 16 | 4864 | 16711696 | 2 |
| 187 | 174 | 4 | 2 | 2 | 2 | 168 | 185408011545258 | 1 | 7999 | 1 | 1 | 31 | 43694 | 10786 | 16 | 13057 | 15794176 | 2 |
| 188 | 174 | 4 | 2 | 2 | 2 | 168 | 187466732579464 | 1 | 14143 | 1 | 1 | 31 | 43759 | 48043 | 81 | 13057 | 7340032 | 2 |
| 189 | 174 | 4 | 2 | 2 | 2 | 168 | 187607606685408 | 1 | 32767 | 1 | 1 | 31 | 43694 | 13058 | 81 | 4864 | 5308433 | 2 |
| 190 | 174 | 4 | 2 | 2 | 2 | 168 | 211655988396224 | 1 | 32767 | 1 | 1 | 31 | 43759 | 10786 | 16 | 769 | 7340080 | 2 |
| 191 | 174 | 4 | 2 | 2 | 2 | 168 | 211655988412330 | 1 | 1301 | 1 | 1 | 31 | 43694 | 8963 | 81 | 1792 | 7667712 | 2 |
| 192 | 174 | 4 | 2 | 2 | 2 | 168 | 211660030669040 | 1 | 14143 | 1 | 1 | 31 | 43759 | 10786 | 81 | 13057 | 3342352 | 2 |
| 193 | 174 | 4 | 2 | 2 | 2 | 168 | 211660283314175 | 1 | 21887 | 1 | 1 | 31 | 43694 | 8706 | 16 | 4864 | 3145744 | 2 |
| 194 | 174 | 4 | 2 | 2 | 2 | 168 | 211930866313920 | 1 | 14335 | 1 | 19 | 31 | 43759 | 48043 | 16 | 13057 | 5242896 | 2 |
| 195 | 174 | 4 | 2 | 2 | 2 | 168 | 211934095917248 | 1 | 22367 | 1 | 1 | 31 | 43759 | 15155 | 81 | 13057 | 16711696 | 2 |
| 196 | 174 | 4 | 2 | 2 | 2 | 168 | 211934297771208 | 1 | 7999 | 1 | 1 | 31 | 43759 | 10786 | 16 | 4353 | 1048576 | 2 |
| 197 | 174 | 4 | 2 | 2 | 2 | 168 | 211934297771208 | 1 | 8191 | 1 | 1 | 31 | 43694 | 8963 | 81 | 4881 | 7340032 | 2 |
| 198 | 174 | 4 | 2 | 2 | 2 | 168 | 220730327353536 | 1 | 21887 | 1 | 1 | 31 | 43759 | 10786 | 16 | 769 | 3211264 | 2 |
| 199 | 174 | 4 | 2 | 2 | 2 | 168 | 220731254243327 | 1 | 7999 | 1 | 1 | 31 | 43759 | 8994 | 81 | 13057 | 7536640 | 2 |
| 200 | 174 | 4 | 2 | 2 | 2 | 168 | 224853558873216 | 1 | 1807 | 1 | 1 | 31 | 43759 | 47931 | 16 | 4864 | 15728656 | 2 |
| 201 | 174 | 4 | 2 | 2 | 2 | 168 | 247115238403788 | 1 | 22527 | 1 | 1 | 31 | 43759 | 8963 | 81 | 4881 | 16711729 | 2 |
| 202 | 174 | 4 | 2 | 2 | 2 | 168 | 247119244750540 | 1 | 32767 | 1 | 1 | 31 | 43759 | 48043 | 16 | 14080 | 3211281 | 2 |
| 203 | 174 | 4 | 2 | 2 | 2 | 168 | 247119533309951 | 1 | 1301 | 1 | 1 | 31 | 43759 | 10786 | 81 | 5393 | 7667712 | 2 |
| 204 | 174 | 4 | 2 | 2 | 2 | 168 | 247252677361632 | 1 | 14143 | 1 | 1 | 31 | 43759 | 10786 | 16 | 769 | 16711729 | 2 |
| 205 | 174 | 4 | 2 | 2 | 2 | 168 | 258110354681548 | 1 | 32767 | 1 | 1 | 31 | 43759 | 8963 | 81 | 769 | 5308433 | 2 |
| 206 | 174 | 4 | 2 | 2 | 2 | 168 | 258110354684656 | 1 | 16255 | 1 | 1 | 31 | 43694 | 47931 | 81 | 4864 | 7340080 | 2 |
| 207 | 174 | 4 | 2 | 2 | 2 | 168 | 258114293131968 | 1 | 2047 | 1 | 21 | 31 | 43759 | 2819 | 16 | 4353 | 7340032 | 2 |
| 208 | 174 | 4 | 2 | 2 | 2 | 168 | 258114293131968 | 1 | 3871 | 1 | 1 | 31 | 43759 | 13091 | 16 | 4881 | 3211281 | 2 |
| 209 | 174 | 4 | 2 | 2 | 2 | 168 | 262645840150527 | 1 | 32767 | 1 | 1 | 31 | 43759 | 47931 | 16 | 13057 | 5308433 | 2 |
| 210 | 174 | 4 | 2 | 2 | 2 | 168 | 262650134069232 | 1 | 30591 | 1 | 1 | 31 | 43694 | 15155 | 81 | 4881 | 7340080 | 2 |
| 211 | 174 | 4 | 2 | 2 | 2 | 168 | 262650135052287 | 1 | 14335 | 1 | 1 | 31 | 43759 | 8963 | 16 | 4881 | 16711729 | 2 |
| 212 | 174 | 4 | 2 | 2 | 2 | 168 | 264436588802288 | 1 | 22367 | 1 | 1 | 31 | 43759 | 10786 | 16 | 4864 | 3342385 | 2 |
| 213 | 174 | 4 | 2 | 2 | 2 | 168 | 264569985498348 | 1 | 3967 | 1 | 1 | 31 | 43759 | 2819 | 81 | 4881 | 15728656 | 2 |
| 214 | 174 | 4 | 2 | 2 | 2 | 168 | 264849158307839 | 1 | 1807 | 1 | 19 | 31 | 43759 | 48043 | 16 | 4864 | 7340080 | 2 |

| Model No. | Functions in integer form at the nodes of the network | | | | | | | | | | | | | | | | | |
|---|---|---|---|---|---|---|---|---|---|---|---|---|---|---|---|---|---|---|
| | CK | ARR1 | SHY2 | AUXIAA | ARF | ARF10 | ARF5 | XAL1 | PLT | AUX | SCR | SHR | MIR166 | PHB | JKD | MGP | WOX5 | CLE40 |
| 215 | 174 | 4 | 2 | 2 | 2 | 168 | 264849158307839 | 1 | 8031 | 1 | 1 | 31 | 43759 | 8963 | 81 | 13057 | 15728656 | 2 |
| 216 | 174 | 4 | 2 | 2 | 2 | 168 | 264913582882814 | 1 | 8191 | 1 | 1 | 31 | 43759 | 10786 | 16 | 4881 | 16187443 | 2 |
| 217 | 174 | 4 | 2 | 2 | 2 | 168 | 264915730362608 | 1 | 30591 | 1 | 1 | 31 | 43759 | 47931 | 16 | 4864 | 3211264 | 2 |
| 218 | 174 | 4 | 2 | 2 | 2 | 168 | 264916443921144 | 1 | 21887 | 1 | 1 | 31 | 43694 | 10786 | 81 | 769 | 5242896 | 2 |
| 219 | 174 | 4 | 2 | 2 | 2 | 168 | 264916812493040 | 1 | 14335 | 1 | 1 | 31 | 43694 | 8963 | 81 | 769 | 15728656 | 2 |
| 220 | 174 | 4 | 2 | 2 | 2 | 168 | 264917575859952 | 1 | 22391 | 1 | 19 | 31 | 43694 | 48043 | 16 | 13057 | 5308416 | 2 |
| 221 | 174 | 4 | 2 | 2 | 2 | 168 | 264917876670462 | 1 | 16255 | 1 | 1 | 31 | 43759 | 47931 | 81 | 4864 | 1048576 | 2 |
| 222 | 174 | 4 | 2 | 2 | 2 | 168 | 273709675904240 | 1 | 24447 | 1 | 1 | 31 | 43759 | 15155 | 81 | 4864 | 16187443 | 2 |
| 223 | 174 | 4 | 2 | 2 | 2 | 168 | 280311022812912 | 1 | 8191 | 1 | 1 | 31 | 43759 | 13091 | 16 | 13057 | 15728656 | 2 |
| 224 | 174 | 4 | 2 | 2 | 2 | 168 | 281200098869216 | 1 | 8191 | 1 | 1 | 31 | 43694 | 47931 | 81 | 4864 | 7340032 | 2 |
| 225 | 174 | 4 | 2 | 2 | 2 | 168 | 281410264956912 | 1 | 16255 | 1 | 1 | 31 | 43759 | 8963 | 81 | 769 | 5308433 | 2 |
| 226 | 174 | 4 | 2 | 2 | 2 | 168 | 9223513326951115434 | 1 | 1301 | 1 | 1 | 31 | 43694 | 10786 | 16 | 4353 | 5242896 | 2 |
| 227 | 174 | 4 | 2 | 2 | 2 | 168 | 9223513326951115434 | 1 | 21847 | 1 | 1 | 31 | 43759 | 10786 | 16 | 4881 | 1114128 | 2 |
| 228 | 174 | 4 | 2 | 2 | 2 | 168 | 9223513326951115434 | 1 | 5461 | 1 | 1 | 31 | 43694 | 8963 | 81 | 13057 | 7340080 | 2 |
| 229 | 174 | 4 | 2 | 2 | 2 | 168 | 9223513326962256554 | 1 | 5631 | 1 | 1 | 31 | 43759 | 48043 | 16 | 5393 | 15794176 | 2 |
| 230 | 174 | 4 | 2 | 2 | 2 | 168 | 9223513327534918860 | 1 | 22367 | 1 | 1 | 31 | 43694 | 13091 | 81 | 769 | 3211312 | 2 |
| 231 | 174 | 4 | 2 | 2 | 2 | 168 | 9223513328377200639 | 1 | 8191 | 1 | 1 | 31 | 43694 | 47931 | 81 | 4881 | 5570576 | 2 |
| 232 | 174 | 4 | 2 | 2 | 2 | 168 | 9223548648065900704 | 1 | 3967 | 1 | 1 | 31 | 43759 | 10786 | 16 | 13057 | 3342385 | 2 |
| 233 | 174 | 4 | 2 | 2 | 2 | 168 | 9223548650204954623 | 1 | 8031 | 1 | 1 | 31 | 43694 | 15155 | 16 | 13057 | 15728656 | 2 |
| 234 | 174 | 4 | 2 | 2 | 2 | 168 | 9223583970405445872 | 1 | 14199 | 1 | 1 | 31 | 43759 | 47931 | 16 | 4864 | 7667712 | 2 |
| 235 | 174 | 4 | 2 | 2 | 2 | 168 | 9223583970573221887 | 1 | 14335 | 1 | 1 | 31 | 43759 | 10786 | 16 | 13057 | 15728656 | 2 |
| 236 | 174 | 4 | 2 | 2 | 2 | 168 | 9259542123273824896 | 1 | 32767 | 1 | 1 | 31 | 43759 | 8994 | 81 | 4881 | 3211264 | 2 |
| 237 | 174 | 4 | 2 | 2 | 2 | 168 | 9259542125152891120 | 1 | 30591 | 1 | 1 | 31 | 43759 | 13091 | 16 | 4864 | 1114128 | 2 |
| 238 | 174 | 4 | 2 | 2 | 2 | 168 | 9259542125404553215 | 1 | 24447 | 1 | 1 | 31 | 43759 | 48043 | 16 | 4881 | 7340080 | 2 |
| 239 | 174 | 4 | 2 | 2 | 2 | 168 | 9259550920444792000 | 1 | 22527 | 1 | 1 | 31 | 43759 | 8706 | 16 | 4864 | 3145744 | 2 |
| 240 | 174 | 4 | 2 | 2 | 2 | 168 | 9259577308184879264 | 1 | 8031 | 1 | 1 | 31 | 43694 | 15155 | 81 | 4881 | 5308433 | 2 |
| 241 | 174 | 4 | 2 | 2 | 2 | 168 | 9259577308925984716 | 1 | 30591 | 1 | 1 | 31 | 43759 | 15155 | 16 | 4881 | 7340080 | 2 |
| 242 | 174 | 4 | 2 | 2 | 2 | 168 | 9259612492018008256 | 1 | 14335 | 1 | 1 | 31 | 43694 | 47931 | 81 | 13057 | 15794176 | 2 |
| 243 | 174 | 4 | 2 | 2 | 2 | 168 | 9259612492556988576 | 1 | 14199 | 1 | 1 | 31 | 43759 | 10786 | 16 | 769 | 15794176 | 2 |
| 244 | 174 | 4 | 2 | 2 | 2 | 168 | 9259612492556988576 | 1 | 5631 | 1 | 1 | 31 | 43759 | 10786 | 81 | 4864 | 5242896 | 2 |
| 245 | 174 | 4 | 2 | 2 | 2 | 168 | 9259625687437593804 | 1 | 21887 | 1 | 1 | 31 | 43694 | 8706 | 16 | 1792 | 5242896 | 2 |
| 246 | 174 | 4 | 2 | 2 | 2 | 168 | 9259681762328510400 | 1 | 2047 | 1 | 1 | 31 | 43694 | 10786 | 81 | 13057 | 3342385 | 2 |
| 247 | 174 | 4 | 2 | 2 | 2 | 168 | 9259681763136962544 | 1 | 2047 | 1 | 1 | 31 | 43759 | 13091 | 16 | 4864 | 1048576 | 2 |
| 248 | 174 | 4 | 2 | 2 | 2 | 168 | 9799982875347561120 | 1 | 2047 | 1 | 19 | 31 | 43694 | 47931 | 16 | 4881 | 7536640 | 2 |
| 249 | 174 | 4 | 2 | 2 | 2 | 168 | 9799982876790358015 | 1 | 7999 | 1 | 1 | 31 | 43694 | 10786 | 16 | 769 | 3342385 | 2 |
| 250 | 174 | 4 | 2 | 2 | 2 | 168 | 9800106503129006079 | 1 | 8031 | 1 | 1 | 31 | 43694 | 48043 | 16 | 3841 | 3145744 | 2 |
| 251 | 174 | 4 | 2 | 2 | 2 | 168 | 9836011671804510192 | 1 | 30591 | 1 | 1 | 31 | 43694 | 8963 | 16 | 4881 | 7340080 | 2 |
| 252 | 174 | 4 | 2 | 2 | 2 | 168 | 9836086714045628400 | 1 | 16255 | 1 | 1 | 31 | 43759 | 2562 | 16 | 4881 | 5242896 | 2 |
| 253 | 174 | 4 | 2 | 2 | 2 | 168 | 9836142995297075184 | 1 | 30591 | 1 | 1 | 31 | 43694 | 10786 | 16 | 4881 | 16711696 | 2 |
| 254 | 174 | 4 | 2 | 2 | 2 | 224 | 11529391521727049968 | 1 | 32767 | 1 | 1 | 31 | 43759 | 2819 | 81 | 4881 | 1114128 | 2 |
| 255 | 174 | 4 | 2 | 2 | 2 | 224 | 11529391521727049968 | 1 | 8031 | 1 | 1 | 31 | 43694 | 10786 | 16 | 13057 | 5570576 | 2 |
| 256 | 174 | 4 | 2 | 2 | 2 | 224 | 11529391659083164908 | 1 | 32767 | 1 | 1 | 31 | 43759 | 10786 | 16 | 4864 | 7667712 | 2 |
| 257 | 174 | 4 | 2 | 2 | 2 | 224 | 11529391659096534252 | 1 | 1807 | 1 | 1 | 31 | 43694 | 48043 | 81 | 769 | 7340032 | 2 |
| 258 | 174 | 4 | 2 | 2 | 2 | 224 | 11529479963409445104 | 1 | 5631 | 1 | 1 | 31 | 43694 | 8706 | 81 | 4864 | 3211264 | 2 |
| 259 | 174 | 4 | 2 | 2 | 2 | 224 | 11529479963422028016 | 1 | 22527 | 1 | 1 | 31 | 43694 | 10786 | 81 | 769 | 1114128 | 2 |

| Model No. | Functions in integer form at the nodes of the network | | | | | | | | | | | | | | | | | |
|---|---|---|---|---|---|---|---|---|---|---|---|---|---|---|---|---|---|---|
| | CK | ARR1 | SHY2 | AUXIAA | ARF | ARF10 | ARF5 | XAL1 | PLT | AUX | SCR | SHR | MIR166 | PHB | JKD | MGP | WOX5 | CLE40 |
| 260 | 174 | 4 | 2 | 2 | 2 | 224 | 11565420317396496576 | 1 | 32767 | 1 | 1 | 31 | 43759 | 47931 | 16 | 769 | 1048576 | 2 |
| 261 | 174 | 4 | 2 | 2 | 2 | 224 | 11565420317396500428 | 1 | 7999 | 1 | 1 | 31 | 43759 | 47931 | 16 | 4881 | 3145744 | 2 |
| 262 | 174 | 4 | 2 | 2 | 2 | 224 | 11565508554550997184 | 1 | 32767 | 1 | 1 | 31 | 43759 | 48043 | 16 | 13057 | 16711729 | 2 |
| 263 | 174 | 4 | 2 | 2 | 2 | 224 | 11574436450353064608 | 1 | 16255 | 1 | 1 | 31 | 43759 | 15155 | 16 | 4881 | 16711729 | 2 |
| 264 | 174 | 4 | 2 | 2 | 2 | 224 | 11574515889900417264 | 1 | 21887 | 1 | 1 | 31 | 43694 | 13091 | 81 | 769 | 16711729 | 2 |
| 265 | 174 | 4 | 2 | 2 | 2 | 224 | 12150897202622954736 | 1 | 3967 | 1 | 1 | 31 | 43694 | 10786 | 16 | 13057 | 3211281 | 2 |
| 266 | 174 | 4 | 2 | 2 | 2 | 224 | 12150989905230626800 | 1 | 21887 | 1 | 1 | 31 | 43694 | 48043 | 81 | 4881 | 5570576 | 2 |
| 267 | 174 | 4 | 2 | 2 | 2 | 224 | 12249978456032455338 | 1 | 32767 | 1 | 1 | 31 | 43694 | 47931 | 81 | 4881 | 5570576 | 2 |
| 268 | 174 | 4 | 2 | 2 | 2 | 224 | 12249978593471408808 | 1 | 8191 | 1 | 1 | 31 | 43759 | 13091 | 81 | 769 | 5570576 | 2 |
| 269 | 174 | 4 | 2 | 2 | 2 | 224 | 12249978593480321704 | 1 | 14143 | 1 | 19 | 31 | 43759 | 48043 | 16 | 5393 | 7340032 | 2 |
| 270 | 174 | 4 | 2 | 2 | 2 | 224 | 12249978593480321704 | 1 | 32767 | 1 | 1 | 31 | 43759 | 10786 | 81 | 4881 | 5308433 | 2 |
| 271 | 174 | 4 | 2 | 2 | 2 | 224 | 12250049101012466416 | 1 | 16255 | 1 | 1 | 31 | 43694 | 47931 | 16 | 4864 | 15728656 | 2 |
| 272 | 174 | 4 | 2 | 2 | 2 | 224 | 12250066897999036156 | 1 | 32767 | 1 | 1 | 31 | 43694 | 15155 | 16 | 769 | 3211312 | 2 |
| 273 | 174 | 4 | 2 | 2 | 2 | 224 | 12286100986428194800 | 1 | 7999 | 1 | 1 | 31 | 43759 | 10786 | 81 | 13057 | 15728656 | 2 |
| 274 | 174 | 4 | 2 | 2 | 2 | 224 | 12286100986428850175 | 1 | 24447 | 1 | 19 | 31 | 43694 | 48043 | 16 | 769 | 5308433 | 2 |
| 275 | 174 | 4 | 2 | 2 | 2 | 224 | 12295014589755599528 | 1 | 32767 | 1 | 1 | 31 | 43759 | 10786 | 16 | 4864 | 1048576 | 2 |
| 276 | 174 | 4 | 2 | 2 | 2 | 224 | 13835269714693115016 | 1 | 14143 | 1 | 1 | 31 | 43759 | 8706 | 16 | 769 | 3342352 | 2 |
| 277 | 174 | 4 | 2 | 2 | 2 | 224 | 13835269715297104032 | 1 | 14335 | 1 | 1 | 31 | 43694 | 48043 | 81 | 4864 | 15728656 | 2 |
| 278 | 174 | 4 | 2 | 2 | 2 | 224 | 13835269715548766122 | 1 | 1301 | 1 | 1 | 31 | 43759 | 47931 | 16 | 13057 | 5570576 | 2 |
| 279 | 174 | 4 | 2 | 2 | 2 | 224 | 13835269990074346218 | 1 | 3967 | 1 | 1 | 31 | 43694 | 47931 | 81 | 13057 | 16187392 | 2 |
| 280 | 174 | 4 | 2 | 2 | 2 | 224 | 13835305036504166058 | 1 | 14199 | 1 | 1 | 31 | 43694 | 15155 | 16 | 5393 | 5308433 | 2 |
| 281 | 174 | 4 | 2 | 2 | 2 | 224 | 13835305037108146336 | 1 | 14199 | 1 | 1 | 31 | 43759 | 15155 | 16 | 13057 | 16187443 | 2 |
| 282 | 174 | 4 | 2 | 2 | 2 | 224 | 13835305037123875056 | 1 | 16255 | 1 | 1 | 31 | 43759 | 13091 | 81 | 769 | 5308416 | 2 |
| 283 | 174 | 4 | 2 | 2 | 2 | 224 | 13835305037123875056 | 1 | 30591 | 1 | 1 | 31 | 43759 | 8963 | 81 | 4864 | 5308416 | 2 |
| 284 | 174 | 4 | 2 | 2 | 2 | 224 | 13871298511519150320 | 1 | 30591 | 1 | 1 | 31 | 43759 | 15155 | 16 | 4881 | 15794176 | 2 |
| 285 | 174 | 4 | 2 | 2 | 2 | 224 | 13871351426575957930 | 1 | 3967 | 1 | 1 | 31 | 43759 | 48043 | 16 | 769 | 7536691 | 2 |
| 286 | 174 | 4 | 2 | 2 | 2 | 224 | 13889313185112050888 | 1 | 30591 | 1 | 1 | 31 | 43759 | 10786 | 16 | 769 | 1114128 | 2 |
| 287 | 174 | 4 | 2 | 2 | 2 | 224 | 13889313185615375082 | 1 | 8191 | 1 | 1 | 31 | 43694 | 15155 | 81 | 4881 | 5308416 | 2 |
| 288 | 174 | 4 | 2 | 2 | 2 | 224 | 13889313185615380479 | 1 | 8191 | 1 | 19 | 31 | 43694 | 48043 | 16 | 769 | 5242896 | 2 |
| 289 | 174 | 4 | 2 | 2 | 2 | 224 | 13889348506721775840 | 1 | 7999 | 1 | 1 | 31 | 43694 | 15155 | 81 | 4881 | 5242896 | 2 |
| 290 | 174 | 4 | 2 | 2 | 2 | 224 | 13889359365207554800 | 1 | 14335 | 1 | 1 | 31 | 43759 | 10786 | 81 | 4881 | 16187443 | 2 |
| 291 | 174 | 4 | 2 | 2 | 2 | 224 | 13889365961603611336 | 1 | 16255 | 1 | 1 | 31 | 43759 | 47931 | 81 | 769 | 3342352 | 2 |
| 292 | 174 | 4 | 2 | 2 | 2 | 224 | 13889365961603611336 | 1 | 8191 | 1 | 1 | 31 | 43759 | 10786 | 81 | 769 | 16187443 | 2 |
| 293 | 174 | 4 | 2 | 2 | 2 | 224 | 13889382454682255328 | 1 | 30591 | 1 | 1 | 31 | 43694 | 8963 | 16 | 13057 | 15728656 | 2 |
| 294 | 174 | 4 | 2 | 2 | 2 | 224 | 13889382454847930304 | 1 | 30591 | 1 | 1 | 31 | 43759 | 15155 | 81 | 13057 | 16187443 | 2 |
| 295 | 174 | 4 | 2 | 2 | 2 | 224 | 141289391685760 | 1 | 21887 | 1 | 1 | 31 | 43759 | 8963 | 16 | 769 | 7536691 | 2 |
| 296 | 174 | 4 | 2 | 2 | 2 | 224 | 141289525905536 | 1 | 21847 | 1 | 1 | 31 | 43759 | 8963 | 81 | 4353 | 15728656 | 2 |
| 297 | 174 | 4 | 2 | 2 | 2 | 224 | 141289928564864 | 1 | 14143 | 1 | 1 | 31 | 43759 | 48043 | 81 | 4353 | 3342352 | 2 |
| 298 | 174 | 4 | 2 | 2 | 2 | 224 | 141291270762624 | 1 | 5631 | 1 | 1 | 31 | 43694 | 13091 | 81 | 13057 | 1048576 | 2 |
| 299 | 174 | 4 | 2 | 2 | 2 | 224 | 14465782733441386696 | 1 | 8191 | 1 | 1 | 31 | 43694 | 15155 | 16 | 1792 | 7667712 | 2 |
| 300 | 174 | 4 | 2 | 2 | 2 | 224 | 14465787131555007680 | 1 | 21887 | 1 | 1 | 31 | 43694 | 10786 | 81 | 4881 | 7667712 | 2 |
| 301 | 174 | 4 | 2 | 2 | 2 | 224 | 14465787131555794124 | 1 | 1807 | 1 | 1 | 31 | 43694 | 15155 | 81 | 13057 | 7667712 | 2 |
| 302 | 174 | 4 | 2 | 2 | 2 | 224 | 14699974037287783628 | 1 | 14199 | 1 | 1 | 31 | 43694 | 13091 | 81 | 4353 | 5242896 | 2 |
| 303 | 174 | 4 | 2 | 2 | 2 | 224 | 14699974038143434656 | 1 | 5461 | 1 | 1 | 31 | 43694 | 10786 | 16 | 4353 | 5570576 | 2 |
| 304 | 174 | 4 | 2 | 2 | 2 | 224 | 14699974038160146431 | 1 | 3967 | 1 | 1 | 31 | 43694 | 13091 | 81 | 13057 | 7667712 | 2 |

| Model No. | Functions in integer form at the nodes of the network | | | | | | | | | | | | | | | | | |
|---|---|---|---|---|---|---|---|---|---|---|---|---|---|---|---|---|---|---|
| | CK | ARR1 | SHY2 | AUXIAA | ARF | ARF10 | ARF5 | XAL1 | PLT | AUX | SCR | SHR | MIR166 | PHB | JKD | MGP | WOX5 | CLE40 |
| 305 | 174 | 4 | 2 | 2 | 2 | 224 | 14699974312165690568 | 1 | 7999 | 1 | 1 | 31 | 43759 | 13091 | 81 | 4881 | 5570576 | 2 |
| 306 | 174 | 4 | 2 | 2 | 2 | 224 | 14700009359680401066 | 1 | 8031 | 1 | 1 | 31 | 43759 | 10786 | 16 | 13057 | 3342385 | 2 |
| 307 | 174 | 4 | 2 | 2 | 2 | 224 | 14700027294882266874 | 1 | 1287 | 1 | 1 | 31 | 43694 | 8963 | 81 | 1792 | 5308416 | 2 |
| 308 | 174 | 4 | 2 | 2 | 2 | 224 | 14736002834315136136 | 1 | 21887 | 1 | 1 | 31 | 43759 | 10786 | 81 | 4881 | 5308433 | 2 |
| 309 | 174 | 4 | 2 | 2 | 2 | 224 | 14736059047711080447 | 1 | 14143 | 1 | 1 | 31 | 43694 | 10786 | 16 | 4881 | 5570576 | 2 |
| 310 | 174 | 4 | 2 | 2 | 2 | 224 | 14754017507706728170 | 1 | 5631 | 1 | 1 | 31 | 43694 | 8706 | 16 | 4881 | 1048576 | 2 |
| 311 | 174 | 4 | 2 | 2 | 2 | 224 | 14754017508279255024 | 1 | 2047 | 1 | 1 | 31 | 43759 | 48043 | 81 | 769 | 7340080 | 2 |
| 312 | 174 | 4 | 2 | 2 | 2 | 224 | 14754017508279910399 | 1 | 14335 | 1 | 1 | 31 | 43694 | 15155 | 16 | 4881 | 15728656 | 2 |
| 313 | 174 | 4 | 2 | 2 | 2 | 224 | 14754073788958703610 | 1 | 8191 | 1 | 1 | 31 | 43759 | 15155 | 81 | 4881 | 16187392 | 2 |
| 314 | 174 | 4 | 2 | 2 | 2 | 224 | 150085627316352 | 1 | 14143 | 1 | 1 | 31 | 43759 | 10786 | 81 | 769 | 1048576 | 2 |
| 315 | 174 | 4 | 2 | 2 | 2 | 224 | 150085627316352 | 1 | 14335 | 1 | 1 | 31 | 43694 | 10786 | 16 | 13057 | 1048576 | 2 |
| 316 | 174 | 4 | 2 | 2 | 2 | 224 | 150085627316352 | 1 | 14335 | 1 | 1 | 31 | 43759 | 10786 | 16 | 4864 | 1048576 | 2 |
| 317 | 174 | 4 | 2 | 2 | 2 | 224 | 150085627316352 | 1 | 14335 | 1 | 1 | 31 | 43759 | 10786 | 81 | 13057 | 1048576 | 2 |
| 318 | 174 | 4 | 2 | 2 | 2 | 224 | 150085627316352 | 1 | 16255 | 1 | 1 | 31 | 43694 | 10786 | 16 | 4881 | 1048576 | 2 |
| 319 | 174 | 4 | 2 | 2 | 2 | 224 | 150085627316352 | 1 | 1807 | 1 | 1 | 31 | 43759 | 10786 | 81 | 13057 | 1048576 | 2 |
| 320 | 174 | 4 | 2 | 2 | 2 | 224 | 150085627316352 | 1 | 2047 | 1 | 1 | 31 | 43759 | 10786 | 16 | 769 | 1048576 | 2 |
| 321 | 174 | 4 | 2 | 2 | 2 | 224 | 150085627316352 | 1 | 21887 | 1 | 1 | 31 | 43694 | 10786 | 16 | 769 | 1048576 | 2 |
| 322 | 174 | 4 | 2 | 2 | 2 | 224 | 150085627316352 | 1 | 22367 | 1 | 1 | 31 | 43694 | 10786 | 16 | 4864 | 1048576 | 2 |
| 323 | 174 | 4 | 2 | 2 | 2 | 224 | 150085627316352 | 1 | 22367 | 1 | 1 | 31 | 43759 | 10786 | 81 | 4881 | 1048576 | 2 |
| 324 | 174 | 4 | 2 | 2 | 2 | 224 | 150085627316352 | 1 | 22527 | 1 | 1 | 31 | 43759 | 10786 | 16 | 13057 | 1048576 | 2 |
| 325 | 174 | 4 | 2 | 2 | 2 | 224 | 150085627316352 | 1 | 22527 | 1 | 1 | 31 | 43759 | 10786 | 81 | 13057 | 1048576 | 2 |
| 326 | 174 | 4 | 2 | 2 | 2 | 224 | 150085627316352 | 1 | 24447 | 1 | 1 | 31 | 43694 | 10786 | 81 | 4881 | 1048576 | 2 |
| 327 | 174 | 4 | 2 | 2 | 2 | 224 | 150085627316352 | 1 | 30591 | 1 | 1 | 31 | 43694 | 10786 | 81 | 769 | 1048576 | 2 |
| 328 | 174 | 4 | 2 | 2 | 2 | 224 | 150085627316352 | 1 | 30591 | 1 | 1 | 31 | 43694 | 8706 | 16 | 4864 | 3211312 | 2 |
| 329 | 174 | 4 | 2 | 2 | 2 | 224 | 150085627316352 | 1 | 32767 | 1 | 1 | 31 | 43694 | 10786 | 16 | 769 | 1048576 | 2 |
| 330 | 174 | 4 | 2 | 2 | 2 | 224 | 150085627316352 | 1 | 32767 | 1 | 1 | 31 | 43694 | 10786 | 81 | 13057 | 1048576 | 2 |
| 331 | 174 | 4 | 2 | 2 | 2 | 224 | 150085627316352 | 1 | 32767 | 1 | 1 | 31 | 43694 | 47931 | 81 | 13057 | 1048576 | 2 |
| 332 | 174 | 4 | 2 | 2 | 2 | 224 | 150085627316352 | 1 | 32767 | 1 | 1 | 31 | 43694 | 8706 | 16 | 769 | 1048576 | 2 |
| 333 | 174 | 4 | 2 | 2 | 2 | 224 | 150085627316352 | 1 | 32767 | 1 | 1 | 31 | 43694 | 8963 | 81 | 4864 | 1114128 | 2 |
| 334 | 174 | 4 | 2 | 2 | 2 | 224 | 150085627316352 | 1 | 32767 | 1 | 1 | 31 | 43694 | 8963 | 81 | 4881 | 7536640 | 2 |
| 335 | 174 | 4 | 2 | 2 | 2 | 224 | 150085627316352 | 1 | 32767 | 1 | 1 | 31 | 43759 | 10786 | 16 | 4881 | 1048576 | 2 |
| 336 | 174 | 4 | 2 | 2 | 2 | 224 | 150085627316352 | 1 | 32767 | 1 | 1 | 31 | 43759 | 10786 | 81 | 13057 | 1048576 | 2 |
| 337 | 174 | 4 | 2 | 2 | 2 | 224 | 150085627316352 | 1 | 32767 | 1 | 1 | 31 | 43759 | 10786 | 81 | 769 | 1048576 | 2 |
| 338 | 174 | 4 | 2 | 2 | 2 | 224 | 150085627316352 | 1 | 32767 | 1 | 1 | 31 | 43759 | 13091 | 81 | 13057 | 1048576 | 2 |
| 339 | 174 | 4 | 2 | 2 | 2 | 224 | 150085627316352 | 1 | 32767 | 1 | 1 | 31 | 43759 | 48043 | 16 | 4881 | 1048576 | 2 |
| 340 | 174 | 4 | 2 | 2 | 2 | 224 | 150085627316352 | 1 | 32767 | 1 | 1 | 31 | 43759 | 48043 | 81 | 4864 | 7340032 | 2 |
| 341 | 174 | 4 | 2 | 2 | 2 | 224 | 150085627316352 | 1 | 32767 | 1 | 1 | 31 | 43759 | 8963 | 16 | 4864 | 16711696 | 2 |
| 342 | 174 | 4 | 2 | 2 | 2 | 224 | 150085627316352 | 1 | 32767 | 1 | 1 | 31 | 43759 | 8963 | 81 | 769 | 1048576 | 2 |
| 343 | 174 | 4 | 2 | 2 | 2 | 224 | 150085627316352 | 1 | 3871 | 1 | 1 | 31 | 43759 | 10786 | 16 | 4353 | 1048576 | 2 |
| 344 | 174 | 4 | 2 | 2 | 2 | 224 | 150085627316352 | 1 | 3871 | 1 | 1 | 31 | 43759 | 15155 | 16 | 4881 | 1048576 | 2 |
| 345 | 174 | 4 | 2 | 2 | 2 | 224 | 150085627316352 | 1 | 3967 | 1 | 1 | 31 | 43759 | 10786 | 81 | 769 | 1048576 | 2 |
| 346 | 174 | 4 | 2 | 2 | 2 | 224 | 150085627316352 | 1 | 5631 | 1 | 1 | 31 | 43694 | 10786 | 16 | 13057 | 1048576 | 2 |
| 347 | 174 | 4 | 2 | 2 | 2 | 224 | 150085627316352 | 1 | 5631 | 1 | 1 | 31 | 43694 | 10786 | 16 | 769 | 1048576 | 2 |
| 348 | 174 | 4 | 2 | 2 | 2 | 224 | 150085627316352 | 1 | 5631 | 1 | 1 | 31 | 43694 | 10786 | 81 | 13057 | 1048576 | 2 |
| 349 | 174 | 4 | 2 | 2 | 2 | 224 | 150085627316352 | 1 | 5631 | 1 | 1 | 31 | 43694 | 10786 | 81 | 4881 | 1048576 | 2 |

| Model No. | Functions in integer form at the nodes of the network | | | | | | | | | | | | | | | | | |
|---|---|---|---|---|---|---|---|---|---|---|---|---|---|---|---|---|---|---|
| | CK | ARR1 | SHY2 | AUXIAA | ARF | ARF10 | ARF5 | XAL1 | PLT | AUX | SCR | SHR | MIR166 | PHB | JKD | MGP | WOX5 | CLE40 |
| 350 | 174 | 4 | 2 | 2 | 2 | 224 | 150085627316352 | 1 | 5631 | 1 | 1 | 31 | 43759 | 10786 | 81 | 4881 | 1048576 | 2 |
| 351 | 174 | 4 | 2 | 2 | 2 | 224 | 150085627316352 | 1 | 5631 | 1 | 1 | 31 | 43759 | 47931 | 81 | 4881 | 1048576 | 2 |
| 352 | 174 | 4 | 2 | 2 | 2 | 224 | 150085627316352 | 1 | 7999 | 1 | 1 | 31 | 43694 | 10786 | 16 | 13057 | 1048576 | 2 |
| 353 | 174 | 4 | 2 | 2 | 2 | 224 | 150085627316352 | 1 | 8191 | 1 | 1 | 31 | 43759 | 10786 | 16 | 4864 | 1048576 | 2 |
| 354 | 174 | 4 | 2 | 2 | 2 | 224 | 150085627316352 | 1 | 8191 | 1 | 1 | 31 | 43759 | 10786 | 16 | 769 | 1048576 | 2 |
| 355 | 174 | 4 | 2 | 2 | 2 | 224 | 150086199847584 | 1 | 1807 | 1 | 1 | 31 | 43694 | 10786 | 16 | 769 | 1114128 | 2 |
| 356 | 174 | 4 | 2 | 2 | 2 | 224 | 150086199847584 | 1 | 5631 | 1 | 1 | 31 | 43694 | 13091 | 16 | 13057 | 1048576 | 2 |
| 357 | 174 | 4 | 2 | 2 | 2 | 224 | 16141148321451409407 | 1 | 1807 | 1 | 1 | 31 | 43694 | 47931 | 16 | 4864 | 3211264 | 2 |
| 358 | 174 | 4 | 2 | 2 | 2 | 224 | 16141165981851513072 | 1 | 7999 | 1 | 1 | 31 | 43759 | 10786 | 81 | 4864 | 5308433 | 2 |
| 359 | 174 | 4 | 2 | 2 | 2 | 224 | 16141165982105268464 | 1 | 21887 | 1 | 1 | 31 | 43694 | 10786 | 81 | 769 | 5308433 | 2 |
| 360 | 174 | 4 | 2 | 2 | 2 | 224 | 16141165982120997104 | 1 | 5631 | 1 | 1 | 31 | 43759 | 10786 | 16 | 4881 | 15794176 | 2 |
| 361 | 174 | 4 | 2 | 2 | 2 | 224 | 16186184317483938032 | 1 | 32767 | 1 | 1 | 31 | 43694 | 10786 | 81 | 13057 | 16187443 | 2 |
| 362 | 174 | 4 | 2 | 2 | 2 | 224 | 16186201634759180202 | 1 | 8031 | 1 | 1 | 31 | 43694 | 13091 | 81 | 4881 | 16187392 | 2 |
| 363 | 174 | 4 | 2 | 2 | 2 | 224 | 16195191379033387200 | 1 | 5631 | 1 | 1 | 31 | 43759 | 48043 | 81 | 4881 | 7667712 | 2 |
| 364 | 174 | 4 | 2 | 2 | 2 | 224 | 16195191379033387200 | 1 | 8031 | 1 | 1 | 31 | 43694 | 10786 | 16 | 769 | 3211281 | 2 |
| 365 | 174 | 4 | 2 | 2 | 2 | 224 | 16195191516740776176 | 1 | 24447 | 1 | 1 | 31 | 43759 | 48043 | 81 | 4881 | 15794176 | 2 |
| 366 | 174 | 4 | 2 | 2 | 2 | 224 | 16195208971455102924 | 1 | 5631 | 1 | 1 | 31 | 43694 | 48043 | 81 | 13057 | 7536691 | 2 |
| 367 | 174 | 4 | 2 | 2 | 2 | 224 | 16204198715729178336 | 1 | 24447 | 1 | 1 | 31 | 43759 | 15155 | 81 | 4881 | 7340080 | 2 |
| 368 | 174 | 4 | 2 | 2 | 2 | 224 | 16204198715729182688 | 1 | 3871 | 1 | 1 | 31 | 43694 | 10786 | 16 | 769 | 1114128 | 2 |
| 369 | 174 | 4 | 2 | 2 | 2 | 224 | 16204214109161455600 | 1 | 14335 | 1 | 1 | 31 | 43694 | 10786 | 81 | 13057 | 7536640 | 2 |
| 370 | 174 | 4 | 2 | 2 | 2 | 224 | 16204216308437352447 | 1 | 3967 | 1 | 1 | 31 | 43694 | 47931 | 81 | 13057 | 15728656 | 2 |
| 371 | 174 | 4 | 2 | 2 | 2 | 224 | 16915778314696720320 | 1 | 5631 | 1 | 1 | 31 | 43694 | 2819 | 16 | 769 | 7340080 | 2 |
| 372 | 174 | 4 | 2 | 2 | 2 | 224 | 16915796113109090303 | 1 | 8031 | 1 | 1 | 31 | 43694 | 15155 | 16 | 769 | 3342352 | 2 |
| 373 | 174 | 4 | 2 | 2 | 2 | 224 | 16915801404797353983 | 1 | 22367 | 1 | 1 | 31 | 43759 | 13091 | 16 | 769 | 15794176 | 2 |
| 374 | 174 | 4 | 2 | 2 | 2 | 224 | 17050909255888732074 | 1 | 16255 | 1 | 1 | 31 | 43759 | 10786 | 81 | 13057 | 5308433 | 2 |
| 375 | 174 | 4 | 2 | 2 | 2 | 224 | 17050909255888732074 | 1 | 30591 | 1 | 1 | 31 | 43694 | 47931 | 81 | 4881 | 7340032 | 2 |
| 376 | 174 | 4 | 2 | 2 | 2 | 224 | 17050909256179515391 | 1 | 14335 | 1 | 1 | 31 | 43694 | 48043 | 81 | 769 | 16711696 | 2 |
| 377 | 174 | 4 | 2 | 2 | 2 | 224 | 17050909256179515391 | 1 | 32767 | 1 | 1 | 31 | 43694 | 13091 | 81 | 4881 | 3211264 | 2 |
| 378 | 174 | 4 | 2 | 2 | 2 | 224 | 17149970031145189360 | 1 | 5461 | 1 | 1 | 31 | 43694 | 8706 | 16 | 769 | 3211281 | 2 |
| 379 | 174 | 4 | 2 | 2 | 2 | 224 | 17149970031145189360 | 1 | 5461 | 1 | 1 | 31 | 43694 | 8963 | 16 | 4881 | 15728656 | 2 |
| 380 | 174 | 4 | 2 | 2 | 2 | 224 | 17149987691781095166 | 1 | 22367 | 1 | 1 | 31 | 43759 | 15155 | 16 | 4864 | 15728656 | 2 |
| 381 | 174 | 4 | 2 | 2 | 2 | 224 | 17149987692050710512 | 1 | 5461 | 1 | 1 | 31 | 43759 | 8706 | 16 | 4864 | 1048576 | 2 |
| 382 | 174 | 4 | 2 | 2 | 2 | 224 | 17294087005675778208 | 1 | 2047 | 1 | 1 | 31 | 43759 | 48043 | 81 | 4353 | 7340032 | 2 |
| 383 | 174 | 4 | 2 | 2 | 2 | 224 | 17294087005675778240 | 1 | 21847 | 1 | 1 | 31 | 43759 | 8994 | 16 | 4881 | 7536640 | 2 |
| 384 | 174 | 4 | 2 | 2 | 2 | 224 | 17294087280721459912 | 1 | 1301 | 1 | 1 | 31 | 43694 | 47931 | 16 | 14080 | 16711729 | 2 |
| 385 | 174 | 4 | 2 | 2 | 2 | 224 | 17294087418008367344 | 1 | 3967 | 1 | 1 | 31 | 43759 | 10786 | 81 | 4864 | 1048576 | 2 |
| 386 | 174 | 4 | 2 | 2 | 2 | 224 | 17294087418261012479 | 1 | 32767 | 1 | 1 | 31 | 43759 | 13091 | 16 | 4864 | 5308416 | 2 |
| 387 | 174 | 4 | 2 | 2 | 2 | 224 | 17294087486846335224 | 1 | 7999 | 1 | 1 | 31 | 43694 | 13091 | 16 | 14080 | 3145744 | 2 |
| 388 | 174 | 4 | 2 | 2 | 2 | 224 | 17294087486846335224 | 1 | 8031 | 1 | 1 | 31 | 43759 | 13091 | 16 | 5393 | 7667712 | 2 |
| 389 | 174 | 4 | 2 | 2 | 2 | 224 | 17330115802703130784 | 1 | 24447 | 1 | 19 | 31 | 43759 | 47931 | 16 | 4881 | 1048576 | 2 |
| 390 | 174 | 4 | 2 | 2 | 2 | 224 | 17330115802703130816 | 1 | 24447 | 1 | 1 | 31 | 43759 | 8963 | 81 | 13057 | 7340080 | 2 |
| 391 | 174 | 4 | 2 | 2 | 2 | 224 | 17330116077833224140 | 1 | 22527 | 1 | 1 | 31 | 43694 | 8994 | 81 | 4881 | 5242896 | 2 |
| 392 | 174 | 4 | 2 | 2 | 2 | 224 | 17330116077841022975 | 1 | 1807 | 1 | 1 | 31 | 43694 | 10786 | 81 | 4881 | 5570576 | 2 |
| 393 | 174 | 4 | 2 | 2 | 2 | 224 | 17339123414276829424 | 1 | 14143 | 1 | 1 | 31 | 43694 | 10786 | 81 | 4881 | 15794176 | 2 |
| 394 | 174 | 4 | 2 | 2 | 2 | 224 | 17348130613536813296 | 1 | 24447 | 1 | 1 | 31 | 43759 | 10786 | 16 | 13057 | 7667712 | 2 |

| Model No. | Functions in integer form at the nodes of the network | | | | | | | | | | | | | | | | | |
|---|---|---|---|---|---|---|---|---|---|---|---|---|---|---|---|---|---|---|
| | CK | ARR1 | SHY2 | AUXIAA | ARF | ARF10 | ARF5 | XAL1 | PLT | AUX | SCR | SHR | MIR166 | PHB | JKD | MGP | WOX5 | CLE40 |
| 395 | 174 | 4 | 2 | 2 | 2 | 224 | 17357137881762562047 | 1 | 8191 | 1 | 1 | 31 | 43759 | 8963 | 81 | 769 | 16187392 | 2 |
| 396 | 174 | 4 | 2 | 2 | 2 | 224 | 17361641481273146104 | 1 | 32767 | 1 | 1 | 31 | 43759 | 10786 | 16 | 4881 | 5570576 | 2 |
| 397 | 174 | 4 | 2 | 2 | 2 | 224 | 17361650277432753392 | 1 | 14143 | 1 | 1 | 31 | 43694 | 10786 | 81 | 769 | 15728656 | 2 |
| 398 | 174 | 4 | 2 | 2 | 2 | 224 | 17361652476254682876 | 1 | 22367 | 1 | 1 | 31 | 43694 | 10786 | 81 | 4881 | 15794176 | 2 |
| 399 | 174 | 4 | 2 | 2 | 2 | 224 | 176475658580208 | 1 | 8191 | 1 | 1 | 31 | 43759 | 8706 | 16 | 4864 | 1048576 | 2 |
| 400 | 174 | 4 | 2 | 2 | 2 | 224 | 176612276494496 | 1 | 5461 | 1 | 1 | 31 | 43694 | 15155 | 16 | 769 | 5308433 | 2 |
| 401 | 174 | 4 | 2 | 2 | 2 | 224 | 17938111029669068016 | 1 | 8191 | 1 | 1 | 31 | 43759 | 10786 | 81 | 4881 | 5308433 | 2 |
| 402 | 174 | 4 | 2 | 2 | 2 | 224 | 18082232814848245744 | 1 | 24447 | 1 | 1 | 31 | 43759 | 47931 | 81 | 13057 | 3211312 | 2 |
| 403 | 174 | 4 | 2 | 2 | 2 | 224 | 18226348003007987696 | 1 | 3967 | 1 | 19 | 31 | 43694 | 48043 | 16 | 4881 | 1048576 | 2 |
| 404 | 174 | 4 | 2 | 2 | 2 | 224 | 18374967409170775944 | 1 | 21887 | 1 | 1 | 31 | 43759 | 47931 | 81 | 4881 | 3342352 | 2 |
| 405 | 174 | 4 | 2 | 2 | 2 | 224 | 18374967409170775944 | 1 | 5631 | 1 | 1 | 31 | 43694 | 15155 | 16 | 4881 | 7340032 | 2 |
| 406 | 174 | 4 | 2 | 2 | 2 | 224 | 18374967409170775968 | 1 | 8031 | 1 | 1 | 31 | 43759 | 10786 | 81 | 4881 | 16711729 | 2 |
| 407 | 174 | 4 | 2 | 2 | 2 | 224 | 18374967409170775978 | 1 | 3871 | 1 | 1 | 31 | 43694 | 47931 | 81 | 4864 | 3211281 | 2 |
| 408 | 174 | 4 | 2 | 2 | 2 | 224 | 18374967409170775978 | 1 | 5461 | 1 | 1 | 31 | 43759 | 48043 | 81 | 1281 | 1114128 | 2 |
| 409 | 174 | 4 | 2 | 2 | 2 | 224 | 18374967409170776048 | 1 | 3967 | 1 | 1 | 31 | 43759 | 10786 | 16 | 13057 | 5308433 | 2 |
| 410 | 174 | 4 | 2 | 2 | 2 | 224 | 18374967409179688840 | 1 | 16255 | 1 | 1 | 31 | 43759 | 13058 | 81 | 1281 | 7340032 | 2 |
| 411 | 174 | 4 | 2 | 2 | 2 | 224 | 18374967409181261728 | 1 | 16255 | 1 | 1 | 31 | 43694 | 47931 | 16 | 769 | 15794176 | 2 |
| 412 | 174 | 4 | 2 | 2 | 2 | 224 | 18374967409181917098 | 1 | 8031 | 1 | 1 | 31 | 43694 | 8706 | 16 | 4864 | 3211281 | 2 |
| 413 | 174 | 4 | 2 | 2 | 2 | 224 | 18374967409186504688 | 1 | 21887 | 1 | 1 | 31 | 43694 | 8994 | 81 | 769 | 15794176 | 2 |
| 414 | 174 | 4 | 2 | 2 | 2 | 224 | 18374967546618642344 | 1 | 14335 | 1 | 1 | 31 | 43694 | 10786 | 16 | 13057 | 3211312 | 2 |
| 415 | 174 | 4 | 2 | 2 | 2 | 224 | 18374967546622312416 | 1 | 8191 | 1 | 1 | 31 | 43759 | 13091 | 16 | 4864 | 3145744 | 2 |
| 416 | 174 | 4 | 2 | 2 | 2 | 224 | 18374967684048682976 | 1 | 32767 | 1 | 1 | 31 | 43759 | 2562 | 16 | 4864 | 3145744 | 2 |
| 417 | 174 | 4 | 2 | 2 | 2 | 224 | 18374967821487636462 | 1 | 1301 | 1 | 1 | 31 | 43759 | 13091 | 16 | 1281 | 5308433 | 2 |
| 418 | 174 | 4 | 2 | 2 | 2 | 224 | 18374967821502316512 | 1 | 22527 | 1 | 1 | 31 | 43759 | 2819 | 16 | 4881 | 3211264 | 2 |
| 419 | 174 | 4 | 2 | 2 | 2 | 224 | 18374967890207113208 | 1 | 24447 | 1 | 1 | 31 | 43694 | 10786 | 16 | 13057 | 1048576 | 2 |
| 420 | 174 | 4 | 2 | 2 | 2 | 224 | 18374967890207113214 | 1 | 5461 | 1 | 1 | 31 | 43759 | 8994 | 81 | 21761 | 5308433 | 2 |
| 421 | 174 | 4 | 2 | 2 | 2 | 224 | 18410996206198128544 | 1 | 14143 | 1 | 1 | 31 | 43759 | 15155 | 16 | 13057 | 5308433 | 2 |
| 422 | 174 | 4 | 2 | 2 | 2 | 224 | 18410996343637606314 | 1 | 32767 | 1 | 1 | 31 | 43694 | 10786 | 81 | 769 | 3342352 | 2 |
| 423 | 174 | 4 | 2 | 2 | 2 | 224 | 18410996343644422128 | 1 | 24447 | 1 | 1 | 31 | 43759 | 15155 | 16 | 4864 | 3211264 | 2 |
| 424 | 174 | 4 | 2 | 2 | 2 | 224 | 18410996481081016268 | 1 | 8031 | 1 | 1 | 31 | 43694 | 47931 | 81 | 13057 | 7340080 | 2 |
| 425 | 174 | 4 | 2 | 2 | 2 | 224 | 18420003542893920168 | 1 | 3871 | 1 | 1 | 31 | 43694 | 47931 | 16 | 22272 | 16187392 | 2 |
| 426 | 174 | 4 | 2 | 2 | 2 | 224 | 18420003542898114544 | 1 | 21887 | 1 | 19 | 31 | 43759 | 47931 | 16 | 4881 | 15794176 | 2 |
| 427 | 174 | 4 | 2 | 2 | 2 | 224 | 18420003817771827184 | 1 | 30591 | 1 | 1 | 31 | 43759 | 48043 | 16 | 4881 | 16187392 | 2 |
| 428 | 174 | 4 | 2 | 2 | 2 | 224 | 18429010879591809008 | 1 | 2047 | 1 | 1 | 31 | 43694 | 10786 | 81 | 4881 | 3211281 | 2 |
| 429 | 174 | 4 | 2 | 2 | 2 | 224 | 18429011017030762464 | 1 | 24447 | 1 | 1 | 31 | 43694 | 10786 | 16 | 769 | 15728656 | 2 |
| 430 | 174 | 4 | 2 | 2 | 2 | 224 | 185405148299263 | 1 | 32767 | 1 | 1 | 31 | 43759 | 43522 | 16 | 13057 | 7536640 | 2 |
| 431 | 174 | 4 | 2 | 2 | 2 | 224 | 187469595847338 | 1 | 21887 | 1 | 1 | 31 | 43759 | 10786 | 81 | 13057 | 16711696 | 2 |
| 432 | 174 | 4 | 2 | 2 | 2 | 224 | 187471027503103 | 1 | 16255 | 1 | 1 | 31 | 43694 | 10786 | 16 | 4881 | 5308416 | 2 |
| 433 | 174 | 4 | 2 | 2 | 2 | 224 | 211659218010240 | 1 | 5631 | 1 | 1 | 31 | 43759 | 10786 | 16 | 769 | 1048576 | 2 |
| 434 | 174 | 4 | 2 | 2 | 2 | 224 | 211933013786816 | 1 | 2047 | 1 | 1 | 31 | 43694 | 10786 | 81 | 4881 | 15794176 | 2 |
| 435 | 174 | 4 | 2 | 2 | 2 | 224 | 211933550665920 | 1 | 1807 | 1 | 1 | 31 | 43759 | 13091 | 81 | 4353 | 3211264 | 2 |
| 436 | 174 | 4 | 2 | 2 | 2 | 224 | 224850127945727 | 1 | 32767 | 1 | 1 | 31 | 43759 | 15155 | 16 | 14080 | 15728656 | 2 |
| 437 | 174 | 4 | 2 | 2 | 2 | 224 | 225127868460768 | 1 | 2047 | 1 | 1 | 31 | 43694 | 48043 | 16 | 4881 | 5242896 | 2 |
| 438 | 174 | 4 | 2 | 2 | 2 | 224 | 246977799450864 | 1 | 2047 | 1 | 1 | 31 | 43759 | 48043 | 81 | 4864 | 15794176 | 2 |
| 439 | 174 | 4 | 2 | 2 | 2 | 224 | 246981841711344 | 1 | 3871 | 1 | 1 | 31 | 43694 | 15155 | 16 | 4353 | 15794176 | 2 |

| Model No. | Functions in integer form at the nodes of the network | | | | | | | | | | | | | | | | | |
|---|---|---|---|---|---|---|---|---|---|---|---|---|---|---|---|---|---|---|
| | CK | ARR1 | SHY2 | AUXIAA | ARF | ARF10 | ARF5 | XAL1 | PLT | AUX | SCR | SHR | MIR166 | PHB | JKD | MGP | WOX5 | CLE40 |
| 440 | 174 | 4 | 2 | 2 | 2 | 224 | 260176233889791 | 1 | 7999 | 1 | 1 | 31 | 43759 | 47931 | 81 | 4864 | 15794176 | 2 |
| 441 | 174 | 4 | 2 | 2 | 2 | 224 | 264436581462144 | 1 | 5631 | 1 | 1 | 31 | 43694 | 47931 | 81 | 4864 | 3342385 | 2 |
| 442 | 174 | 4 | 2 | 2 | 2 | 224 | 264436841447423 | 1 | 1807 | 1 | 1 | 31 | 43759 | 10786 | 81 | 769 | 3342385 | 2 |
| 443 | 174 | 4 | 2 | 2 | 2 | 224 | 264569985498280 | 1 | 8191 | 1 | 1 | 31 | 43759 | 10786 | 16 | 769 | 16711729 | 2 |
| 444 | 174 | 4 | 2 | 2 | 2 | 224 | 264848904614112 | 1 | 8031 | 1 | 1 | 31 | 43694 | 10786 | 81 | 769 | 15728656 | 2 |
| 445 | 174 | 4 | 2 | 2 | 2 | 224 | 264915730362608 | 1 | 5631 | 1 | 1 | 31 | 43759 | 15155 | 16 | 769 | 3211281 | 2 |
| 446 | 174 | 4 | 2 | 2 | 2 | 224 | 264917340975344 | 1 | 3967 | 1 | 1 | 31 | 43759 | 10786 | 16 | 13057 | 7667712 | 2 |
| 447 | 174 | 4 | 2 | 2 | 2 | 224 | 264917575859952 | 1 | 32767 | 1 | 1 | 31 | 43759 | 8963 | 81 | 4864 | 16187392 | 2 |
| 448 | 174 | 4 | 2 | 2 | 2 | 224 | 273713852381424 | 1 | 32767 | 1 | 1 | 31 | 43759 | 8963 | 16 | 769 | 1114128 | 2 |
| 449 | 174 | 4 | 2 | 2 | 2 | 224 | 275912135081724 | 1 | 1807 | 1 | 19 | 31 | 43759 | 48043 | 16 | 22272 | 1048576 | 2 |
| 450 | 174 | 4 | 2 | 2 | 2 | 224 | 278110551473392 | 1 | 21887 | 1 | 1 | 31 | 43759 | 10786 | 81 | 769 | 7667712 | 2 |
| 451 | 174 | 4 | 2 | 2 | 2 | 224 | 280929507540864 | 1 | 14143 | 1 | 1 | 31 | 43759 | 15155 | 81 | 769 | 5308416 | 2 |
| 452 | 174 | 4 | 2 | 2 | 2 | 224 | 280929515864063 | 1 | 3967 | 1 | 1 | 31 | 43694 | 8706 | 81 | 22272 | 1048576 | 2 |
| 453 | 174 | 4 | 2 | 2 | 2 | 224 | 281337537822704 | 1 | 8191 | 1 | 1 | 31 | 43759 | 10786 | 16 | 13057 | 3211281 | 2 |
| 454 | 174 | 4 | 2 | 2 | 2 | 224 | 281410265874430 | 1 | 30591 | 1 | 1 | 31 | 43694 | 10786 | 16 | 4881 | 5308433 | 2 |
| 455 | 174 | 4 | 2 | 2 | 2 | 224 | 9223513326380681352 | 1 | 14143 | 1 | 1 | 31 | 43759 | 10786 | 16 | 4353 | 7340032 | 2 |
| 456 | 174 | 4 | 2 | 2 | 2 | 224 | 9223513326951115434 | 1 | 1301 | 1 | 1 | 31 | 43759 | 47931 | 81 | 769 | 3211264 | 2 |
| 457 | 174 | 4 | 2 | 2 | 2 | 224 | 9223513328377200639 | 1 | 22391 | 1 | 1 | 31 | 43694 | 47931 | 81 | 4353 | 5242896 | 2 |
| 458 | 174 | 4 | 2 | 2 | 2 | 224 | 9259542125404553215 | 1 | 30591 | 1 | 1 | 31 | 43694 | 8706 | 16 | 4864 | 7536640 | 2 |
| 459 | 174 | 4 | 2 | 2 | 2 | 224 | 9259577446359957408 | 1 | 2047 | 1 | 1 | 31 | 43759 | 48043 | 16 | 769 | 16187392 | 2 |
| 460 | 174 | 4 | 2 | 2 | 2 | 224 | 9259612767600574400 | 1 | 16255 | 1 | 1 | 31 | 43694 | 47931 | 16 | 769 | 16187443 | 2 |
| 461 | 174 | 4 | 2 | 2 | 2 | 224 | 9259625686696525728 | 1 | 5631 | 1 | 1 | 31 | 43694 | 8963 | 16 | 13057 | 3145744 | 2 |
| 462 | 174 | 4 | 2 | 2 | 2 | 224 | 9259625687235480768 | 1 | 5631 | 1 | 1 | 31 | 43694 | 10786 | 81 | 4881 | 16711696 | 2 |
| 463 | 174 | 4 | 2 | 2 | 2 | 224 | 9259681763136962544 | 1 | 32767 | 1 | 1 | 31 | 43759 | 2819 | 81 | 4881 | 1048576 | 2 |
| 464 | 174 | 4 | 2 | 2 | 2 | 224 | 9799982874777127048 | 1 | 5461 | 1 | 1 | 31 | 43759 | 10786 | 81 | 13057 | 7536691 | 2 |
| 465 | 174 | 4 | 2 | 2 | 2 | 224 | 9799982875347561120 | 1 | 21847 | 1 | 1 | 31 | 43694 | 15155 | 81 | 4353 | 16711696 | 2 |
| 466 | 174 | 4 | 2 | 2 | 2 | 224 | 9799982875917995200 | 1 | 14199 | 1 | 1 | 31 | 43694 | 10786 | 16 | 769 | 3342352 | 2 |
| 467 | 174 | 4 | 2 | 2 | 2 | 224 | 9799982876790358015 | 1 | 14143 | 1 | 1 | 31 | 43694 | 8706 | 16 | 22272 | 16187443 | 2 |
| 468 | 174 | 4 | 2 | 2 | 2 | 224 | 9800018196588178090 | 1 | 14335 | 1 | 1 | 31 | 43694 | 8706 | 16 | 4881 | 3145744 | 2 |
| 469 | 174 | 4 | 2 | 2 | 2 | 224 | 9800018197158603424 | 1 | 1287 | 1 | 1 | 31 | 43694 | 10786 | 81 | 4353 | 15794176 | 2 |
| 470 | 174 | 4 | 2 | 2 | 2 | 224 | 9800053518399229132 | 1 | 21887 | 1 | 1 | 31 | 43694 | 47931 | 16 | 13057 | 16187443 | 2 |
| 471 | 174 | 4 | 2 | 2 | 2 | 224 | 9800053518399229132 | 1 | 21887 | 1 | 1 | 31 | 43759 | 48043 | 16 | 1792 | 5308416 | 2 |
| 472 | 174 | 4 | 2 | 2 | 2 | 224 | 9800053518408141000 | 1 | 32767 | 1 | 1 | 31 | 43759 | 10786 | 81 | 4881 | 1048576 | 2 |
| 473 | 174 | 4 | 2 | 2 | 2 | 224 | 9800053518969666288 | 1 | 21887 | 1 | 1 | 31 | 43759 | 47931 | 81 | 13057 | 3211264 | 2 |
| 474 | 174 | 4 | 2 | 2 | 2 | 224 | 9800053518969666288 | 1 | 3871 | 1 | 1 | 31 | 43694 | 47931 | 16 | 4864 | 5308416 | 2 |
| 475 | 174 | 4 | 2 | 2 | 2 | 224 | 9800106501124716792 | 1 | 30591 | 1 | 1 | 31 | 43759 | 48043 | 16 | 769 | 3211312 | 2 |
| 476 | 174 | 4 | 2 | 2 | 2 | 224 | 9836011671804510192 | 1 | 24447 | 1 | 1 | 31 | 43759 | 48043 | 16 | 4864 | 5308416 | 2 |
| 477 | 174 | 4 | 2 | 2 | 2 | 236 | 11529391521711321280 | 1 | 3967 | 1 | 1 | 31 | 43694 | 47931 | 81 | 4864 | 7340032 | 2 |
| 478 | 174 | 4 | 2 | 2 | 2 | 236 | 11529391658881835232 | 1 | 14335 | 1 | 1 | 31 | 43759 | 10786 | 81 | 4881 | 16187443 | 2 |
| 479 | 174 | 4 | 2 | 2 | 2 | 236 | 11529391659083164908 | 1 | 14199 | 1 | 1 | 31 | 43694 | 48043 | 16 | 3841 | 3342385 | 2 |
| 480 | 174 | 4 | 2 | 2 | 2 | 236 | 11529462165077549248 | 1 | 1807 | 1 | 1 | 31 | 43759 | 13091 | 16 | 4864 | 3342385 | 2 |
| 481 | 174 | 4 | 2 | 2 | 2 | 236 | 11529462165349134576 | 1 | 1807 | 1 | 1 | 31 | 43759 | 48043 | 81 | 4864 | 15728656 | 2 |
| 482 | 174 | 4 | 2 | 2 | 2 | 236 | 11565420456434270207 | 1 | 1807 | 1 | 1 | 31 | 43694 | 47931 | 16 | 769 | 16711696 | 2 |
| 483 | 174 | 4 | 2 | 2 | 2 | 236 | 11565420456437612543 | 1 | 8191 | 1 | 1 | 31 | 43759 | 8963 | 16 | 769 | 16711696 | 2 |
| 484 | 174 | 4 | 2 | 2 | 2 | 236 | 11574427654092270248 | 1 | 1287 | 1 | 1 | 31 | 43759 | 8963 | 81 | 769 | 15794176 | 2 |

| Model No. | Functions in integer form at the nodes of the network | | | | | | | | | | | | | | | | | |
|---|---|---|---|---|---|---|---|---|---|---|---|---|---|---|---|---|---|---|
| | CK | ARR1 | SHY2 | AUXIAA | ARF | ARF10 | ARF5 | XAL1 | PLT | AUX | SCR | SHR | MIR166 | PHB | JKD | MGP | WOX5 | CLE40 |
| 485 | 174 | 4 | 2 | 2 | 2 | 236 | 11574427654226487464 | 1 | 2047 | 1 | 1 | 31 | 43759 | 10786 | 81 | 13057 | 15794176 | 2 |
| 486 | 174 | 4 | 2 | 2 | 2 | 236 | 11574427655166025952 | 1 | 1807 | 1 | 1 | 31 | 43694 | 13091 | 16 | 769 | 5308433 | 2 |
| 487 | 174 | 4 | 2 | 2 | 2 | 236 | 11574427655170224352 | 1 | 16255 | 1 | 1 | 31 | 43694 | 47931 | 81 | 4881 | 1048576 | 2 |
| 488 | 174 | 4 | 2 | 2 | 2 | 236 | 11574427655367355628 | 1 | 22527 | 1 | 1 | 31 | 43759 | 10786 | 81 | 769 | 5308416 | 2 |
| 489 | 174 | 4 | 2 | 2 | 2 | 236 | 11574427655367360416 | 1 | 5631 | 1 | 1 | 31 | 43759 | 8963 | 81 | 4881 | 5308416 | 2 |
| 490 | 174 | 4 | 2 | 2 | 2 | 236 | 11574427655372341228 | 1 | 32767 | 1 | 1 | 31 | 43759 | 13091 | 16 | 13057 | 5308416 | 2 |
| 491 | 174 | 4 | 2 | 2 | 2 | 236 | 11574427655686127596 | 1 | 8031 | 1 | 1 | 31 | 43759 | 8994 | 81 | 13057 | 7340032 | 2 |
| 492 | 174 | 4 | 2 | 2 | 2 | 236 | 11574436450185291944 | 1 | 30591 | 1 | 1 | 31 | 43694 | 10786 | 81 | 13057 | 7340032 | 2 |
| 493 | 174 | 4 | 2 | 2 | 2 | 236 | 11574498024436531199 | 1 | 22527 | 1 | 1 | 31 | 43759 | 13091 | 81 | 4881 | 15794176 | 2 |
| 494 | 174 | 4 | 2 | 2 | 2 | 236 | 11574515959966265584 | 1 | 24447 | 1 | 1 | 31 | 43694 | 15155 | 81 | 4881 | 7340080 | 2 |
| 495 | 174 | 4 | 2 | 2 | 2 | 236 | 11574532107696930796 | 1 | 21887 | 1 | 19 | 31 | 43694 | 47931 | 16 | 4881 | 3211281 | 2 |
| 496 | 174 | 4 | 2 | 2 | 2 | 236 | 12249978456032455338 | 1 | 32767 | 1 | 1 | 31 | 43759 | 10786 | 16 | 4881 | 3211312 | 2 |
| 497 | 174 | 4 | 2 | 2 | 2 | 236 | 12249978456043596458 | 1 | 8191 | 1 | 19 | 31 | 43759 | 48043 | 16 | 13057 | 15794176 | 2 |
| 498 | 174 | 4 | 2 | 2 | 2 | 236 | 12249978457475252223 | 1 | 22367 | 1 | 1 | 31 | 43694 | 48043 | 16 | 1792 | 15728656 | 2 |
| 499 | 174 | 4 | 2 | 2 | 2 | 236 | 12250049099654557388 | 1 | 14143 | 1 | 1 | 31 | 43694 | 48043 | 16 | 3841 | 7667712 | 2 |
| 500 | 174 | 4 | 2 | 2 | 2 | 236 | 12250049099654557388 | 1 | 22367 | 1 | 1 | 31 | 43694 | 8706 | 16 | 769 | 5308416 | 2 |
| 501 | 174 | 4 | 2 | 2 | 2 | 236 | 12250049099665697514 | 1 | 7999 | 1 | 1 | 31 | 43694 | 15155 | 16 | 21761 | 16187392 | 2 |
| 502 | 174 | 4 | 2 | 2 | 2 | 236 | 12250049100740881088 | 1 | 16255 | 1 | 1 | 31 | 43694 | 15155 | 81 | 4864 | 1114128 | 2 |
| 503 | 174 | 4 | 2 | 2 | 2 | 236 | 12250066898007948024 | 1 | 22367 | 1 | 1 | 31 | 43694 | 48043 | 16 | 14080 | 5570576 | 2 |
| 504 | 174 | 4 | 2 | 2 | 2 | 236 | 12286100986428194800 | 1 | 21887 | 1 | 1 | 31 | 43694 | 8994 | 81 | 13057 | 7667712 | 2 |
| 505 | 174 | 4 | 2 | 2 | 2 | 236 | 12286100987856355264 | 1 | 22527 | 1 | 1 | 31 | 43759 | 2562 | 16 | 4881 | 16187443 | 2 |
| 506 | 174 | 4 | 2 | 2 | 2 | 236 | 12286100987860484095 | 1 | 1807 | 1 | 1 | 31 | 43759 | 10786 | 16 | 22272 | 7340032 | 2 |
| 507 | 174 | 4 | 2 | 2 | 2 | 236 | 12295014590900666352 | 1 | 7999 | 1 | 1 | 31 | 43694 | 48043 | 16 | 769 | 15728656 | 2 |
| 508 | 174 | 4 | 2 | 2 | 2 | 236 | 13835269990074346218 | 1 | 14335 | 1 | 1 | 31 | 43694 | 8963 | 81 | 769 | 7536691 | 2 |
| 509 | 174 | 4 | 2 | 2 | 2 | 236 | 13835305036302840048 | 1 | 22367 | 1 | 1 | 31 | 43694 | 10786 | 16 | 769 | 16711729 | 2 |
| 510 | 174 | 4 | 2 | 2 | 2 | 236 | 13835305036315418848 | 1 | 5631 | 1 | 1 | 31 | 43759 | 10786 | 81 | 769 | 5308433 | 2 |
| 511 | 174 | 4 | 2 | 2 | 2 | 236 | 13835305036504166058 | 1 | 3967 | 1 | 1 | 31 | 43759 | 2819 | 81 | 4353 | 5308433 | 2 |
| 512 | 174 | 4 | 2 | 2 | 2 | 236 | 13871298511519137984 | 1 | 22367 | 1 | 1 | 31 | 43694 | 8706 | 16 | 769 | 3145744 | 2 |
| 513 | 174 | 4 | 2 | 2 | 2 | 236 | 13871351425520431344 | 1 | 14143 | 1 | 1 | 31 | 43759 | 13091 | 16 | 13057 | 16711696 | 2 |
| 514 | 174 | 4 | 2 | 2 | 2 | 236 | 13871351426581528575 | 1 | 8191 | 1 | 1 | 31 | 43694 | 47931 | 16 | 14080 | 15794176 | 2 |
| 515 | 174 | 4 | 2 | 2 | 2 | 236 | 13871351769926267120 | 1 | 24447 | 1 | 1 | 31 | 43759 | 47931 | 16 | 4881 | 3211312 | 2 |
| 516 | 174 | 4 | 2 | 2 | 2 | 236 | 13889313184910731968 | 1 | 14143 | 1 | 1 | 31 | 43694 | 15155 | 81 | 4881 | 3145744 | 2 |
| 517 | 174 | 4 | 2 | 2 | 2 | 236 | 13889313185112575180 | 1 | 30591 | 1 | 1 | 31 | 43759 | 10786 | 16 | 4881 | 3211281 | 2 |
| 518 | 174 | 4 | 2 | 2 | 2 | 236 | 13889321981003745480 | 1 | 32767 | 1 | 1 | 31 | 43759 | 13091 | 16 | 1792 | 1114128 | 2 |
| 519 | 174 | 4 | 2 | 2 | 2 | 236 | 13889321981137963200 | 1 | 32767 | 1 | 1 | 31 | 43694 | 47931 | 81 | 769 | 15728656 | 2 |
| 520 | 174 | 4 | 2 | 2 | 2 | 236 | 13889348369819689152 | 1 | 32767 | 1 | 1 | 31 | 43694 | 15155 | 81 | 4864 | 1114128 | 2 |
| 521 | 174 | 4 | 2 | 2 | 2 | 236 | 13889366099716272368 | 1 | 8031 | 1 | 1 | 31 | 43759 | 8963 | 81 | 4881 | 5308416 | 2 |
| 522 | 174 | 4 | 2 | 2 | 2 | 236 | 13889366167829151738 | 1 | 8191 | 1 | 1 | 31 | 43759 | 8994 | 81 | 4881 | 7340032 | 2 |
| 523 | 174 | 4 | 2 | 2 | 2 | 236 | 141287244212864 | 1 | 3967 | 1 | 1 | 31 | 43694 | 8963 | 81 | 4864 | 16187443 | 2 |
| 524 | 174 | 4 | 2 | 2 | 2 | 236 | 141289391685760 | 1 | 5461 | 1 | 1 | 31 | 43759 | 15155 | 16 | 13057 | 16711696 | 2 |
| 525 | 174 | 4 | 2 | 2 | 2 | 236 | 141289525905536 | 1 | 22367 | 1 | 1 | 31 | 43694 | 47931 | 81 | 4881 | 3211281 | 2 |
| 526 | 174 | 4 | 2 | 2 | 2 | 236 | 14465782733441387712 | 1 | 7999 | 1 | 1 | 31 | 43759 | 13091 | 16 | 4864 | 3211264 | 2 |
| 527 | 174 | 4 | 2 | 2 | 2 | 236 | 14465782733441387724 | 1 | 8031 | 1 | 1 | 31 | 43759 | 13091 | 16 | 22272 | 5308416 | 2 |
| 528 | 174 | 4 | 2 | 2 | 2 | 236 | 14465787132414787583 | 1 | 16255 | 1 | 1 | 31 | 43759 | 15155 | 16 | 22272 | 15794176 | 2 |
| 529 | 174 | 4 | 2 | 2 | 2 | 236 | 14699974037287783616 | 1 | 1301 | 1 | 1 | 31 | 43694 | 47931 | 16 | 4353 | 16711696 | 2 |

| Model No. | Functions in integer form at the nodes of the network | | | | | | | | | | | | | | | | | |
|---|---|---|---|---|---|---|---|---|---|---|---|---|---|---|---|---|---|---|
| | CK | ARR1 | SHY2 | AUXIAA | ARF | ARF10 | ARF5 | XAL1 | PLT | AUX | SCR | SHR | MIR166 | PHB | JKD | MGP | WOX5 | CLE40 |
| 530 | 174 | 4 | 2 | 2 | 2 | 236 | 14699974312165690568 | 1 | 22527 | 1 | 1 | 31 | 43759 | 2562 | 16 | 4864 | 7536640 | 2 |
| 531 | 174 | 4 | 2 | 2 | 2 | 236 | 14700009359098838256 | 1 | 14199 | 1 | 1 | 31 | 43694 | 13091 | 81 | 4881 | 3211264 | 2 |
| 532 | 174 | 4 | 2 | 2 | 2 | 236 | 14700009359098838256 | 1 | 14199 | 1 | 1 | 31 | 43759 | 48043 | 16 | 4353 | 5308416 | 2 |
| 533 | 174 | 4 | 2 | 2 | 2 | 236 | 14700009359098839039 | 1 | 5631 | 1 | 1 | 31 | 43759 | 13091 | 81 | 769 | 16711696 | 2 |
| 534 | 174 | 4 | 2 | 2 | 2 | 236 | 14700009359646190752 | 1 | 3967 | 1 | 1 | 31 | 43694 | 10786 | 16 | 769 | 16187392 | 2 |
| 535 | 174 | 4 | 2 | 2 | 2 | 236 | 14700009359954476960 | 1 | 1807 | 1 | 1 | 31 | 43694 | 8994 | 81 | 4864 | 3342385 | 2 |
| 536 | 174 | 4 | 2 | 2 | 2 | 236 | 14700027294882266874 | 1 | 22391 | 1 | 1 | 31 | 43759 | 47931 | 81 | 13057 | 16187443 | 2 |
| 537 | 174 | 4 | 2 | 2 | 2 | 236 | 14700027294882266874 | 1 | 24447 | 1 | 1 | 31 | 43759 | 10786 | 81 | 4881 | 3211264 | 2 |
| 538 | 174 | 4 | 2 | 2 | 2 | 236 | 14736003110057017343 | 1 | 22527 | 1 | 1 | 31 | 43759 | 10786 | 81 | 13057 | 1114128 | 2 |
| 539 | 174 | 4 | 2 | 2 | 2 | 236 | 14754055028541030384 | 1 | 5631 | 1 | 1 | 31 | 43694 | 48043 | 81 | 769 | 3145744 | 2 |
| 540 | 174 | 4 | 2 | 2 | 2 | 236 | 150083337243840 | 1 | 3967 | 1 | 1 | 31 | 43759 | 15155 | 81 | 4353 | 5308416 | 2 |
| 541 | 174 | 4 | 2 | 2 | 2 | 236 | 150085627316352 | 1 | 14143 | 1 | 1 | 31 | 43694 | 8706 | 16 | 13057 | 1048576 | 2 |
| 542 | 174 | 4 | 2 | 2 | 2 | 236 | 150085627316352 | 1 | 14335 | 1 | 1 | 31 | 43694 | 10786 | 81 | 769 | 1048576 | 2 |
| 543 | 174 | 4 | 2 | 2 | 2 | 236 | 150085627316352 | 1 | 14335 | 1 | 1 | 31 | 43759 | 10786 | 16 | 4881 | 1048576 | 2 |
| 544 | 174 | 4 | 2 | 2 | 2 | 236 | 150085627316352 | 1 | 16255 | 1 | 1 | 31 | 43694 | 10786 | 16 | 13057 | 1048576 | 2 |
| 545 | 174 | 4 | 2 | 2 | 2 | 236 | 150085627316352 | 1 | 16255 | 1 | 1 | 31 | 43694 | 15155 | 81 | 4353 | 1048576 | 2 |
| 546 | 174 | 4 | 2 | 2 | 2 | 236 | 150085627316352 | 1 | 16255 | 1 | 1 | 31 | 43759 | 10786 | 81 | 769 | 1048576 | 2 |
| 547 | 174 | 4 | 2 | 2 | 2 | 236 | 150085627316352 | 1 | 1807 | 1 | 1 | 31 | 43694 | 10786 | 16 | 4881 | 1048576 | 2 |
| 548 | 174 | 4 | 2 | 2 | 2 | 236 | 150085627316352 | 1 | 1807 | 1 | 1 | 31 | 43694 | 10786 | 81 | 4353 | 1048576 | 2 |
| 549 | 174 | 4 | 2 | 2 | 2 | 236 | 150085627316352 | 1 | 1807 | 1 | 1 | 31 | 43759 | 10786 | 16 | 13057 | 1048576 | 2 |
| 550 | 174 | 4 | 2 | 2 | 2 | 236 | 150085627316352 | 1 | 2047 | 1 | 1 | 31 | 43759 | 10786 | 16 | 4881 | 1048576 | 2 |
| 551 | 174 | 4 | 2 | 2 | 2 | 236 | 150085627316352 | 1 | 21887 | 1 | 1 | 31 | 43759 | 15155 | 16 | 769 | 1048576 | 2 |
| 552 | 174 | 4 | 2 | 2 | 2 | 236 | 150085627316352 | 1 | 22367 | 1 | 1 | 31 | 43694 | 10786 | 16 | 4881 | 1048576 | 2 |
| 553 | 174 | 4 | 2 | 2 | 2 | 236 | 150085627316352 | 1 | 22391 | 1 | 1 | 31 | 43759 | 10786 | 16 | 13057 | 3145744 | 2 |
| 554 | 174 | 4 | 2 | 2 | 2 | 236 | 150085627316352 | 1 | 24447 | 1 | 1 | 31 | 43694 | 10786 | 16 | 769 | 1048576 | 2 |
| 555 | 174 | 4 | 2 | 2 | 2 | 236 | 150085627316352 | 1 | 24447 | 1 | 1 | 31 | 43694 | 10786 | 81 | 769 | 1048576 | 2 |
| 556 | 174 | 4 | 2 | 2 | 2 | 236 | 150085627316352 | 1 | 24447 | 1 | 1 | 31 | 43694 | 47931 | 16 | 4881 | 7340080 | 2 |
| 557 | 174 | 4 | 2 | 2 | 2 | 236 | 150085627316352 | 1 | 24447 | 1 | 1 | 31 | 43694 | 47931 | 81 | 13057 | 1048576 | 2 |
| 558 | 174 | 4 | 2 | 2 | 2 | 236 | 150085627316352 | 1 | 30591 | 1 | 1 | 31 | 43694 | 10786 | 81 | 769 | 1048576 | 2 |
| 559 | 174 | 4 | 2 | 2 | 2 | 236 | 150085627316352 | 1 | 30591 | 1 | 1 | 31 | 43759 | 10786 | 16 | 4881 | 1048576 | 2 |
| 560 | 174 | 4 | 2 | 2 | 2 | 236 | 150085627316352 | 1 | 32767 | 1 | 1 | 31 | 43694 | 10786 | 16 | 13057 | 1048576 | 2 |
| 561 | 174 | 4 | 2 | 2 | 2 | 236 | 150085627316352 | 1 | 32767 | 1 | 1 | 31 | 43694 | 10786 | 81 | 13057 | 1048576 | 2 |
| 562 | 174 | 4 | 2 | 2 | 2 | 236 | 150085627316352 | 1 | 32767 | 1 | 1 | 31 | 43694 | 10786 | 81 | 769 | 1048576 | 2 |
| 563 | 174 | 4 | 2 | 2 | 2 | 236 | 150085627316352 | 1 | 32767 | 1 | 1 | 31 | 43694 | 13091 | 16 | 4864 | 1048576 | 2 |
| 564 | 174 | 4 | 2 | 2 | 2 | 236 | 150085627316352 | 1 | 32767 | 1 | 1 | 31 | 43694 | 13091 | 16 | 4881 | 1048576 | 2 |
| 565 | 174 | 4 | 2 | 2 | 2 | 236 | 150085627316352 | 1 | 32767 | 1 | 1 | 31 | 43694 | 48043 | 81 | 13057 | 1048576 | 2 |
| 566 | 174 | 4 | 2 | 2 | 2 | 236 | 150085627316352 | 1 | 32767 | 1 | 1 | 31 | 43759 | 10786 | 16 | 769 | 1048576 | 2 |
| 567 | 174 | 4 | 2 | 2 | 2 | 236 | 150085627316352 | 1 | 32767 | 1 | 1 | 31 | 43759 | 10786 | 81 | 4881 | 1048576 | 2 |
| 568 | 174 | 4 | 2 | 2 | 2 | 236 | 150085627316352 | 1 | 32767 | 1 | 1 | 31 | 43759 | 13091 | 81 | 769 | 1048576 | 2 |
| 569 | 174 | 4 | 2 | 2 | 2 | 236 | 150085627316352 | 1 | 32767 | 1 | 1 | 31 | 43759 | 2562 | 16 | 4881 | 15794176 | 2 |
| 570 | 174 | 4 | 2 | 2 | 2 | 236 | 150085627316352 | 1 | 32767 | 1 | 1 | 31 | 43759 | 8706 | 16 | 4864 | 1048576 | 2 |
| 571 | 174 | 4 | 2 | 2 | 2 | 236 | 150085627316352 | 1 | 3871 | 1 | 1 | 31 | 43694 | 10786 | 16 | 4864 | 1048576 | 2 |
| 572 | 174 | 4 | 2 | 2 | 2 | 236 | 150085627316352 | 1 | 3871 | 1 | 1 | 31 | 43759 | 10786 | 16 | 4864 | 3342352 | 2 |
| 573 | 174 | 4 | 2 | 2 | 2 | 236 | 150085627316352 | 1 | 3967 | 1 | 1 | 31 | 43694 | 10786 | 81 | 13057 | 1048576 | 2 |
| 574 | 174 | 4 | 2 | 2 | 2 | 236 | 150085627316352 | 1 | 3967 | 1 | 1 | 31 | 43759 | 10786 | 81 | 769 | 1048576 | 2 |

| Model No. | Functions in integer form at the nodes of the network | | | | | | | | | | | | | | | | | |
|---|---|---|---|---|---|---|---|---|---|---|---|---|---|---|---|---|---|---|
| | CK | ARR1 | SHY2 | AUXIAA | ARF | ARF10 | ARF5 | XAL1 | PLT | AUX | SCR | SHR | MIR166 | PHB | JKD | MGP | WOX5 | CLE40 |
| 575 | 174 | 4 | 2 | 2 | 2 | 236 | 150085627316352 | 1 | 3967 | 1 | 1 | 31 | 43759 | 13091 | 16 | 13057 | 1048576 | 2 |
| 576 | 174 | 4 | 2 | 2 | 2 | 236 | 150085627316352 | 1 | 7999 | 1 | 1 | 31 | 43694 | 10786 | 16 | 769 | 1048576 | 2 |
| 577 | 174 | 4 | 2 | 2 | 2 | 236 | 150085627316352 | 1 | 8031 | 1 | 1 | 31 | 43759 | 10786 | 81 | 4864 | 1048576 | 2 |
| 578 | 174 | 4 | 2 | 2 | 2 | 236 | 150085627316352 | 1 | 8191 | 1 | 1 | 31 | 43694 | 10786 | 16 | 13057 | 1048576 | 2 |
| 579 | 174 | 4 | 2 | 2 | 2 | 236 | 150085627316352 | 1 | 8191 | 1 | 1 | 31 | 43694 | 10786 | 16 | 4353 | 1048576 | 2 |
| 580 | 174 | 4 | 2 | 2 | 2 | 236 | 16141148321451409407 | 1 | 32767 | 1 | 1 | 31 | 43759 | 47931 | 16 | 4864 | 7536640 | 2 |
| 581 | 174 | 4 | 2 | 2 | 2 | 236 | 16141165982373642239 | 1 | 21887 | 1 | 1 | 31 | 43694 | 15155 | 16 | 1792 | 3145744 | 2 |
| 582 | 174 | 4 | 2 | 2 | 2 | 236 | 16141165982373642239 | 1 | 3967 | 1 | 1 | 31 | 43694 | 8963 | 81 | 4881 | 7340080 | 2 |
| 583 | 174 | 4 | 2 | 2 | 2 | 236 | 16186184042337595050 | 1 | 22367 | 1 | 1 | 31 | 43694 | 48043 | 81 | 4881 | 1048576 | 2 |
| 584 | 174 | 4 | 2 | 2 | 2 | 236 | 16186201978389459184 | 1 | 1807 | 1 | 1 | 31 | 43759 | 48043 | 81 | 13057 | 5242896 | 2 |
| 585 | 174 | 4 | 2 | 2 | 2 | 236 | 16195191516474433760 | 1 | 14143 | 1 | 1 | 31 | 43759 | 15155 | 81 | 769 | 1048576 | 2 |
| 586 | 174 | 4 | 2 | 2 | 2 | 236 | 16195191516474433760 | 1 | 1807 | 1 | 1 | 31 | 43759 | 10786 | 16 | 769 | 1114128 | 2 |
| 587 | 174 | 4 | 2 | 2 | 2 | 236 | 16195208971455102924 | 1 | 2047 | 1 | 1 | 31 | 43694 | 2562 | 16 | 13057 | 16187443 | 2 |
| 588 | 174 | 4 | 2 | 2 | 2 | 236 | 16195208971491012848 | 1 | 3871 | 1 | 1 | 31 | 43759 | 48043 | 81 | 13057 | 3211312 | 2 |
| 589 | 174 | 4 | 2 | 2 | 2 | 236 | 16204198715729178336 | 1 | 22527 | 1 | 1 | 31 | 43759 | 47931 | 81 | 13057 | 3145744 | 2 |
| 590 | 174 | 4 | 2 | 2 | 2 | 236 | 16204216376869584880 | 1 | 21887 | 1 | 1 | 31 | 43759 | 2562 | 16 | 13057 | 5308416 | 2 |
| 591 | 174 | 4 | 2 | 2 | 2 | 236 | 16915796113109090303 | 1 | 24447 | 1 | 1 | 31 | 43694 | 8706 | 16 | 4881 | 15728656 | 2 |
| 592 | 174 | 4 | 2 | 2 | 2 | 236 | 17050906300922002684 | 1 | 14335 | 1 | 1 | 31 | 43759 | 47931 | 16 | 3841 | 5308433 | 2 |
| 593 | 174 | 4 | 2 | 2 | 2 | 236 | 17050906301242015743 | 1 | 8031 | 1 | 1 | 31 | 43759 | 48043 | 16 | 13057 | 5308433 | 2 |
| 594 | 174 | 4 | 2 | 2 | 2 | 236 | 17149987691765497855 | 1 | 8191 | 1 | 1 | 31 | 43694 | 8963 | 16 | 13057 | 3211312 | 2 |
| 595 | 174 | 4 | 2 | 2 | 2 | 236 | 17213020425657843696 | 1 | 8191 | 1 | 1 | 31 | 43759 | 2819 | 81 | 13057 | 1114128 | 2 |
| 596 | 174 | 4 | 2 | 2 | 2 | 236 | 17294087005675778208 | 1 | 32767 | 1 | 1 | 31 | 43759 | 2819 | 16 | 4864 | 5570576 | 2 |
| 597 | 174 | 4 | 2 | 2 | 2 | 236 | 17294087005675778288 | 1 | 8031 | 1 | 1 | 31 | 43694 | 47931 | 16 | 4881 | 5308433 | 2 |
| 598 | 174 | 4 | 2 | 2 | 2 | 236 | 17294087280721459912 | 1 | 2047 | 1 | 1 | 31 | 43694 | 13091 | 16 | 4864 | 3145744 | 2 |
| 599 | 174 | 4 | 2 | 2 | 2 | 236 | 17294087280721459912 | 1 | 3871 | 1 | 1 | 31 | 43694 | 13091 | 81 | 4353 | 15794176 | 2 |
| 600 | 174 | 4 | 2 | 2 | 2 | 236 | 17294087280732601066 | 1 | 22527 | 1 | 1 | 31 | 43759 | 15155 | 16 | 14080 | 16711696 | 2 |
| 601 | 174 | 4 | 2 | 2 | 2 | 236 | 17294087417992638704 | 1 | 5461 | 1 | 1 | 31 | 43694 | 48043 | 16 | 13057 | 3211281 | 2 |
| 602 | 174 | 4 | 2 | 2 | 2 | 236 | 17294087418007318752 | 1 | 14143 | 1 | 1 | 31 | 43694 | 10786 | 16 | 769 | 7340032 | 2 |
| 603 | 174 | 4 | 2 | 2 | 2 | 236 | 17294087418261012479 | 1 | 1301 | 1 | 1 | 31 | 43694 | 8963 | 81 | 769 | 16187443 | 2 |
| 604 | 174 | 4 | 2 | 2 | 2 | 236 | 17294087486855248120 | 1 | 14143 | 1 | 1 | 31 | 43759 | 15155 | 16 | 1281 | 3211264 | 2 |
| 605 | 174 | 4 | 2 | 2 | 2 | 236 | 17294087486947000062 | 1 | 14335 | 1 | 19 | 31 | 43694 | 48043 | 16 | 4881 | 7340032 | 2 |
| 606 | 174 | 4 | 2 | 2 | 2 | 236 | 17294087486947000062 | 1 | 3967 | 1 | 1 | 31 | 43694 | 47931 | 16 | 14080 | 1048576 | 2 |
| 607 | 174 | 4 | 2 | 2 | 2 | 236 | 17294087486962597630 | 1 | 8031 | 1 | 1 | 31 | 43759 | 47931 | 16 | 4881 | 16187443 | 2 |
| 608 | 174 | 4 | 2 | 2 | 2 | 236 | 17330115802703134719 | 1 | 21887 | 1 | 1 | 31 | 43694 | 8963 | 81 | 13057 | 16711729 | 2 |
| 609 | 174 | 4 | 2 | 2 | 2 | 236 | 17330116077841022975 | 1 | 14335 | 1 | 1 | 31 | 43759 | 47931 | 81 | 4881 | 16187392 | 2 |
| 610 | 174 | 4 | 2 | 2 | 2 | 236 | 17339123139398925480 | 1 | 7999 | 1 | 1 | 31 | 43694 | 10786 | 81 | 13057 | 1048576 | 2 |
| 611 | 174 | 4 | 2 | 2 | 2 | 236 | 17339123139600777130 | 1 | 21887 | 1 | 1 | 31 | 43759 | 15155 | 81 | 769 | 1048576 | 2 |
| 612 | 174 | 4 | 2 | 2 | 2 | 236 | 17339123139600777130 | 1 | 5461 | 1 | 1 | 31 | 43759 | 15155 | 81 | 4881 | 16711729 | 2 |
| 613 | 174 | 4 | 2 | 2 | 2 | 236 | 17348130476094716616 | 1 | 22367 | 1 | 1 | 31 | 43694 | 8706 | 16 | 769 | 3211312 | 2 |
| 614 | 174 | 4 | 2 | 2 | 2 | 236 | 17348130613535764704 | 1 | 32767 | 1 | 21 | 31 | 43759 | 2819 | 16 | 4881 | 16187392 | 2 |
| 615 | 174 | 4 | 2 | 2 | 2 | 236 | 17348130613536813296 | 1 | 24447 | 1 | 1 | 31 | 43759 | 8963 | 16 | 769 | 15728656 | 2 |
| 616 | 174 | 4 | 2 | 2 | 2 | 236 | 17348130682255241456 | 1 | 32767 | 1 | 1 | 31 | 43759 | 47931 | 81 | 4881 | 7536691 | 2 |
| 617 | 174 | 4 | 2 | 2 | 2 | 236 | 17357137881511031024 | 1 | 8031 | 1 | 1 | 31 | 43694 | 15155 | 81 | 4881 | 3211264 | 2 |
| 618 | 174 | 4 | 2 | 2 | 2 | 236 | 17361641481339731192 | 1 | 1807 | 1 | 1 | 31 | 43694 | 13091 | 16 | 4881 | 5308433 | 2 |
| 619 | 174 | 4 | 2 | 2 | 2 | 236 | 17361641481339731706 | 1 | 1301 | 1 | 1 | 31 | 43694 | 10762 | 81 | 769 | 7340032 | 2 |

| Model No. | Functions in integer form at the nodes of the network | | | | | | | | | | | | | | | | | |
|---|---|---|---|---|---|---|---|---|---|---|---|---|---|---|---|---|---|---|
| | CK | ARR1 | SHY2 | AUXIAA | ARF | ARF10 | ARF5 | XAL1 | PLT | AUX | SCR | SHR | MIR166 | PHB | JKD | MGP | WOX5 | CLE40 |
| 620 | 174 | 4 | 2 | 2 | 2 | 236 | 17361641481390587900 | 1 | 30591 | 1 | 1 | 31 | 43759 | 8706 | 16 | 769 | 15794176 | 2 |
| 621 | 174 | 4 | 2 | 2 | 2 | 236 | 17361652476389423864 | 1 | 7999 | 1 | 1 | 31 | 43694 | 47931 | 16 | 65281 | 1048576 | 2 |
| 622 | 174 | 4 | 2 | 2 | 2 | 236 | 17361657973947564024 | 1 | 32767 | 1 | 1 | 31 | 43759 | 8994 | 81 | 769 | 16711696 | 2 |
| 623 | 174 | 4 | 2 | 2 | 2 | 236 | 176471616301704 | 1 | 3967 | 1 | 1 | 31 | 43759 | 10786 | 81 | 4864 | 5570576 | 2 |
| 624 | 174 | 4 | 2 | 2 | 2 | 236 | 176474479569578 | 1 | 3967 | 1 | 1 | 31 | 43694 | 47931 | 16 | 4353 | 15794176 | 2 |
| 625 | 174 | 4 | 2 | 2 | 2 | 236 | 17938111029669068016 | 1 | 5631 | 1 | 1 | 31 | 43694 | 8963 | 81 | 769 | 5308433 | 2 |
| 626 | 174 | 4 | 2 | 2 | 2 | 236 | 17938111029669068799 | 1 | 22367 | 1 | 1 | 31 | 43694 | 48043 | 81 | 4881 | 16187392 | 2 |
| 627 | 174 | 4 | 2 | 2 | 2 | 236 | 17938113228725877488 | 1 | 7999 | 1 | 1 | 31 | 43694 | 13091 | 81 | 769 | 7340032 | 2 |
| 628 | 174 | 4 | 2 | 2 | 2 | 236 | 18370464290595471344 | 1 | 30591 | 1 | 1 | 31 | 43759 | 48043 | 16 | 4881 | 7340032 | 2 |
| 629 | 174 | 4 | 2 | 2 | 2 | 236 | 18374967409170775978 | 1 | 22367 | 1 | 1 | 31 | 43694 | 48043 | 16 | 13057 | 3211264 | 2 |
| 630 | 174 | 4 | 2 | 2 | 2 | 236 | 18374967409170775978 | 1 | 8191 | 1 | 1 | 31 | 43759 | 43522 | 81 | 769 | 3342385 | 2 |
| 631 | 174 | 4 | 2 | 2 | 2 | 236 | 18374967409170776000 | 1 | 16255 | 1 | 1 | 31 | 43694 | 8963 | 81 | 13057 | 16711696 | 2 |
| 632 | 174 | 4 | 2 | 2 | 2 | 236 | 18374967409170776000 | 1 | 30591 | 1 | 1 | 31 | 43759 | 47931 | 16 | 769 | 1048576 | 2 |
| 633 | 174 | 4 | 2 | 2 | 2 | 236 | 18374967409170776063 | 1 | 22527 | 1 | 1 | 31 | 43694 | 8994 | 81 | 4881 | 16711696 | 2 |
| 634 | 174 | 4 | 2 | 2 | 2 | 236 | 18374967409181917098 | 1 | 8031 | 1 | 1 | 31 | 43694 | 13091 | 16 | 1792 | 7536691 | 2 |
| 635 | 174 | 4 | 2 | 2 | 2 | 236 | 18374967409184145356 | 1 | 8031 | 1 | 1 | 31 | 43759 | 8994 | 16 | 1281 | 16187392 | 2 |
| 636 | 174 | 4 | 2 | 2 | 2 | 236 | 18374967546609729448 | 1 | 14199 | 1 | 1 | 31 | 43759 | 47931 | 81 | 4881 | 7536691 | 2 |
| 637 | 174 | 4 | 2 | 2 | 2 | 236 | 18374967684059824106 | 1 | 24447 | 1 | 1 | 31 | 43694 | 8963 | 81 | 4881 | 16187443 | 2 |
| 638 | 174 | 4 | 2 | 2 | 2 | 236 | 18374967821487636462 | 1 | 1287 | 1 | 1 | 31 | 43694 | 13091 | 16 | 1281 | 3145744 | 2 |
| 639 | 174 | 4 | 2 | 2 | 2 | 236 | 18374967821487636479 | 1 | 22391 | 1 | 1 | 31 | 43759 | 8994 | 16 | 21761 | 7667712 | 2 |
| 640 | 174 | 4 | 2 | 2 | 2 | 236 | 18374967821502316512 | 1 | 2047 | 1 | 1 | 31 | 43759 | 2819 | 81 | 769 | 7536691 | 2 |
| 641 | 174 | 4 | 2 | 2 | 2 | 236 | 18374967821503234030 | 1 | 30591 | 1 | 1 | 31 | 43759 | 15155 | 81 | 769 | 3211312 | 2 |
| 642 | 174 | 4 | 2 | 2 | 2 | 236 | 18374967890221793264 | 1 | 7999 | 1 | 19 | 31 | 43694 | 48043 | 16 | 4864 | 16187392 | 2 |
| 643 | 174 | 4 | 2 | 2 | 2 | 236 | 18374967890221793264 | 1 | 8031 | 1 | 1 | 31 | 43759 | 8963 | 81 | 4864 | 5308433 | 2 |
| 644 | 174 | 4 | 2 | 2 | 2 | 236 | 18374967890222710782 | 1 | 1807 | 1 | 19 | 31 | 43759 | 48043 | 16 | 4864 | 3211281 | 2 |
| 645 | 174 | 4 | 2 | 2 | 2 | 236 | 18410996206198128554 | 1 | 3967 | 1 | 1 | 31 | 43759 | 8994 | 81 | 769 | 15728656 | 2 |
| 646 | 174 | 4 | 2 | 2 | 2 | 236 | 18410996206198128639 | 1 | 3967 | 1 | 1 | 31 | 43694 | 47931 | 16 | 4864 | 7536691 | 2 |
| 647 | 174 | 4 | 2 | 2 | 2 | 236 | 18420003542898114544 | 1 | 32767 | 1 | 1 | 31 | 43759 | 48043 | 16 | 769 | 3342385 | 2 |
| 648 | 174 | 4 | 2 | 2 | 2 | 236 | 18420003817772482542 | 1 | 3967 | 1 | 1 | 31 | 43759 | 13091 | 16 | 769 | 16711729 | 2 |
| 649 | 174 | 4 | 2 | 2 | 2 | 236 | 18429011017031811056 | 1 | 1807 | 1 | 1 | 31 | 43694 | 10786 | 81 | 4881 | 5570576 | 2 |
| 650 | 174 | 4 | 2 | 2 | 2 | 236 | 187466732601280 | 1 | 1287 | 1 | 1 | 31 | 43694 | 10786 | 16 | 4353 | 3145744 | 2 |
| 651 | 174 | 4 | 2 | 2 | 2 | 236 | 187607607471852 | 1 | 14143 | 1 | 1 | 31 | 43694 | 15155 | 81 | 769 | 3211264 | 2 |
| 652 | 174 | 4 | 2 | 2 | 2 | 236 | 211655988399308 | 1 | 3967 | 1 | 1 | 31 | 43759 | 48043 | 81 | 4353 | 7340032 | 2 |
| 653 | 174 | 4 | 2 | 2 | 2 | 236 | 211930866313920 | 1 | 5631 | 1 | 1 | 31 | 43759 | 13091 | 16 | 4864 | 15794176 | 2 |
| 654 | 174 | 4 | 2 | 2 | 2 | 236 | 211933550665920 | 1 | 14143 | 1 | 1 | 31 | 43759 | 10786 | 81 | 769 | 1114128 | 2 |
| 655 | 174 | 4 | 2 | 2 | 2 | 236 | 211933718440640 | 1 | 30591 | 1 | 1 | 31 | 43759 | 47931 | 16 | 13057 | 16187392 | 2 |
| 656 | 174 | 4 | 2 | 2 | 2 | 236 | 211933718440640 | 1 | 3871 | 1 | 1 | 31 | 43759 | 47931 | 81 | 13057 | 3342352 | 2 |
| 657 | 174 | 4 | 2 | 2 | 2 | 236 | 211933718440640 | 1 | 3967 | 1 | 1 | 31 | 43759 | 10786 | 16 | 4864 | 1048576 | 2 |
| 658 | 174 | 4 | 2 | 2 | 2 | 236 | 220726959327432 | 1 | 32767 | 1 | 1 | 31 | 43694 | 2819 | 81 | 4881 | 15794176 | 2 |
| 659 | 174 | 4 | 2 | 2 | 2 | 236 | 220726959328448 | 1 | 16255 | 1 | 1 | 31 | 43694 | 2819 | 81 | 4864 | 7536640 | 2 |
| 660 | 174 | 4 | 2 | 2 | 2 | 236 | 220730327353536 | 1 | 32767 | 1 | 1 | 31 | 43694 | 48043 | 81 | 4881 | 7667712 | 2 |
| 661 | 174 | 4 | 2 | 2 | 2 | 236 | 220731169372912 | 1 | 24447 | 1 | 1 | 31 | 43759 | 2562 | 16 | 769 | 3342385 | 2 |
| 662 | 174 | 4 | 2 | 2 | 2 | 236 | 224850127945727 | 1 | 8191 | 1 | 1 | 31 | 43759 | 10786 | 16 | 13057 | 7340032 | 2 |
| 663 | 174 | 4 | 2 | 2 | 2 | 236 | 247252677361632 | 1 | 14335 | 1 | 1 | 31 | 43759 | 8963 | 16 | 4864 | 3342352 | 2 |
| 664 | 174 | 4 | 2 | 2 | 2 | 236 | 247252677361632 | 1 | 22527 | 1 | 21 | 31 | 43759 | 2819 | 16 | 4881 | 1048576 | 2 |

| Model No. | Functions in integer form at the nodes of the network | | | | | | | | | | | | | | | | | |
|---|---|---|---|---|---|---|---|---|---|---|---|---|---|---|---|---|---|---|
| | CK | ARR1 | SHY2 | AUXIAA | ARF | ARF10 | ARF5 | XAL1 | PLT | AUX | SCR | SHR | MIR166 | PHB | JKD | MGP | WOX5 | CLE40 |
| 665 | 174 | 4 | 2 | 2 | 2 | 236 | 260171938987936 | 1 | 30591 | 1 | 1 | 31 | 43759 | 13091 | 81 | 4864 | 5308433 | 2 |
| 666 | 174 | 4 | 2 | 2 | 2 | 236 | 260176233889791 | 1 | 30591 | 1 | 21 | 31 | 43759 | 2819 | 16 | 769 | 3211264 | 2 |
| 667 | 174 | 4 | 2 | 2 | 2 | 236 | 262649847803616 | 1 | 22367 | 1 | 1 | 31 | 43694 | 47931 | 16 | 4864 | 7536691 | 2 |
| 668 | 174 | 4 | 2 | 2 | 2 | 236 | 264432546541728 | 1 | 30591 | 1 | 1 | 31 | 43694 | 8963 | 81 | 4864 | 7340032 | 2 |
| 669 | 174 | 4 | 2 | 2 | 2 | 236 | 264573421419756 | 1 | 3871 | 1 | 1 | 31 | 43694 | 47931 | 81 | 769 | 1048576 | 2 |
| 670 | 174 | 4 | 2 | 2 | 2 | 236 | 264844863406062 | 1 | 32767 | 1 | 1 | 31 | 43759 | 43522 | 81 | 13057 | 3211312 | 2 |
| 671 | 174 | 4 | 2 | 2 | 2 | 236 | 264913582881008 | 1 | 21887 | 1 | 1 | 31 | 43759 | 13091 | 81 | 4864 | 5308416 | 2 |
| 672 | 174 | 4 | 2 | 2 | 2 | 236 | 264915864582384 | 1 | 3967 | 1 | 1 | 31 | 43759 | 47931 | 81 | 4864 | 7340032 | 2 |
| 673 | 174 | 4 | 2 | 2 | 2 | 236 | 264917014347000 | 1 | 3967 | 1 | 1 | 31 | 43759 | 8963 | 16 | 4881 | 5570576 | 2 |
| 674 | 174 | 4 | 2 | 2 | 2 | 236 | 264917589229308 | 1 | 14143 | 1 | 1 | 31 | 43759 | 47931 | 16 | 5393 | 15794176 | 2 |
| 675 | 174 | 4 | 2 | 2 | 2 | 236 | 264917589229308 | 1 | 30591 | 1 | 1 | 31 | 43694 | 8994 | 81 | 769 | 7667712 | 2 |
| 676 | 174 | 4 | 2 | 2 | 2 | 236 | 273709675903738 | 1 | 22527 | 1 | 19 | 31 | 43759 | 48043 | 16 | 769 | 5308416 | 2 |
| 677 | 174 | 4 | 2 | 2 | 2 | 236 | 273709675904240 | 1 | 3967 | 1 | 1 | 31 | 43759 | 2562 | 16 | 4864 | 7340032 | 2 |
| 678 | 174 | 4 | 2 | 2 | 2 | 236 | 273709675904252 | 1 | 21887 | 1 | 1 | 31 | 43759 | 47931 | 16 | 14080 | 7536640 | 2 |
| 679 | 174 | 4 | 2 | 2 | 2 | 236 | 278107722415352 | 1 | 22527 | 1 | 1 | 31 | 43759 | 10786 | 16 | 13057 | 5308433 | 2 |
| 680 | 174 | 4 | 2 | 2 | 2 | 236 | 278107722415352 | 1 | 24447 | 1 | 1 | 31 | 43759 | 15155 | 16 | 769 | 5308416 | 2 |
| 681 | 174 | 4 | 2 | 2 | 2 | 236 | 280306745671664 | 1 | 32767 | 1 | 21 | 31 | 43694 | 8994 | 16 | 4881 | 7340080 | 2 |
| 682 | 174 | 4 | 2 | 2 | 2 | 236 | 280929507540864 | 1 | 24447 | 1 | 1 | 31 | 43759 | 2819 | 16 | 769 | 1048576 | 2 |
| 683 | 174 | 4 | 2 | 2 | 2 | 236 | 281066095837164 | 1 | 14143 | 1 | 1 | 31 | 43694 | 10786 | 16 | 769 | 5308416 | 2 |
| 684 | 174 | 4 | 2 | 2 | 2 | 236 | 281200098869192 | 1 | 8191 | 1 | 1 | 31 | 43694 | 10786 | 81 | 13057 | 16711729 | 2 |
| 685 | 174 | 4 | 2 | 2 | 2 | 236 | 281408547913720 | 1 | 3871 | 1 | 1 | 31 | 43694 | 10786 | 16 | 13057 | 3342385 | 2 |
| 686 | 174 | 4 | 2 | 2 | 2 | 236 | 9223513326389594248 | 1 | 5631 | 1 | 1 | 31 | 43694 | 10786 | 16 | 13057 | 15728656 | 2 |
| 687 | 174 | 4 | 2 | 2 | 2 | 236 | 9223513326793826464 | 1 | 14143 | 1 | 1 | 31 | 43694 | 47931 | 81 | 769 | 5308416 | 2 |
| 688 | 174 | 4 | 2 | 2 | 2 | 236 | 9223513326951115434 | 1 | 14335 | 1 | 1 | 31 | 43694 | 47931 | 16 | 13057 | 7667712 | 2 |
| 689 | 174 | 4 | 2 | 2 | 2 | 236 | 9223513326951115434 | 1 | 21847 | 1 | 1 | 31 | 43694 | 48043 | 16 | 21761 | 3211281 | 2 |
| 690 | 174 | 4 | 2 | 2 | 2 | 236 | 9223513326962256554 | 1 | 14143 | 1 | 1 | 31 | 43694 | 47931 | 81 | 4881 | 7340032 | 2 |
| 691 | 174 | 4 | 2 | 2 | 2 | 236 | 9223513328377200639 | 1 | 5631 | 1 | 1 | 31 | 43694 | 15155 | 16 | 5393 | 7667712 | 2 |
| 692 | 174 | 4 | 2 | 2 | 2 | 236 | 9223548649332604927 | 1 | 14199 | 1 | 1 | 31 | 43759 | 15155 | 16 | 4864 | 16711696 | 2 |
| 693 | 174 | 4 | 2 | 2 | 2 | 236 | 9223583970954887360 | 1 | 14335 | 1 | 1 | 31 | 43694 | 15155 | 16 | 4864 | 1114128 | 2 |
| 694 | 174 | 4 | 2 | 2 | 2 | 236 | 9223583970954887360 | 1 | 8031 | 1 | 1 | 31 | 43694 | 8994 | 81 | 4864 | 7340032 | 2 |
| 695 | 174 | 4 | 2 | 2 | 2 | 236 | 9223583971157003468 | 1 | 3871 | 1 | 1 | 31 | 43759 | 48043 | 16 | 769 | 5308433 | 2 |
| 696 | 174 | 4 | 2 | 2 | 2 | 236 | 9223636953122009328 | 1 | 21847 | 1 | 1 | 31 | 43759 | 15155 | 81 | 769 | 15794176 | 2 |
| 697 | 174 | 4 | 2 | 2 | 2 | 236 | 9223636954732560383 | 1 | 14143 | 1 | 1 | 31 | 43759 | 47931 | 16 | 14080 | 16187443 | 2 |
| 698 | 174 | 4 | 2 | 2 | 2 | 236 | 9259542123273833600 | 1 | 14143 | 1 | 1 | 31 | 43694 | 8994 | 16 | 4881 | 16187443 | 2 |
| 699 | 174 | 4 | 2 | 2 | 2 | 236 | 9259542123273842816 | 1 | 16255 | 1 | 1 | 31 | 43694 | 48043 | 81 | 4864 | 16187392 | 2 |
| 700 | 174 | 4 | 2 | 2 | 2 | 236 | 9259542125152891120 | 1 | 16255 | 1 | 1 | 31 | 43759 | 8706 | 16 | 769 | 15794176 | 2 |
| 701 | 174 | 4 | 2 | 2 | 2 | 236 | 9259550919366838408 | 1 | 7999 | 1 | 1 | 31 | 43694 | 10786 | 16 | 4864 | 15794176 | 2 |
| 702 | 174 | 4 | 2 | 2 | 2 | 236 | 9259550921505898495 | 1 | 32767 | 1 | 1 | 31 | 43759 | 2562 | 16 | 4881 | 1048576 | 2 |
| 703 | 174 | 4 | 2 | 2 | 2 | 236 | 9259577309784965119 | 1 | 1807 | 1 | 1 | 31 | 43759 | 13091 | 81 | 4881 | 16187443 | 2 |
| 704 | 174 | 4 | 2 | 2 | 2 | 236 | 9259577445789510304 | 1 | 22527 | 1 | 1 | 31 | 43759 | 48043 | 16 | 13057 | 15794176 | 2 |
| 705 | 174 | 4 | 2 | 2 | 2 | 236 | 9259577445789510304 | 1 | 5631 | 1 | 1 | 31 | 43759 | 8994 | 81 | 4881 | 16187443 | 2 |
| 706 | 174 | 4 | 2 | 2 | 2 | 236 | 9259577447215595424 | 1 | 8031 | 1 | 1 | 31 | 43759 | 15155 | 16 | 4881 | 16187392 | 2 |
| 707 | 174 | 4 | 2 | 2 | 2 | 236 | 9259588303466834560 | 1 | 8191 | 1 | 1 | 31 | 43759 | 15155 | 81 | 4864 | 7536640 | 2 |
| 708 | 174 | 4 | 2 | 2 | 2 | 236 | 9259588303840149440 | 1 | 7999 | 1 | 1 | 31 | 43694 | 13091 | 16 | 4864 | 5570576 | 2 |
| 709 | 174 | 4 | 2 | 2 | 2 | 236 | 9259588304901242879 | 1 | 32767 | 1 | 1 | 31 | 43694 | 8963 | 16 | 4881 | 3211281 | 2 |

| Model No. | Functions in integer form at the nodes of the network | | | | | | | | | | | | | | | | | |
|---|---|---|---|---|---|---|---|---|---|---|---|---|---|---|---|---|---|---|
| | CK | ARR1 | SHY2 | AUXIAA | ARF | ARF10 | ARF5 | XAL1 | PLT | AUX | SCR | SHR | MIR166 | PHB | JKD | MGP | WOX5 | CLE40 |
| 710 | 174 | 4 | 2 | 2 | 2 | 236 | 9259588304901242879 | 1 | 3967 | 1 | 1 | 31 | 43694 | 13091 | 81 | 4864 | 7667712 | 2 |
| 711 | 174 | 4 | 2 | 2 | 2 | 236 | 9259612494157053951 | 1 | 14335 | 1 | 1 | 31 | 43759 | 15155 | 81 | 4881 | 5242896 | 2 |
| 712 | 174 | 4 | 2 | 2 | 2 | 236 | 9259612494157053951 | 1 | 1807 | 1 | 1 | 31 | 43759 | 10786 | 16 | 769 | 7667712 | 2 |
| 713 | 174 | 4 | 2 | 2 | 2 | 236 | 9259612768170986688 | 1 | 2047 | 1 | 1 | 31 | 43759 | 10786 | 81 | 4864 | 7340032 | 2 |
| 714 | 174 | 4 | 2 | 2 | 2 | 236 | 9259612768774975680 | 1 | 32767 | 1 | 1 | 31 | 43759 | 43522 | 16 | 769 | 5242896 | 2 |
| 715 | 174 | 4 | 2 | 2 | 2 | 236 | 9259625686696525728 | 1 | 3871 | 1 | 1 | 31 | 43759 | 47931 | 16 | 769 | 7536691 | 2 |
| 716 | 174 | 4 | 2 | 2 | 2 | 236 | 9259625686696525728 | 1 | 8191 | 1 | 1 | 31 | 43759 | 8706 | 16 | 4881 | 1114128 | 2 |
| 717 | 174 | 4 | 2 | 2 | 2 | 236 | 9259625687235480768 | 1 | 32767 | 1 | 19 | 31 | 43759 | 48043 | 16 | 4881 | 5308416 | 2 |
| 718 | 174 | 4 | 2 | 2 | 2 | 236 | 9259665270715187199 | 1 | 8191 | 1 | 1 | 31 | 43759 | 2819 | 81 | 13057 | 3342385 | 2 |
| 719 | 174 | 4 | 2 | 2 | 2 | 236 | 9259681761250574335 | 1 | 22527 | 1 | 1 | 31 | 43694 | 8706 | 16 | 769 | 3211281 | 2 |
| 720 | 174 | 4 | 2 | 2 | 2 | 236 | 9799982875358702250 | 1 | 14335 | 1 | 1 | 31 | 43759 | 8994 | 81 | 4864 | 7340032 | 2 |
| 721 | 174 | 4 | 2 | 2 | 2 | 236 | 9799982875358702250 | 1 | 1807 | 1 | 1 | 31 | 43694 | 8963 | 81 | 4864 | 5308416 | 2 |
| 722 | 174 | 4 | 2 | 2 | 2 | 236 | 9799982876790358015 | 1 | 21887 | 1 | 1 | 31 | 43694 | 15155 | 81 | 4881 | 3342385 | 2 |
| 723 | 174 | 4 | 2 | 2 | 2 | 236 | 9800018197158603424 | 1 | 1287 | 1 | 1 | 31 | 43759 | 10786 | 81 | 4353 | 1114128 | 2 |
| 724 | 174 | 4 | 2 | 2 | 2 | 236 | 9800018198601400319 | 1 | 22527 | 1 | 1 | 31 | 43694 | 2819 | 81 | 13057 | 5308416 | 2 |
| 725 | 174 | 4 | 2 | 2 | 2 | 236 | 9800053518969666288 | 1 | 1287 | 1 | 1 | 31 | 43759 | 8963 | 81 | 4864 | 5570576 | 2 |
| 726 | 174 | 4 | 2 | 2 | 2 | 236 | 9800106501124716792 | 1 | 1807 | 1 | 1 | 31 | 43694 | 13091 | 16 | 13057 | 15794176 | 2 |
| 727 | 174 | 4 | 2 | 2 | 2 | 236 | 9836049194643619839 | 1 | 3871 | 1 | 1 | 31 | 43694 | 47931 | 16 | 1792 | 7536640 | 2 |
| 728 | 174 | 4 | 2 | 2 | 2 | 236 | 9836086715476934640 | 1 | 22527 | 1 | 1 | 31 | 43694 | 8994 | 81 | 769 | 15794176 | 2 |
| 729 | 174 | 4 | 2 | 2 | 2 | 236 | 9836142995297075184 | 1 | 30591 | 1 | 1 | 31 | 43694 | 15155 | 81 | 769 | 16187443 | 2 |
| 730 | 174 | 4 | 2 | 2 | 2 | 236 | 9836142996728381424 | 1 | 2047 | 1 | 1 | 31 | 43694 | 13091 | 81 | 13057 | 5308416 | 2 |
| 731 | 174 | 4 | 2 | 2 | 2 | 248 | 11529391520379609248 | 1 | 21887 | 1 | 1 | 31 | 43759 | 10786 | 16 | 769 | 15728656 | 2 |
| 732 | 174 | 4 | 2 | 2 | 2 | 248 | 11529391521962983372 | 1 | 5461 | 1 | 1 | 31 | 43759 | 10786 | 16 | 4881 | 7667712 | 2 |
| 733 | 174 | 4 | 2 | 2 | 2 | 248 | 11529391657942296744 | 1 | 16255 | 1 | 1 | 31 | 43694 | 47931 | 16 | 1792 | 16187392 | 2 |
| 734 | 174 | 4 | 2 | 2 | 2 | 248 | 11529391658881835232 | 1 | 22527 | 1 | 1 | 31 | 43694 | 8963 | 16 | 4864 | 16711729 | 2 |
| 735 | 174 | 4 | 2 | 2 | 2 | 248 | 11529462165077549248 | 1 | 3871 | 1 | 1 | 31 | 43694 | 47931 | 81 | 4881 | 1048576 | 2 |
| 736 | 174 | 4 | 2 | 2 | 2 | 248 | 11529479962344091888 | 1 | 7999 | 1 | 1 | 31 | 43759 | 8706 | 16 | 4864 | 15794176 | 2 |
| 737 | 174 | 4 | 2 | 2 | 2 | 248 | 11565420317396478634 | 1 | 30591 | 1 | 1 | 31 | 43694 | 47931 | 16 | 22272 | 3342352 | 2 |
| 738 | 174 | 4 | 2 | 2 | 2 | 248 | 11565420317396478634 | 1 | 8191 | 1 | 1 | 31 | 43759 | 8963 | 81 | 4864 | 7536640 | 2 |
| 739 | 174 | 4 | 2 | 2 | 2 | 248 | 11565420456184967408 | 1 | 14335 | 1 | 1 | 31 | 43759 | 10786 | 16 | 769 | 7536691 | 2 |
| 740 | 174 | 4 | 2 | 2 | 2 | 248 | 11565420456184967408 | 1 | 22367 | 1 | 1 | 31 | 43694 | 15155 | 16 | 769 | 3211264 | 2 |
| 741 | 174 | 4 | 2 | 2 | 2 | 248 | 11565508553375154175 | 1 | 21887 | 1 | 1 | 31 | 43759 | 48043 | 16 | 4881 | 16187392 | 2 |
| 742 | 174 | 4 | 2 | 2 | 2 | 248 | 11565508760965218303 | 1 | 32767 | 1 | 1 | 31 | 43759 | 43522 | 16 | 4881 | 15794176 | 2 |
| 743 | 174 | 4 | 2 | 2 | 2 | 248 | 11574427654092270248 | 1 | 1807 | 1 | 1 | 31 | 43759 | 8963 | 81 | 4881 | 7536640 | 2 |
| 744 | 174 | 4 | 2 | 2 | 2 | 248 | 11574427654260566698 | 1 | 24447 | 1 | 1 | 31 | 43694 | 8963 | 16 | 13057 | 3211312 | 2 |
| 745 | 174 | 4 | 2 | 2 | 2 | 248 | 11574436450185292458 | 1 | 7999 | 1 | 1 | 31 | 43759 | 10786 | 16 | 4881 | 3342385 | 2 |
| 746 | 174 | 4 | 2 | 2 | 2 | 248 | 11574438649208548008 | 1 | 30591 | 1 | 1 | 31 | 43759 | 10786 | 16 | 769 | 5242896 | 2 |
| 747 | 174 | 4 | 2 | 2 | 2 | 248 | 11574438650488614636 | 1 | 24447 | 1 | 1 | 31 | 43759 | 8963 | 81 | 4881 | 3145744 | 2 |
| 748 | 174 | 4 | 2 | 2 | 2 | 248 | 11574498023914397920 | 1 | 8031 | 1 | 1 | 31 | 43694 | 8963 | 81 | 4881 | 15794176 | 2 |
| 749 | 174 | 4 | 2 | 2 | 2 | 248 | 11574498297882144480 | 1 | 21887 | 1 | 1 | 31 | 43759 | 15155 | 81 | 13057 | 16711729 | 2 |
| 750 | 174 | 4 | 2 | 2 | 2 | 248 | 11574511216976003071 | 1 | 3871 | 1 | 1 | 31 | 43759 | 48043 | 16 | 13057 | 15728656 | 2 |
| 751 | 174 | 4 | 2 | 2 | 2 | 248 | 11574515891500482559 | 1 | 24447 | 1 | 19 | 31 | 43694 | 47931 | 16 | 4881 | 16711696 | 2 |
| 752 | 174 | 4 | 2 | 2 | 2 | 248 | 11574515958788194300 | 1 | 2047 | 1 | 1 | 31 | 43694 | 10786 | 81 | 13057 | 3145744 | 2 |
| 753 | 174 | 4 | 2 | 2 | 2 | 248 | 11574532107831672744 | 1 | 14199 | 1 | 1 | 31 | 43694 | 10786 | 81 | 769 | 7536691 | 2 |
| 754 | 174 | 4 | 2 | 2 | 2 | 248 | 11574532108971999136 | 1 | 22391 | 1 | 1 | 31 | 43694 | 8706 | 16 | 13057 | 1114128 | 2 |

| Model No. | Functions in integer form at the nodes of the network | | | | | | | | | | | | | | | | | |
|---|---|---|---|---|---|---|---|---|---|---|---|---|---|---|---|---|---|---|
| | CK | ARR1 | SHY2 | AUXIAA | ARF | ARF10 | ARF5 | XAL1 | PLT | AUX | SCR | SHR | MIR166 | PHB | JKD | MGP | WOX5 | CLE40 |
| 755 | 174 | 4 | 2 | 2 | 2 | 248 | 12249978456032455338 | 1 | 22527 | 1 | 1 | 31 | 43759 | 47931 | 16 | 22272 | 16711696 | 2 |
| 756 | 174 | 4 | 2 | 2 | 2 | 248 | 12249978457458540480 | 1 | 5461 | 1 | 1 | 31 | 43694 | 10786 | 81 | 13057 | 1048576 | 2 |
| 757 | 174 | 4 | 2 | 2 | 2 | 248 | 12249978593471408808 | 1 | 8031 | 1 | 1 | 31 | 43759 | 48043 | 16 | 5393 | 5242896 | 2 |
| 758 | 174 | 4 | 2 | 2 | 2 | 248 | 12249978593479797408 | 1 | 16255 | 1 | 1 | 31 | 43694 | 10786 | 81 | 13057 | 1048576 | 2 |
| 759 | 174 | 4 | 2 | 2 | 2 | 248 | 12249978594612276960 | 1 | 1301 | 1 | 1 | 31 | 43759 | 13091 | 16 | 4881 | 1114128 | 2 |
| 760 | 174 | 4 | 2 | 2 | 2 | 248 | 12250049099654557388 | 1 | 7999 | 1 | 1 | 31 | 43759 | 48043 | 16 | 22272 | 16187443 | 2 |
| 761 | 174 | 4 | 2 | 2 | 2 | 248 | 12250049099654561791 | 1 | 21847 | 1 | 1 | 31 | 43759 | 47931 | 16 | 5393 | 7667712 | 2 |
| 762 | 174 | 4 | 2 | 2 | 2 | 248 | 12250049099665697514 | 1 | 16255 | 1 | 1 | 31 | 43759 | 47931 | 16 | 13057 | 7536691 | 2 |
| 763 | 174 | 4 | 2 | 2 | 2 | 248 | 12250049099665697514 | 1 | 32767 | 1 | 1 | 31 | 43759 | 13091 | 81 | 769 | 7536691 | 2 |
| 764 | 174 | 4 | 2 | 2 | 2 | 248 | 12250049100740881088 | 1 | 5631 | 1 | 1 | 31 | 43759 | 10786 | 81 | 4864 | 3211264 | 2 |
| 765 | 174 | 4 | 2 | 2 | 2 | 248 | 12250049100808777420 | 1 | 30591 | 1 | 1 | 31 | 43694 | 8706 | 16 | 4864 | 5308416 | 2 |
| 766 | 174 | 4 | 2 | 2 | 2 | 248 | 12250049101012466416 | 1 | 22367 | 1 | 1 | 31 | 43694 | 48043 | 81 | 4864 | 5242896 | 2 |
| 767 | 174 | 4 | 2 | 2 | 2 | 248 | 12250066898007948024 | 1 | 3871 | 1 | 1 | 31 | 43759 | 15155 | 16 | 4864 | 16711696 | 2 |
| 768 | 174 | 4 | 2 | 2 | 2 | 248 | 12250066899085359856 | 1 | 1807 | 1 | 1 | 31 | 43759 | 8994 | 81 | 4353 | 5308416 | 2 |
| 769 | 174 | 4 | 2 | 2 | 2 | 248 | 12286007253059807880 | 1 | 8191 | 1 | 1 | 31 | 43759 | 47931 | 81 | 13057 | 3211281 | 2 |
| 770 | 174 | 4 | 2 | 2 | 2 | 248 | 12295014590901452799 | 1 | 2047 | 1 | 1 | 31 | 43694 | 47931 | 81 | 13057 | 15728656 | 2 |
| 771 | 174 | 4 | 2 | 2 | 2 | 248 | 12295108391841890300 | 1 | 22527 | 1 | 1 | 31 | 43759 | 13091 | 16 | 13057 | 5308433 | 2 |
| 772 | 174 | 4 | 2 | 2 | 2 | 248 | 13835269714491785408 | 1 | 3871 | 1 | 1 | 31 | 43759 | 47931 | 81 | 4881 | 16187443 | 2 |
| 773 | 174 | 4 | 2 | 2 | 2 | 248 | 13835269714491785408 | 1 | 3967 | 1 | 1 | 31 | 43759 | 47931 | 81 | 4864 | 16187443 | 2 |
| 774 | 174 | 4 | 2 | 2 | 2 | 248 | 13835269715548766122 | 1 | 14143 | 1 | 1 | 31 | 43759 | 48043 | 81 | 4353 | 1048576 | 2 |
| 775 | 174 | 4 | 2 | 2 | 2 | 248 | 13835269989512825032 | 1 | 22527 | 1 | 1 | 31 | 43759 | 15155 | 16 | 4864 | 5570576 | 2 |
| 776 | 174 | 4 | 2 | 2 | 2 | 248 | 13871298511519141000 | 1 | 7999 | 1 | 1 | 31 | 43694 | 8963 | 81 | 769 | 15794176 | 2 |
| 777 | 174 | 4 | 2 | 2 | 2 | 248 | 13871298787209703664 | 1 | 21887 | 1 | 1 | 31 | 43759 | 13091 | 81 | 4881 | 7340032 | 2 |
| 778 | 174 | 4 | 2 | 2 | 2 | 248 | 13871298787462348799 | 1 | 8191 | 1 | 1 | 31 | 43694 | 48043 | 16 | 14080 | 7667712 | 2 |
| 779 | 174 | 4 | 2 | 2 | 2 | 248 | 13871351425718091690 | 1 | 3871 | 1 | 1 | 31 | 43694 | 47931 | 16 | 22272 | 15794176 | 2 |
| 780 | 174 | 4 | 2 | 2 | 2 | 248 | 13871351769315475455 | 1 | 7999 | 1 | 1 | 31 | 43759 | 47931 | 81 | 4881 | 7536640 | 2 |
| 781 | 174 | 4 | 2 | 2 | 2 | 248 | 13889321981003746508 | 1 | 16255 | 1 | 1 | 31 | 43694 | 10786 | 81 | 13057 | 7536640 | 2 |
| 782 | 174 | 4 | 2 | 2 | 2 | 248 | 13889321981137963200 | 1 | 8191 | 1 | 1 | 31 | 43759 | 13091 | 16 | 13057 | 7667712 | 2 |
| 783 | 174 | 4 | 2 | 2 | 2 | 248 | 13889326379050257608 | 1 | 32767 | 1 | 1 | 31 | 43759 | 10786 | 16 | 13057 | 3342385 | 2 |
| 784 | 174 | 4 | 2 | 2 | 2 | 248 | 13889348370088128704 | 1 | 21887 | 1 | 1 | 31 | 43694 | 10786 | 81 | 4881 | 15794176 | 2 |
| 785 | 174 | 4 | 2 | 2 | 2 | 248 | 13889348507778744288 | 1 | 32767 | 1 | 1 | 31 | 43694 | 8706 | 16 | 4881 | 16711696 | 2 |
| 786 | 174 | 4 | 2 | 2 | 2 | 248 | 13889359365103741632 | 1 | 2047 | 1 | 1 | 31 | 43694 | 8963 | 16 | 769 | 3342352 | 2 |
| 787 | 174 | 4 | 2 | 2 | 2 | 248 | 13889359365106494186 | 1 | 8031 | 1 | 1 | 31 | 43759 | 15155 | 16 | 13057 | 3211264 | 2 |
| 788 | 174 | 4 | 2 | 2 | 2 | 248 | 13889365961603611336 | 1 | 22527 | 1 | 1 | 31 | 43759 | 13091 | 81 | 13057 | 3342385 | 2 |
| 789 | 174 | 4 | 2 | 2 | 2 | 248 | 13889366168331942640 | 1 | 3967 | 1 | 1 | 31 | 43759 | 15155 | 16 | 13057 | 5242896 | 2 |
| 790 | 174 | 4 | 2 | 2 | 2 | 248 | 141290096339584 | 1 | 30591 | 1 | 1 | 31 | 43759 | 15155 | 81 | 769 | 16711696 | 2 |
| 791 | 174 | 4 | 2 | 2 | 2 | 248 | 141290465443968 | 1 | 8031 | 1 | 1 | 31 | 43694 | 8994 | 16 | 4881 | 5308433 | 2 |
| 792 | 174 | 4 | 2 | 2 | 2 | 248 | 141291522424704 | 1 | 5461 | 1 | 1 | 31 | 43694 | 10786 | 16 | 769 | 16711696 | 2 |
| 793 | 174 | 4 | 2 | 2 | 2 | 248 | 141291522424704 | 1 | 7999 | 1 | 1 | 31 | 43694 | 15155 | 16 | 13057 | 5308433 | 2 |
| 794 | 174 | 4 | 2 | 2 | 2 | 248 | 14465782733441400831 | 1 | 8191 | 1 | 1 | 31 | 43759 | 15155 | 16 | 22272 | 5570576 | 2 |
| 795 | 174 | 4 | 2 | 2 | 2 | 248 | 14465787131488423116 | 1 | 32767 | 1 | 1 | 31 | 43694 | 47931 | 16 | 13057 | 15794176 | 2 |
| 796 | 174 | 4 | 2 | 2 | 2 | 248 | 14699974037287783560 | 1 | 5631 | 1 | 1 | 31 | 43694 | 10786 | 16 | 4864 | 7340080 | 2 |
| 797 | 174 | 4 | 2 | 2 | 2 | 248 | 14699974312165690568 | 1 | 22367 | 1 | 1 | 31 | 43694 | 8994 | 81 | 1281 | 7340032 | 2 |
| 798 | 174 | 4 | 2 | 2 | 2 | 248 | 14700009359098834602 | 1 | 2047 | 1 | 1 | 31 | 43759 | 47931 | 16 | 21761 | 3145744 | 2 |
| 799 | 174 | 4 | 2 | 2 | 2 | 248 | 14700009359098834602 | 1 | 24447 | 1 | 1 | 31 | 43694 | 13091 | 81 | 4881 | 1048576 | 2 |

| Model No. | Functions in integer form at the nodes of the network | | | | | | | | | | | | | | | | | |
|---|---|---|---|---|---|---|---|---|---|---|---|---|---|---|---|---|---|---|
| | CK | ARR1 | SHY2 | AUXIAA | ARF | ARF10 | ARF5 | XAL1 | PLT | AUX | SCR | SHR | MIR166 | PHB | JKD | MGP | WOX5 | CLE40 |
| 800 | 174 | 4 | 2 | 2 | 2 | 248 | 14700009359098834602 | 1 | 5461 | 1 | 1 | 31 | 43694 | 10786 | 81 | 4881 | 7667712 | 2 |
| 801 | 174 | 4 | 2 | 2 | 2 | 248 | 14700009359098838256 | 1 | 1807 | 1 | 1 | 31 | 43694 | 48043 | 81 | 4864 | 16711696 | 2 |
| 802 | 174 | 4 | 2 | 2 | 2 | 248 | 14700009359112203500 | 1 | 8031 | 1 | 1 | 31 | 43759 | 15155 | 16 | 769 | 3211281 | 2 |
| 803 | 174 | 4 | 2 | 2 | 2 | 248 | 14700009359971188735 | 1 | 32767 | 1 | 1 | 31 | 43694 | 10786 | 16 | 13057 | 1048576 | 2 |
| 804 | 174 | 4 | 2 | 2 | 2 | 248 | 14700027294882266874 | 1 | 3871 | 1 | 1 | 31 | 43759 | 8706 | 16 | 4864 | 16711729 | 2 |
| 805 | 174 | 4 | 2 | 2 | 2 | 248 | 14736002834315136192 | 1 | 14143 | 1 | 1 | 31 | 43694 | 15155 | 16 | 4881 | 16187443 | 2 |
| 806 | 174 | 4 | 2 | 2 | 2 | 248 | 14736003109193567436 | 1 | 3871 | 1 | 1 | 31 | 43759 | 10786 | 81 | 4881 | 5570576 | 2 |
| 807 | 174 | 4 | 2 | 2 | 2 | 248 | 14736003109197237440 | 1 | 22367 | 1 | 1 | 31 | 43694 | 47931 | 16 | 4864 | 5308416 | 2 |
| 808 | 174 | 4 | 2 | 2 | 2 | 248 | 14736059046851313648 | 1 | 2047 | 1 | 1 | 31 | 43694 | 8994 | 16 | 769 | 15728656 | 2 |
| 809 | 174 | 4 | 2 | 2 | 2 | 248 | 14736059046852100095 | 1 | 32767 | 1 | 1 | 31 | 43694 | 13091 | 81 | 13057 | 7340080 | 2 |
| 810 | 174 | 4 | 2 | 2 | 2 | 248 | 14754055028541812462 | 1 | 32767 | 1 | 1 | 31 | 43759 | 8706 | 16 | 13057 | 1048576 | 2 |
| 811 | 174 | 4 | 2 | 2 | 2 | 248 | 14754055029113548512 | 1 | 5631 | 1 | 1 | 31 | 43759 | 15155 | 81 | 4881 | 7536640 | 2 |
| 812 | 174 | 4 | 2 | 2 | 2 | 248 | 14754073789530701808 | 1 | 8031 | 1 | 1 | 31 | 43759 | 10786 | 16 | 769 | 3211281 | 2 |
| 813 | 174 | 4 | 2 | 2 | 2 | 248 | 150085627316352 | 1 | 14143 | 1 | 1 | 31 | 43694 | 10786 | 16 | 4864 | 1048576 | 2 |
| 814 | 174 | 4 | 2 | 2 | 2 | 248 | 150085627316352 | 1 | 14143 | 1 | 1 | 31 | 43694 | 10786 | 16 | 769 | 1048576 | 2 |
| 815 | 174 | 4 | 2 | 2 | 2 | 248 | 150085627316352 | 1 | 14143 | 1 | 1 | 31 | 43759 | 10786 | 81 | 4881 | 1048576 | 2 |
| 816 | 174 | 4 | 2 | 2 | 2 | 248 | 150085627316352 | 1 | 16255 | 1 | 1 | 31 | 43694 | 10786 | 81 | 13057 | 1048576 | 2 |
| 817 | 174 | 4 | 2 | 2 | 2 | 248 | 150085627316352 | 1 | 1807 | 1 | 1 | 31 | 43694 | 10786 | 16 | 4881 | 1048576 | 2 |
| 818 | 174 | 4 | 2 | 2 | 2 | 248 | 150085627316352 | 1 | 2047 | 1 | 1 | 31 | 43694 | 10786 | 16 | 4881 | 1048576 | 2 |
| 819 | 174 | 4 | 2 | 2 | 2 | 248 | 150085627316352 | 1 | 21887 | 1 | 1 | 31 | 43759 | 10786 | 16 | 4864 | 1048576 | 2 |
| 820 | 174 | 4 | 2 | 2 | 2 | 248 | 150085627316352 | 1 | 21887 | 1 | 1 | 31 | 43759 | 10786 | 81 | 4864 | 1048576 | 2 |
| 821 | 174 | 4 | 2 | 2 | 2 | 248 | 150085627316352 | 1 | 22367 | 1 | 1 | 31 | 43694 | 10786 | 81 | 4881 | 1048576 | 2 |
| 822 | 174 | 4 | 2 | 2 | 2 | 248 | 150085627316352 | 1 | 22367 | 1 | 1 | 31 | 43694 | 8706 | 16 | 4881 | 1048576 | 2 |
| 823 | 174 | 4 | 2 | 2 | 2 | 248 | 150085627316352 | 1 | 22527 | 1 | 1 | 31 | 43759 | 8706 | 16 | 4864 | 16187443 | 2 |
| 824 | 174 | 4 | 2 | 2 | 2 | 248 | 150085627316352 | 1 | 24447 | 1 | 1 | 31 | 43694 | 10786 | 16 | 769 | 1048576 | 2 |
| 825 | 174 | 4 | 2 | 2 | 2 | 248 | 150085627316352 | 1 | 24447 | 1 | 1 | 31 | 43694 | 10786 | 81 | 769 | 1048576 | 2 |
| 826 | 174 | 4 | 2 | 2 | 2 | 248 | 150085627316352 | 1 | 24447 | 1 | 1 | 31 | 43694 | 15155 | 16 | 4881 | 1048576 | 2 |
| 827 | 174 | 4 | 2 | 2 | 2 | 248 | 150085627316352 | 1 | 32767 | 1 | 1 | 31 | 43694 | 10786 | 16 | 13057 | 1048576 | 2 |
| 828 | 174 | 4 | 2 | 2 | 2 | 248 | 150085627316352 | 1 | 32767 | 1 | 1 | 31 | 43694 | 10786 | 81 | 13057 | 1048576 | 2 |
| 829 | 174 | 4 | 2 | 2 | 2 | 248 | 150085627316352 | 1 | 32767 | 1 | 1 | 31 | 43694 | 10786 | 81 | 769 | 1048576 | 2 |
| 830 | 174 | 4 | 2 | 2 | 2 | 248 | 150085627316352 | 1 | 32767 | 1 | 1 | 31 | 43694 | 47931 | 81 | 4881 | 1048576 | 2 |
| 831 | 174 | 4 | 2 | 2 | 2 | 248 | 150085627316352 | 1 | 32767 | 1 | 1 | 31 | 43694 | 8706 | 16 | 769 | 1048576 | 2 |
| 832 | 174 | 4 | 2 | 2 | 2 | 248 | 150085627316352 | 1 | 32767 | 1 | 1 | 31 | 43694 | 8963 | 81 | 4881 | 1048576 | 2 |
| 833 | 174 | 4 | 2 | 2 | 2 | 248 | 150085627316352 | 1 | 32767 | 1 | 1 | 31 | 43694 | 8994 | 81 | 4864 | 16711729 | 2 |
| 834 | 174 | 4 | 2 | 2 | 2 | 248 | 150085627316352 | 1 | 32767 | 1 | 1 | 31 | 43759 | 10786 | 16 | 13057 | 1048576 | 2 |
| 835 | 174 | 4 | 2 | 2 | 2 | 248 | 150085627316352 | 1 | 32767 | 1 | 1 | 31 | 43759 | 10786 | 16 | 769 | 1048576 | 2 |
| 836 | 174 | 4 | 2 | 2 | 2 | 248 | 150085627316352 | 1 | 32767 | 1 | 1 | 31 | 43759 | 10786 | 81 | 13057 | 1048576 | 2 |
| 837 | 174 | 4 | 2 | 2 | 2 | 248 | 150085627316352 | 1 | 32767 | 1 | 1 | 31 | 43759 | 47931 | 81 | 4864 | 1048576 | 2 |
| 838 | 174 | 4 | 2 | 2 | 2 | 248 | 150085627316352 | 1 | 3967 | 1 | 1 | 31 | 43759 | 48043 | 81 | 4881 | 16187443 | 2 |
| 839 | 174 | 4 | 2 | 2 | 2 | 248 | 150085627316352 | 1 | 7999 | 1 | 1 | 31 | 43694 | 10786 | 16 | 4881 | 1048576 | 2 |
| 840 | 174 | 4 | 2 | 2 | 2 | 248 | 150085627316352 | 1 | 8031 | 1 | 1 | 31 | 43694 | 10786 | 81 | 769 | 1048576 | 2 |
| 841 | 174 | 4 | 2 | 2 | 2 | 248 | 150085627316352 | 1 | 8031 | 1 | 1 | 31 | 43694 | 15155 | 81 | 4353 | 7667712 | 2 |
| 842 | 174 | 4 | 2 | 2 | 2 | 248 | 150085627316352 | 1 | 8031 | 1 | 1 | 31 | 43759 | 15155 | 16 | 4881 | 1048576 | 2 |
| 843 | 174 | 4 | 2 | 2 | 2 | 248 | 150085627316352 | 1 | 8191 | 1 | 1 | 31 | 43694 | 10786 | 81 | 4881 | 1048576 | 2 |
| 844 | 174 | 4 | 2 | 2 | 2 | 248 | 150085627316352 | 1 | 8191 | 1 | 1 | 31 | 43694 | 10786 | 81 | 769 | 1048576 | 2 |

| Model No. | Functions in integer form at the nodes of the network | | | | | | | | | | | | | | | | | |
|---|---|---|---|---|---|---|---|---|---|---|---|---|---|---|---|---|---|---|
| | CK | ARR1 | SHY2 | AUXIAA | ARF | ARF10 | ARF5 | XAL1 | PLT | AUX | SCR | SHR | MIR166 | PHB | JKD | MGP | WOX5 | CLE40 |
| 845 | 174 | 4 | 2 | 2 | 2 | 248 | 150085627316352 | 1 | 8191 | 1 | 1 | 31 | 43759 | 10786 | 16 | 4864 | 1048576 | 2 |
| 846 | 174 | 4 | 2 | 2 | 2 | 248 | 150085627316352 | 1 | 8191 | 1 | 1 | 31 | 43759 | 10786 | 16 | 769 | 1048576 | 2 |
| 847 | 174 | 4 | 2 | 2 | 2 | 248 | 16141148321166192366 | 1 | 14335 | 1 | 1 | 31 | 43694 | 15155 | 16 | 4864 | 7536691 | 2 |
| 848 | 174 | 4 | 2 | 2 | 2 | 248 | 16141148321166192366 | 1 | 22527 | 1 | 1 | 31 | 43759 | 2819 | 81 | 4881 | 15728656 | 2 |
| 849 | 174 | 4 | 2 | 2 | 2 | 248 | 16141148321181789934 | 1 | 3871 | 1 | 1 | 31 | 43759 | 47931 | 16 | 21761 | 7536640 | 2 |
| 850 | 174 | 4 | 2 | 2 | 2 | 248 | 16141148321215475952 | 1 | 14143 | 1 | 1 | 31 | 43759 | 15155 | 81 | 4864 | 5308433 | 2 |
| 851 | 174 | 4 | 2 | 2 | 2 | 248 | 16195191379033391103 | 1 | 16255 | 1 | 1 | 31 | 43759 | 13091 | 81 | 13057 | 7667712 | 2 |
| 852 | 174 | 4 | 2 | 2 | 2 | 248 | 16195208971743657983 | 1 | 32767 | 1 | 1 | 31 | 43759 | 47931 | 16 | 22272 | 16187392 | 2 |
| 853 | 174 | 4 | 2 | 2 | 2 | 248 | 16195209177379959024 | 1 | 8191 | 1 | 1 | 31 | 43759 | 10786 | 16 | 769 | 5308433 | 2 |
| 854 | 174 | 4 | 2 | 2 | 2 | 248 | 16204214109414096895 | 1 | 3871 | 1 | 1 | 31 | 43694 | 48043 | 81 | 4881 | 7536640 | 2 |
| 855 | 174 | 4 | 2 | 2 | 2 | 248 | 16204232800825442286 | 1 | 5631 | 1 | 1 | 31 | 43694 | 48043 | 81 | 769 | 7667712 | 2 |
| 856 | 174 | 4 | 2 | 2 | 2 | 248 | 16915778314696714986 | 1 | 8191 | 1 | 1 | 31 | 43759 | 10786 | 16 | 13057 | 3211264 | 2 |
| 857 | 174 | 4 | 2 | 2 | 2 | 248 | 16915796113043946234 | 1 | 22527 | 1 | 1 | 31 | 43694 | 15155 | 16 | 13057 | 7667712 | 2 |
| 858 | 174 | 4 | 2 | 2 | 2 | 248 | 16915796113312774896 | 1 | 24447 | 1 | 1 | 31 | 43759 | 8963 | 81 | 4881 | 15728656 | 2 |
| 859 | 174 | 4 | 2 | 2 | 2 | 248 | 16915796113393516528 | 1 | 22367 | 1 | 1 | 31 | 43759 | 10786 | 81 | 769 | 3211264 | 2 |
| 860 | 174 | 4 | 2 | 2 | 2 | 248 | 17050909256179515391 | 1 | 8191 | 1 | 1 | 31 | 43759 | 15155 | 16 | 1792 | 5308416 | 2 |
| 861 | 174 | 4 | 2 | 2 | 2 | 248 | 17149970031161901055 | 1 | 14143 | 1 | 1 | 31 | 43694 | 15155 | 81 | 4864 | 3342352 | 2 |
| 862 | 174 | 4 | 2 | 2 | 2 | 248 | 17149987691781095166 | 1 | 14335 | 1 | 1 | 31 | 43759 | 10786 | 81 | 4881 | 7536691 | 2 |
| 863 | 174 | 4 | 2 | 2 | 2 | 248 | 17294087005675778240 | 1 | 3967 | 1 | 1 | 31 | 43694 | 48043 | 16 | 4864 | 5242896 | 2 |
| 864 | 174 | 4 | 2 | 2 | 2 | 248 | 17294087005688361152 | 1 | 8031 | 1 | 1 | 31 | 43759 | 8706 | 16 | 13057 | 1114128 | 2 |
| 865 | 174 | 4 | 2 | 2 | 2 | 248 | 17294087005944152063 | 1 | 21847 | 1 | 1 | 31 | 43694 | 8963 | 81 | 1281 | 15794176 | 2 |
| 866 | 174 | 4 | 2 | 2 | 2 | 248 | 17294087143114731744 | 1 | 16255 | 1 | 1 | 31 | 43759 | 2562 | 16 | 4864 | 7340032 | 2 |
| 867 | 174 | 4 | 2 | 2 | 2 | 248 | 17294087143127314656 | 1 | 1807 | 1 | 1 | 31 | 43694 | 15155 | 16 | 4864 | 16187443 | 2 |
| 868 | 174 | 4 | 2 | 2 | 2 | 248 | 17294087280553685216 | 1 | 14143 | 1 | 1 | 31 | 43759 | 13091 | 16 | 4864 | 3342352 | 2 |
| 869 | 174 | 4 | 2 | 2 | 2 | 248 | 17294087418007318752 | 1 | 32767 | 1 | 1 | 31 | 43759 | 10786 | 81 | 13057 | 3342385 | 2 |
| 870 | 174 | 4 | 2 | 2 | 2 | 248 | 17330115802703130816 | 1 | 3967 | 1 | 1 | 31 | 43694 | 10786 | 16 | 769 | 1048576 | 2 |
| 871 | 174 | 4 | 2 | 2 | 2 | 248 | 17330115940144181408 | 1 | 3967 | 1 | 1 | 31 | 43694 | 10786 | 81 | 13057 | 1048576 | 2 |
| 872 | 174 | 4 | 2 | 2 | 2 | 248 | 17330116077841022975 | 1 | 22367 | 1 | 1 | 31 | 43694 | 8963 | 81 | 4881 | 7536640 | 2 |
| 873 | 174 | 4 | 2 | 2 | 2 | 248 | 17348130613536813296 | 1 | 16255 | 1 | 1 | 31 | 43694 | 10786 | 81 | 4881 | 16711696 | 2 |
| 874 | 174 | 4 | 2 | 2 | 2 | 248 | 17361641481306701823 | 1 | 8031 | 1 | 1 | 31 | 43759 | 13091 | 81 | 13057 | 3342352 | 2 |
| 875 | 174 | 4 | 2 | 2 | 2 | 248 | 17361641481373286142 | 1 | 21847 | 1 | 1 | 31 | 43759 | 10786 | 16 | 13057 | 5308416 | 2 |
| 876 | 174 | 4 | 2 | 2 | 2 | 248 | 17361641481374203902 | 1 | 16255 | 1 | 1 | 31 | 43759 | 2819 | 81 | 769 | 7667712 | 2 |
| 877 | 174 | 4 | 2 | 2 | 2 | 248 | 17361641481374203902 | 1 | 21887 | 1 | 1 | 31 | 43759 | 15155 | 16 | 4881 | 7667712 | 2 |
| 878 | 174 | 4 | 2 | 2 | 2 | 248 | 17361650277231425784 | 1 | 30591 | 1 | 1 | 31 | 43759 | 48043 | 16 | 65281 | 7340032 | 2 |
| 879 | 174 | 4 | 2 | 2 | 2 | 248 | 176471616299168 | 1 | 16255 | 1 | 1 | 31 | 43694 | 15155 | 81 | 4864 | 15728656 | 2 |
| 880 | 174 | 4 | 2 | 2 | 2 | 248 | 176474309042304 | 1 | 22527 | 1 | 1 | 31 | 43759 | 8994 | 81 | 13057 | 16187443 | 2 |
| 881 | 174 | 4 | 2 | 2 | 2 | 248 | 176611747995808 | 1 | 14199 | 1 | 1 | 31 | 43759 | 10786 | 16 | 4881 | 7536691 | 2 |
| 882 | 174 | 4 | 2 | 2 | 2 | 248 | 176612276494496 | 1 | 14335 | 1 | 1 | 31 | 43694 | 15155 | 81 | 769 | 5570576 | 2 |
| 883 | 174 | 4 | 2 | 2 | 2 | 248 | 17938111029669068784 | 1 | 30591 | 1 | 1 | 31 | 43759 | 8994 | 81 | 4881 | 15794176 | 2 |
| 884 | 174 | 4 | 2 | 2 | 2 | 248 | 17938111029669068799 | 1 | 24447 | 1 | 1 | 31 | 43759 | 8963 | 81 | 769 | 7536640 | 2 |
| 885 | 174 | 4 | 2 | 2 | 2 | 248 | 17938115427749789695 | 1 | 7999 | 1 | 1 | 31 | 43759 | 10786 | 81 | 13057 | 3342352 | 2 |
| 886 | 174 | 4 | 2 | 2 | 2 | 248 | 17938118726284017648 | 1 | 22367 | 1 | 1 | 31 | 43759 | 8963 | 16 | 4881 | 5570576 | 2 |
| 887 | 174 | 4 | 2 | 2 | 2 | 248 | 17938118726368886783 | 1 | 32767 | 1 | 1 | 31 | 43694 | 48043 | 16 | 4881 | 15728656 | 2 |
| 888 | 174 | 4 | 2 | 2 | 2 | 248 | 18082232814932131824 | 1 | 22527 | 1 | 1 | 31 | 43694 | 15155 | 81 | 13057 | 3211264 | 2 |
| 889 | 174 | 4 | 2 | 2 | 2 | 248 | 18370463191067066367 | 1 | 8031 | 1 | 1 | 31 | 43694 | 13091 | 16 | 4881 | 3211264 | 2 |

| Model No. | Functions in integer form at the nodes of the network | | | | | | | | | | | | | | | | | |
|---|---|---|---|---|---|---|---|---|---|---|---|---|---|---|---|---|---|---|
| | CK | ARR1 | SHY2 | AUXIAA | ARF | ARF10 | ARF5 | XAL1 | PLT | AUX | SCR | SHR | MIR166 | PHB | JKD | MGP | WOX5 | CLE40 |
| 890 | 174 | 4 | 2 | 2 | 2 | 248 | 18374967409170775944 | 1 | 14335 | 1 | 1 | 31 | 43694 | 8963 | 81 | 5393 | 5242896 | 2 |
| 891 | 174 | 4 | 2 | 2 | 2 | 248 | 18374967409170775978 | 1 | 3967 | 1 | 1 | 31 | 43694 | 8963 | 16 | 5393 | 5242896 | 2 |
| 892 | 174 | 4 | 2 | 2 | 2 | 248 | 18374967409170776012 | 1 | 21887 | 1 | 1 | 31 | 43759 | 13091 | 16 | 21761 | 7340032 | 2 |
| 893 | 174 | 4 | 2 | 2 | 2 | 248 | 18374967409170776063 | 1 | 14199 | 1 | 1 | 31 | 43759 | 48043 | 81 | 769 | 3342352 | 2 |
| 894 | 174 | 4 | 2 | 2 | 2 | 248 | 18374967409170776063 | 1 | 22527 | 1 | 1 | 31 | 43694 | 13091 | 16 | 22272 | 5242896 | 2 |
| 895 | 174 | 4 | 2 | 2 | 2 | 248 | 18374967409179688840 | 1 | 1807 | 1 | 1 | 31 | 43694 | 15155 | 16 | 4353 | 15794176 | 2 |
| 896 | 174 | 4 | 2 | 2 | 2 | 248 | 18374967409181917098 | 1 | 24447 | 1 | 1 | 31 | 43694 | 8994 | 81 | 4864 | 15794176 | 2 |
| 897 | 174 | 4 | 2 | 2 | 2 | 248 | 18374967546609729516 | 1 | 8031 | 1 | 1 | 31 | 43759 | 15155 | 16 | 13057 | 1048576 | 2 |
| 898 | 174 | 4 | 2 | 2 | 2 | 248 | 18374967684048682952 | 1 | 1301 | 1 | 1 | 31 | 43694 | 13091 | 81 | 4353 | 5308433 | 2 |
| 899 | 174 | 4 | 2 | 2 | 2 | 248 | 18374967684048682952 | 1 | 22527 | 1 | 1 | 31 | 43694 | 15155 | 16 | 13057 | 3342385 | 2 |
| 900 | 174 | 4 | 2 | 2 | 2 | 248 | 18374967684048682952 | 1 | 3967 | 1 | 1 | 31 | 43694 | 13091 | 16 | 13057 | 15728656 | 2 |
| 901 | 174 | 4 | 2 | 2 | 2 | 248 | 18374967684048682986 | 1 | 8031 | 1 | 1 | 31 | 43759 | 13091 | 16 | 1792 | 3211312 | 2 |
| 902 | 174 | 4 | 2 | 2 | 2 | 248 | 18374967684057071552 | 1 | 8191 | 1 | 1 | 31 | 43759 | 8706 | 16 | 13057 | 3342385 | 2 |
| 903 | 174 | 4 | 2 | 2 | 2 | 248 | 18374967821487636462 | 1 | 32767 | 1 | 1 | 31 | 43759 | 43522 | 81 | 769 | 1048576 | 2 |
| 904 | 174 | 4 | 2 | 2 | 2 | 248 | 18374967821487636464 | 1 | 16255 | 1 | 1 | 31 | 43759 | 13091 | 16 | 4864 | 5242896 | 2 |
| 905 | 174 | 4 | 2 | 2 | 2 | 248 | 18374967821487636479 | 1 | 5461 | 1 | 1 | 31 | 43694 | 10786 | 81 | 5393 | 16711696 | 2 |
| 906 | 174 | 4 | 2 | 2 | 2 | 248 | 18374967821503234030 | 1 | 16255 | 1 | 1 | 31 | 43694 | 8963 | 16 | 22272 | 5308433 | 2 |
| 907 | 174 | 4 | 2 | 2 | 2 | 248 | 18374967890222710782 | 1 | 21887 | 1 | 1 | 31 | 43694 | 8706 | 16 | 13057 | 5308416 | 2 |
| 908 | 174 | 4 | 2 | 2 | 2 | 248 | 18410996206198128576 | 1 | 5631 | 1 | 1 | 31 | 43759 | 10786 | 81 | 4864 | 3145744 | 2 |
| 909 | 174 | 4 | 2 | 2 | 2 | 248 | 18410996206198128588 | 1 | 2047 | 1 | 1 | 31 | 43759 | 8963 | 81 | 4881 | 16187392 | 2 |
| 910 | 174 | 4 | 2 | 2 | 2 | 248 | 18410996206198128624 | 1 | 1807 | 1 | 1 | 31 | 43759 | 48043 | 81 | 4864 | 3145744 | 2 |
| 911 | 174 | 4 | 2 | 2 | 2 | 248 | 18410996343642062847 | 1 | 14143 | 1 | 1 | 31 | 43759 | 48043 | 16 | 769 | 7667712 | 2 |
| 912 | 174 | 4 | 2 | 2 | 2 | 248 | 18410996343644422128 | 1 | 3967 | 1 | 1 | 31 | 43759 | 47931 | 16 | 4864 | 1048576 | 2 |
| 913 | 174 | 4 | 2 | 2 | 2 | 248 | 18410996481084358655 | 1 | 22527 | 1 | 1 | 31 | 43694 | 47931 | 81 | 4881 | 5308416 | 2 |
| 914 | 174 | 4 | 2 | 2 | 2 | 248 | 18410996687241805808 | 1 | 22391 | 1 | 1 | 31 | 43759 | 10786 | 16 | 13057 | 15794176 | 2 |
| 915 | 174 | 4 | 2 | 2 | 2 | 248 | 18429011017028665328 | 1 | 16255 | 1 | 1 | 31 | 43694 | 47931 | 16 | 13057 | 3211281 | 2 |
| 916 | 174 | 4 | 2 | 2 | 2 | 248 | 18429011017028665328 | 1 | 21887 | 1 | 1 | 31 | 43759 | 13091 | 16 | 4881 | 5570576 | 2 |
| 917 | 174 | 4 | 2 | 2 | 2 | 248 | 18429011085748666362 | 1 | 21887 | 1 | 1 | 31 | 43694 | 47931 | 16 | 13057 | 3145744 | 2 |
| 918 | 174 | 4 | 2 | 2 | 2 | 248 | 185409443201023 | 1 | 1287 | 1 | 1 | 31 | 43759 | 15155 | 81 | 769 | 7340032 | 2 |
| 919 | 174 | 4 | 2 | 2 | 2 | 248 | 185409443201023 | 1 | 16255 | 1 | 1 | 31 | 43694 | 15155 | 16 | 4864 | 3145744 | 2 |
| 920 | 174 | 4 | 2 | 2 | 2 | 248 | 187466732579464 | 1 | 32767 | 1 | 21 | 31 | 43759 | 770 | 81 | 13057 | 3342385 | 2 |
| 921 | 174 | 4 | 2 | 2 | 2 | 248 | 187607607471852 | 1 | 14335 | 1 | 1 | 31 | 43694 | 8963 | 81 | 4881 | 5308433 | 2 |
| 922 | 174 | 4 | 2 | 2 | 2 | 248 | 211930866313920 | 1 | 14143 | 1 | 1 | 31 | 43759 | 10786 | 16 | 4864 | 5308416 | 2 |
| 923 | 174 | 4 | 2 | 2 | 2 | 248 | 211934297771208 | 1 | 14335 | 1 | 1 | 31 | 43759 | 8963 | 81 | 769 | 15728656 | 2 |
| 924 | 174 | 4 | 2 | 2 | 2 | 248 | 211934297771208 | 1 | 2047 | 1 | 1 | 31 | 43694 | 10786 | 16 | 769 | 16187392 | 2 |
| 925 | 174 | 4 | 2 | 2 | 2 | 248 | 211934903333088 | 1 | 3967 | 1 | 1 | 31 | 43694 | 8963 | 81 | 13057 | 16711729 | 2 |
| 926 | 174 | 4 | 2 | 2 | 2 | 248 | 211935155650538 | 1 | 14143 | 1 | 1 | 31 | 43759 | 8706 | 16 | 1281 | 7536640 | 2 |
| 927 | 174 | 4 | 2 | 2 | 2 | 248 | 220726959328448 | 1 | 14199 | 1 | 1 | 31 | 43759 | 10786 | 16 | 4353 | 5308416 | 2 |
| 928 | 174 | 4 | 2 | 2 | 2 | 248 | 224850127932620 | 1 | 32767 | 1 | 1 | 31 | 43694 | 8963 | 81 | 769 | 7667712 | 2 |
| 929 | 174 | 4 | 2 | 2 | 2 | 248 | 224850127945632 | 1 | 14335 | 1 | 1 | 31 | 43759 | 2819 | 81 | 4353 | 5308416 | 2 |
| 930 | 174 | 4 | 2 | 2 | 2 | 248 | 224850127945632 | 1 | 16255 | 1 | 1 | 31 | 43694 | 8963 | 81 | 4353 | 15794176 | 2 |
| 931 | 174 | 4 | 2 | 2 | 2 | 248 | 224850127945632 | 1 | 32767 | 1 | 1 | 31 | 43759 | 47931 | 16 | 4864 | 15794176 | 2 |
| 932 | 174 | 4 | 2 | 2 | 2 | 248 | 224850127945632 | 1 | 3967 | 1 | 1 | 31 | 43759 | 8963 | 81 | 4353 | 15728656 | 2 |
| 933 | 174 | 4 | 2 | 2 | 2 | 248 | 224854422847487 | 1 | 30591 | 1 | 1 | 31 | 43694 | 47931 | 16 | 22272 | 16187443 | 2 |
| 934 | 174 | 4 | 2 | 2 | 2 | 248 | 247119533309951 | 1 | 8191 | 1 | 1 | 31 | 43694 | 13091 | 16 | 1792 | 7340032 | 2 |

| Model No. | Functions in integer form at the nodes of the network | | | | | | | | | | | | | | | | | |
|---|---|---|---|---|---|---|---|---|---|---|---|---|---|---|---|---|---|---|
| | CK | ARR1 | SHY2 | AUXIAA | ARF | ARF10 | ARF5 | XAL1 | PLT | AUX | SCR | SHR | MIR166 | PHB | JKD | MGP | WOX5 | CLE40 |
| 935 | 174 | 4 | 2 | 2 | 2 | 248 | 247256670334688 | 1 | 21847 | 1 | 1 | 31 | 43759 | 48043 | 16 | 4881 | 16187443 | 2 |
| 936 | 174 | 4 | 2 | 2 | 2 | 248 | 247256670334688 | 1 | 21887 | 1 | 1 | 31 | 43759 | 48043 | 16 | 4864 | 15794176 | 2 |
| 937 | 174 | 4 | 2 | 2 | 2 | 248 | 258114649587711 | 1 | 8191 | 1 | 1 | 31 | 43759 | 8963 | 16 | 14080 | 15794176 | 2 |
| 938 | 174 | 4 | 2 | 2 | 2 | 248 | 262650134069232 | 1 | 16255 | 1 | 1 | 31 | 43694 | 8963 | 81 | 4881 | 16187443 | 2 |
| 939 | 174 | 4 | 2 | 2 | 2 | 248 | 264432546545663 | 1 | 5631 | 1 | 1 | 31 | 43759 | 10786 | 81 | 4864 | 1048576 | 2 |
| 940 | 174 | 4 | 2 | 2 | 2 | 248 | 264436581462144 | 1 | 32767 | 1 | 1 | 31 | 43759 | 13091 | 81 | 4864 | 5308433 | 2 |
| 941 | 174 | 4 | 2 | 2 | 2 | 248 | 264913582882808 | 1 | 32767 | 1 | 1 | 31 | 43694 | 8963 | 81 | 4881 | 16711729 | 2 |
| 942 | 174 | 4 | 2 | 2 | 2 | 248 | 264913582882814 | 1 | 8031 | 1 | 1 | 31 | 43759 | 15155 | 81 | 13057 | 7536691 | 2 |
| 943 | 174 | 4 | 2 | 2 | 2 | 248 | 264915864582384 | 1 | 3967 | 1 | 1 | 31 | 43759 | 8963 | 81 | 13057 | 1114128 | 2 |
| 944 | 174 | 4 | 2 | 2 | 2 | 248 | 264917353558256 | 1 | 30591 | 1 | 1 | 31 | 43759 | 47931 | 81 | 4864 | 5242896 | 2 |
| 945 | 174 | 4 | 2 | 2 | 2 | 248 | 264917587001082 | 1 | 3871 | 1 | 1 | 31 | 43759 | 47931 | 16 | 4353 | 7340032 | 2 |
| 946 | 174 | 4 | 2 | 2 | 2 | 248 | 264917617799408 | 1 | 24447 | 1 | 1 | 31 | 43694 | 10786 | 81 | 13057 | 3342352 | 2 |
| 947 | 174 | 4 | 2 | 2 | 2 | 248 | 278107722415352 | 1 | 32767 | 1 | 1 | 31 | 43694 | 15155 | 81 | 13057 | 7340032 | 2 |
| 948 | 174 | 4 | 2 | 2 | 2 | 248 | 278107722415856 | 1 | 16255 | 1 | 1 | 31 | 43759 | 10786 | 16 | 769 | 3211264 | 2 |
| 949 | 174 | 4 | 2 | 2 | 2 | 248 | 280306745671664 | 1 | 16255 | 1 | 1 | 31 | 43759 | 13091 | 81 | 4864 | 7536691 | 2 |
| 950 | 174 | 4 | 2 | 2 | 2 | 248 | 280925220962208 | 1 | 21847 | 1 | 1 | 31 | 43759 | 10786 | 81 | 4353 | 5308416 | 2 |
| 951 | 174 | 4 | 2 | 2 | 2 | 248 | 281200098869216 | 1 | 16255 | 1 | 1 | 31 | 43759 | 8963 | 16 | 13057 | 7340032 | 2 |
| 952 | 174 | 4 | 2 | 2 | 2 | 248 | 281202962137066 | 1 | 24447 | 1 | 1 | 31 | 43759 | 47931 | 81 | 4881 | 1114128 | 2 |
| 953 | 174 | 4 | 2 | 2 | 2 | 248 | 9223513326380681352 | 1 | 8031 | 1 | 1 | 31 | 43694 | 8963 | 81 | 4881 | 7667712 | 2 |
| 954 | 174 | 4 | 2 | 2 | 2 | 248 | 9223513326951115434 | 1 | 5631 | 1 | 1 | 31 | 43759 | 8994 | 16 | 769 | 7667712 | 2 |
| 955 | 174 | 4 | 2 | 2 | 2 | 248 | 9223513327521549516 | 1 | 1287 | 1 | 1 | 31 | 43759 | 47931 | 81 | 4353 | 7340080 | 2 |
| 956 | 174 | 4 | 2 | 2 | 2 | 248 | 9223513327521549516 | 1 | 24447 | 1 | 1 | 31 | 43694 | 8706 | 16 | 13057 | 5308416 | 2 |
| 957 | 174 | 4 | 2 | 2 | 2 | 248 | 9223548648065900704 | 1 | 21847 | 1 | 1 | 31 | 43759 | 47931 | 81 | 4353 | 15794176 | 2 |
| 958 | 174 | 4 | 2 | 2 | 2 | 248 | 9223548649131274480 | 1 | 32767 | 1 | 1 | 31 | 43759 | 8963 | 16 | 769 | 16711696 | 2 |
| 959 | 174 | 4 | 2 | 2 | 2 | 248 | 9223548650204954623 | 1 | 3871 | 1 | 1 | 31 | 43694 | 8963 | 81 | 1792 | 16187443 | 2 |
| 960 | 174 | 4 | 2 | 2 | 2 | 248 | 9223583970002783436 | 1 | 14199 | 1 | 1 | 31 | 43694 | 13091 | 16 | 14080 | 3211264 | 2 |
| 961 | 174 | 4 | 2 | 2 | 2 | 248 | 9223636952719359999 | 1 | 1287 | 1 | 1 | 31 | 43759 | 48043 | 81 | 1281 | 16711729 | 2 |
| 962 | 174 | 4 | 2 | 2 | 2 | 248 | 9223636953122009328 | 1 | 30591 | 1 | 1 | 31 | 43694 | 13091 | 16 | 769 | 7340032 | 2 |
| 963 | 174 | 4 | 2 | 2 | 2 | 248 | 9223636953122009328 | 1 | 32767 | 1 | 1 | 31 | 43759 | 13091 | 81 | 4881 | 7536640 | 2 |
| 964 | 174 | 4 | 2 | 2 | 2 | 248 | 9223636953671463152 | 1 | 7999 | 1 | 19 | 31 | 43759 | 48043 | 16 | 4864 | 3145744 | 2 |
| 965 | 174 | 4 | 2 | 2 | 2 | 248 | 9223636954732560383 | 1 | 22367 | 1 | 1 | 31 | 43759 | 15155 | 16 | 4864 | 16711729 | 2 |
| 966 | 174 | 4 | 2 | 2 | 2 | 248 | 9259542123273824896 | 1 | 32767 | 1 | 1 | 31 | 43759 | 48043 | 16 | 769 | 5242896 | 2 |
| 967 | 174 | 4 | 2 | 2 | 2 | 248 | 9259542124347572416 | 1 | 2047 | 1 | 1 | 31 | 43759 | 15155 | 81 | 4864 | 16711729 | 2 |
| 968 | 174 | 4 | 2 | 2 | 2 | 248 | 9259550920444792000 | 1 | 3967 | 1 | 1 | 31 | 43759 | 10786 | 16 | 4881 | 3211281 | 2 |
| 969 | 174 | 4 | 2 | 2 | 2 | 248 | 9259577446158626976 | 1 | 16255 | 1 | 1 | 31 | 43759 | 48043 | 81 | 769 | 3211264 | 2 |
| 970 | 174 | 4 | 2 | 2 | 2 | 248 | 9259588303301159584 | 1 | 32767 | 1 | 1 | 31 | 43759 | 2562 | 16 | 769 | 15728656 | 2 |
| 971 | 174 | 4 | 2 | 2 | 2 | 248 | 9259588303469587114 | 1 | 5631 | 1 | 1 | 31 | 43759 | 47931 | 16 | 4864 | 7340032 | 2 |
| 972 | 174 | 4 | 2 | 2 | 2 | 248 | 9259612493298060492 | 1 | 8191 | 1 | 1 | 31 | 43759 | 48043 | 81 | 4881 | 5242896 | 2 |
| 973 | 174 | 4 | 2 | 2 | 2 | 248 | 9259612493904408816 | 1 | 8191 | 1 | 1 | 31 | 43694 | 48043 | 16 | 769 | 16711729 | 2 |
| 974 | 174 | 4 | 2 | 2 | 2 | 248 | 9259612767600574400 | 1 | 24447 | 1 | 1 | 31 | 43759 | 43522 | 16 | 4881 | 15794176 | 2 |
| 975 | 174 | 4 | 2 | 2 | 2 | 248 | 9259612767600574400 | 1 | 30591 | 1 | 1 | 31 | 43694 | 15155 | 16 | 4864 | 5308416 | 2 |
| 976 | 174 | 4 | 2 | 2 | 2 | 248 | 9259612769026637760 | 1 | 7999 | 1 | 1 | 31 | 43759 | 13091 | 81 | 4864 | 16187392 | 2 |
| 977 | 174 | 4 | 2 | 2 | 2 | 248 | 9259665268576153840 | 1 | 8191 | 1 | 1 | 31 | 43694 | 48043 | 16 | 4864 | 16711696 | 2 |
| 978 | 174 | 4 | 2 | 2 | 2 | 248 | 9259665749746712560 | 1 | 22367 | 1 | 1 | 31 | 43759 | 10786 | 16 | 13057 | 16711729 | 2 |
| 979 | 174 | 4 | 2 | 2 | 2 | 248 | 9259665750149361904 | 1 | 2047 | 1 | 1 | 31 | 43694 | 47931 | 81 | 4881 | 16711696 | 2 |

| Model No. | Functions in integer form at the nodes of the network | | | | | | | | | | | | | | | | | |
|---|---|---|---|---|---|---|---|---|---|---|---|---|---|---|---|---|---|---|
| | CK | ARR1 | SHY2 | AUXIAA | ARF | ARF10 | ARF5 | XAL1 | PLT | AUX | SCR | SHR | MIR166 | PHB | JKD | MGP | WOX5 | CLE40 |
| 980 | 174 | 4 | 2 | 2 | 2 | 248 | 9799982874777127048 | 1 | 3871 | 1 | 1 | 31 | 43694 | 47931 | 81 | 4353 | 5308416 | 2 |
| 981 | 174 | 4 | 2 | 2 | 2 | 248 | 9799982876773646320 | 1 | 7999 | 1 | 1 | 31 | 43759 | 8963 | 81 | 769 | 16187392 | 2 |
| 982 | 174 | 4 | 2 | 2 | 2 | 248 | 9800018197158603424 | 1 | 7999 | 1 | 1 | 31 | 43759 | 8963 | 81 | 4864 | 16711729 | 2 |
| 983 | 174 | 4 | 2 | 2 | 2 | 248 | 9800018197729049840 | 1 | 22367 | 1 | 1 | 31 | 43759 | 2562 | 16 | 769 | 3342352 | 2 |
| 984 | 174 | 4 | 2 | 2 | 2 | 248 | 9800053518408141000 | 1 | 3871 | 1 | 19 | 31 | 43694 | 48043 | 16 | 22272 | 16711696 | 2 |
| 985 | 174 | 4 | 2 | 2 | 2 | 248 | 9800053520412442623 | 1 | 8031 | 1 | 1 | 31 | 43694 | 10786 | 16 | 4864 | 5308433 | 2 |
| 986 | 174 | 4 | 2 | 2 | 2 | 248 | 9800106502256655600 | 1 | 14199 | 1 | 1 | 31 | 43759 | 13091 | 81 | 4864 | 16711729 | 2 |
| 987 | 174 | 4 | 2 | 2 | 2 | 248 | 9836049192639310506 | 1 | 7999 | 1 | 1 | 31 | 43759 | 13091 | 16 | 769 | 7340080 | 2 |
| 988 | 174 | 4 | 2 | 2 | 2 | 248 | 9836049193211308704 | 1 | 7999 | 1 | 1 | 31 | 43694 | 10786 | 81 | 769 | 16187443 | 2 |
| 989 | 174 | 4 | 2 | 2 | 2 | 248 | 9836086714618137792 | 1 | 2047 | 1 | 1 | 31 | 43694 | 8963 | 16 | 4881 | 5308416 | 2 |
| 990 | 174 | 4 | 2 | 2 | 2 | 248 | 9836086714618924236 | 1 | 14143 | 1 | 1 | 31 | 43759 | 15155 | 81 | 4881 | 16187443 | 2 |

### SUPPLEMENTARY FIGURES

**(a) Network Structure**

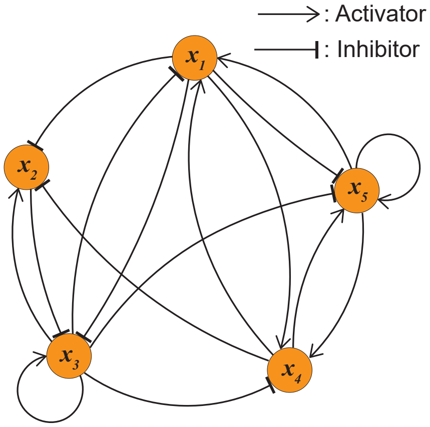

**(b) Boolean functions $f_i$ at each node $x_i$**

$$f_1 = \bar{x}_3 \wedge x_4 \wedge x_5$$
$$f_2 = \bar{x}_1 \vee (x_3 \wedge \bar{x}_4)$$
$$f_3 = \bar{x}_1 \wedge (\bar{x}_2 \vee x_3)$$
$$f_4 = x_1 \vee (\bar{x}_3 \wedge x_5)$$
$$f_5 = (\bar{x}_1 \wedge \bar{x}_3 \wedge x_4) \vee x_5$$

**(c) State transitions**

$\mathbf{X}(t) = (x_1(t), x_2(t), x_3(t), x_4(t), x_5(t))$

| $\mathbf{X}(t)$ | $\longrightarrow$ | $\mathbf{X}(t+1)$ |
|---|---|---|
| 00000 (0) | ⟶ | 01100 (12) |
| 00001 (1) | ⟶ | 01111 (15) |
| 00010 (2) | ⟶ | 01101 (13) |
| 00011 (3) | ⟶ | 11111 (31) |
| 00100 (4) | ⟶ | 01100 (12) |
| 00101 (5) | ⟶ | 01101 (13) |
| 00110 (6) | ⟶ | 01100 (12) |
| 00111 (7) | ⟶ | 01101 (13) |
| 01000 (8) | ⟶ | 01000 (8) |
| 01001 (9) | ⟶ | 01011 (11) |
| 01010 (10) | ⟶ | 01001 (9) |
| 01011 (11) | ⟶ | 11011 (27) |
| 01100 (12) | ⟶ | 01100 (12) |
| 01101 (13) | ⟶ | 01101 (13) |
| 01110 (14) | ⟶ | 01100 (12) |
| 01111 (15) | ⟶ | 01101 (13) |
| 10000 (16) | ⟶ | 00010 (2) |
| 10001 (17) | ⟶ | 00011 (3) |
| 10010 (18) | ⟶ | 00010 (2) |
| 10011 (19) | ⟶ | 10011 (19) |
| 10100 (20) | ⟶ | 01010 (10) |
| 10101 (21) | ⟶ | 01011 (11) |
| 10110 (22) | ⟶ | 00010 (2) |
| 10111 (23) | ⟶ | 00011 (3) |
| 11000 (24) | ⟶ | 00010 (2) |
| 11001 (25) | ⟶ | 00011 (3) |
| 11010 (26) | ⟶ | 00010 (2) |
| 11011 (27) | ⟶ | 10011 (19) |
| 11100 (28) | ⟶ | 01010 (10) |
| 11101 (29) | ⟶ | 01011 (11) |
| 11110 (30) | ⟶ | 00010 (2) |
| 11111 (31) | ⟶ | 00011 (3) |

**(d) State transition graph**

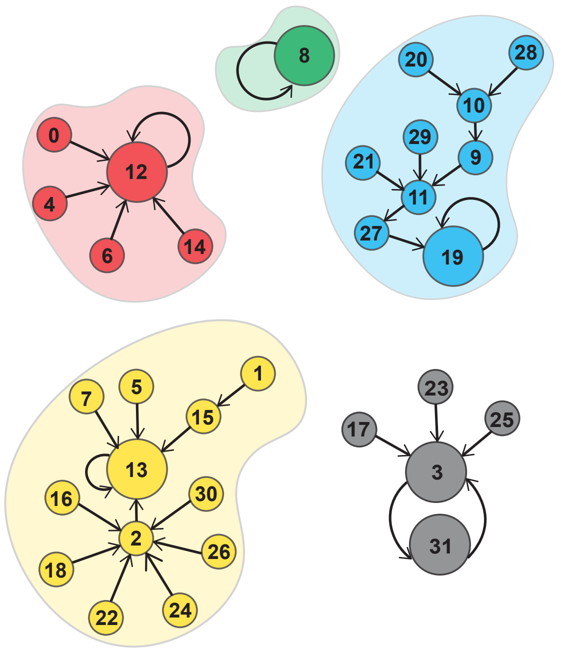

**(e) Obtained attractors**

| | $x_1$ | $x_2$ | $x_3$ | $x_4$ | $x_5$ |
|---|---|---|---|---|---|
| Att 1 | 0 | 1 | 0 | 0 | 0 |
| Att 2 | 0 | 1 | 1 | 0 | 0 |
| Att 3 | 1 | 0 | 0 | 1 | 1 |
| Att 4 | 0 | 1 | 1 | 0 | 1 |
| Att 5 | 0 | 0 | 0 | 1 | 1 |
| | 1 | 1 | 1 | 1 | 1 |

FIG. S1. **Toy Boolean network.** **(a)** The network structure of a toy Boolean model consisting of 5 genes and 16 edges. **(b)** Boolean function (BF) at each node specified as a Boolean expression of its input variables. **(c)** The state to state transitions for all $2^5 = 32$ states when synchronously using the Boolean functions or logic rules provided in (b). **(d)** The state to state transitions in (c) lead to trajectories which either converge to fixed point attractors (as shown in colors green, blue, red and yellow) or cyclic attractors (as shown in grey). **(e)** The attractor states of this model. The cyclic attractor consists of 2 states shown in grey.

**(a) Gene regulatory network and attractors of *Arabidopsis thaliana* Root Stem Cell Niche (RSCN)** (Azpeitia *et al.* *2010*)

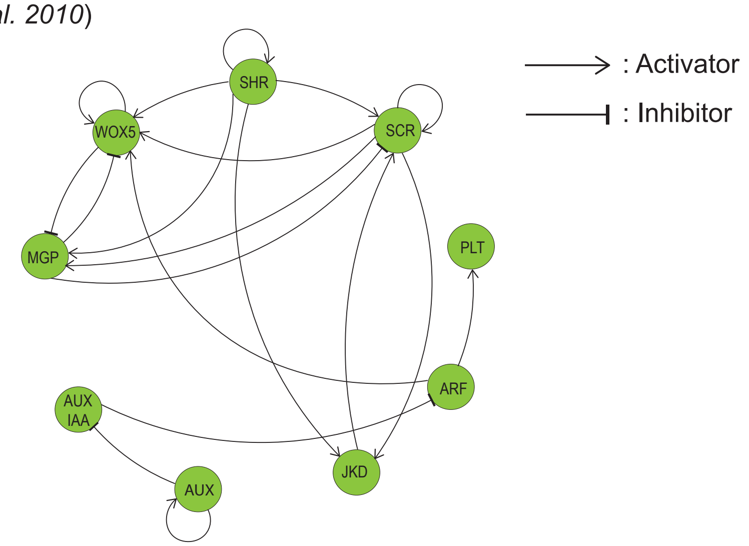

| | SCR | PLT | ARF | AUXIAA | AUX | SHR | JKD | MGP | WOX5 |
|---|---|---|---|---|---|---|---|---|---|
| **QC** | 1 | 1 | 1 | 0 | 1 | 1 | 1 | 0 | 1 |
| **VI** | 0 | 1 | 1 | 0 | 1 | 1 | 0 | 0 | 0 |
| **CEI** | 1 | 1 | 1 | 0 | 1 | 1 | 1 | 1 | 0 |
| **CEpI** | 0 | 1 | 1 | 0 | 1 | 0 | 0 | 0 | 0 |

**(b) Gene regulatory network and attractors of Pancreas cell differentiation** (Zhou *et al.* *2016*)

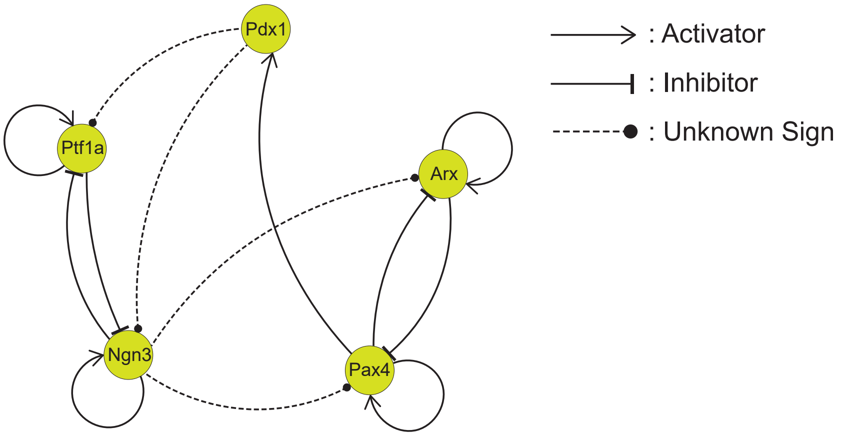

| | Pdx1 | Ptf1a | Ngn3 | Pax4 | Arx |
|---|---|---|---|---|---|
| **Exocrine** | 0 | 1 | 0 | 0 | 0 |
| **β/δ cell progenitor** | 1 | 0 | 1 | 1 | 0 |
| **α/PP cell progenitor** | 0 | 0 | 1 | 0 | 1 |

FIG. S2. **Model systems used to construct ensembles of Boolean models.** **(a)** *Arabidopsis thaliana* RSCN Boolean GRN and its attractors. The network is constructed using regulatory interactions obtained from the BFs of *model A* in [12]. Here, QC: Quiescent center, VI: Vascular initials, CEI: Cortex-Endodermis initials, CEpI: Columella epidermis initials (CEpI) **(b)** Pancreas cell differentiation model Boolean GRN and its attractors.

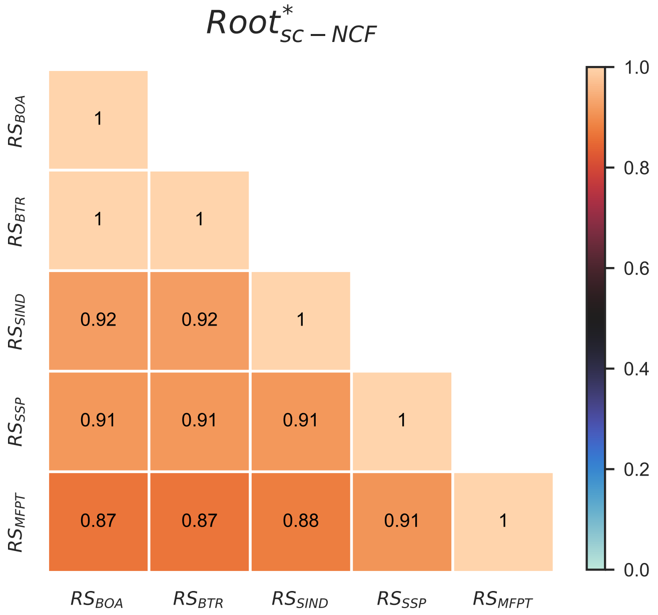

FIG. S3. **Pearson correlation between different pairs of relative stability measures for the ensemble $Root^*_{sc-NCF}$.** The rows and columns correspond to choices for the relative stability measures. These 5 measures are based on size of basin of attraction ($RS_{BOA}$), basin transition rates ($RS_{BTR}$), a stability index ($RS_{SIND}$), steady state probabilities ($RS_{SSP}$) and mean first passage times ($RS_{MFPT}$). The heatmap indicates the value of the Pearson correlation coefficient between pairs of these measures. Note that these measures are computed by exact means across all pairs of biological fixed points, for all 170 models in this ensemble $Root^*_{sc-NCF}$ using a noise intensity parameter value of 1%. The upper triangular portion of the heatmap is not displayed as the heatmap entries constitute a symmetric matrix. Furthermore, $RS_{BOA}$ and $RS_{BTR}$ are perfectly correlated, an observation which we prove theoretically by showing that $RS_{BOA}$ and $RS_{BTR}$ are in fact equivalent.

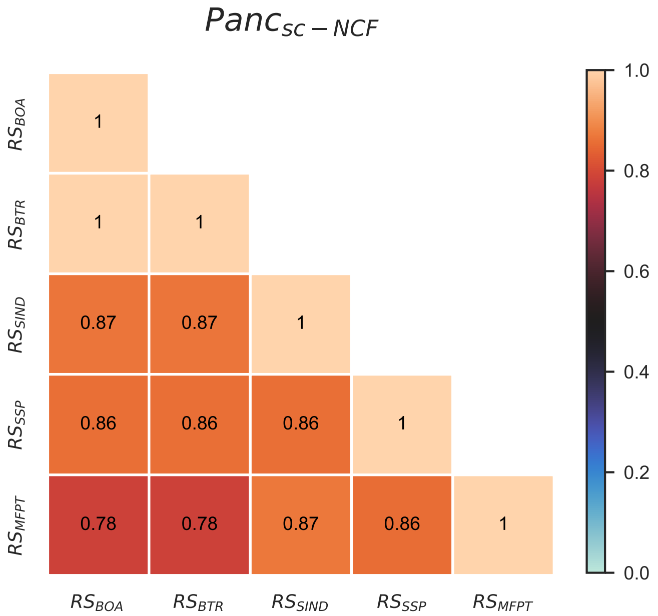

FIG. S4. **Pearson correlation between different pairs of relative stability measures for the ensemble $Panc_{sc-NCF}$** The rows and columns correspond to choices for the relative stability measures. These 5 measures are based on size of basin of attraction ($RS_{BOA}$), basin transition rates ($RS_{BTR}$), a stability index ($RS_{SIND}$), steady state probabilities ($RS_{SSP}$) and mean first passage times ($RS_{MFPT}$). The heatmap indicates the value of the Pearson correlation coefficient between pairs of these measures. Note that these measures are computed by exact means across all pairs of biological fixed points, for all 3600 models in this ensemble $Panc_{sc-NCF}$ using a noise intensity parameter value of 1%. The upper triangular portion of the heatmap is not displayed as the heatmap entries constitute a symmetric matrix. Furthermore, $RS_{BOA}$ and $RS_{BTR}$ are perfectly correlated, an observation which we prove theoretically by showing that $RS_{BOA}$ and $RS_{BTR}$ are in fact equivalent.

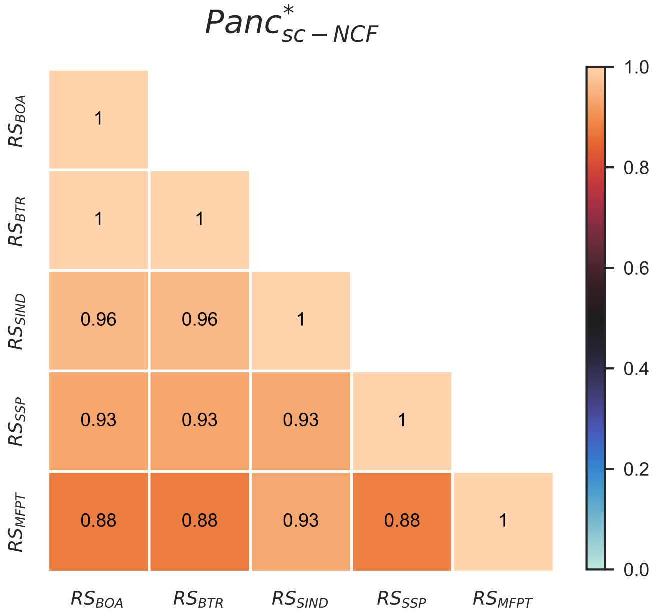

FIG. S5. **Pearson correlation between different pairs of relative stability measures for the ensemble $Panc^*_{sc-NCF}$.** The rows and columns correspond to choices for the relative stability measures. These 5 measures are based on size of basin of attraction ($RS_{BOA}$), basin transition rates ($RS_{BTR}$), a stability index ($RS_{SIND}$), steady state probabilities ($RS_{SSP}$) and mean first passage times ($RS_{MFPT}$). The heatmap indicates the value of the Pearson correlation coefficient between pairs of these measures. Note that these measures are computed by exact means across all pairs of biological fixed points, for all 109 models in this ensemble $Panc^*_{sc-NCF}$ using a noise intensity parameter value of 1%. The upper triangular portion of the heatmap is not displayed as the heatmap entries constitute a symmetric matrix. Furthermore, $RS_{BOA}$ and $RS_{BTR}$ are perfectly correlated, an observation which we prove theoretically by showing that $RS_{BOA}$ and $RS_{BTR}$ are in fact equivalent.

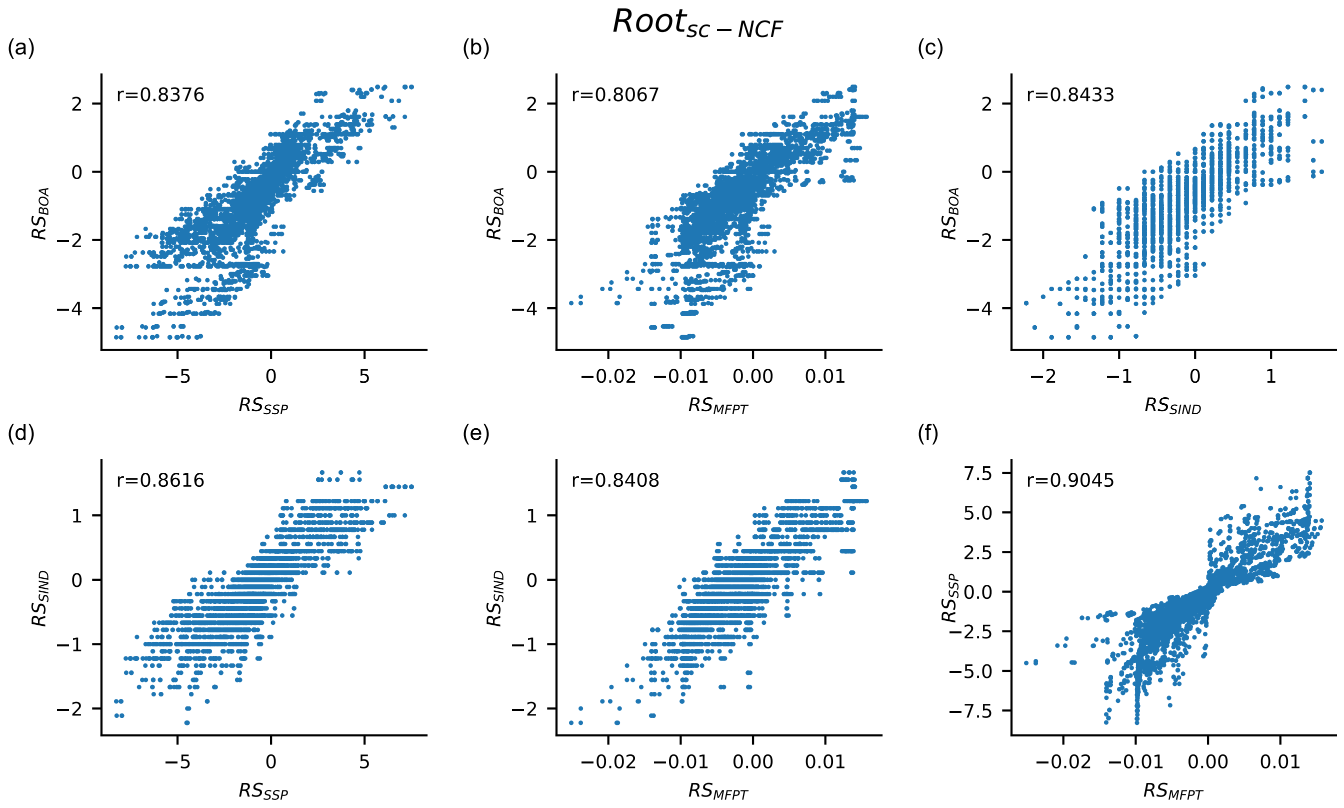

FIG. S6. **Scatter plots displaying values of relative stability in the ensemble $\boldsymbol{Root_{sc-NCF}}$.** Each subfigure from (a) to (f) is a scatter plot where the $x$ and $y$ axes are for different measures of relative stability based on size of basin of attraction ($RS_{BOA}$), basin transition rate ($RS_{BTR}$), stability index ($RS_{SIND}$), steady state probability ($RS_{SSP}$) and mean first passage time ($RS_{MFPT}$). These measures have been computed by the exact method for all pair of biological fixed points, for all 1275 models belonging to the ensemble $Root_{sc-NCF}$, at 1% noise. Of the 10 possible scatter plots for distinct pairs of the 5 relative stability measures, only 6 are shown here as $RS_{BOA}$ and $RS_{BTR}$ are equivalent. The Pearson correlation coefficient (r) for each scatter plot is computed and reported in the plot. These plots indicate that the correlation between the different relative stability measures is quite strong.

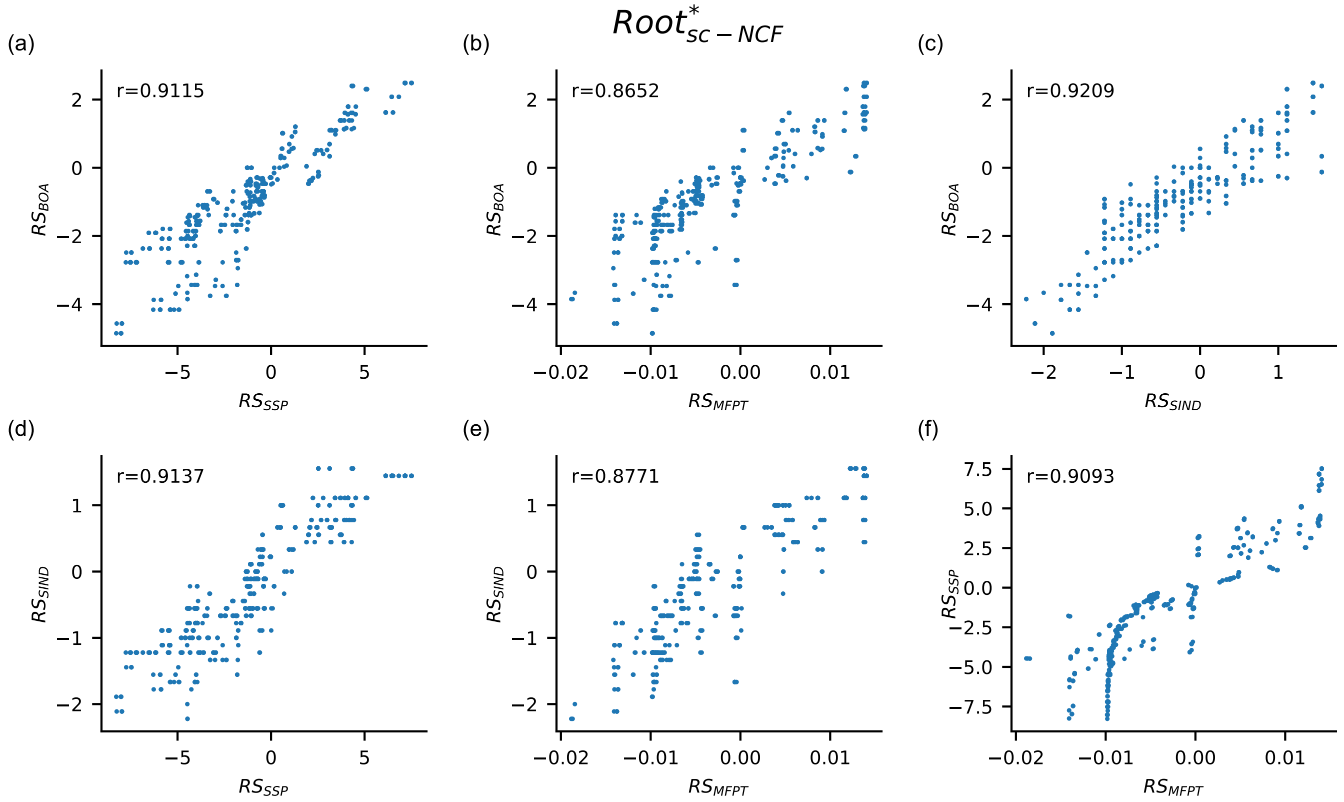

FIG. S7. **Scatter plots displaying values of relative stability in the ensemble $Root^*_{sc-NCF}$.** Each subfigure from (a) to (f) is a scatter plot where the $x$ and $y$ axes are for different measures of relative stability based on size of basin of attraction ($RS_{BOA}$), basin transition rate ($RS_{BTR}$), stability index ($RS_{SIND}$), steady state probability ($RS_{SSP}$) and mean first passage time ($RS_{MFPT}$). These measures have been computed by the exact method for all pair of biological fixed points, for all 170 models belonging to the ensemble $Root^*_{sc-NCF}$, at 1% noise. Of the 10 possible scatter plots for distinct pairs of the 5 relative stability measures, only 6 are shown here as $RS_{BOA}$ and $RS_{BTR}$ are equivalent. The Pearson correlation coefficient (r) for each scatter plot is computed and reported in the plot. These plots indicate that the correlation between the different relative stability measures is quite strong.

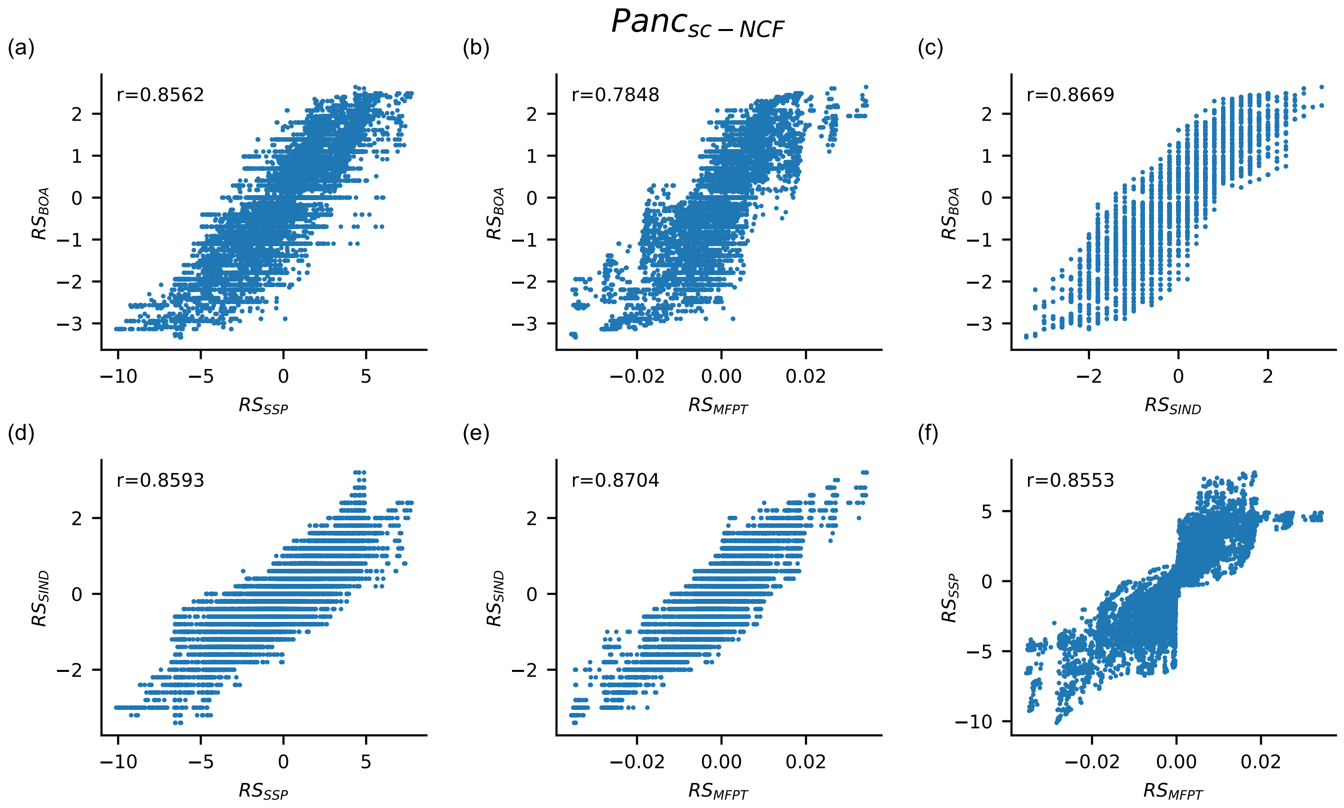

FIG. S8. **Scatter plots displaying values of relative stability in the ensemble $Panc_{sc-NCF}$.** Each subfigure from (a) to (f) is a scatter plot where the $x$ and $y$ axes are for different measures of relative stability based on size of basin of attraction ($RS_{BOA}$), basin transition rate ($RS_{BTR}$), stability index ($RS_{SIND}$), steady state probability ($RS_{SSP}$) and mean first passage time ($RS_{MFPT}$). These measures have been computed by the exact method for all pair of biological fixed points, for all 3600 models belonging to the ensemble $Panc_{sc-NCF}$, at 1% noise. Of the 10 possible scatter plots for distinct pairs of the 5 relative stability measures, only 6 are shown here as $RS_{BOA}$ and $RS_{BTR}$ are equivalent. The Pearson correlation coefficient (r) for each scatter plot is computed and reported in the plot. These plots indicate that the correlation between the different relative stability measures is quite strong.

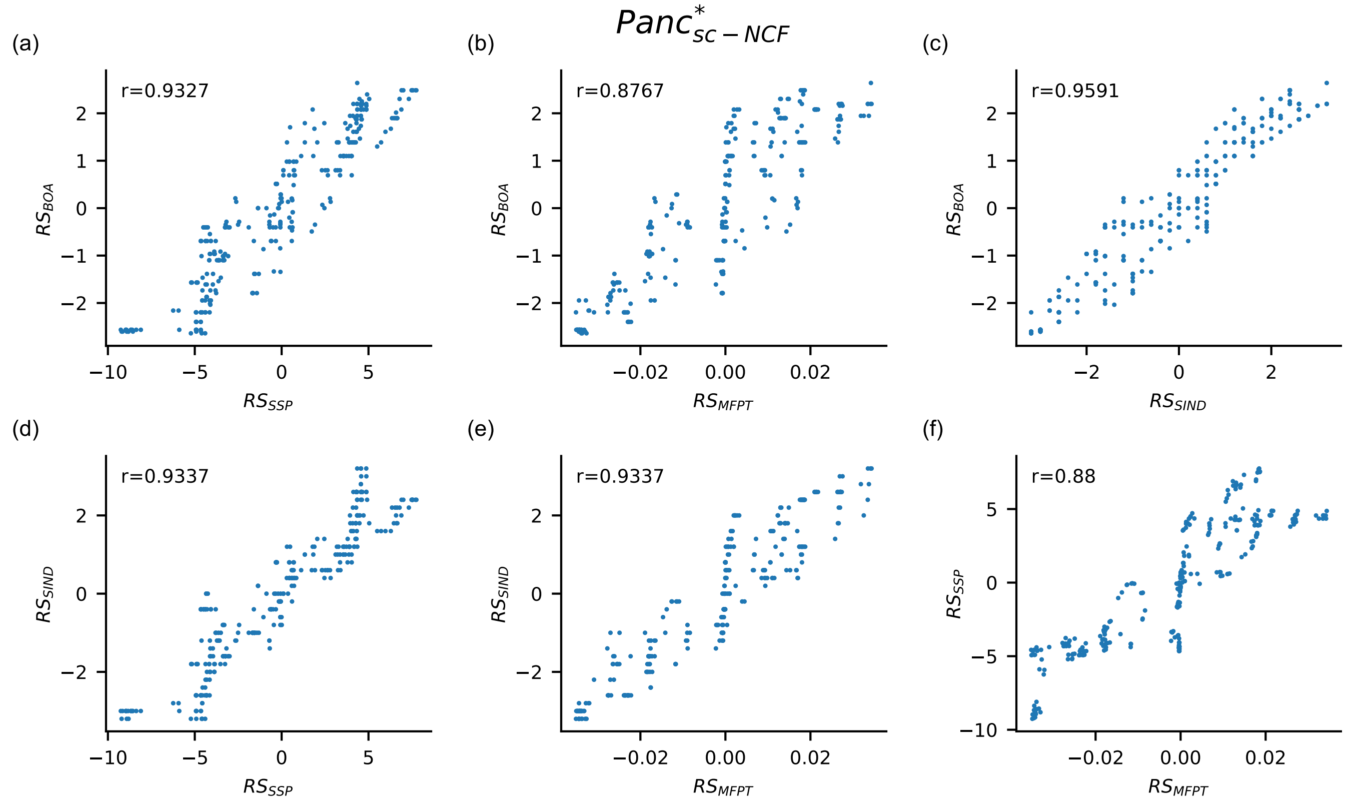

FIG. S9. **Scatter plots displaying values of relative stability in the ensemble $\boldsymbol{Panc^*_{sc-NCF}}$.** Each subfigure from (a) to (f) is a scatter plot where the $x$ and $y$ axes are for different measures of relative stability based on size of basin of attraction ($RS_{BOA}$), basin transition rate ($RS_{BTR}$), stability index ($RS_{SIND}$), steady state probability ($RS_{SSP}$) and mean first passage time ($RS_{MFPT}$). These measures have been computed by the exact method for all pair of biological fixed points, for all 109 models belonging to the ensemble $Panc^*_{sc-NCF}$, at 1% noise. Of the 10 possible scatter plots for distinct pairs of the 5 relative stability measures, only 6 are shown here as $RS_{BOA}$ and $RS_{BTR}$ are equivalent. The Pearson correlation coefficient (r) for each scatter plot is computed and reported in the plot. These plots indicate that the correlation between the different relative stability measures is quite strong.

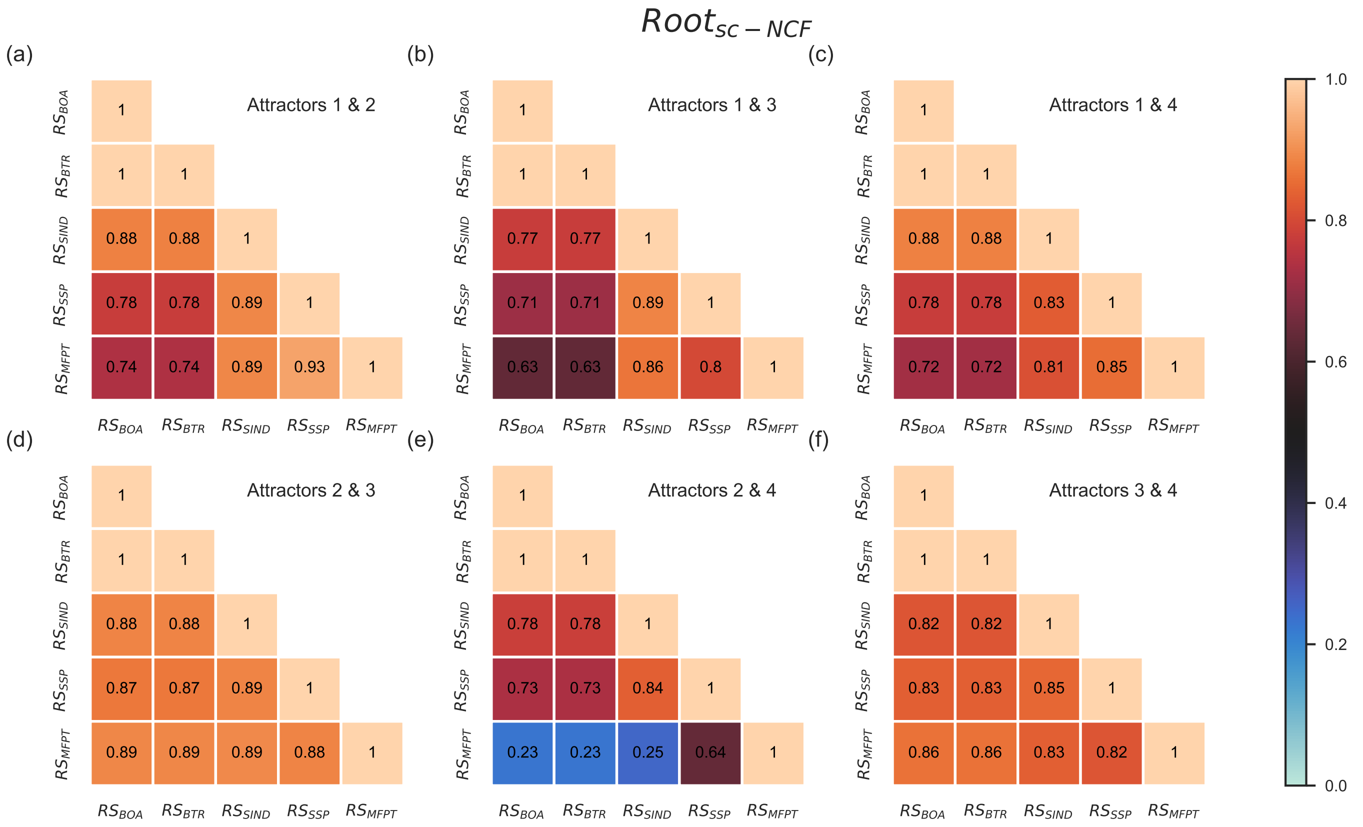

FIG. S10. **Pearson correlation between different pairs of relative stability measures for a given pair of fixed points for the ensemble $Root_{sc-NCF}$.** The rows and columns of all heatmaps correspond to choices for the relative stability measures. The 5 measures of relative stability are based on size of basin of attraction ($RS_{BOA}$), basin transition rates ($RS_{BTR}$), a stability index ($RS_{SIND}$), steady state probabilities ($RS_{SSP}$) and mean first passage times ($RS_{MFPT}$). The heatmaps indicate the Pearson correlation coefficient between pairs of these measures. For a particular subfigure, these measures are computed by exact means for the pair of biological fixed points specified in that subfigure, for all 1275 models in the ensemble $Root_{sc-NCF}$ using a noise intensity parameter value of 1%. Each biological attractor (fixed point) is numbered as follows. 1: Quiescent center (QC), 2: Vascular initials (VI), 3: Cortex-Endodermis initials (CEI), 4: Columella epidermis initials (CEpI). The upper triangular portion of the heat map is not displayed as the heatmap entries constitute a symmetric matrix. Furthermore, $RS_{BOA}$ and $RS_{BTR}$ are perfectly correlated, an observation which we prove theoretically by showing that $RS_{BOA}$ and $RS_{BTR}$ are in fact equivalent.

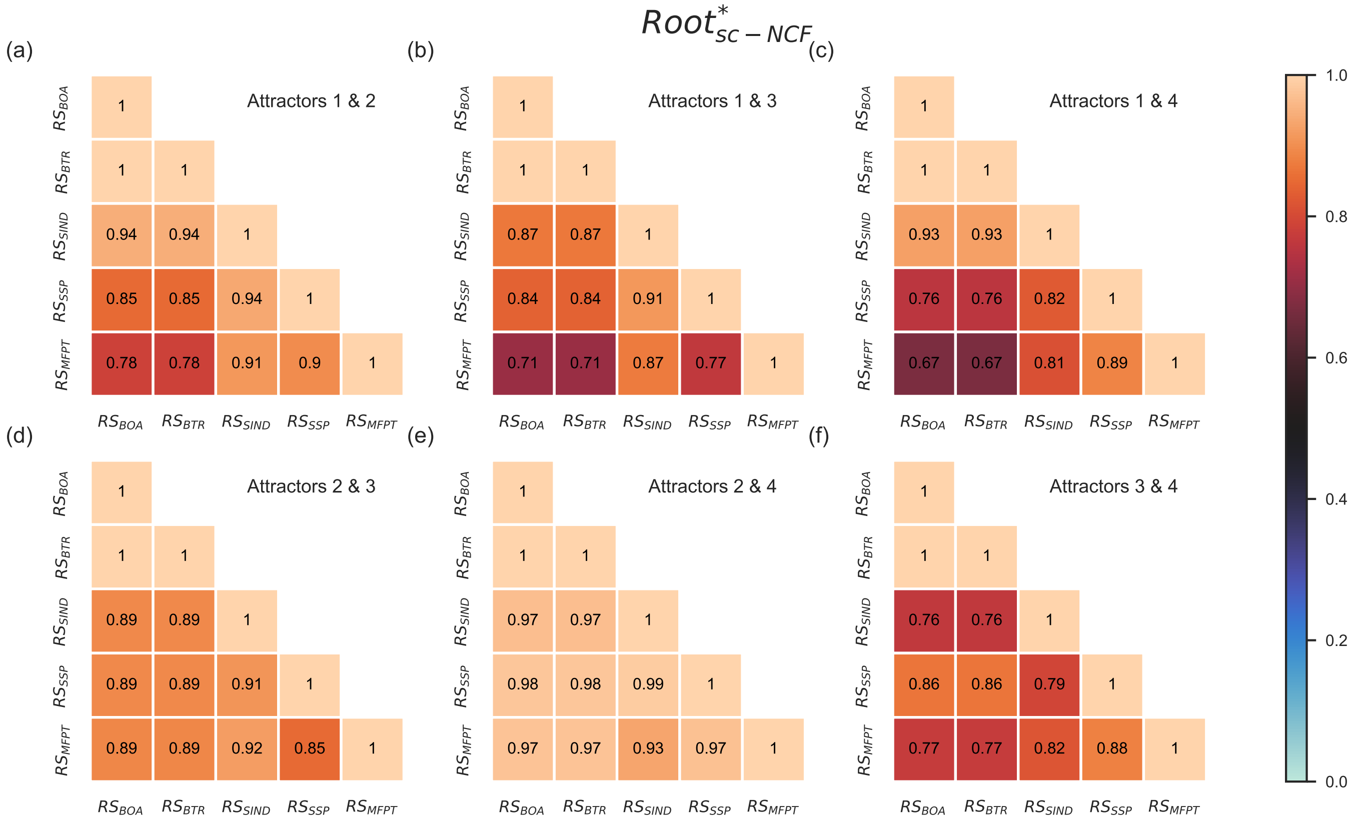

FIG. S11. **Pearson correlation between different pairs of relative stability measures for a given pair of fixed points for the ensemble $Root^*_{sc-NCF}$.** The rows and columns of all heatmaps correspond to choices for the relative stability measures. The 5 measures of relative stability are based on size of basin of attraction ($RS_{BOA}$), basin transition rates ($RS_{BTR}$), a stability index ($RS_{SIND}$), steady state probabilities ($RS_{SSP}$) and mean first passage times ($RS_{MFPT}$). The heatmaps indicate the Pearson correlation coefficient between pairs of these measures. For a particular subfigure, these measures are computed by exact means for the pair of biological fixed points specified in that subfigure, for all 170 models in the ensemble $Root^*_{sc-NCF}$ using a noise intensity parameter value of 1%. Each biological attractor (fixed point) is numbered as follows. 1: Quiescent center (QC), 2: Vascular initials (VI), 3: Cortex-Endodermis initials (CEI), 4: Columella epidermis initials (CEpI). The upper triangular portion of the heat map is not displayed as the heatmap entries constitute a symmetric matrix. Furthermore, $RS_{BOA}$ and $RS_{BTR}$ are perfectly correlated, an observation which we prove theoretically by showing that $RS_{BOA}$ and $RS_{BTR}$ are in fact equivalent.

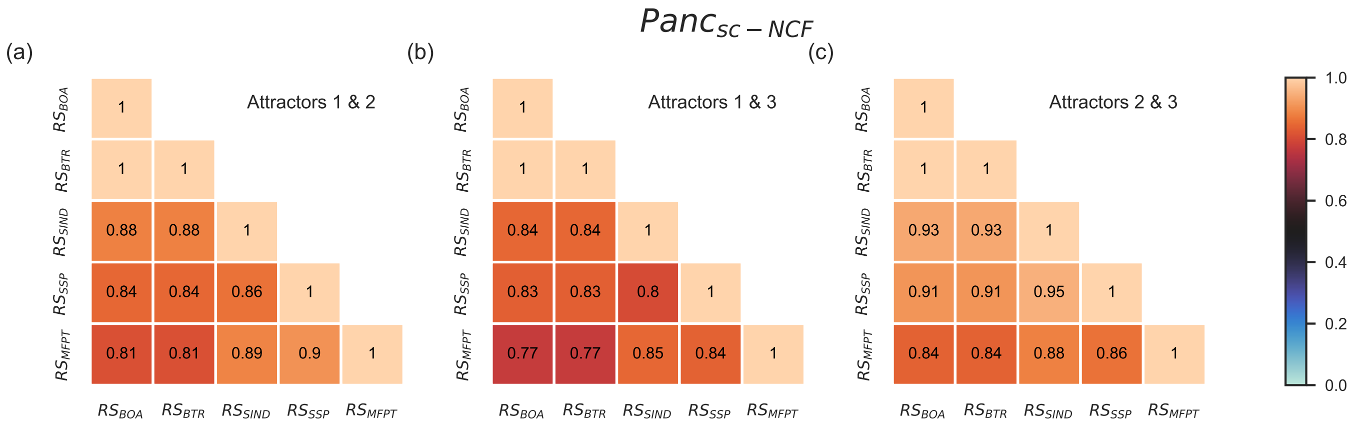

FIG. S12. **Pearson correlation between different pairs of relative stability measures for a given pair of fixed points for the ensemble $Panc_{sc-NCF}$.** The rows and columns of all heatmaps correspond to choices for the relative stability measures. The 5 measures of relative stability are based on size of basin of attraction ($RS_{BOA}$), basin transition rates ($RS_{BTR}$), a stability index ($RS_{SIND}$), steady state probabilities ($RS_{SSP}$) and mean first passage times ($RS_{MFPT}$). The heatmaps indicate the Pearson correlation coefficient between pairs of these measures. For a particular subfigure, these measures are computed by exact means for the pair of biological fixed points specified in that subfigure, for all 3600 models in the ensemble $Panc_{sc-NCF}$ using a noise intensity parameter value of 1%. Each biological attractor (fixed point) is numbered as follows. 1: Exocrine, 2: $\beta/\delta$ progenitor, 3: $\alpha$/PP progenitor. The upper triangular portion of the heat map is not displayed as the heatmap entries constitute a symmetric matrix. Furthermore, $RS_{BOA}$ and $RS_{BTR}$ are perfectly correlated, an observation which we prove theoretically by showing that $RS_{BOA}$ and $RS_{BTR}$ are in fact equivalent.

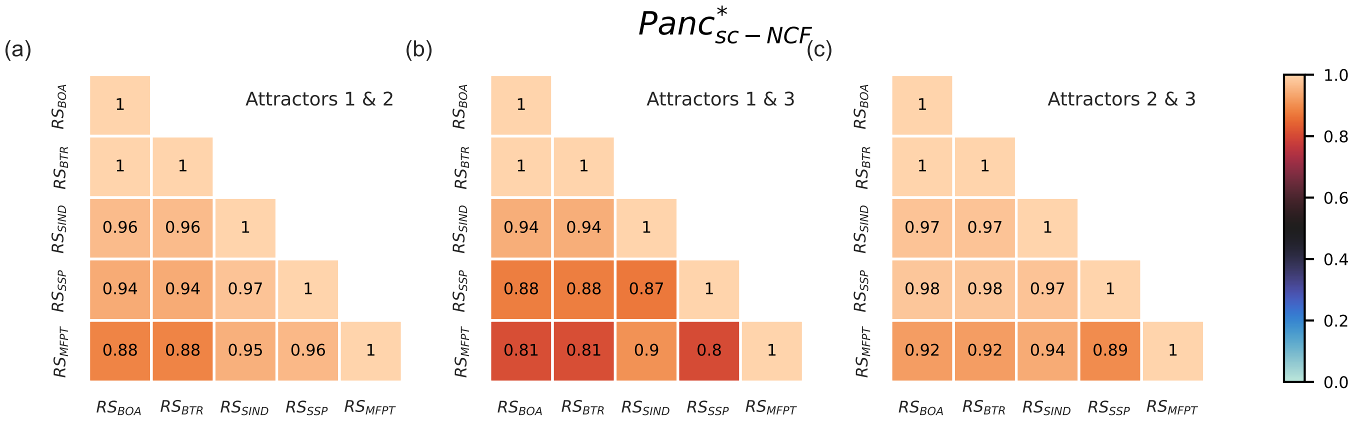

FIG. S13. **Pearson correlation between different pairs of relative stability measures for a given pair of fixed points for the ensemble $Panc^*_{sc-NCF}$.** The rows and columns of all heatmaps correspond to choices for the relative stability measures. The 5 measures of relative stability are based on size of basin of attraction ($RS_{BOA}$), basin transition rates ($RS_{BTR}$), a stability index ($RS_{SIND}$), steady state probabilities ($RS_{SSP}$) and mean first passage times ($RS_{MFPT}$). The heatmaps indicate the Pearson correlation coefficient between pairs of these measures. For a particular subfigure, these measures are computed by exact means for the pair of biological fixed points specified in that subfigure, for all 109 models in the ensemble $Panc^*_{sc-NCF}$ using a noise intensity parameter value of 1%. Each biological attractor (fixed point) is numbered as follows. 1: Exocrine, 2: $\beta/\delta$ progenitor, 3: $\alpha$/PP progenitor. The upper triangular portion of the heat map is not displayed as the heatmap entries constitute a symmetric matrix. Furthermore, $RS_{BOA}$ and $RS_{BTR}$ are perfectly correlated, an observation which we prove theoretically by showing that $RS_{BOA}$ and $RS_{BTR}$ are in fact equivalent.

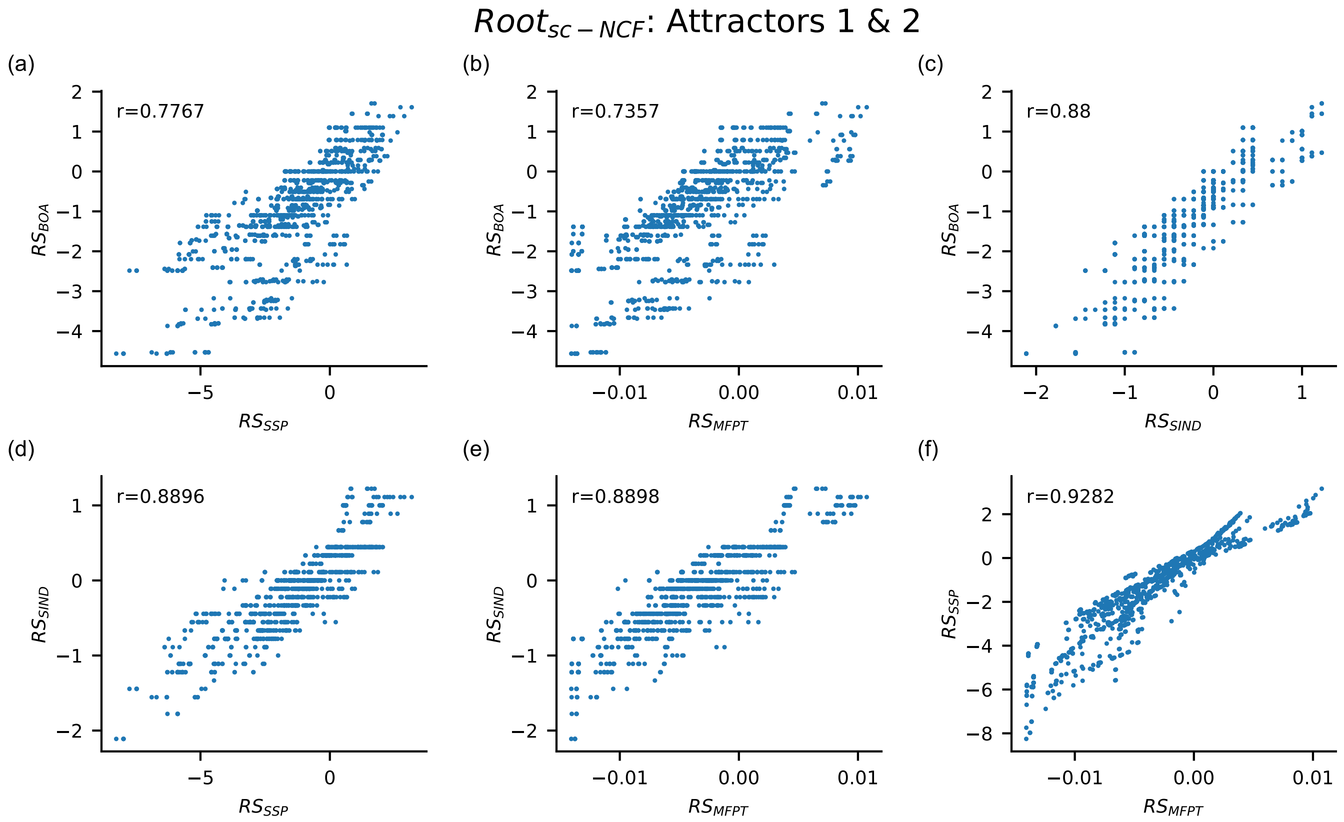

FIG. S14. **Scatter plots between the different pairs of relative stability measures for the pair of attractors 1 and 2 for the ensemble $\boldsymbol{Root_{sc-NCF}}$.** Each subfigure from (a) to (f) is a scatter plot where the $x$ and $y$ axes are for different measures of relative stability based on size of basin of attraction ($RS_{BOA}$), basin transition rate ($RS_{BTR}$), stability index ($RS_{SIND}$), steady state probability ($RS_{SSP}$) and mean first passage time ($RS_{MFPT}$). All these measures have been computed by the exact method for the pair of biological fixed points 1 (Quiescent center (QC)) and 2 (Vascular initials (VI)), for all 1275 models belonging to the ensemble $Root_{sc-NCF}$, at 1% noise. Of the 10 possible scatter plots for distinct pairs of the 5 relative stability measures, only 6 are shown here as $RS_{BOA}$ and $RS_{BTR}$ are equivalent. The Pearson correlation coefficient (r) for each scatter plot is computed and reported in the plot.

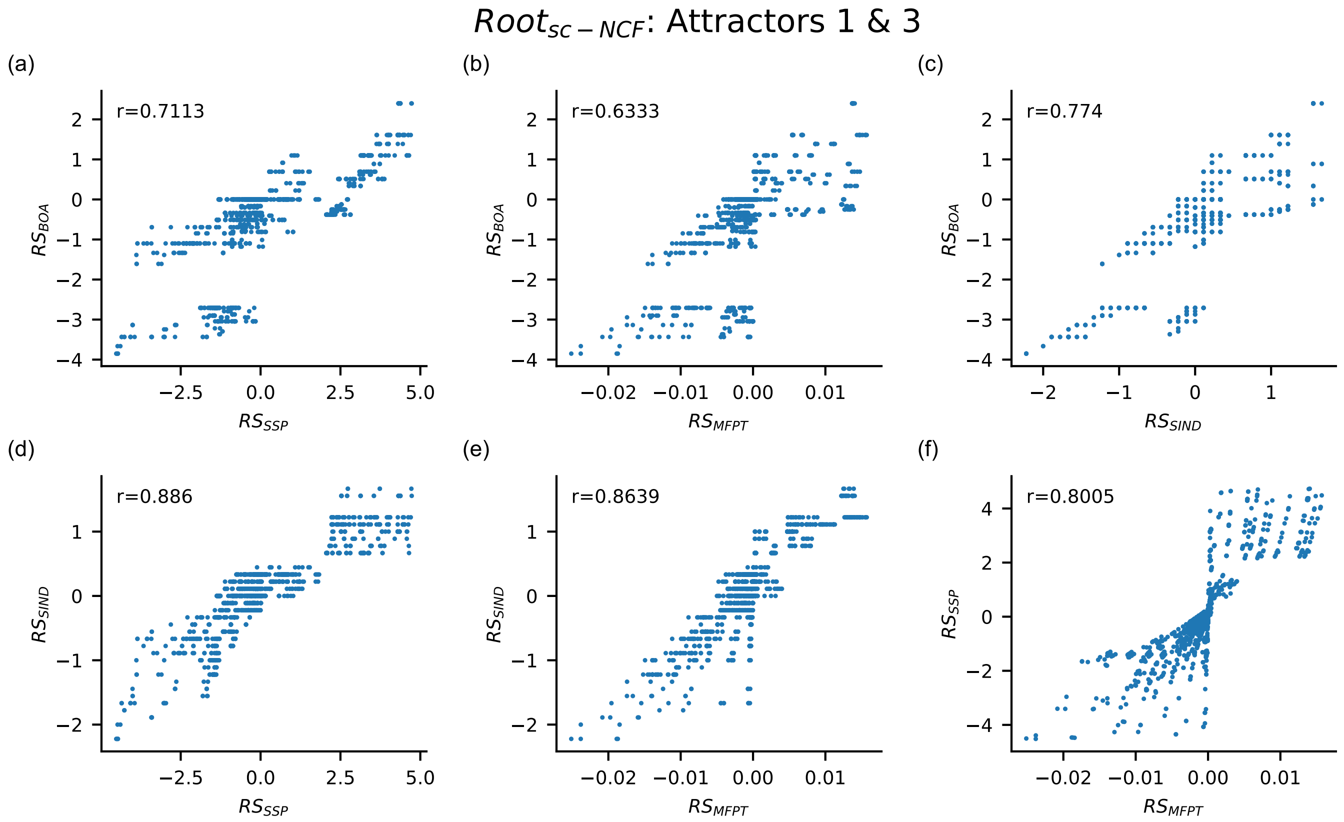

FIG. S15. **Scatter plots between the different pairs of relative stability measures for the pair of attractors** 1 **and** 3 **for the ensemble $Root_{sc-NCF}$.** Each subfigure from (a) to (f) is a scatter plot where the $x$ and $y$ axes are for different measures of relative stability based on size of basin of attraction ($RS_{BOA}$), basin transition rate ($RS_{BTR}$), stability index ($RS_{SIND}$), steady state probability ($RS_{SSP}$) and mean first passage time ($RS_{MFPT}$). All these measures have been computed by the exact method for the pair of biological fixed points 1 (Quiescent center (QC)) and 3 (Cortex-Endodermis initials (CEI)), for all 1275 models belonging to the ensemble $Root_{sc-NCF}$, at 1% noise. Of the 10 possible scatter plots for distinct pairs of the 5 relative stability measures, only 6 are shown here as $RS_{BOA}$ and $RS_{BTR}$ are equivalent. The Pearson correlation coefficient (r) for each scatter plot is computed and reported in the plot.

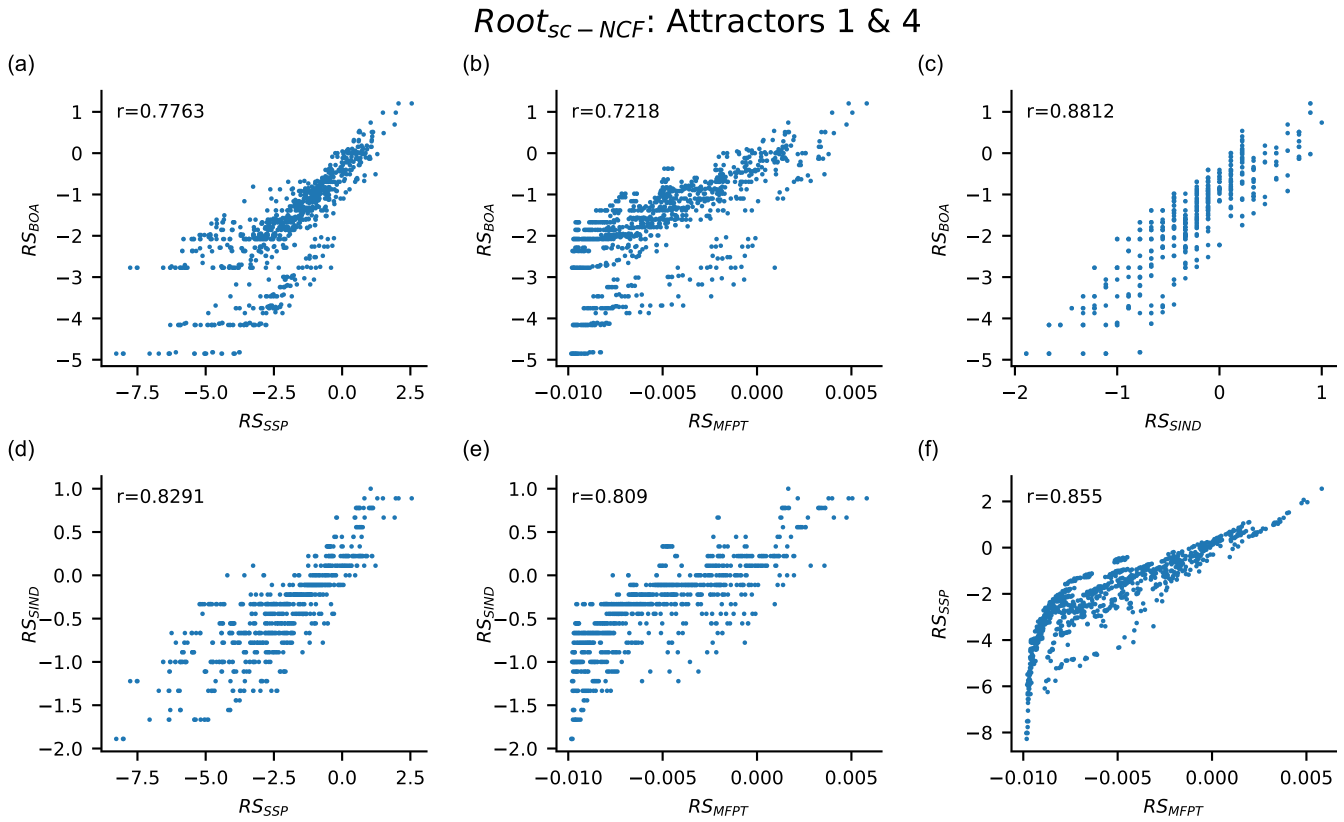

FIG. S16. **Scatter plots between the different pairs of relative stability measures for the pair of attractors** 1 **and** 4 **for the ensemble** $\boldsymbol{Root_{sc-NCF}}$. Each subfigure from (a) to (f) is a scatter plot where the $x$ and $y$ axes are for different measures of relative stability based on size of basin of attraction ($RS_{BOA}$), basin transition rate ($RS_{BTR}$), stability index ($RS_{SIND}$), steady state probability ($RS_{SSP}$) and mean first passage time ($RS_{MFPT}$). All these measures have been computed by the exact method for the pair of biological fixed points 1 (Quiescent center (QC)) and 4 (Columella epidermis initials (CEpI)), for all 1275 models belonging to the ensemble $Root_{sc-NCF}$, at 1% noise. Of the 10 possible scatter plots for distinct pairs of the 5 relative stability measures, only 6 are shown here as $RS_{BOA}$ and $RS_{BTR}$ are equivalent. The Pearson correlation coefficient (r) for each scatter plot is computed and reported in the plot.

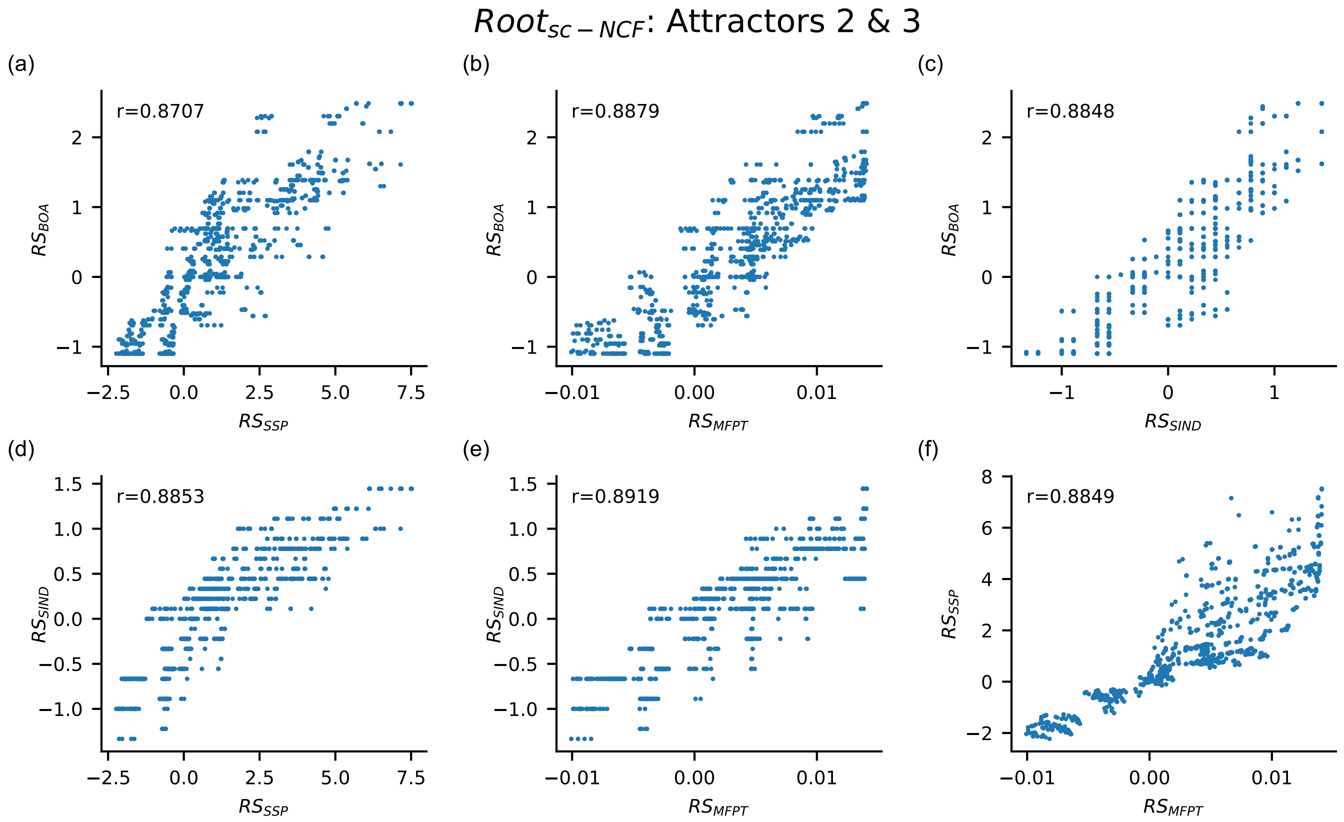

FIG. S17. **Scatter plots between the different pairs of relative stability measures for the pair of attractors 2 and 3 for the ensemble $\boldsymbol{Root_{sc-NCF}}$.** Each subfigure from (a) to (f) is a scatter plot where the $x$ and $y$ axes are for different measures of relative stability based on size of basin of attraction ($RS_{BOA}$), basin transition rate ($RS_{BTR}$), stability index ($RS_{SIND}$), steady state probability ($RS_{SSP}$) and mean first passage time ($RS_{MFPT}$). All these measures have been computed by the exact method for the pair of biological fixed points 2 (Vascular initials (VI)) and 3 (Cortex-Endodermis initials (CEI)), for all 1275 models belonging to the ensemble $Root_{sc-NCF}$, at 1% noise. Of the 10 possible scatter plots for distinct pairs of the 5 relative stability measures, only 6 are shown here as $RS_{BOA}$ and $RS_{BTR}$ are equivalent. The Pearson correlation coefficient (r) for each scatter plot is computed and reported in the plot.

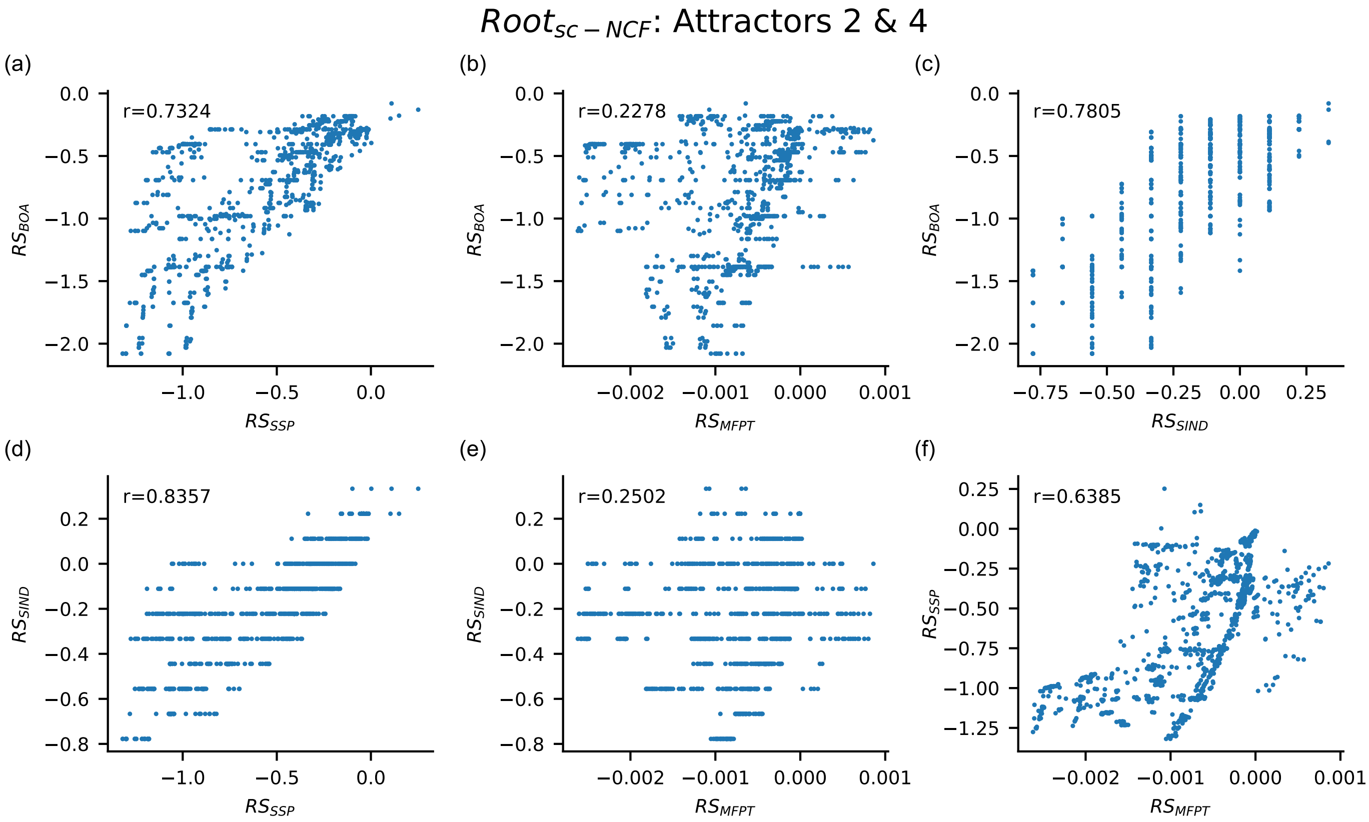

FIG. S18. **Scatter plots between the different pairs of relative stability measures for the pair of attractors** 2 **and** 4 **for the ensemble** $\boldsymbol{Root_{sc-NCF}}$. Each subfigure from (a) to (f) is a scatter plot where the $x$ and $y$ axes are for different measures of relative stability based on size of basin of attraction ($RS_{BOA}$), basin transition rate ($RS_{BTR}$), stability index ($RS_{SIND}$), steady state probability ($RS_{SSP}$) and mean first passage time ($RS_{MFPT}$). All these measures have been computed by the exact method for the pair of biological fixed points 2 (Vascular initials (VI)) and 4 (Columella epidermis initials (CEpI)), for all 1275 models belonging to the ensemble $Root_{sc-NCF}$, at 1% noise. Of the 10 possible scatter plots for distinct pairs of the 5 relative stability measures, only 6 are shown here as $RS_{BOA}$ and $RS_{BTR}$ are equivalent. The Pearson correlation coefficient (r) for each scatter plot is computed and reported in the plot.

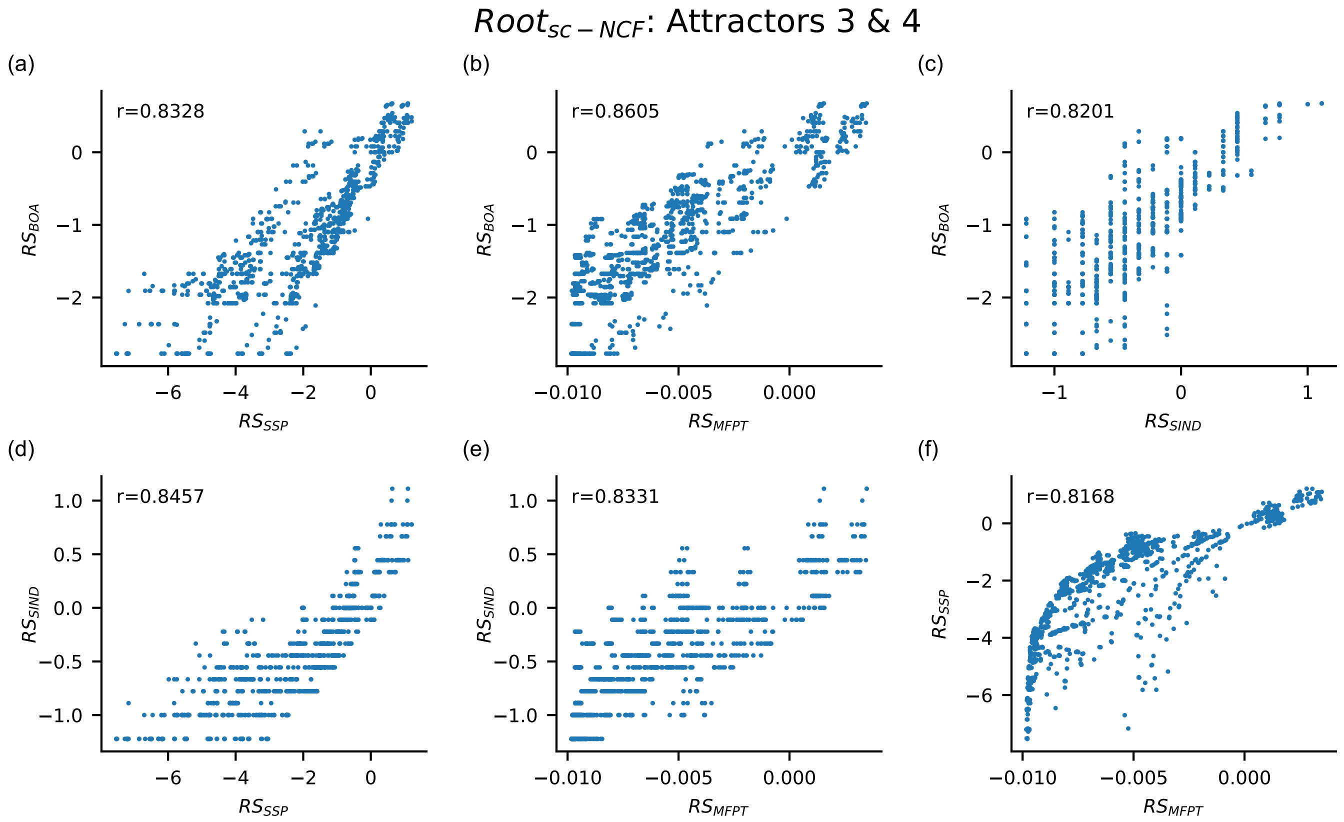

FIG. S19. **Scatter plots between the different pairs of relative stability measures for the pair of attractors 3 and 4 for the ensemble $\boldsymbol{Root_{sc-NCF}}$.** Each subfigure from (a) to (f) is a scatter plot where the $x$ and $y$ axes are for different measures of relative stability based on size of basin of attraction ($RS_{BOA}$), basin transition rate ($RS_{BTR}$), stability index ($RS_{SIND}$), steady state probability ($RS_{SSP}$) and mean first passage time ($RS_{MFPT}$). All these measures have been computed by the exact method for the pair of biological fixed points 3 (Cortex-Endodermis initials (CEI)) and 4 (Columella epidermis initials (CEpI)), for all 1275 models belonging to the ensemble $Root_{sc-NCF}$, at 1% noise. Of the 10 possible scatter plots for distinct pairs of the 5 relative stability measures, only 6 are shown here as $RS_{BOA}$ and $RS_{BTR}$ are equivalent. The Pearson correlation coefficient (r) for each scatter plot is computed and reported in the plot.

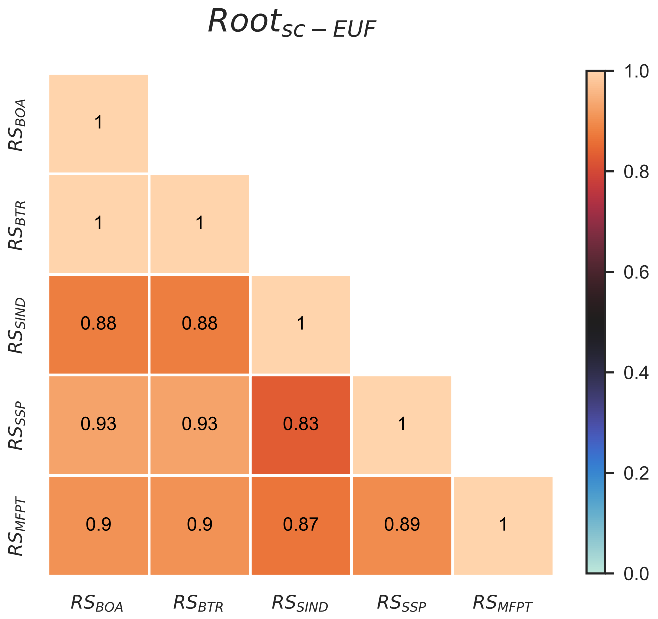

FIG. S20. **Pearson correlation between different pairs of relative stability measures for the ensemble $Root_{sc-EUF}$.** The rows and columns correspond to choices for the relative stability measures. These 5 measures are based on size of basin of attraction ($RS_{BOA}$), basin transition rates ($RS_{BTR}$), a stability index ($RS_{SIND}$), steady state probabilities ($RS_{SSP}$) and mean first passage times ($RS_{MFPT}$). The heatmap indicates the value of the Pearson correlation coefficient between pairs of these measures. Note that these measures are computed by exact means across all pairs of biological fixed points, for all 36600 models in this ensemble $Root_{sc-EUF}$ using a noise intensity parameter value of 1%. The upper triangular portion of the heatmap is not displayed as the heatmap entries constitute a symmetric matrix. Furthermore, $RS_{BOA}$ and $RS_{BTR}$ are perfectly correlated, an observation which we prove theoretically by showing that $RS_{BOA}$ and $RS_{BTR}$ are in fact equivalent.

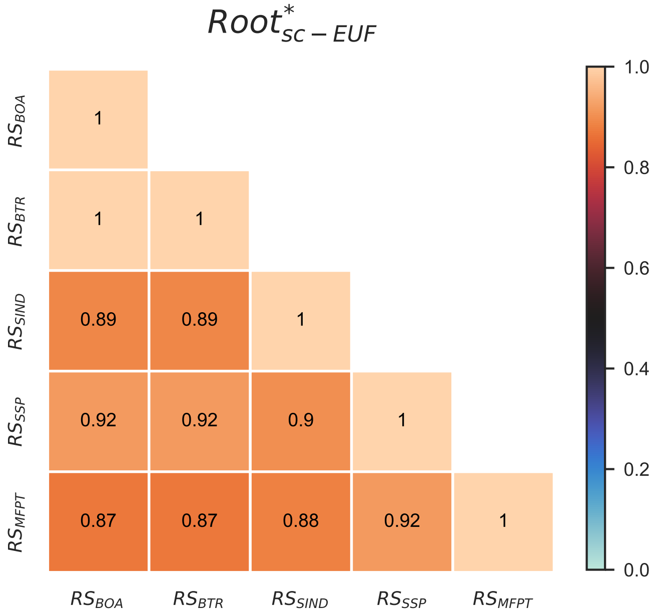

FIG. S21. **Pearson correlation between different pairs of relative stability measures for the ensemble $Root^{*}_{sc-EUF}$.** The rows and columns correspond to choices for the relative stability measures. These 5 measures are based on size of basin of attraction ($RS_{BOA}$), basin transition rates ($RS_{BTR}$), a stability index ($RS_{SIND}$), steady state probabilities ($RS_{SSP}$) and mean first passage times ($RS_{MFPT}$). The heatmap indicates the value of the Pearson correlation coefficient between pairs of these measures. Note that these measures are computed by exact means across all pairs of biological fixed points, for all 1400 models in this ensemble $Root^{*}_{sc-EUF}$ using a noise intensity parameter value of 1%. The upper triangular portion of the heatmap is not displayed as the heatmap entries constitute a symmetric matrix. Furthermore, $RS_{BOA}$ and $RS_{BTR}$ are perfectly correlated, an observation which we prove theoretically by showing that $RS_{BOA}$ and $RS_{BTR}$ are in fact equivalent.

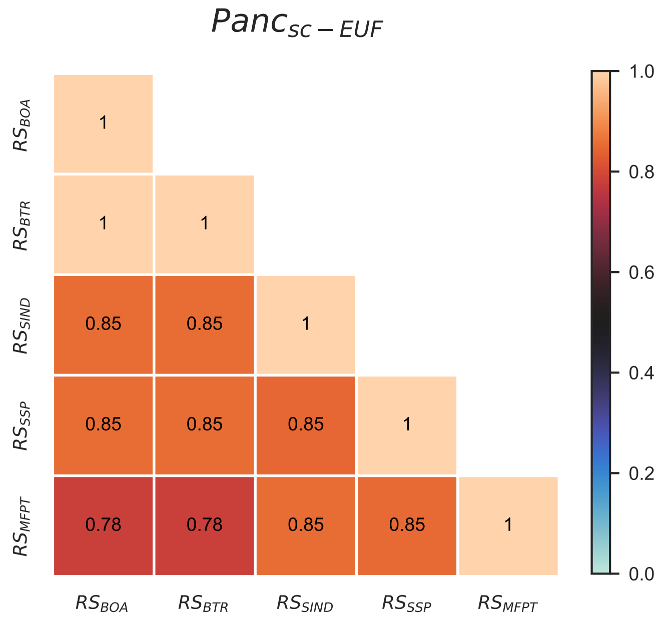

FIG. S22. **Pearson correlation between different pairs of relative stability measures for the ensemble $\boldsymbol{Panc_{sc-EUF}}$** The rows and columns correspond to choices for the relative stability measures. These 5 measures are based on size of basin of attraction ($RS_{BOA}$), basin transition rates ($RS_{BTR}$), a stability index ($RS_{SIND}$), steady state probabilities ($RS_{SSP}$) and mean first passage times ($RS_{MFPT}$). The heatmap indicates the value of the Pearson correlation coefficient between pairs of these measures. Note that these measures are computed by exact means across all pairs of biological fixed points, for all 7056 models in this ensemble $Panc_{sc-EUF}$ using a noise intensity parameter value of 1%. The upper triangular portion of the heatmap is not displayed as the heatmap entries constitute a symmetric matrix. Furthermore, $RS_{BOA}$ and $RS_{BTR}$ are perfectly correlated, an observation which we prove theoretically by showing that $RS_{BOA}$ and $RS_{BTR}$ are in fact equivalent.

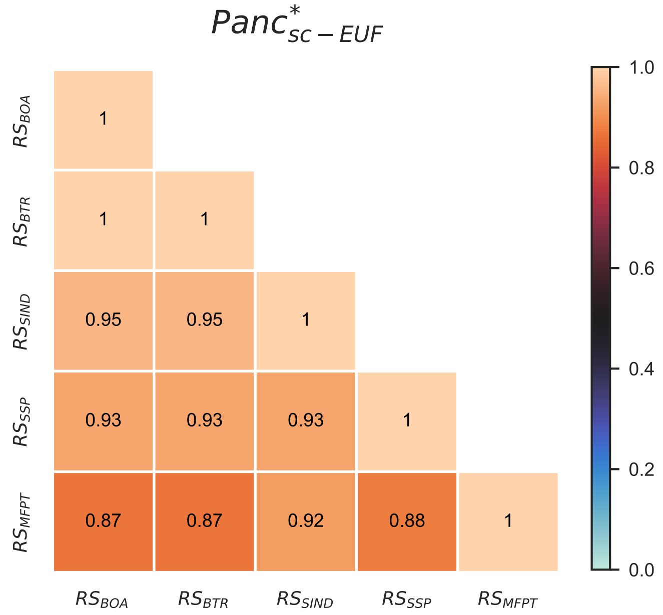

FIG. S23. **Pearson correlation between different pairs of relative stability measures for the ensemble $\boldsymbol{Panc^*_{sc-EUF}}$**. The rows and columns correspond to choices for the relative stability measures. These 5 measures are based on size of basin of attraction ($RS_{BOA}$), basin transition rates ($RS_{BTR}$), a stability index ($RS_{SIND}$), steady state probabilities ($RS_{SSP}$) and mean first passage times ($RS_{MFPT}$). The heatmap indicates the value of the Pearson correlation coefficient between pairs of these measures. Note that these measures are computed by exact means across all pairs of biological fixed points, for all 159 models in this ensemble $Panc^*_{sc-EUF}$ using a noise intensity parameter value of 1%. The upper triangular portion of the heatmap is not displayed as the heatmap entries constitute a symmetric matrix. Furthermore, $RS_{BOA}$ and $RS_{BTR}$ are perfectly correlated, an observation which we prove theoretically by showing that $RS_{BOA}$ and $RS_{BTR}$ are in fact equivalent.

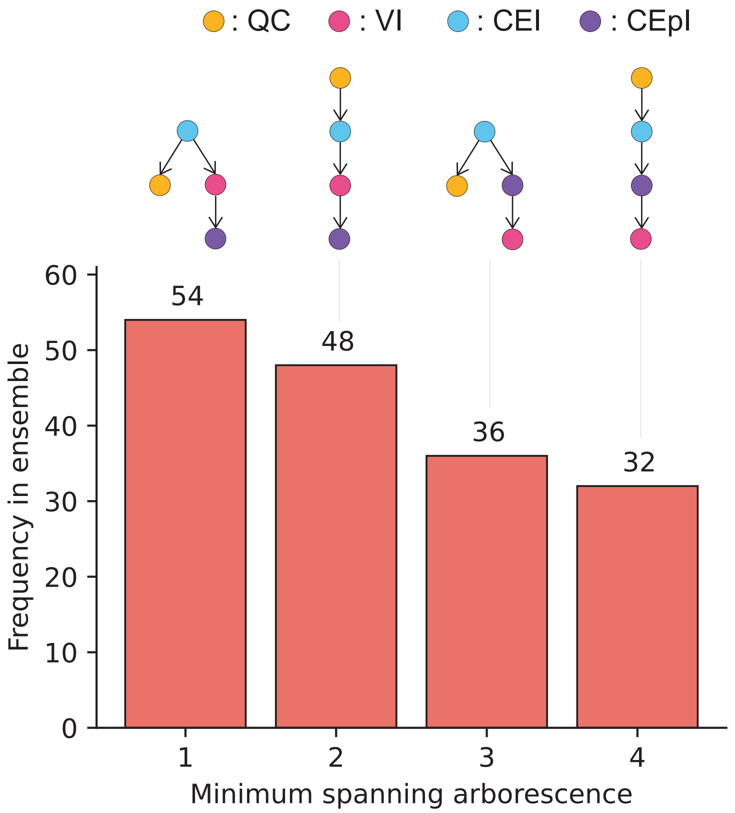

FIG. S24. **Frequency distribution of the minimum spanning arborescences (MSA) for the ensemble $\boldsymbol{Root^*_{sc-NCF}}$.** The MSA for a Boolean model is constructed from a complete digraph whose vertices are biological fixed points and directed edges are the MFPTs (see Methods for more details). The $x$ axis labels the different MSAs that occur in the ensemble $Root^*_{sc-NCF}$. Of the 64 possible (labeled and oriented) trees for 4 fixed points, only 4 occur in the $Root^*_{sc-NCF}$. The $y$ axis is the frequency of each of these trees among the 170 models of the $Root^*_{sc-NCF}$ ensemble. The biological fixed points of the $Root^*_{sc-NCF}$ ensemble are as follows. QC: Quiescent center, VI: Vascular initials, CEI: Cortex-Endodermis initials and CEpI: Columella epidermis initials.

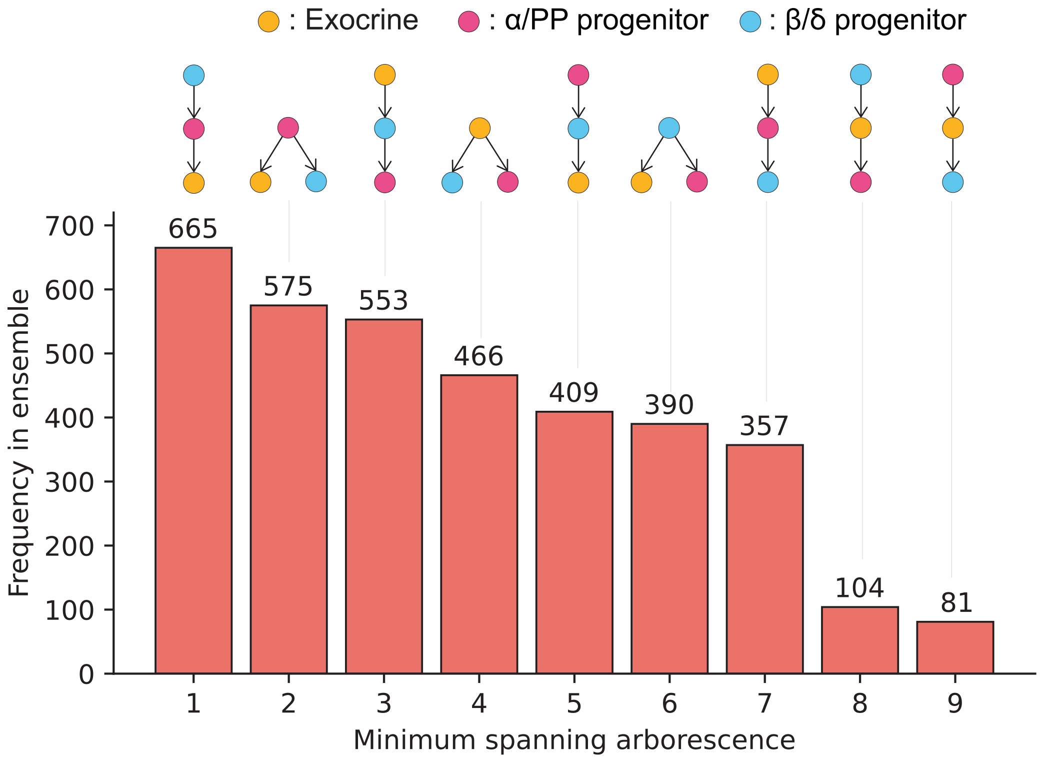

FIG. S25. **Frequency distribution of the minimum spanning arborescences (MSA) for the ensemble $\boldsymbol{Panc_{sc-NCF}}$.** The MSA for a Boolean model is constructed from a complete digraph whose vertices are biological fixed points and directed edges are the MFPTs (see Methods for more details). The $x$ axis labels the different MSAs that occur in the ensemble $Panc_{sc-NCF}$. Of the 9 possible (labeled and oriented) trees for 3 fixed points, all 9 occur in the $Panc_{sc-NCF}$. The $y$ axis is the frequency of each of these trees among the 3600 models of the $Panc_{sc-NCF}$ ensemble.

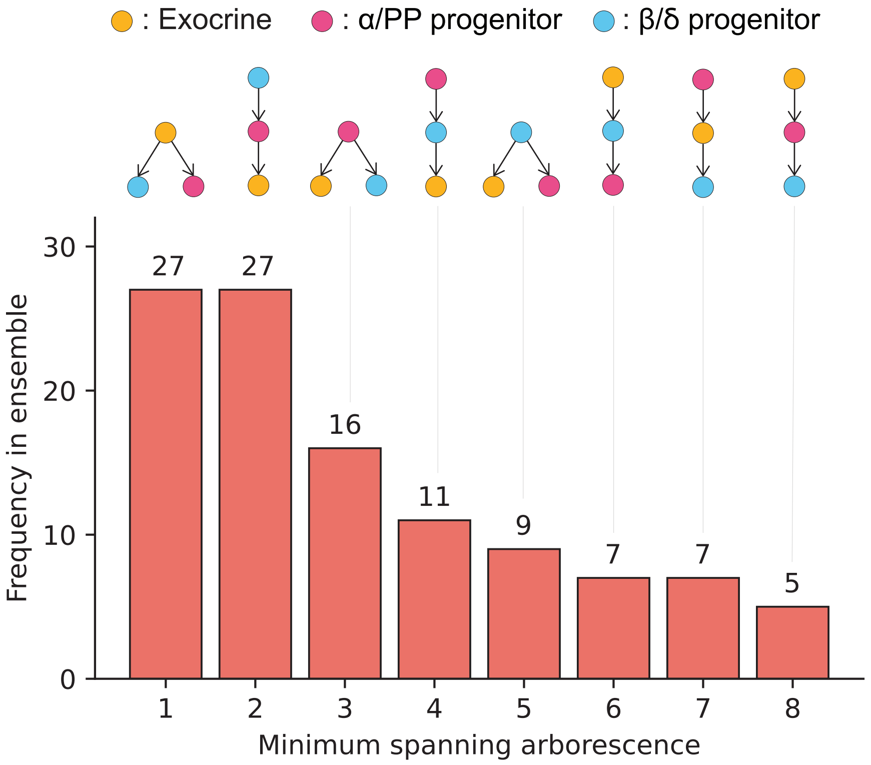

FIG. S26. **Frequency distribution of the minimum spanning arborescences (MSA) for the ensemble $Panc^*_{sc-NCF}$.** The MSA for a Boolean model is constructed from a complete digraph whose vertices are biological fixed points and directed edges are the MFPTs (see Methods for more details). The $x$ axis labels the different MSAs that occur in the ensemble $Panc^*_{sc-NCF}$. Of the 9 possible (labeled and oriented) trees for 3 fixed points, only 8 occur in the $Panc^*_{sc-NCF}$. The $y$ axis is the frequency of each of these trees among the 109 models of the $Panc^*_{sc-NCF}$ ensemble.

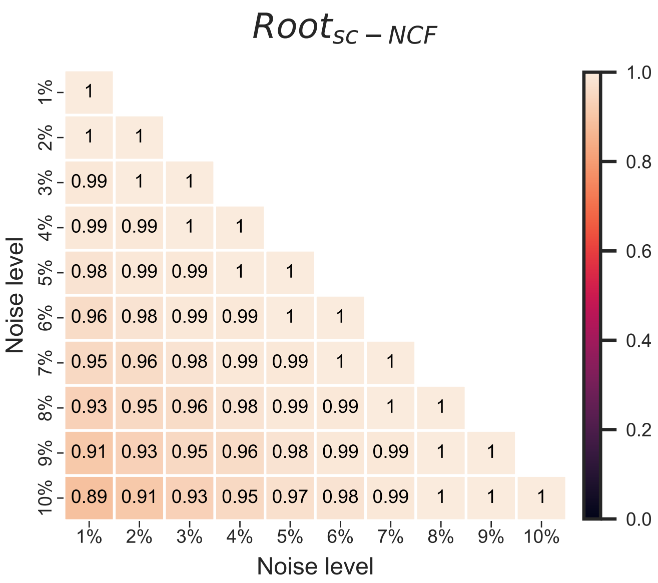

| | 1% | 2% | 3% | 4% | 5% | 6% | 7% | 8% | 9% | 10% |
|---|---|---|---|---|---|---|---|---|---|---|
| 1% | 1 | | | | | | | | | |
| 2% | 1 | 1 | | | | | | | | |
| 3% | 0.99 | 1 | 1 | | | | | | | |
| 4% | 0.99 | 0.99 | 1 | 1 | | | | | | |
| 5% | 0.98 | 0.99 | 0.99 | 1 | 1 | | | | | |
| 6% | 0.96 | 0.98 | 0.99 | 0.99 | 1 | 1 | | | | |
| 7% | 0.95 | 0.96 | 0.98 | 0.99 | 0.99 | 1 | 1 | | | |
| 8% | 0.93 | 0.95 | 0.96 | 0.98 | 0.99 | 0.99 | 1 | 1 | | |
| 9% | 0.91 | 0.93 | 0.95 | 0.96 | 0.98 | 0.99 | 0.99 | 1 | 1 | |
| 10% | 0.89 | 0.91 | 0.93 | 0.95 | 0.97 | 0.98 | 0.99 | 1 | 1 | 1 |

FIG. S27. **Pearson correlation between $RS_{MFPT}$ values computed by exact methods for different pairs of noise values for the ensemble $Root_{sc-NCF}$.** Rows and columns correspond to the noise intensities ranging from 1% to 10%. The heatmap gives the value of the Pearson correlation coefficient of $RS_{MFPT}$ values when considering all pairs of biological fixed points and all 1275 models within the ensemble $Root_{sc-NCF}$, for different pairs of noise intensities. The upper triangular portion of the heatmap is not displayed because it constitutes a symmetric matrix. The correlation between the $RS_{MFPT}$ for different values of noise is found to be very strong even for pairs of noise values which have a large difference.

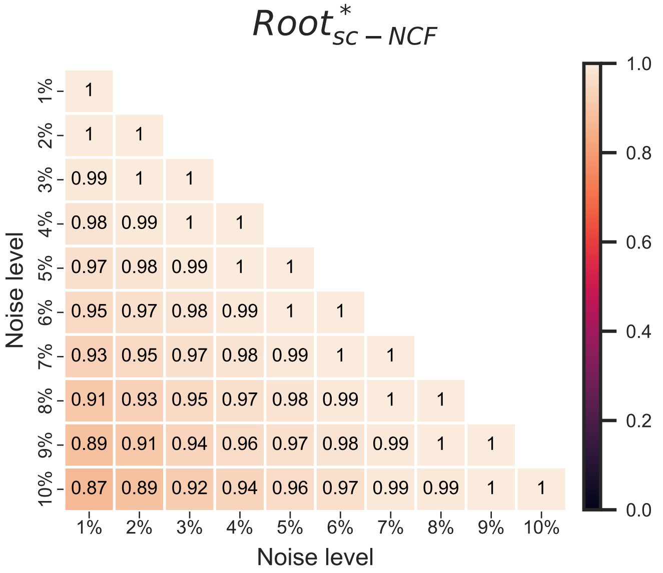

FIG. S28. **Pearson correlation between $RS_{MFPT}$ values computed by exact methods for different pairs of noise values for the ensemble $\boldsymbol{Root^*_{sc-NCF}}$.** Rows and columns correspond to the noise intensities ranging from 1% to 10%. The heatmap gives the value of the Pearson correlation coefficient of $RS_{MFPT}$ values when considering all pairs of biological fixed points and all 170 models within the ensemble $Root^*_{sc-NCF}$, for different pairs of noise intensities. The upper triangular portion of the heatmap is not displayed because it constitutes a symmetric matrix. The correlation between the $RS_{MFPT}$ for different values of noise is found to be very strong even for pairs of noise values which have a large difference.

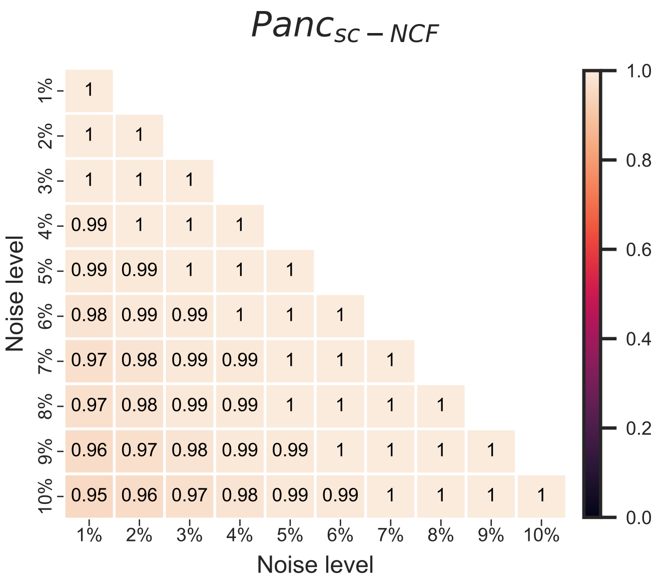

FIG. S29. **Pearson correlation between $RS_{MFPT}$ values computed by exact methods for different pairs of noise values for the ensemble $\boldsymbol{Panc_{sc-NCF}}$.** Rows and columns correspond to the noise intensities ranging from 1% to 10%. The heatmap gives the value of the Pearson correlation coefficient of $RS_{MFPT}$ values when considering all pairs of biological fixed points and all 3600 models within the ensemble $Panc_{sc-NCF}$, for different pairs of noise intensities. The upper triangular portion of the heatmap is not displayed because it constitutes a symmetric matrix. The correlation between the $RS_{MFPT}$ for different values of noise is found to be very strong even for pairs of noise values which have a large difference.

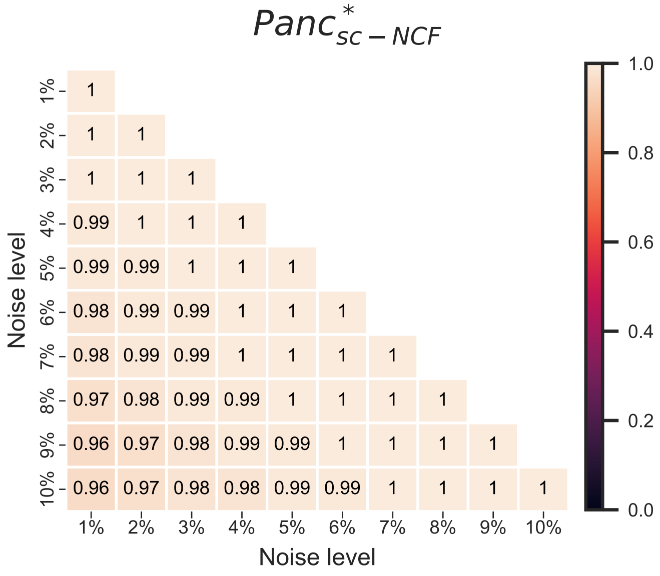

FIG. S30. **Pearson correlation between $RS_{MFPT}$ values computed by exact methods for different pairs of noise values for the ensemble $\boldsymbol{Panc^*_{sc-NCF}}$.** Rows and columns correspond to the noise intensities ranging from 1% to 10%. The heatmap gives the value of the Pearson correlation coefficient of $RS_{MFPT}$ values when considering all pairs of biological fixed points and all 109 models within the ensemble $Panc^*_{sc-NCF}$, for different pairs of noise intensities. The upper triangular portion of the heatmap is not displayed because it constitutes a symmetric matrix. The correlation between the $RS_{MFPT}$ for different values of noise is found to be very strong even for pairs of noise values which have a large difference.

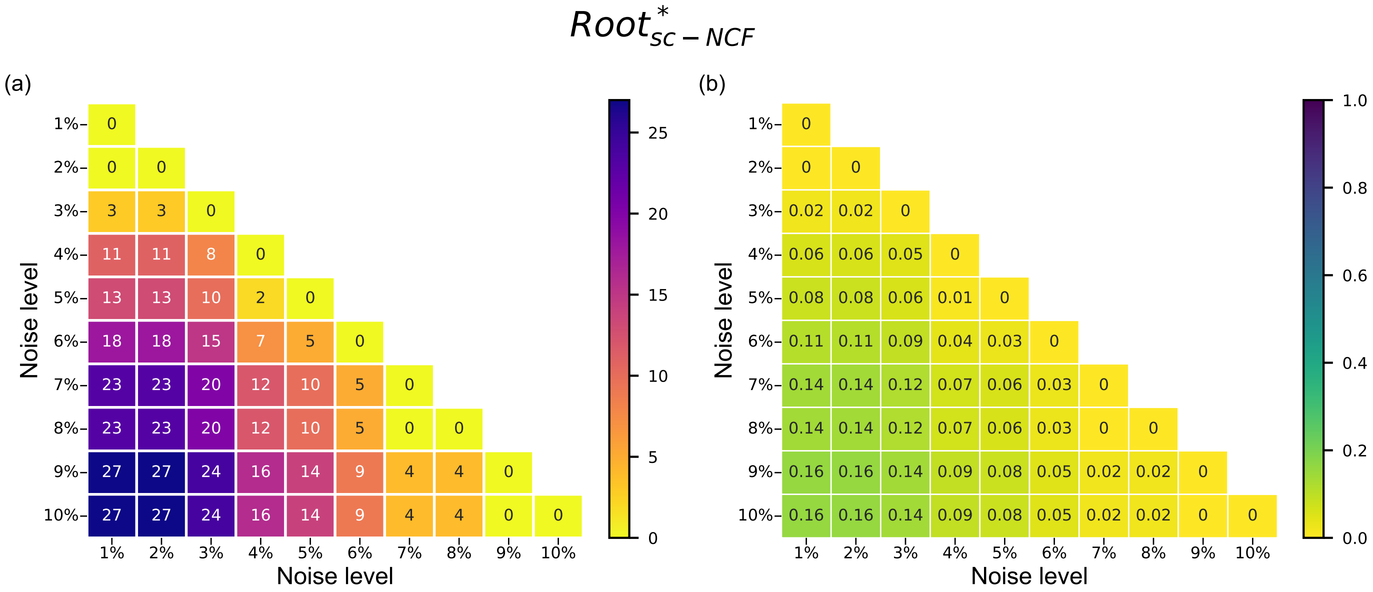

FIG. S31. **Number and fraction of models which differ in at least one comparison of partial ordering of the different biological fixed points when considering two different noise values, in the ensemble $\boldsymbol{Root^*_{sc-NCF}}$.** The (partial) order of two fixed point is specified via the MFPT values for going from one to the other, computed here using an exact method. Rows and columns correspond to the noise intensity. The heatmap (a) gives the number of models (out of a total of 170 models in the ensemble $Root^*_{sc-NCF}$) that differ in at least one (partial) order across pairs of biological fixed points. The heatmap (b) provides the same information but using the fraction of such models. The upper triangular portions of the heatmaps are not displayed because they constitute a symmetric matrix.

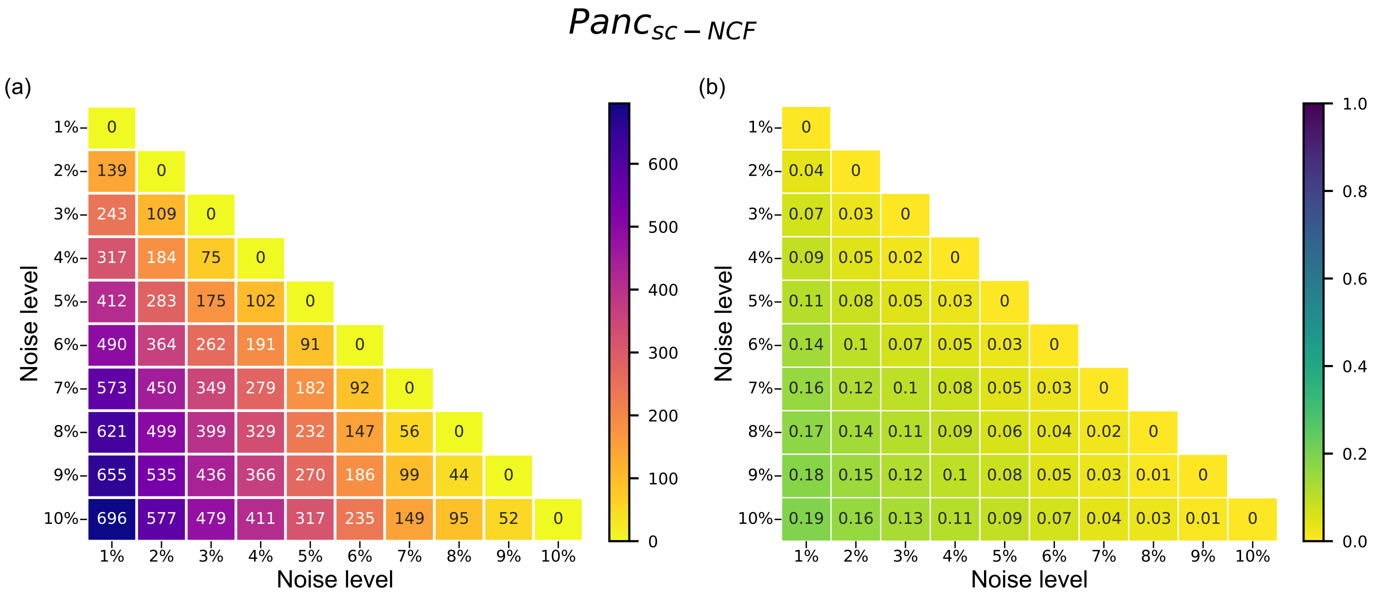

FIG. S32. **Number and fraction of models which differ in at least one comparison of partial ordering of the different biological fixed points when considering two different noise values, in the ensemble $Panc_{sc-NCF}$.** The (partial) order of two fixed point is specified via the MFPT values for going from one to the other, computed here using an exact method. Rows and columns correspond to the noise intensity. The heatmap (a) gives the number of models (out of a total of 3600 models in the ensemble $Panc_{sc-NCF}$) that differ in at least one (partial) order across pairs of biological fixed points. The heatmap (b) provides the same information but using the fraction of such models. The upper triangular portions of the heatmaps are not displayed because they constitute a symmetric matrix.

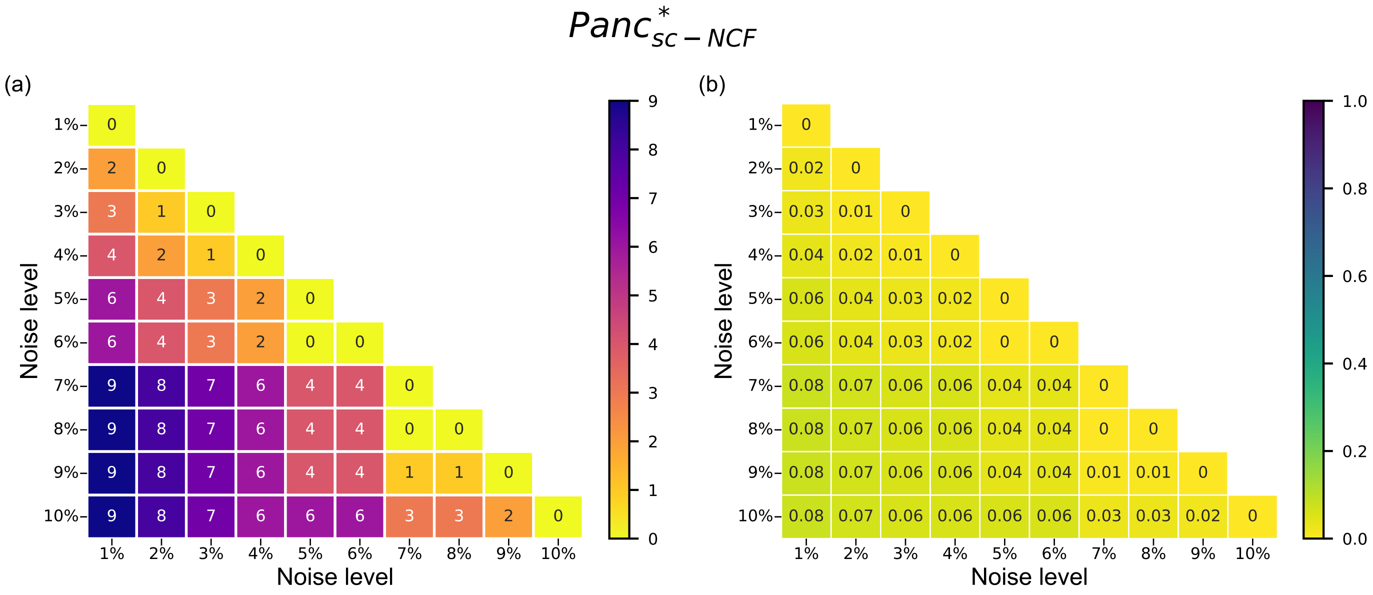

FIG. S33. **Number and fraction of models which differ in at least one comparison of partial ordering of the different biological fixed points when considering two different noise values, in the ensemble $Panc^{*}_{sc-NCF}$.** The (partial) order of two fixed point is specified via the MFPT values for going from one to the other, computed here using an exact method. Rows and columns correspond to the noise intensity. The heatmap (a) gives the number of models (out of a total of 109 models in the ensemble $Panc^{*}_{sc-NCF}$) that differ in at least one (partial) order across pairs of biological fixed points. The heatmap (b) provides the same information but using the fraction of such models. The upper triangular portions of the heatmaps are not displayed because they constitute a symmetric matrix.

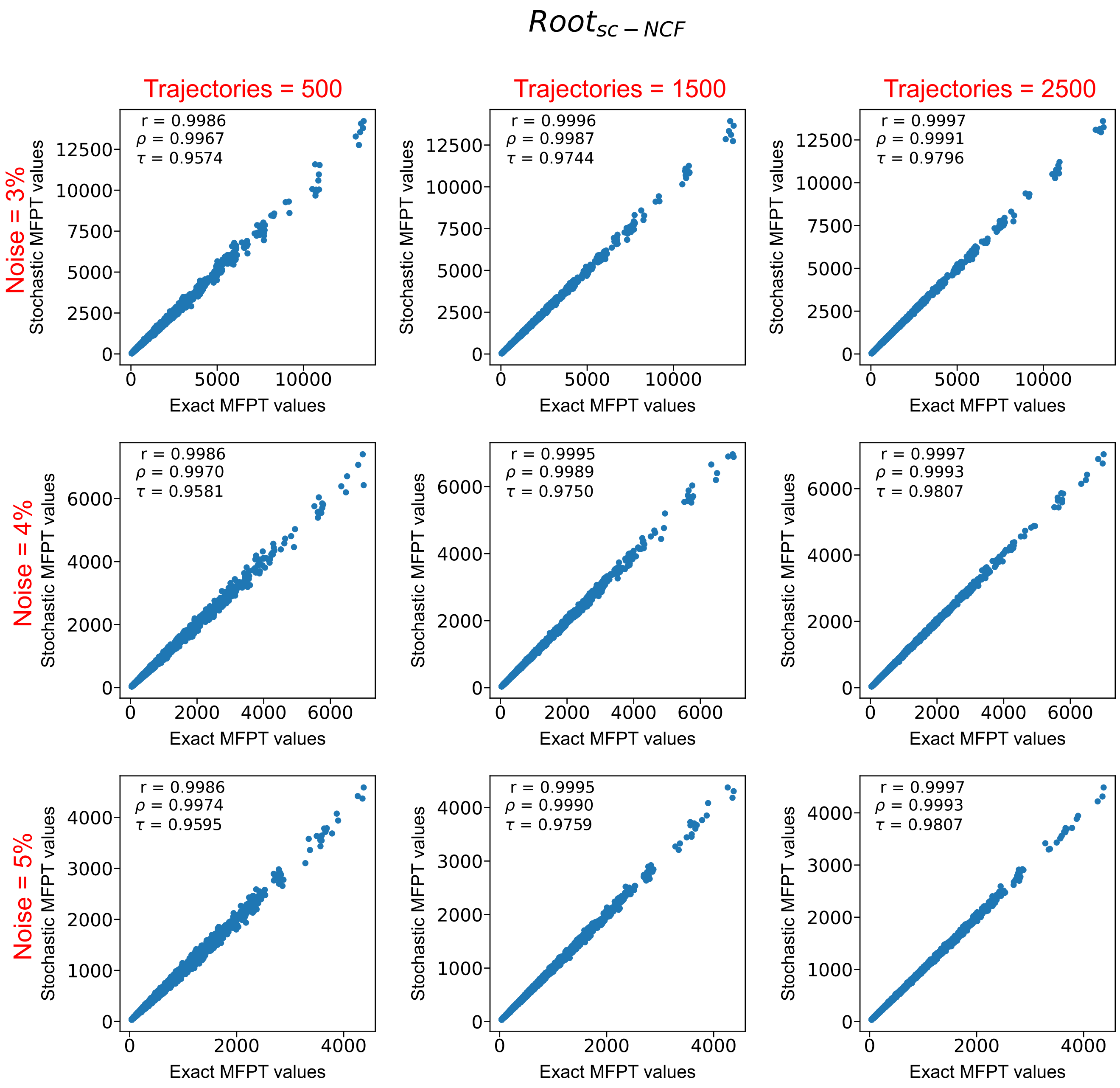

FIG. S34. **Correlation between the MFPT obtained via the exact method versus the proposed stochastic method using the ensemble $Root_{sc-NCF}$.** The $x$ and $y$ axis of all scatter plots represent the MFPT from the biological fixed point $v$ to the biological fixed point $u$ (denoted by $M_{uv}$) computed via exact and stochastic means respectively, for all pairs of fixed points and for all 1275 models belonging to the ensemble $Root_{sc-NCF}$. Each scatter plot is generated for a particular noise (3%, 4% or 5%) and number of trajectories (500, 1500 or 2500) going from fixed point $v$ to fixed point $u$. The exact and stochastic MFPT values are strongly correlated as can be seen from the 3 measures of correlation, namely, Pearson correlation coefficient (r), Spearman rank correlation ($\rho$) and Kendall rank correlation ($\tau$). It can be seen that at a fixed noise as the number of trajectories are increased from 500 to 2500, the correlation becomes stronger across all 3 correlation measures.

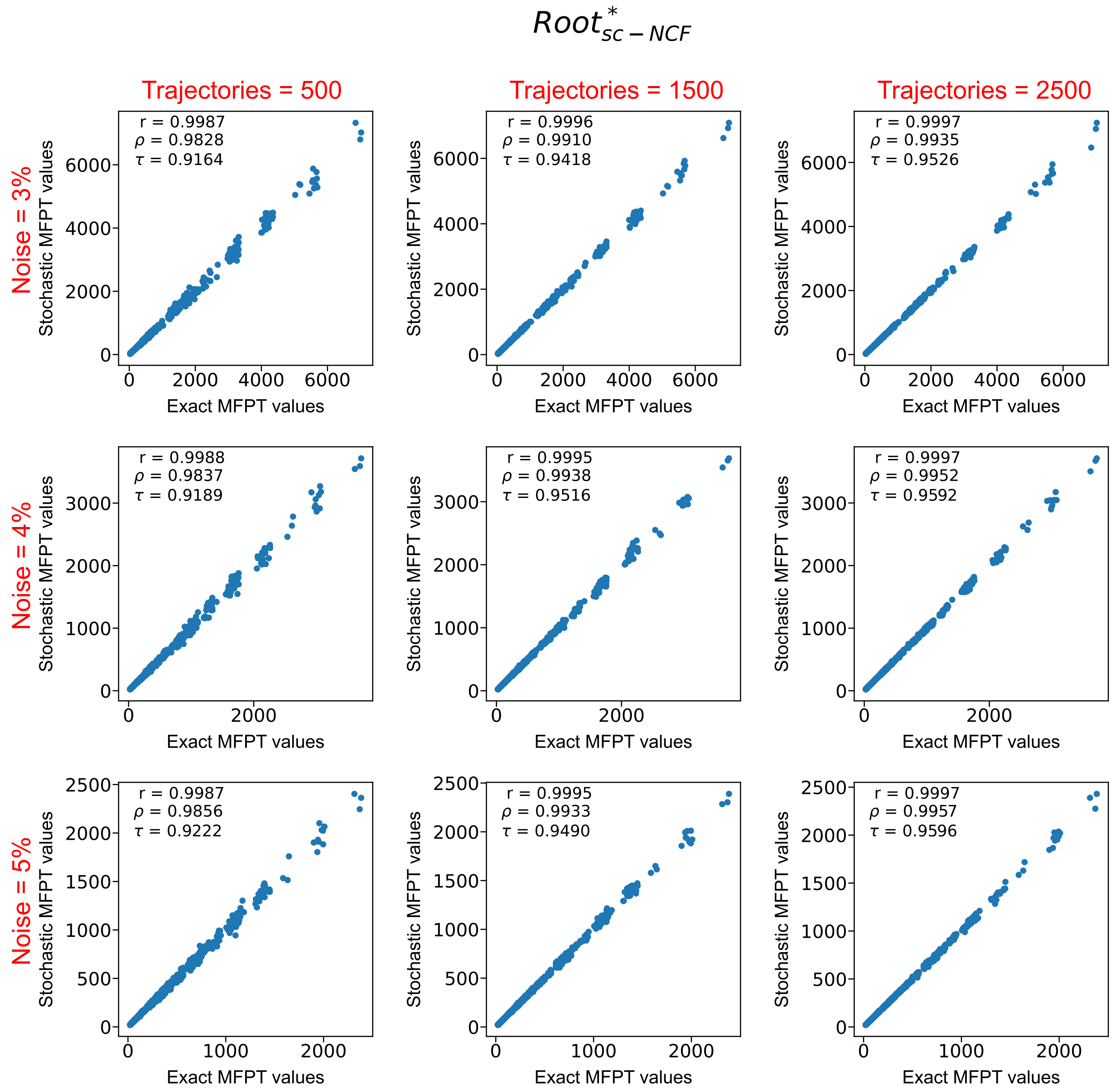

FIG. S35. **Correlation between the MFPT obtained via the exact method versus the proposed stochastic method using the ensemble $Root^*_{sc-NCF}$.** The $x$ and $y$ axis of all scatter plots represent the MFPT from the biological fixed point $v$ to the biological fixed point $u$ (denoted by $M_{uv}$) computed via exact and stochastic means respectively, for all pairs of fixed points and for all 170 models belonging to the ensemble $Root^*_{sc-NCF}$. Each scatter plot is generated for a particular noise (3%, 4% or 5%) and number of trajectories (500, 1500 or 2500) going from fixed point $v$ to fixed point $u$. The exact and stochastic MFPT values are strongly correlated as can be seen from the 3 measures of correlation, namely, Pearson correlation coefficient (r), Spearman rank correlation ($\rho$) and Kendall rank correlation ($\tau$). It can be seen that at a fixed noise as the number of trajectories are increased from 500 to 2500, the correlation becomes stronger across all 3 correlation measures.

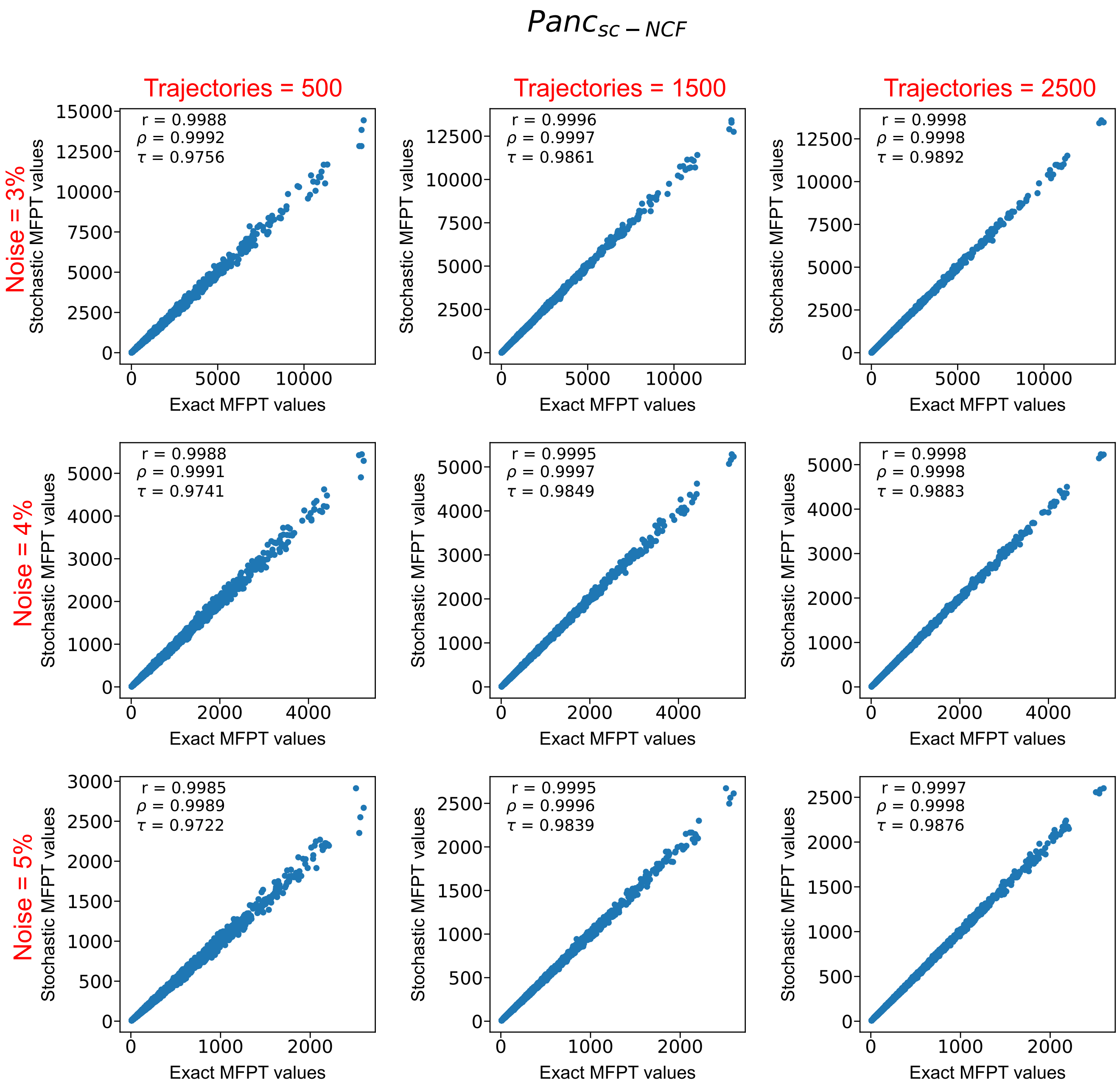

FIG. S36. **Correlation between the MFPT obtained via the exact method versus the proposed stochastic method using the ensemble $Panc_{sc-NCF}$.** The $x$ and $y$ axis of all scatter plots represent the MFPT from the biological fixed point $v$ to the biological fixed point $u$ (denoted by $M_{uv}$) computed via exact and stochastic means respectively, for all pairs of fixed points and for all 3600 models belonging to the ensemble $Panc_{sc-NCF}$. Each scatter plot is generated for a particular noise (3%, 4% or 5%) and number of trajectories (500, 1500 or 2500) going from fixed point $v$ to fixed point $u$. The exact and stochastic MFPT values are strongly correlated as can be seen from the 3 measures of correlation, namely, Pearson correlation coefficient (r), Spearman rank correlation ($\rho$) and Kendall rank correlation ($\tau$). It can be seen that at a fixed noise as the number of trajectories are increased from 500 to 2500, the correlation becomes stronger across all 3 correlation measures.

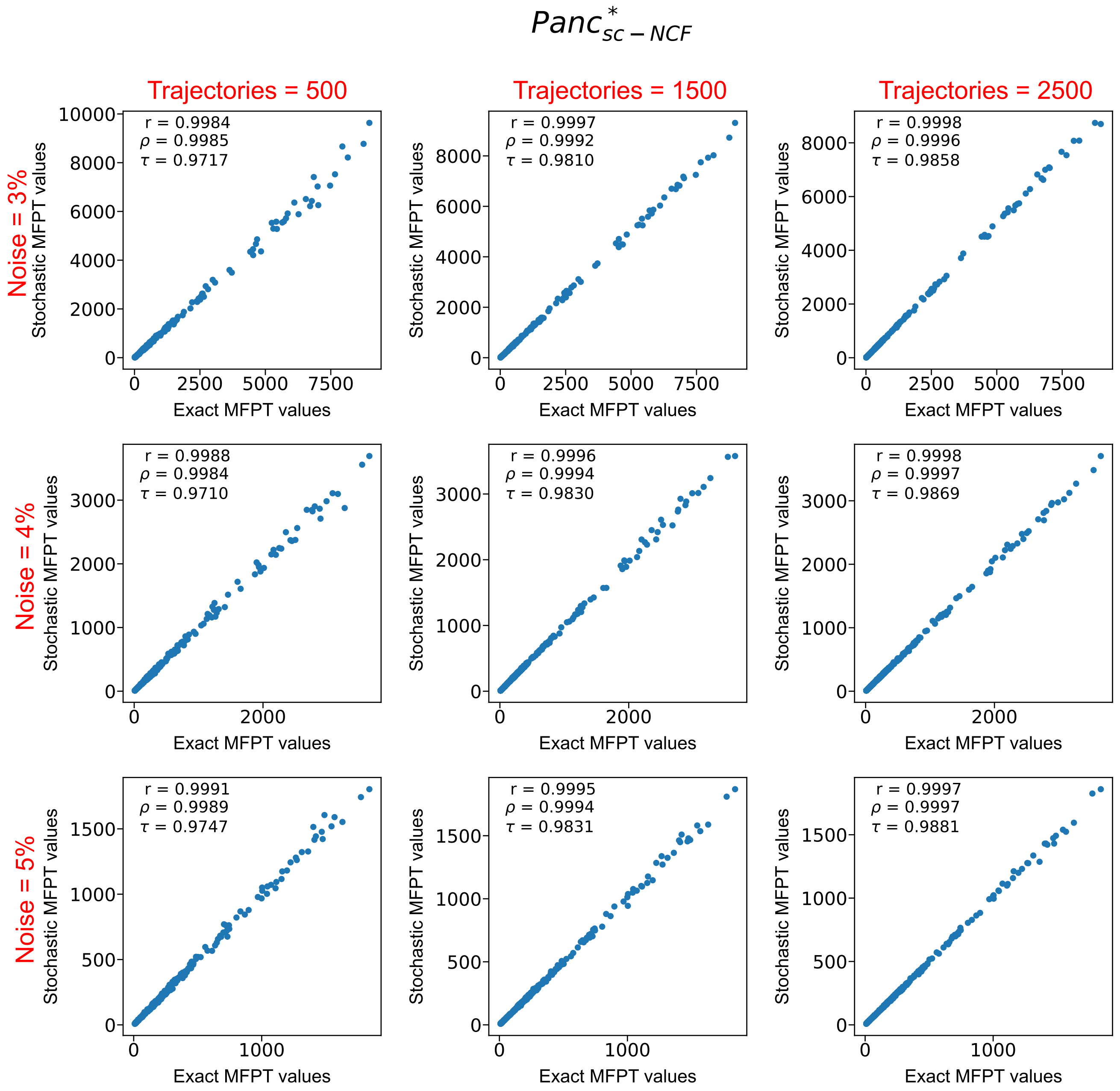

FIG. S37. **Correlation between the MFPT obtained via the exact method versus the proposed stochastic method using the ensemble $Panc^*_{sc-NCF}$.** The $x$ and $y$ axis of all scatter plots represent the MFPT from the biological fixed point $v$ to the biological fixed point $u$ (denoted by $M_{uv}$) computed via exact and stochastic means respectively, for all pairs of fixed points and for all 109 models belonging to the ensemble $Panc^*_{sc-NCF}$. Each scatter plot is generated for a particular noise (3%, 4% or 5%) and number of trajectories (500, 1500 or 2500) going from fixed point $v$ to fixed point $u$. The exact and stochastic MFPT values are strongly correlated as can be seen from the 3 measures of correlation, namely, Pearson correlation coefficient (r), Spearman rank correlation ($\rho$) and Kendall rank correlation ($\tau$). It can be seen that at a fixed noise as the number of trajectories are increased from 500 to 2500, the correlation becomes stronger across all 3 correlation measures.

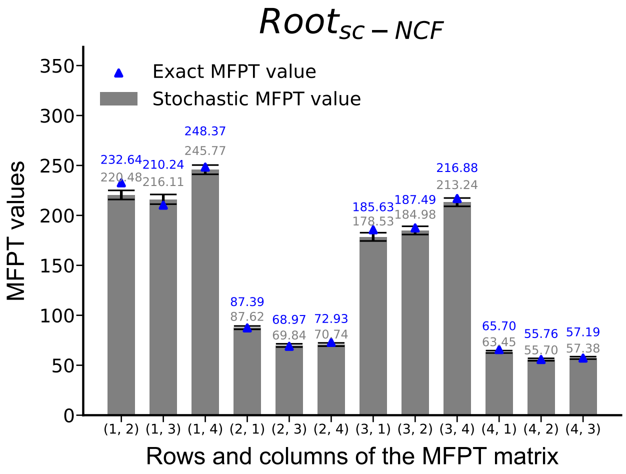

FIG. S38. **Barplot of the mean first passage time (MFPT) from one biological fixed point to another, computed via the stochastic method for a model taken from the ensemble $\boldsymbol{Root_{sc-NCF}}$.** The $x$-axis labels the rows and columns of the MFPT matrix entries, so for instance $(1, 3)$ denotes the case of matrix element $M_{13}$ when going from fixed point 3 to fixed point 1. The numbering of the fixed points are as follows. 1: Quiescent center (QC), 2: Vascular initials (VI), 3: Cortex-Endodermis initials (CEI) and 4: Columella epidermis initials (CEpI). The $y$-axis represents the associated mean first passage time (MFPT). It is computed following the stochastic approach as described in Methods, averaging over 2500 different trajectories of the dynamics starting from one fixed point and stopping as soon as the other fixed point is reached when using the rules for a particular Boolean model in the ensemble $Root_{sc-NCF}$, at 5% noise level. The tiny error bars indicate that the statistical error in the estimation of the MFPT value is very low. For comparison, the MFPT values obtained via the exact method are displayed via blue triangles (numerical values are provided above the bars in blue). The MFPT obtained via the proposed stochastic method is very close to that obtained via exact means. The grouping of bars according to the target fixed point visually illustrates the ease or difficulty of reaching a particular biological fixed point from the other three. For instance, in this particular Boolean model, it is difficult to go to the QC starting from any other fixed point and it is relatively easy to go to the CEpI starting from any other fixed point.

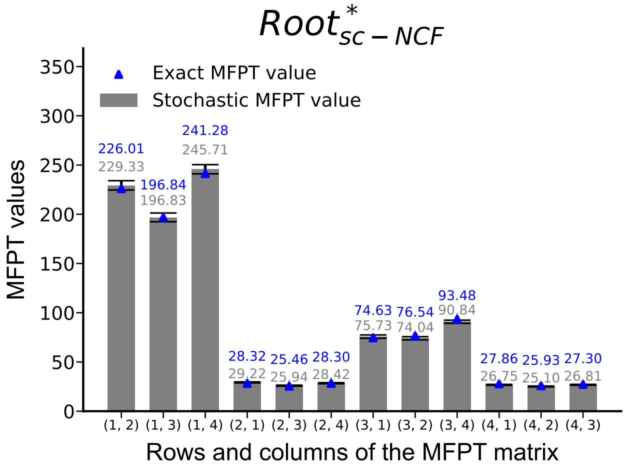

FIG. S39. **Barplot of the mean first passage time (MFPT) from one biological fixed point to another, computed via stochastic methods for a model specific to the ensemble $Root^*_{sc-NCF}$.** The $x$-axis labels the rows and columns of the MFPT matrix entries, so for instance $(1, 3)$ denotes the case of matrix element $M_{13}$ when going from fixed point 3 to fixed point 1. The numbering of the fixed points are as follows. 1: Quiescent center (QC), 2: Vascular initials (VI), 3: Cortex-Endodermis initials (CEI) and 4: Columella epidermis initials (CEpI). The $y$-axis represents the associated mean first passage time (MFPT). It is computed following our stochastic approach as described in Methods, averaging over 2500 different trajectories of the dynamics starting from one fixed point and stopping as soon as the other fixed point is reached when using the rules for a particular Boolean model in the ensemble $Root^*_{sc-NCF}$, at 5% noise level. The tiny error bars indicate that the statistical error in the estimation of the MFPT value is very low. For comparison, the MFPT values obtained via the exact method are displayed via blue triangles (numerical values are provided above the bars in blue). The MFPT obtained via the proposed stochastic method is very close to that obtained via exact means. The grouping of bars with similar height visually illustrates the difficulty of reaching a particular biological fixed point from any other one. For instance, in this particular Boolean model, it is difficult to go to the QC starting from any other fixed point and it is quite easy to go to the CEpI starting from any other fixed point.

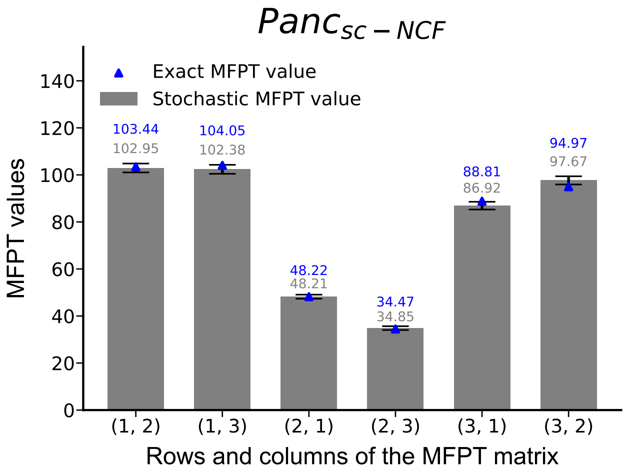

FIG. S40. **Barplot of the mean first passage time (MFPT) from one biological fixed point to another, computed via stochastic methods for a model specific to the ensemble $\boldsymbol{Panc_{sc-NCF}}$.** The $x$-axis labels the rows and columns of the MFPT matrix entries, so for instance $(1, 3)$ denotes the case of matrix element $M_{13}$ when going from fixed point 3 to fixed point 1. The numbering of the fixed points are as follows. 1: Exocrine, 2: $\beta/\delta$ progenitor, 3: $\alpha$/PP progenitor. The $y$-axis represents the associated mean first passage time (MFPT). It is computed following our stochastic approach as described in Methods, averaging over 2500 different trajectories of the dynamics starting from one fixed point and stopping as soon as the other fixed point is reached when using the rules for a particular Boolean model in the ensemble $Panc_{sc-NCF}$, at 5% noise level. The tiny error bars indicate that the statistical error in the estimation of the MFPT value is very low. For comparison, the MFPT values obtained via the exact method are displayed via blue triangles (numerical values are provided above the bars in blue). The MFPT obtained via the proposed stochastic method is very close to that obtained via exact means. The grouping of bars with similar height visually illustrates the difficulty of reaching a particular biological fixed point from any other one. For instance, in this particular Boolean model, it is difficult to go to the Exocrine fixed point starting from any other fixed point and it is relatively easier to go to the $\beta/\delta$ progenitor starting from any other fixed point.

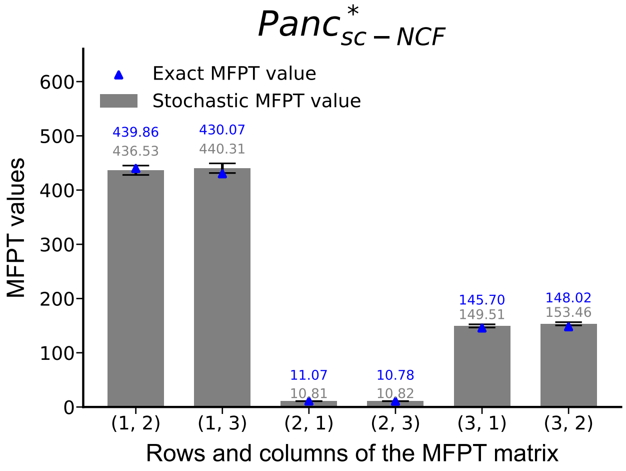

FIG. S41. **Barplot of the mean first passage time (MFPT) from one biological fixed point to another, computed via stochastic methods for a model specific to the ensemble $Panc^*_{sc-NCF}$.** The $x$-axis labels the rows and columns of the MFPT matrix entries, so for instance $(1, 3)$ denotes the case of matrix element $M_{13}$ when going from fixed point 3 to fixed point 1. The numbering of the fixed points are as follows. 1: Exocrine, 2: $\beta/\delta$ progenitor, 3: $\alpha$/PP progenitor. The $y$-axis represents the associated mean first passage time (MFPT). It is computed following our stochastic approach as described in Methods, averaging over 2500 different trajectories of the dynamics starting from one fixed point and stopping as soon as the other fixed point is reached when using the rules for a particular Boolean model in the ensemble $Panc^*_{sc-NCF}$, at 5% noise level. The tiny error bars indicate that the statistical error in the estimation of the MFPT value is very low. For comparison, the MFPT values obtained via the exact method are displayed via blue triangles (numerical values are provided above the bars in blue). The MFPT obtained via the proposed stochastic method is very close to that obtained via exact means. The grouping of bars with similar height visually illustrates the difficulty of reaching a particular biological fixed point from any other one. For instance, in this particular Boolean model, it is difficult to go to the Exocrine fixed point starting from any other fixed point and it is relatively easier to go to the $\beta/\delta$ progenitor starting from any other fixed point.

**(a) Gene regulatory network and attractors of *Arabidopsis thaliana* root stem cell niche** (Azpeitia *et al. 2013*)

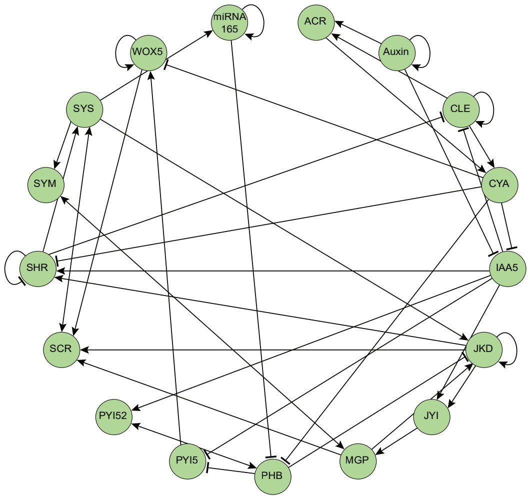

| | SHR | SCR | JKD | MGP | miRNA 165 | PHB | Auxin | IAA5 | WOX5 | CLE | ACR | SYS | CYA | PYI5 | JYI | SYM | PYI52 |
|---|---|---|---|---|---|---|---|---|---|---|---|---|---|---|---|---|---|
| LCC | 0 | 0 | 0 | 0 | 0 | 0 | 1 | 0 | 0 | 1 | 1 | 0 | 1 | 1 | 0 | 0 | 0 |
| CLEI | 0 | 0 | 0 | 0 | 1 | 0 | 1 | 0 | 0 | 1 | 1 | 0 | 1 | 1 | 0 | 0 | 0 |
| QC | 1 | 1 | 1 | 1 | 1 | 0 | 1 | 0 | 1 | 0 | 0 | 1 | 0 | 1 | 0 | 0 | 0 |
| CEI | 1 | 1 | 1 | 1 | 1 | 0 | 1 | 0 | 0 | 0 | 0 | 1 | 0 | 1 | 0 | 1 | 0 |
| Non-biological attractor | 1 | 1 | 1 | 1 | 1 | 0 | 1 | 0 | 1 | 0 | 0 | 1 | 0 | 1 | 0 | 1 | 0 |

**(b) Gene regulatory network and attractors of *Arabidopsis thaliana* root stem cell niche**
(García-Gómez *et al. 2017*)

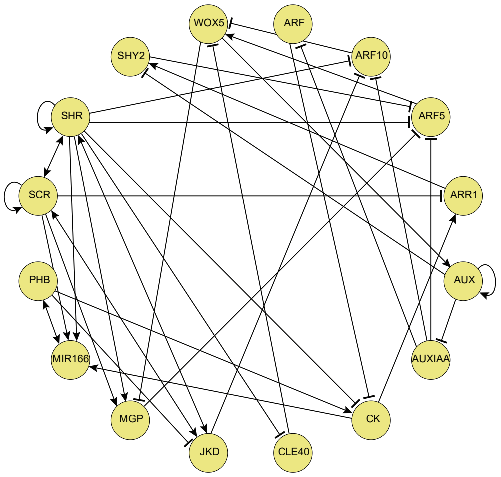

| | CK | ARR1 | SHY2 | AUXI AA | ARF | ARF10 | ARF5 | AUXIN | SCR | SHR | MIR 166 | PHB | JKD | MGP | WOX5 | CLE40 |
|---|---|---|---|---|---|---|---|---|---|---|---|---|---|---|---|---|
| QC | 0 | 0 | 0 | 0 | 1 | 0 | 1 | 1 | 1 | 1 | 1 | 0 | 1 | 0 | 1 | 0 |
| Endodermis PD | 0 | 0 | 0 | 0 | 1 | 0 | 0 | 1 | 1 | 1 | 1 | 0 | 1 | 1 | 0 | 0 |
| P. pro-vascular PD | 0 | 0 | 0 | 0 | 1 | 1 | 1 | 1 | 0 | 1 | 1 | 0 | 0 | 0 | 0 | 0 |
| C. pro-vascular PD | 0 | 0 | 0 | 0 | 1 | 1 | 1 | 1 | 0 | 1 | 0 | 1 | 0 | 0 | 0 | 0 |
| Root cap2 | 1 | 1 | 0 | 0 | 1 | 1 | 1 | 1 | 0 | 0 | 1 | 0 | 0 | 0 | 0 | 1 |

FIG. S42. **(a)** 2013 model of the *Arabidopsis thaliana* RSCN Boolean GRN and its fixed points with AUX = 1. This model has 17 nodes and 42 edges of which 5 nodes are intermediate nodes. The network is constructed using regulatory interactions obtained from the BFs of *model 4* in [27]. Here, QC: Quiescent center, CEI: Cortex-endodermis initials, LCC: Lateral root-cap and CLEI: Columella and lateral root-cap-epidermis initials. **(b)** 2017 model of the *Arabidopsis thaliana* RAM Boolean GRN and its fixed points with AUX = 1. This model has 16 nodes and 39 edges. The network is constructed using regulatory interactions obtained from the BFs of the *GHRN1 model* in [28].

### Gene regulatory network and attractors of *Arabidopsis thaliana* root stem cell niche
(García-Gómez *et al. 2020*)

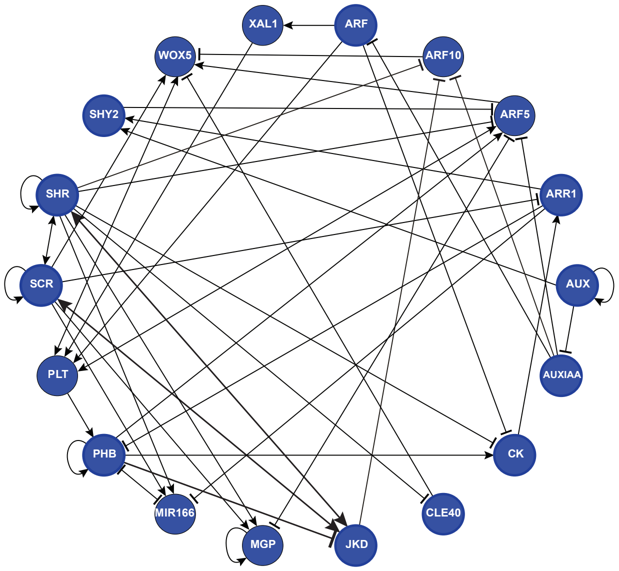

| gene | CK | ARR1 | SHY2 | AUXIAA | ARF | ARF10 | ARF5 | XAL1 | PLT | AUX | SCR | SHR | MIR166 | PHB | JKD | MGP | WOX5 | CLE40 |
|---|---|---|---|---|---|---|---|---|---|---|---|---|---|---|---|---|---|---|
| QC | 0 | 0 | 0 | 0 | 1 | 0 | 1 | 1 | 1 | 1 | 1 | 1 | 1 | 0 | 1 | 0 | 1 | 0 |
| CEI/ Endodermis PD | 0 | 0 | 0 | 0 | 1 | 0 | 0 | 1 | 1 | 1 | 1 | 1 | 1 | 0 | 1 | 1 | 0 | 0 |
| P .Pro-vascular PD | 0 | 0 | 0 | 0 | 1 | 1 | 1 | 1 | 1 | 1 | 0 | 1 | 1 | 0 | 0 | 0 | 0 | 0 |
| C. Pro-vascular PD | 0 | 0 | 0 | 0 | 1 | 1 | 1 | 1 | 1 | 1 | 0 | 1 | 0 | 1 | 0 | 0 | 0 | 0 |
| C. Pro-vascular TD2 | 1 | 1 | 0 | 0 | 1 | 1 | 1 | 1 | 1 | 1 | 0 | 0 | 0 | 1 | 0 | 0 | 0 | 1 |
| Columella 1 | 1 | 1 | 0 | 0 | 1 | 1 | 1 | 1 | 1 | 1 | 0 | 0 | 1 | 0 | 0 | 0 | 0 | 1 |

FIG. S43. 2020 model of the *Arabidopsis thaliana* RSCN Boolean GRN and its fixed points with AUX = 1. This network has 18 nodes and 51 edges. The network is constructed using regulatory interactions obtained from the BFs of the model in [13]. Here, QC: Quiescent center, CEI: Cortex-endodermis initials, P. Pro-vascular PD: Peripheral Pro-vascular initials, C. Pro-vascular PD: Central Pro-vascular initials, C. Pro-vascular TD2: Transition domain, Columella 1: Columella initials.
